# Vagal Signaling Decline in Age-related Macular Degeneration Drives Spleen-Dependent Retinal Inflammation

**DOI:** 10.64898/2026.02.03.703339

**Authors:** Kaitryn E. Ronning, Christophe Roubeix, Caroline Nous, Sébastien Augustin, Frédéric Blond, Foteini Argyriou, Christopher Rue Molbech, Katalin Veres, Jesper Mehlsen, Marie Bak, Emmanuel L. Gautier, Henrik Toft Sørensen, Xavier Guillonneau, Cécile Delarasse, Torben Lykke Sørensen, Florian Sennlaub

## Abstract

Under relaxed physiological conditions, vagal innervation via the splenic nerve restrains the release of inflammatory cytokines from splenic macrophages and helps preserve systemic homeostasis, constituting the efferent cholinergic anti-inflammatory arm of the inflammatory reflex. Our analysis of Danish National Patient Registry data revealed that vagotomy for peptic ulcer disease, particularly truncal vagotomy, markedly increased the risk of developing age-related macular degeneration (AMD). Consistently, experimental truncal vagotomy or splenic denervation in mice exacerbated laser- and light-induced subretinal inflammation, both models of late AMD. The pro-inflammatory effects of vagotomy were abolished by concurrent splenectomy or pharmacologically augmenting acetylcholine signaling. Mechanistically, vagotomy activated a pro-inflammatory transcriptional program in splenic monocytes while suppressing tissue-retention genes *Cxcr4* and *Fn1*, leading to enhanced monocyte egress from the spleen and increased infiltration into the injured retina. In the laser-injured retina, single-cell RNA sequencing (scRNAseq) of mononuclear phagocytes revealed a broad set of vagotomy-induced transcripts in infiltrating monocytes and activated microglia. Remarkably, more than one-third of these were normalized by splenectomy. Together, these findings identify a vagus nerve–spleen–retina axis that connects stress and vagal tone to AMD pathogenesis.

**Graphical abstract:** 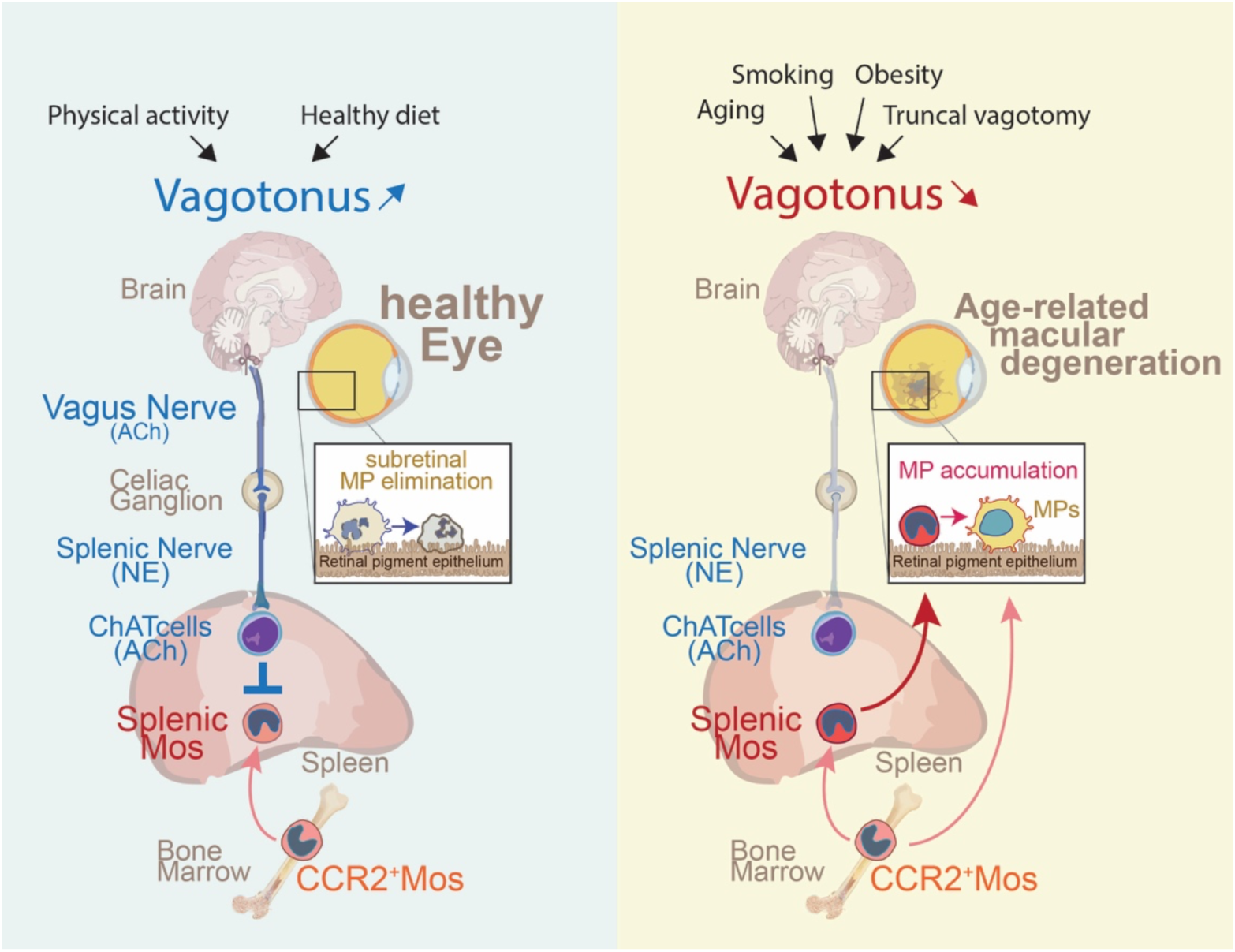

**In Brief:** Using population-level human data and complementary mouse models, this study demonstrates that loss of vagal signaling enhances subretinal inflammation and increases susceptibility to age-related macular degeneration (AMD). Mechanistically, vagotomy reprograms splenic monocytes toward a pro-inflammatory state, promoting their egress and infiltration into the injured retina, where they amplify local inflammation. These effects identify a vagus nerve–spleen–retina axis that links reduced vagal tone to AMD pathogenesis.

**Highlights:**

- Vagotomy increases AMD risk in humans and exacerbates subretinal inflammation in mouse models of late-stage AMD.
- Loss of vagal signaling drives pro-inflammatory reprogramming and enhanced egress of splenic monocytes via suppression of tissue-retention genes.
- Single-cell transcriptomics reveals vagotomy-induced inflammatory signatures in retinal monocytes and microglia that are largely reversed by splenectomy.

## Background

Age-related macular degeneration (AMD) arises from age, genetic, and environmental factors. Early disease is marked by subretinal deposits (pseudodrusen and soft drusen), and progression leads to debilitating choroidal neovascularization (CNV, neoAMD) or an extending area of outer retinal degeneration (geographic atrophy, GA) (Guillonneau et al., 2017). VEGF inhibition suppresses CNV, but complement factor 3 or 5 blockade provides little benefit for neurodegeneration in GA, and all require burdensome lifelong repeated injections.

A shared pathogenic axis across AMD forms is chronic inflammation (Guillonneau et al., 2017; Yu et al., 2020). In AMD, microglia (MC) become activated and, along with monocyte-derived macrophages (MdMs), infiltrate the photoreceptor layer that is normally devoid of immune cells (Guillonneau et al., 2017; Lad et al., 2015; Sennlaub et al., 2013). The three major genetic risk factors, APOE2 (Levy et al., 2015), a CFH variant (Calippe et al., 2017), and 10q26-driven HTRA1 overexpression in MPs (Beguier et al., 2020), promote this MP accumulation (Beguier et al., 2020; Calippe et al., 2017; Levy et al., 2015). In early disease MPs may contribute to debris clearance, but in advanced AMD age-related, genetic, and environmental factors promote their pathogenic polarization. In animal models, infiltrating activated MPs release inflammatory cytokines and reactive oxygen species that drive CNV, accelerate photoreceptor loss, and amplify inflammation by recruiting additional MPs (Guillonneau et al., 2017). MdMs in general and particularly those derived from splenic monocytes, are especially pathogenic in these models (Roubeix et al., 2024; Sennlaub et al., 2013).

Evidence from human studies supports similar mechanisms. Circulating monocytes from patients with AMD display elevated CCL2, IL-8, VEGFA, and IL-6 expression, among other transcriptional alterations (Grunin et al., 2016; Lechner et al., 2017). Serum IL-6 levels further correlate with AMD incidence and progression (Guillonneau et al., 2017; Nahavandipour et al., 2020) and aqueous humor IL-6 levels are increased in GA (Huang et al., 2025). Consistent with a pathogenic role of mononuclear phagocyte cytokines, Janus kinase (JAK) inhibitors, which block IL-6, GM-CSF, IFN-γ, and other pro-inflammatory cytokines signaling, are associated with reduced AMD incidence (Hallak et al., 2024). Collectively, these findings highlight both local and systemic inflammation as central drivers of AMD pathogenesis.

One important regulator of the immune system is the inflammatory reflex, a vagus nerve–mediated neuroimmune pathway that senses peripheral inflammation and suppresses excessive cytokine production, thereby maintaining immune homeostasis. In its cholinergic anti-inflammatory efferent arm, parasympathetic signals from the vagus nerve stimulate the splenic nerve at the coeliac ganglion, increasing norepinephrine release in the spleen. This stimulates acetylcholine release from choline acetyltransferase (ChAT)-expressing T cells, which activates α7 nicotinic acetylcholine receptors on splenic macrophages, delivering a tonic inhibitory signal that suppresses macrophage activity and limits the release of proinflammatory cytokines into the circulation (Andersson & Tracey, 2012; Chavan et al., 2017). Disruption of this reflex by truncal vagotomy in animals leads to disproportionate cytokine responses and increased mortality during sterile endotoxic shock, whereas vagal nerve stimulation improves survival (Borovikova et al., 2000). In humans, truncal vagotomy has been linked to an increased incidence of RA (Baker et al., 2025), whereas implantable vagus nerve stimulation has been shown to reduce disease severity and was recently granted FDA premarket approval (PMA number P240039) (Koopman et al., 2016). It remains unclear, however, whether the protective effect of high vagal tone in RA is mediated through regulation of systemic cytokine release or through other, as yet unidentified, anti-inflammatory mechanisms.

Vagal tone diminishes with aging, smoking and obesity (Andersson & Tracey, 2012; Bodin et al., 2017), the main non-genetic AMD risk factors (Fleckenstein et al., 2021), and increases with aerobic exercise, which protects against AMD (Andersson & Tracey, 2012; McGuinness et al., 2017). We recently demonstrated that reduced heart rate variability, reflecting impaired vagal tone, is associated with neoAMD (Molbech et al., 2025). This observation suggests that individuals with AMD may exhibit diminished vagal activity, which could contribute to a dysregulated inflammatory reflex, heightened inflammatory responses, and progression of the disease.

Here we demonstrate that truncal vagotomy is associated with an increased risk of AMD in humans, and that experimental vagotomy or splenic denervation in mice exacerbated subretinal inflammation. These pro-inflammatory effects depended on an intact cholinergic anti-inflammatory pathway and were abolished by splenectomy or by inhibition of acetylcholine degradation. Mechanistically, vagotomy reprogrammed splenic monocytes and promoted their mobilization, leading to exacerbated retinal inflammation, Together, these findings define a vagus–spleen–retina axis that links vagal tone to splenic monocyte mobilization and pathogenic macrophage infiltration in AMD pathogenesis, and highlight potential therapeutic targets.

## Results

### Vagotomy is associated with increased incidence of AMD

To investigate whether loss of vagal tone influences the risk of age-related macular degeneration (AMD), we performed a nationwide cohort study using Danish National Patient Registry data (Fig. 1A). Although advances in medical therapy, including antibiotics and proton pump inhibitors, have largely rendered vagotomy obsolete, the procedure was commonly performed for peptic ulcer disease in Denmark between 1977 and 1995 (Lagoo et al., 2014; Svensson et al., 2015). Two vagotomy procedures were analyzed: truncal vagotomy (TV), which involves cutting the vagus nerves inferior to the esophageal hiatus and denervates major abdominal organs including the spleen, and superselective vagotomy (SSV), which cuts only gastric branches while largely sparing innervation of other abdominal organs, including the spleen (Fig. 1A). These procedures provided two patient groups with similar indications for surgery but differing degrees of systemic vagal denervation.

**Figure 1.**
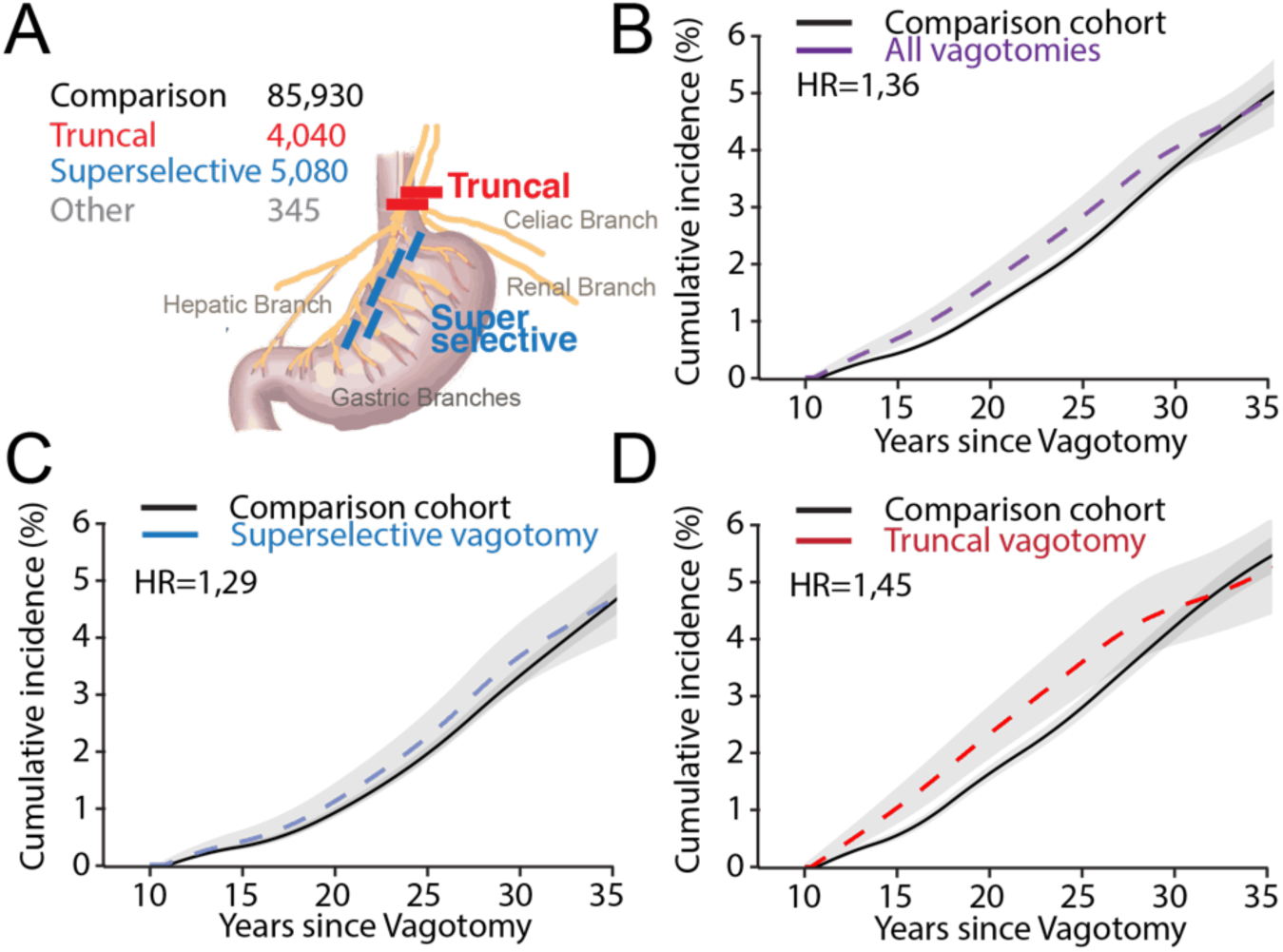
Vagotomy is associated with increased incidence of AMD. (A) Study cohorts included all patients in Denmark who underwent vagotomy between 1977 and 1995 with ≥10 years follow-up (n=9,465), subdivided into truncal vagotomy (TV, n=4,040), superselective vagotomy (SSV, n=5,080), and unspecified type (n=345). Each patient was matched to 10 individuals from the general population (n=85,930) by exact birth year and sex using the Civil Registration System. The schematic illustrates that TV denervates abdominal organs including the celiac branch that innervates the spleen, whereas SSV primarily targets gastric branches and spares extra-gastric innervation. (B) Cumulative incidence of AMD in all vagotomized patients compared to their matched population cohort. (C) Cumulative incidence of AMD in patients with superselective vagotomy compared to their matched cohort. (D) Cumulative incidence of AMD in patients with truncal vagotomy compared to their matched cohort. Follow-up began 10 years after index date as the Civil Registry was only available from 1986; curves were estimated using Kaplan–Meier analysis treating death as a competing risk and smoothed for visualization using splines. Hazard ratios (HR) were obtained from Cox regression adjusted for age, sex, COPD, obesity-related diseases, hypertension, and Charlson Comorbidity Index categories (full details in Table 1 and Supplemental Tables 1–3).

**Table 1.** Incidence rates and hazard ratios of age-related macular degeneration for vagotomy patients surviving 10 years after vagotomy and their comparison cohort. AMD = age-related macular degeneration; COPD = chronic obstructive pulmonary disease; CCI = Charlson comorbidity index.

|  | No. of AMD | Incidence Rate (per 1000 person-years at risk) (95% CI) | Hazard ratio controlled for age and gender by design | Adjusted for age, gender, and COPD | Adjusted for age, gender, and obesity-related diseases | Adjusted for age, gender, and hypertension | Adjusted for age, gender, and CCI | Adjusted for age, gender, COPD, obesity-related diseases, hypertension and CCI |
| --- | --- | --- | --- | --- | --- | --- | --- | --- |
| Comparison cohort | 3070 | 2.40<br>(2.31-2.48) |  |  |  |  |  |  |
| All vagotomy | 360 | 3.00<br>(2.69-3.31) | 1.37<br>(1.22-1.55) | 1.36<br>(1.21-1.53) | 1.37<br>(1.22-1.54) | 1.37<br>(1.22-1.55) | 1.37<br>(1.21-1.54) | 1.36<br>(1.21-1.53) |
| Comparison cohort | 1400 | 2.85<br>(2.70-3.00) |  |  |  |  |  |  |
| Truncal vagotomy | 170 | 3.85<br>(3.29-4.44) | 1.47<br>(1.23-1.74) | 1.46<br>(1.23-1.73) | 1.46<br>(1.23-1.74) | 1.47<br>(1.24-1.74) | 1.46<br>(1.23-1.73) | 1.45<br>(1.22-1.73) |
| Comparison cohort | 1550 | 2.09<br>(1.99-2.19) |  |  |  |  |  |  |
| Superselective vagotomy | 180 | 2.47<br>(2.12-2.85) | 1.31<br>(1.11-1.55) | 1.30<br>(1.10-1.53) | 1.31<br>(1.11-1.55) | 1.31<br>(1.11-1.55) | 1.30<br>(1.10-1.54) | 1.29<br>(1.09-1.53) |

The study cohort included 9,465 patients who underwent vagotomy with at least 10 years of follow-up: 4,040 underwent TV, 5,080 underwent SSV, and 345 had unspecified vagotomy (Supplemental Table 1). A comparison cohort of 85,930 age- and sex-matched individuals from the general population was randomly sampled at a 10:1 ratio for each vagotomized patient. The index date for the comparison cohort was defined as the date of vagotomy in the matched patient. Patients with AMD diagnosed prior to the index date were excluded from all analyses.

During follow-up, vagotomy was associated with a higher cumulative incidence of AMD. After adjustment for age, sex, chronic obstructive pulmonary disease (COPD, as a proxy for smoking), hypertension, obesity-related diseases, and 19 Charlson Comorbidity Index (CCI) categories, the overall adjusted hazard ratio (aHR) for AMD was 1.36 (Table 1, Figure 1B, Supplemental Table 2). When examining TV and SSV separately, SSV was associated with only a modest increase in AMD incidence (2.47 per 1,000 person-years vs. 2.09 in its comparison cohort), whereas TV was associated with a substantially higher incidence (3.85 per 1,000 person-years vs. 2.85 in its comparison cohort) and an aHR of 1.45 (Table 1, Figure 1C-D, Supplemental Table 2).

The lack of a pronounced association in the SSV cohort suggests that the increased risk of AMD observed in TV patients is unlikely to be driven solely by peptic ulcer disease or by vagotomy-induced changes in gastric acidity, or other important peptic ulcer risk factors such as smoking, which are common to both procedures. Instead, the data indicate that broader abdominal denervation, including the splenic branch of the vagus nerve, may underlie the observed increase in AMD incidence.

### Vagotomy promotes inflammation and exacerbates disease severity in mouse models of AMD in a spleen-dependent manner

Our data identified truncal vagotomy as an AMD risk factor, implicating reduced vagal tone in disease pathogenesis, in agreement with evidence that the AMD-risk factors age, smoking, and obesity depress vagotonus, whereas aerobic exercise protects against AMD and increases vagotonus (Andersson & Tracey, 2012; Bodin et al., 2017; Fleckenstein et al., 2021; McGuinness et al., 2017). Physiologically, parasympathetic vagal output activates the adrenergic splenic nerve at the celiac ganglion, triggering acetylcholine release from ChAT⁺ T cells. This provides a tonic inhibitory signal that suppresses proinflammatory cytokine production by splenic macrophages and may exert additional spleen-dependent anti-inflammatory effects (Andersson & Tracey, 2012; Chavan et al., 2017) (Fig. 2A). To model reduced vagal tone, we used left cervical truncal vagotomy, which strongly reduces splenic nerve activity in mice (Carnevale et al., 2016).

**Figure 2.**
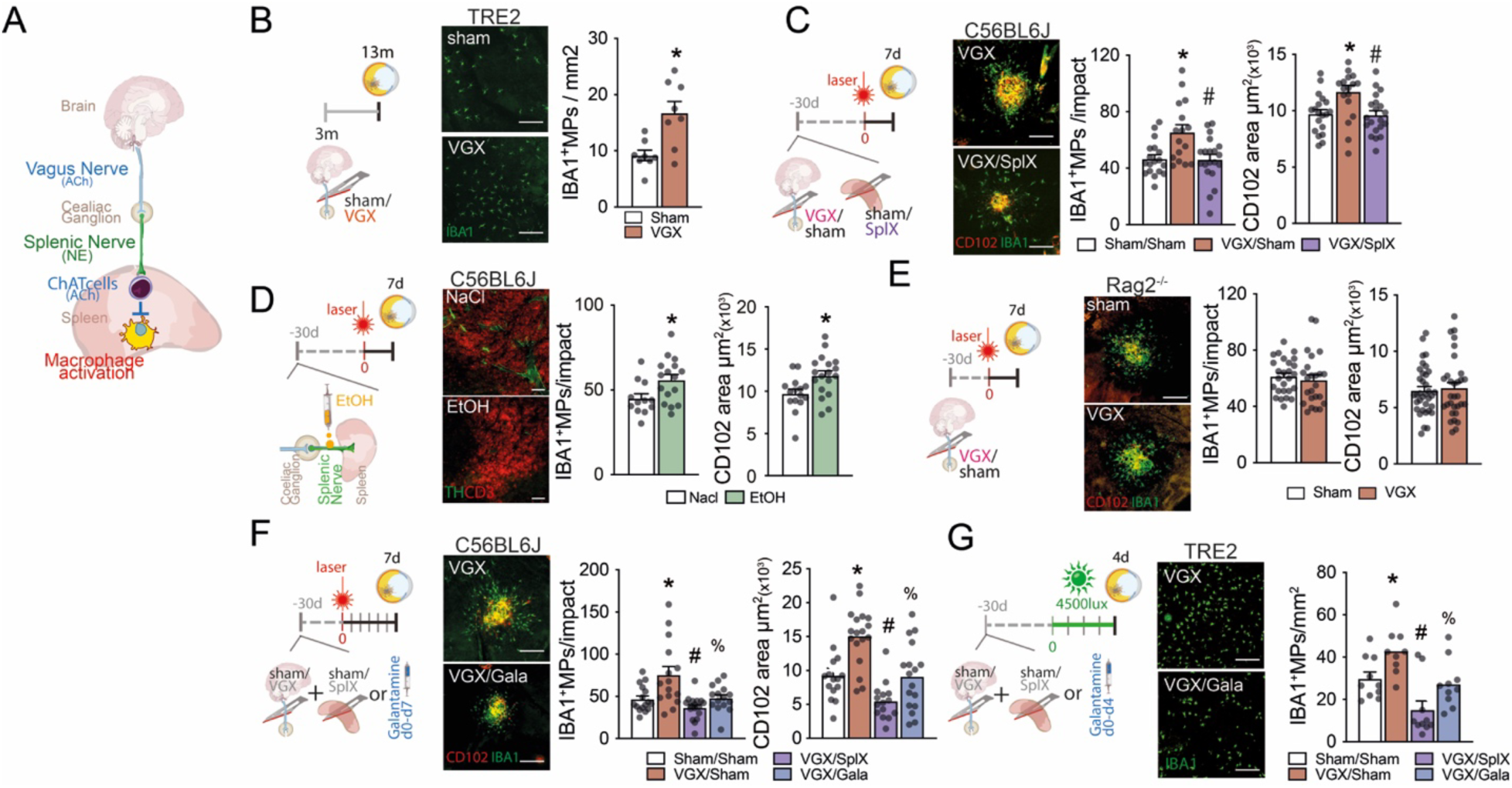
Vagotomy promotes inflammation and exacerbates disease severity in mouse models of AMD in a spleen-dependent manner. (A) Schematic of the parasympathetic vagus–spleen axis: Acetylcholine from vagal efferents activate the splenic nerve at the coeliac ganglion, increasing splenic norepinephrine and driving ChAT⁺ T cell–derived acetylcholine that tonically restrains mononuclear phagocytes (MPs). (B) Representative images and quantification of subretinal IBA1⁺ MPs on IBA1-stained RPE/choroidal flatmounts from TRE2 mice subjected to left cervical truncal vagotomy at 3 months and analyzed at 13 months (n=8 per group; Mann-Whitney test *p=0.0148). (C) Laser-induced choroidal neovascularization (CNV) model in 3-month-old male C57BL/6J mice: vagotomy with or without splenectomy performed 1 month before laser (at 2 months old); quantification of subretinal IBA1⁺ MP counts and CD102⁺ CNV area on IBA1/CD102 double-stained RPE/choroidal flatmounts 7 days after laser injury (n= 16-20 per group; one-way ANOVA/Tukey: IBA1 Sham/Sham versus VGX/Sham *p=0.0057, VGX/Sham versus VGX/SplX #p=0.0034, Sham/Sham versus VGX/SplX not significant (ns); CD102 Sham/Sham versus VGX/Sham *p=0.0123, VGX/Sham versus VGX/SplX #p=0.0068, Sham/Sham versus VGX/SplX ns). (D) Splenic denervation in 2-month-old male C57BL/6J mice by ethanol application at the splenic hiatus: (left) verification by loss of tyrosine-hydroxylase (TH, green)(co-stained for CD3, magenta, and DAPI, blue) immunostaining on spleen sections; (right) quantification of subretinal IBA1⁺ MP counts and CD102⁺ CNV area on IBA1/CD102 double-stained RPE/choroidal flatmounts 7 days after laser injury (laser 1 month post-surgery)(n= 12-18 per group; Mann-Whitney: IBA1 *p=0.0284; CD102 *p=0.0118). (E) Laser model in 2-month-old male Rag2^-/-^ mice (lacking mature lymphocytes including ChAT⁺ T cells): quantification of subretinal IBA1⁺ MPs and CD102⁺ CNV area on IBA1/CD102 double-stained RPE/choroidal flatmounts 7 days after laser injury (n= 24-26 per group; Mann-Whitney: IBA1 and CD102 not statistically significant). (F) Daily intraperitoneal galantamine (an acetylcholinesterase inhibitor, 4 mg/kg) initiated after laser in previously vagotomized 3-month-old male C57BL/6J mice: quantification of IBA1⁺ MP counts and CD102⁺ CNV area on IBA1/CD102 double-stained RPE/choroidal flatmounts 7 days after laser injury (n= 15-17 per group; one-way ANOVA/Tukey: IBA1 Sham/Sham versus VGX/Sham *p=0.005, VGX/Sham versus VGX/SplX #p<0.001, VGX/Sham versus VGX/Gala%p=0.0093; CD102 Sham/Sham versus VGX/Sham *p=0.0009, VGX/Sham versus VGX/SplX #p<0.0001, VGX/Sham versus VGX/Gala %p=0.0004). (G) Bright-light challenge calibrated to induce subretinal MP infiltration in TRE2 but not controls; quantification of IBA1⁺ MPs on IBA1-stained RPE flatmounts from 3-month-old male TRE2 mice after 4 days of light exposure, one month after sham or vagotomy with or without splenectomy or daily galantamine (4 mg/kg)(n=9-10 per group; one-way ANOVA/Tukey: Sham/Sham versus VGX/Sham *p=0.0344, VGX/Sham versus VGX/SplX #p<0.0001, VGX/Sham versus VGX/Gala %p=0.0285). Statistics: data are represented as mean ± SEM, Mann-Whitney test (two-group comparisons) or one-way ANOVA with Tukey’s test (multiple comparisons) as indicated; replicates represent quantifications of eyes or plasma samples from different mice (C57BL/6Js were all males, TRE2 were males and females) from at least three independent cages/experiments. ChAT: choline acetyltransferase; TRE2: targeted-replacement mice expressing human APOE2; MP: mononuclear phagocyte; CNV: choroidal neovascularization. Scale bars: 20µm

In humanized transgenic mice expressing the AMD-risk APOE2 isoform (TRE2 mice), vagotomy at 3 months of age increased age-associated subretinal IBA1⁺ MP accumulation that characterizes this model (Calippe et al., 2017; Levy et al., 2015) measured at 13 months on RPE/choroidal flatmounts (Fig. 2B). In the laser-induced subretinal inflammation and choroidal neovascularization model (the standard model of neoAMD) in 3–month-old male C57BL/6J mice, vagotomy (performed one month before laser to allow for complete recovery) increased IBA1⁺ MP accumulation and CD102⁺ CNV area quantified on RPE/choroidal flatmounts. Concomitant splenectomy normalized both parameters to levels observed in sham-operated controls, demonstrating a spleen-dependent effect (Fig. 2C).

We next performed a series of experiments to determine if the pro-inflammatory effect of vagotomy was mediated via the inflammatory reflex. First, denervating the spleen by ethanol application at the splenic hiatus in 2-month-old male C57BL/6J mice (one month before laser to allow for recovery), confirmed by loss of tyrosine-hydroxylase (TH) staining on spleen sections (Fig. 2D, left), increased laser-induced IBA1⁺ MP accumulation and CD102⁺ CNV on RPE/choroidal flatmounts relative to sham, phenocopying vagotomy and confirming splenic nerve involvement (Fig. 2E, right). Second, vagotomy failed to augment subretinal MPs or CNV after laser in 3-month-old male *Rag2^-/-^*mice, which lack mature lymphocytes including ChAT⁺ T cells but continue to harbor pathogenic splenic monocytes (Roubeix et al., 2024) (Fig. 2E).

Next, mice received daily intraperitoneal injections of the acetylcholinesterase inhibitor galantamine after laser to restore acetylcholine availability diminished by vagotomy and reduced ChAT-cell output. This reversed the pro-inflammatory phenotype, returning subretinal MP counts and CNV size to sham in the laser CNV model, similar to splenectomy (Fig. 2F). Further, 3-month-old male TRE2 mice were exposed to a calibrated bright-light challenge, which induces MP infiltration in TRE2 mice but not in controls and within a shorter time frame than in the experiments shown in Fig. 2B (Calippe et al., 2017; Levy et al., 2015). Quantification of IBA1+ MPs on RPE flatmounts revealed that vagotomy also increased MP accumulation in this model, an effect again abrogated by splenectomy and reversed by galantamine (Fig. 2G).

Collectively, our results demonstrate that disrupting vagus-to-spleen cholinergic signaling through truncal vagotomy amplifies subretinal MP responses and CNV. This effect requires both splenic innervation and lymphocytes, most likely ChAT^+^ T cells, and it can be reversed by enhancing acetylcholine tone, consistent with the cholinergic efferent arm of the inflammatory reflex. Notably, these findings in animal models parallel the finding in the cohort study that truncal vagotomy promoted AMD pathogenesis (Fig.1).

### Loss of vagal tone reshapes the splenic monocyte transcriptome and drives their mobilization

The spleen functions as a major reservoir of CCR2⁺Ly6C^high^ monocytes (SpleMos) (Swirski et al., 2009), which, together with circulating Ly6C^high^ monocytes, are recruited into diseased tissues (Robbins et al., 2012; Swirski et al., 2009) where they differentiate into inflammatory MdMs. We recently demonstrated that SpleMos play a particularly pathogenic role in retinal inflammation in AMD (Roubeix et al., 2024), analogous to their established role in atherosclerosis (Robbins et al., 2012). On this basis, we next examined cellular and molecular changes in the spleen, focusing specifically on potentially pathogenic splenic monocytes following the loss of vagal tone.

Flow cytometric analysis of spleens from 3-month-old mice, performed 30 days after vagotomy, revealed no differences in the numbers of CD45⁺CD11b⁺ MPs, CD45⁺CD11b⁺Ly6C^high^ monocytes (Mos), or CD4⁺ and CD8⁺ T cells compared with sham-operated controls (Fig. 3A, Supplemental Fig. 1). Bulk RNA sequencing of FACS-sorted CD45⁺CD11b⁺Ly6C^high^Ly6G^neg^ splenic Mos identified 508 upregulated and 1860 downregulated genes in splenic Mos from vagotomized mice compared with sham-operated controls (Fig. 3B, Supplemental Table 5). Classical monocyte markers, including *Ccr2*, *Ly6c2*, *Cd14*, and *Sell*, were expressed at comparable levels between groups (Fig. 3C). In contrast, several genes associated with immune defense were upregulated in Mos from vagotomized mice, such as MHC class II-related genes (*H2-Aa*, *CD74*) and pattern recognition receptors (*Clec4g*, *CD209e*) (Fig. 3D). Several pro-inflammatory genes were also upregulated, including *Il1b*, *Lilra5*, *Ccl4*, and *C5ar1* (Fig. 3E). Conversely, among the 1860 downregulated genes were several implicated in monocyte retention. These included two of the most strongly regulated genes, *Fn1*, whose autocrine expression anchors endothelial cells within the extracellular matrix (Cseh et al., 2010), and *Cxcr4*, a central mediator of bone marrow monocyte retention (Chong et al., 2016), along with the integrins *Itga4* and *Itga1* (Fig. 3F).

**Figure 3.**
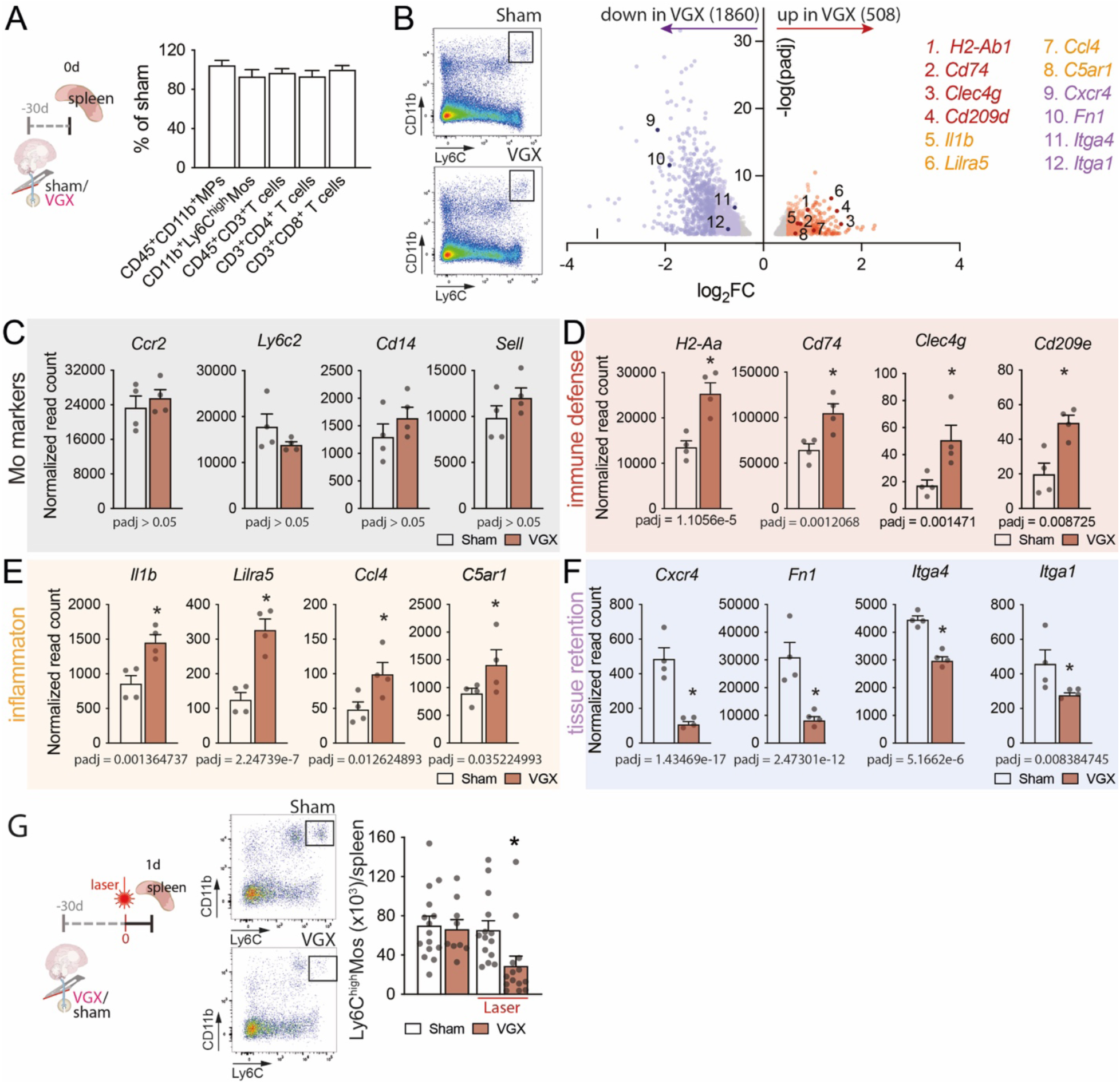
Loss of vagal tone reshapes the splenic monocyte transcriptome and drives their mobilization. (A) Flow cytometric quantification of splenic CD45⁺CD11b⁺ MPs, CD45⁺CD11b⁺Ly6C^high^ monocytes (Mos), CD4⁺ T cells, and CD8⁺ T cells from 3-month-old male mice 30 days after vagotomy (VGX) or sham surgery, expressed as percentage of sham (see Supplemental Fig. 1 for comparisons between sham and VGX). B) Bulk RNA sequencing of FACS-sorted CD45⁺CD11b⁺Ly6G^neg^Ly6C^high^ splenic Mos: (left) representative flow plots; (right) volcano plot showing downregulated (n=1860) and upregulated (n=508) genes in VGX versus sham (right), with selected genes highlighted. (C–F) Normalized read counts of selected genes: (C) classical monocyte markers (*Ccr2*, *Ly6c2*, *Cd14*, *Sell*); (D) immune defense–associated genes (*H2-Aa*, *Cd74*, *Clec4g*, *Cd209e*); (E) inflammatory genes (*Il1b*, *Lilra5*, *Ccl4*, *C5ar1*); (F) tissue retention–associated genes (*Cxcr4*, *Fn1*, *Itga4*, *Itga1*). Adjusted p values are indicated in the figure. (G) Experimental schematic of vagotomy/sham followed by laser injury or no injury 30 days later, with analysis at day 1 (left) and representative flow cytometric plots and quantification of splenic Ly6C^high^ Mos from sham and VGX mice with or without laser (right, n=9-15 per group; Mann-Whitney: Laser Sham versus VGX *p=0.001; no laser Sham versus VGX not statistically significant). Statistics: (G) data are represented as mean ± SEM, Mann-Whitney test (two-group comparisons); (B-F) see methods for RNAseq analysis details; replicates represent spleens from different mice (all males). MP: mononuclear phagocyte; Mo: monocyte; VGX: vagotomy.

To assess how vagal denervation affects monocyte efflux from the spleen, we analyzed spleens of vagotomized mice with or without laser injury at day 1 (Fig. 3G). In the spleen, prior to laser injury, the number of Ly6C^high^ monocytes did not differ between groups, as quantified by flow cytometry. However, one day after laser treatment, spleens from vagotomized mice showed a marked reduction in monocyte numbers, indicating enhanced efflux in response to injury (Fig. 3G).

Together, these findings suggest that disruption of vagal signaling alters the splenic monocyte transcriptome toward enhanced immune defense and a pro-inflammatory profile, potentially influencing their polarization upon infiltration into diseased tissues. Importantly, vagotomy increased the mobilization of splenic Mos before the vagotomy-induced, spleen-dependent increase in subretinal inflammation that we observed (Fig. 2).

### Vagotomy induces transcriptional reprogramming of infiltrating monocytes and activated microglia in the retina in a spleen-dependent manner

Since vagotomy altered the splenic monocyte transcriptome, enhanced their mobilization (Fig. 3), and promoted spleen-dependent accumulation of chorio-retinal MPs (Fig. 2), we next investigated how vagotomy-primed splenic monocytes influence the composition and polarization of resident and infiltrating MPs during sterile chorio-retinal inflammation. We performed single-cell RNA sequencing on sorted CD45⁺CD11b⁺ cells from posterior eye cups from vagotomized or sham operated mice one day after laser injury or without laser-injury (Fig. 4A). Using Seurat for dimensionality reduction with principal component analysis (PCA), UMAP, and graph-based clustering, we identified 11 transcriptionally distinct clusters of MPs based on canonical marker gene expression (Fig. 4B). Under homeostatic non-laser conditions (Fig. 4C, upper panel), these clusters were annotated as resident macrophages (*C1qa*^high^, *Cd68*^high^) of the choroid (*F13a1*^high^) or as microglial cells (MCs, *P2ry12*^high^), as well as circulating monocytes (*Nr4a1*⁺, *Spn*⁺, *Ace*⁺) that segregated into two subsets: patrolling Mos (*Ly6c2*^low^, *Ccr2*^low^) and inflammatory Mos (*Ly6c2*^high^, *Ccr2*^high^). In post-lasered eyes, we additionally observed several subsets of activated MCs, characterized by reduced expression of homeostatic MC genes and upregulation of inflammatory cytokines (*Ccl12*, *Ccl2*) (clusters 2-5, Fig. 4C lower panel). We also detected a large population of infiltrating Mos (cluster 9, *Lgals3*^high^, *Il1b*^high^, *Spn*^low^, *Ace*^low^), as well as early monocyte-derived macrophages distinguished by increased *Itgax* (Cd11c, cluster 10 and 11) expression. It is important to note that the Seurat-defined clusters are based on a broad transcriptional profile, and these genes are shown here only as representative examples.

**Figure 4.**
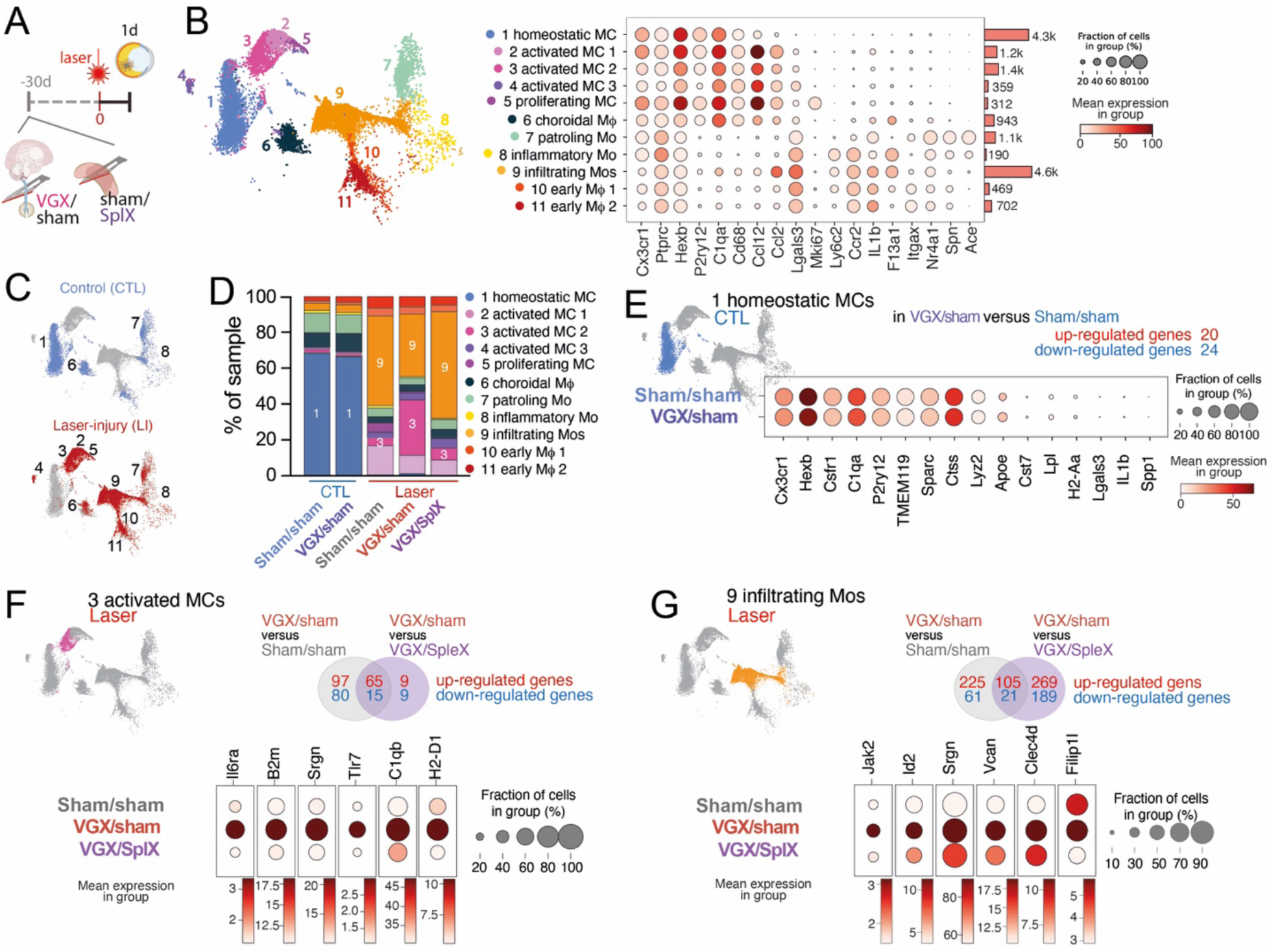
Vagotomy induces transcriptional reprogramming of infiltrating monocytes and activated microglia in the retina in a spleen-dependent manner. (A) CD45⁺CD11b⁺ cells were FACS-sorted from posterior eye segments of mice 1 month after sham or vagotomy (VGX), and from mice 1 day after laser injury that had undergone sham, VGX, or VGX with splenectomy (VGX+SPLX) one month prior. (B) UMAP projection of myeloid cell clusters (left) and dot plot showing expression of representative marker genes (right). (C) Distribution of cells between control (non-lasered, CTL) and lasered samples. (D) Relative proportions of each cluster across experimental conditions. (E) Expression of representative homeostatic and activated microglia genes in the homeostatic microglia cluster from non-lasered sham and VGX samples. (F) Differential gene expression analysis of cluster 3 (activated MC2) between surgery conditions (top), with representative genes shown as dot plots (bottom). (G) Differential gene expression analysis of cluster 9 (infiltrating monocytes, Mos) between surgery conditions (top), with representative genes shown as dot plots (bottom). VGX: vagotomy; SplX: splenectomy; CTL: control; LI: laser injury.

In non-lasered eyes, vagotomy did not alter the overall cell type composition compared to sham-operated controls. The majority of MPs were homeostatic microglia, accompanied by a small cluster of choroidal macrophages and predominantly patrolling monocytes (Fig. 4D). The activation state of homeostatic microglia also remained unchanged, as evidenced by stable expression of homeostatic markers, absence of activation marker expression, and only limited transcriptional changes, with 20 genes upregulated and 24 genes downregulated between vagotomy and sham conditions (Fig. 4E, Supplemental Table 6).

In sharp contrast, the lasered samples were dominated by activated microglia (clusters 2-5) and infiltrating monocytes (clusters 9 and 10; Fig. 4C–D). Within the activated microglial population, several subclusters were detected, but cluster #3 (activated MC 2) was markedly expanded in the vagotomized condition compared to both sham and splenectomy controls, prompting us to focus on this subset (Fig. 4F and G). Relative to sham/sham-operated laser-injured animals, vagotomized/sham laser-injured mice displayed 162 upregulated and 95 downregulated genes in the MC cluster 3 (Fig. 4F). Vagotomized/sham laser-injured mice displayed 74 upregulated and 24 downregulated genes relative to VGX/SpleX -operated laser-injured animals (Supplemental Table 7). Strikingly, the vagotomy-induced deregulation of a large proportion of genes (65/162 and 15/95, respectively) was reversed by splenectomy (Fig. 4F, Supplemental Table 8). The most regulated genes included *Il6ra*, *B2m*, *Srgn*, *Tlr7*, *C1qb*, and *H2-D1*, covering pathways in cytokine signaling, antigen presentation, proteoglycan-mediated granule biology, innate immune sensing, complement activation, transcriptional regulation, and MHC class II presentation. Similarly, in infiltrating monocytes, we identified 330 upregulated and 82 downregulated genes in vagotomized/sham laser-injured mice, of which 105 and 21, respectively, were normalized in VGX/SpleX laser-injured animals (Fig. 4G, Supplemental Tables 9-10). The most regulated genes included *Jak2*, *Id2*, *Srgn*, *Vcan*, *Clec4d*, and *Filip1l*, which are associated with cytokine signaling, transcriptional control, granule biology, extracellular matrix organization, pathogen recognition, cytoskeletal regulation, and immune modulation.

Taken together, vagotomy had minimal impact on the transcriptome of MPs in uninjured eyes. By contrast, following laser injury, which recruits, in part, spleen-derived monocytes (Roubeix et al., 2024), vagotomy induced transcriptional changes in infiltrating monocytes but also activated microglia. Notably, splenectomy reversed a substantial fraction of these transcriptional changes.

## Discussion

Factors known to reduce vagal tone, such as aging, smoking, and obesity (Andersson & Tracey, 2012; Bodin et al., 2017), are also established risk factors for AMD (Fleckenstein et al., 2021). In line with this, we recently reported that reduced heart rate variability, a marker of vagal tone, is associated with AMD (Molbech et al., 2025). In the present study, we demonstrated that the surgical vagotomy, once used in the treatment of peptic ulcers, is associated with an increased risk of AMD (all forms combined), independently of age, smoking, and obesity (Fig.1). Superselective vagotomy was associated with only a modest risk increase compared to the substantial risk increase of truncal vagotomy, supporting the interpretation that the observed association with truncal vagotomy is not attributable to peptic ulcer disease or related risk factors such as smoking, which are similarly distributed between the two patient groups.

Since truncal, but not superselective, vagotomy profoundly reduces vagal tone in the spleen, and truncal vagotomy (but not superselective) has recently been linked to the incidence of autoimmune rheumatoid arthritis (Baker et al., 2025), we hypothesized that the observed association of truncal vagotomy with AMD risk is driven by disruption of the anti-inflammatory effect of the vagus nerve on the spleen. Indeed, experimental cervical truncal vagotomy in mice, which markedly reduces splenic nerve activity (Carnevale et al., 2016), led to increased subretinal inflammation following laser injury, as well as exaggerated age-related MP accumulation observed in TRE2 mice expressing the AMD risk gene *APOE2* (Levy et al., 2015) (Fig. 2B-C). Importantly, splenectomy reversed the vagotomy-induced exaggerated inflammation in laser-induced subretinal inflammation and in light induced inflammation in *APOE2*-expressing mice (Fig. 2F-G), highlighting the spleen’s pivotal role in mediating the proinflammatory effects of vagotomy.

We performed a series of experiments to examine if the pro-inflammatory effect of vagotomy is propagated via the disturbance of the inflammatory reflex (Fig. 2A). Splenic denervation alone was sufficient to amplify the laser-induced inflammatory response (Fig. 2D), suggesting that in our model parasympathetic anti-inflammatory influence outweighs the sympathetic tone transmitted through the splenic nerve, consistent with observations of splenic denervation in the endotoxin shock model (Rosas-Ballina et al., 2008). Acetylcholinesterase inhibitor galantamine administration, to restore acetylcholine levels decreased by both diminished vagotonus in the celiac ganglion and from ChAT^+^ T cells in the spleen, also reversed the effect of vagotomy in both models (Fig 2F-G). Interestingly, use of anticholinergic drugs has previously been associated with AMD (Aldebert et al., 2018), which our findings suggests might be mediated at least in part by a loss of cholinergic signaling in the anti-inflammatory reflex.

The finding that vagotomy failed to augment laser-induced subretinal MPs or CNV in *Rag2^-/-^* mice, which lack mature lymphocytes (Shinkai et al., 1992) including ChAT⁺ T cells but still maintain splenic monocytes (Roubeix et al., 2024), further supports ChAT⁺ T-cell dependence (Fig. 2E). However, in contrast to severe sterile endotoxic shock (Borovikova et al., 2000), we did not observe vagotomy-induced excessive cytokine release by splenic macrophages (data not shown) likely because localized ocular laser injury was not sufficient to increase cytokine release from splenic macrophages contrary to endotoxic shock. Taken together this set of experiments demonstrates that experimental vagotomy in mice aggravates AMD models and confirms the clinical observation of truncal vagotomy association with AMD. Further, it establishes a pivotal role of the spleen in mediating this effect.

In the spleen, under homeostatic conditions and in the absence of laser injury, vagotomy did not alter the numbers of MPs, T cells, or monocytes (Fig. 3A, Supplemental Fig. 1). Given that CD45^+^CD11b⁺Ly6C^high^ splenic monocytes play a particularly pathogenic role in retinal inflammation in AMD (Roubeix et al., 2024), we next analyzed their transcriptome. Several pro-inflammatory genes (*Il1b*, *Lilra5*, *Ccl4*, and *C5ar1*), as well as genes linked to immune defense (MHC class II genes and pattern recognition receptors), were upregulated in splenic monocytes after vagotomy compared to sham-operated animals (Fig. 3B-E). These findings raise the possibility that vagal innervation provides a tonic inhibitory input to splenic monocytes, potentially mediated by currently unidentified signals from splenic macrophages, beyond its previously established effects on macrophage cytokine release (Andersson & Tracey, 2012; Chavan et al., 2017). Notably, the most strongly regulated genes included downregulation of *Cxcr4*, a central mediator of bone marrow monocyte retention (Chong et al., 2016) and Fibronectin 1 (*Fn1*), whose autocrine expression anchors endothelial cells within the extracellular matrix (Cseh et al., 2010), along with fibronectin-binding *Itga4* and collagen-binding *Itga1* (Fig. 3F). Given that the spleen serves as a major reservoir for Ly6C^high^ monocytes (SpleMos) (Swirski et al., 2009), we next tested whether these transcriptional changes translated into altered monocyte egress during inflammation. Flow cytometric analysis revealed a marked reduction in splenic Ly6C^high^ monocytes in vagotomized mice compared with sham-operated controls one day after laser injury, consistent with enhanced egress (Fig. 3G).

Because vagotomy reprograms splenic monocytes, increases their mobilization (Fig. 3), and amplifies spleen-dependent subretinal MP accumulation (Fig. 2), we next examined how these primed cells influence MP composition and polarization during sterile chorio-retinal inflammation. In eyes without laser injury, vagotomy produced only minimal transcriptional alterations in the homeostatic microglia and choroidal macrophages (Fig. 4B-E). In contrast, laser-induced injury led to the infiltration of monocytes and the appearance of several activated microglial subtypes (Fig. 4B-D). Notably, vagotomy selectively increased the relative abundance of activated MC2 (cluster #3) in a spleen-dependent manner, demonstrating that signals derived from the spleen are required for this effect. A plausible possibility is that infiltrating spleMos convey these signals; however, directly and specifically labeling spleMos remains technically challenging. Subcapsular red pulp dye injections, similarly to intravenous injections, also label classical circulating blood monocytes. Likewise, spleMos from transplanted GFP⁺ spleens are replaced by GFP⁻ recipient spleMos before the host has fully recovered from surgery. Finally, photoconversion of subcapsular spleMos in *Cx3cr1^Dendra2^* mice (Miller et al., 2021) proved either inefficient or excessively damaging at the ultraviolet intensities required in our hands. Technical advancements and further studies will be necessary to decipher the specific molecular and signaling pathways by which spleMos or other signals from the spleen modulate microglial activation after vagotomy.

In both infiltrating monocytes and activated MCs, the differentially expressed genes regulated by vagotomy were predominantly upregulated. Strikingly one-third of these changes reverted to baseline after splenectomy, underscoring the necessity of the efferent arm of the inflammatory reflex and the spleen.

Together, our demonstration of the involvement of the inflammatory reflex in AMD pathogenesis points to several novel therapeutic strategies. In addition to pharmacological inhibition of acetylcholinesterase the avoidance of anticholinergic drugs, our findings suggest that AMD patients may benefit from therapeutic vagus nerve stimulation, which has been shown to reduce neuroinflammation and modulate microglial polarization in rodent models of spinal cord injury (Chen et al., 2022) and stroke (Zhang et al., 2021).

## Conclusions

In conclusion, we demonstrate that vagotomy is associated with an increased incidence of AMD in a human patient cohort and exacerbates pathogenic inflammation in mouse models of the disease. Mechanistically, this effect stems from an exaggerated, spleen-dependent inflammatory response that profoundly alters the chorioretinal mononuclear phagocyte response during retinal inflammation. The pro-inflammatory consequences of experimental vagotomy can be mitigated by splenectomy and pharmacological enhancement of parasympathetic signaling. Together, these findings provide mechanistic insight into how systemic factors such as aging and obesity may increase the risk of AMD, and they confirm cholinergic drugs and vagus nerve stimulation as potential therapeutic strategies to dampen excessive inflammation in AMD and other conditions associated with dysregulation of the inflammatory reflex.

## Methods

### Danish National Patient Registry cohort study

We conducted a population-based nationwide cohort study using Danish administrative registry data, following principles similar to those in previous epidemiological studies (Svensson et al., 2015). Denmark’s tax-funded healthcare system provides all residents with free, unlimited access to diagnostic work-up and treatment, including for AMD. The Civil Registration System (CRS) assigns all residents a unique identification number, enabling linkage across national registries (Schmidt et al., 2014; Schmidt et al., 2019).

All patients undergoing vagotomy (TV or SSV) between January 1, 1977 and December 31, 1995 were identified via procedural codes in the Danish National Patient Registry (DNPR; Supplemental Table 4). AMD cases were identified with ICD-8 and ICD-10 codes (Supplemental Table 3). Although surgical procedures are generally coded with high accuracy, no validation studies have specifically assessed the accuracy of DNPR procedure codes for vagotomy, so although unlikely misclassification is therefore possible. However, the most likely misclassification, if any, would be the coding of a vagotomy despite incomplete resection. Thus, misclassification, if any, is expected to bias estimated associations towards the null (Svensson et al., 2015).

For each vagotomized patient, up to ten age- and sex-matched individuals from the general population who had not undergone vagotomy were randomly sampled. The index date for each control was defined as the date of vagotomy in the matched patient. Persons in the comparison cohort who underwent vagotomy after the index date were reclassified into the vagotomy cohort. Individuals with pre-existing AMD were not included in the cohorts.

Comorbidities were assessed via linkage to Danish National Patient Registry data according to 19 major disease categories from the Charlson Comorbidity Index (CCI; Supplemental Table 3). Peptic ulcer disease was excluded from the CCI because it was present in nearly all vagotomized patients. Comorbidity burden was classified as normal (0), moderate (1–2), or severe (3+), according to the number of comorbid categories present.

All participants were followed from the index date until AMD diagnosis, death, emigration, or December 31, 2012, whichever occurred first. Cumulative incidence curves were generated using Kaplan-Meier analysis with death treated as a competing risk and smoothed for visualization using splines. Because proportional hazards assumptions were not met in the first five years, analyses were restricted to patients surviving ≥10 years after vagotomy and their matched controls. Hazard ratios for AMD were estimated with Cox proportional hazards regression, adjusted for age, sex, COPD, hypertension, obesity-related diseases, and CCI category. Sensitivity analyses were performed, stratified by COPD as a proxy for smoking, by obesity-related diseases, and by hypertension. E-values were calculated to assess the potential effects of unmeasured confounders, such as smoking (VanderWeele & Ding, 2017).

The study is registered at Aarhus University’s Record of Processing Activities (j.nr. 2022-0367531, record number 2234). Studies based exclusively on data from the Danish national registers do, by Danish legislation, not require approval from the Danish Health Research Ethics committees.

### Animals

C57BL/6J mice (Jax strain #000664) were purchased from the Janvier Breeding Center. *TRE2* mice, which express the human APOE2 isoform, were engineered using targeted replacement as previously described (Levy et al., 2015). *Rag2^-/-^* mice were purchased from Charles River Laboratories (Jax strain #008449). All mice were negative for the *Crb1^rd8^*, *Pde6b^rd1^*, and *Gnat2^cpfl3^* mutations. Mice were housed under specific pathogen-free conditions in a 12 hr/12 hr light/dark cycle with water and food available ad libitum. All experimental protocols and procedures were approved by the French Ministry of Higher Education, Research and Innovation (APAFIS authorization #19412 2019022215422220).

### Surgeries (vagotomy, splenectomy, splenic denervation, and sham)

Mice received a subcutaneous injection of buprenorphine (0.05 mg/kg) 30 minutes before surgery, then were anesthetized with a mixture of ketamine (80 mg/kg) and xylazine (8 mg/kg). Mice received a second subcutaneous injection of buprenorphine (0.05 mg/kg) the day after surgery. Antiseptic solution (Vetedine, povidone iodine) was applied to all skin sutures immediately after closure. Mice were allowed to fully recover for at least 1 month following surgery before any further experiments.

For vagotomies, a 1 cm midline cervical incision was made, the left vagus nerve was cut, and then the incision was closed with absorbable sutures. For sham vagotomies the left vagus nerve was visualized but not cut, and were otherwise identical.

Splenectomies and sham splenectomies were performed as previously described (Roubeix et al., 2024). In brief, a 1 cm incision was made on the left side of the abdomen under the rib cage, the spleen was removed by cutting the connective tissue and cauterizing the splenic vessels (or only visualized without removal for sham splenectomies), and finally the incisions in the abdominal muscles and skin were each closed with absorbable sutures.

Splenic denervation was achieved using ethanol, similarly to previous studies (Zhang et al., 2020). In brief, an incision was made in the left abdomen as for splenectomy, and then the spleen was isolated from the abdominal cavity using blunt forceps, while cotton pads moistened with physiological saline were used to protect the abdominal opening and organs. Splenic denervation was achieved by applying a cotton swab dipped in absolute ethanol to the vascular trees and corresponding nerve fibers, approximately 7 times, 5-10 seconds per application. For sham denervation the cotton swab was instead dipped in physiological saline solution (0.9% NaCl). Finally, the spleen was returned to the abdominal cavity and incisions were closed as for splenectomies.

### Laser injury and bright light challenge

Laser injury and bright light challenge were performed as previously described (Roubeix et al., 2024). For laser injury, mice received eye drops of mydriaticum and neosynephrine to dilate the pupils, and then were anesthetized with a mixture of ketamine (80 mg/kg) and xylazine (8 mg/kg). Lubrithal was placed on the eye to maintain ocular surface moisture. For immunohistochemistry 4 laser injuries were performed per eye in the mid-periphery using a photocoagulation laser (Vitra Laser, 532 nm, 450 mW, 50 ms, 250 mm). For single-cell RNA sequencing and flow cytometry experiments, 10 laser injuries were performed per eye. For bright light challenge, the *TRE2* mice received daily eyedrops of atropine (1%, Novartis) to dilate their pupils, and were exposed to green LED light (4500 lux) for 4 days.

### Immunohistochemistry

Eyes were fixed in 4% PFA at room temperature (RT) for 45 minutes before storage in PBS and dissection. Anterior segments and lens were removed, and after 4 curvature-relieving cuts were made the retinas and RPE/choroid/sclera were carefully separated and then incubated overnight at RT in PBS with 1% Triton X-100 with the following primary antibodies (dilutions indicated in parentheses): rabbit anti-IBA1 (Wako, 1:400) and rat anti-CD102 (BD Biosciences, 1:200). After 3 PBS rinses, retinas and RPE/choroid/sclera complexes were incubated for 2 hours at RT with Alexa Fluor-conjugated secondary antibodies (Thermo Fisher Scientific, diluted 1:500) and Hoechst (Sigma Aldrich, diluted 1:1000) in PBS. After 3 more PBS rinses, tissues were flatmounted and coverslipped with an aqueous mounting medium (Sigma Aldrich) and imaged using a fluorescence microscope (Leica, DM5500). Quantifications of MP counts and CNV size were completed in FIJI. MPs and CNV size were quantified for individual laser impacts and were then averaged by eye for further statistical analyses and graphical representation.

For spleen cryosections, spleens were flash frozen in liquid nitrogen, embedded in Neg-50 optimal cutting temperature compound (Fischer Scientific), and then 12 μm sections were cut on a Leica CM3050 cryostat. Sections were fixed on the slide with 4% PFA for 15 minutes at RT, blocked for 1 hour at RT in blocking solution (PBS + 0.1% Triton X-100 + 5% normal horse serum), and then incubated overnight at 4 C with rabbit anti-tyrosine hydroxylase (Sigma-Aldrich, 1:200) and human anti-mouse CD3 (REAfinity, PE-conjugated, Miltenyi, 1:100) diluted in blocking solution. After 3 PBS rinses, sections were incubated for 2 hours at RT with Alexa Fluor-conjugated secondary antibodies (Thermo Fisher Scientific, 1:500) and Hoechst (Sigma Aldrich, 1:1000) in blocking solution. After 3 more PBS rinses slides were coverslipped with an aqueous mounting medium (Sigma Aldrich) and imaged using a fluorescence microscope (Leica, DM5500).

### Galantamine treatment

Galantamine, an acetylcholinesterase inhibitor (Sigma Aldrich), was resuspended in physiological 0.9% NaCl solution at a concentration of 1 mg/ml and then mice were given daily IP injections of Galantamine (4 mg/kg) or an equal volume of saline for control.

### Flow cytometry and fluorescence-activated cell sorting

Flow cytometry of eyes and spleens was performed as previously described (Roubeix et al., 2024). In brief, spleens were mechanically dissociated on a 70-µm cell strainer (Miltenyi), red blood cells were lysed using ammonium chloride (StemCell Technologies), and after rinsing in PBS and centrifugation (500 g, 5 min, 4 C), cells were resuspended in PBS. Eyes were dissected to remove posterior connective tissue from the sclera, the anterior segment, and the lens, and the resulting posterior segment eye cups were enzymatically dissociated using Liberase TL in PBS (0.8 Wunsch unit/ml, Sigma-Aldrich) followed by gentle mechanical dissociation using a P1000 pipette. The resulting cell suspension was passed through a 70-µm filter, rinsed with PBS, and resuspended in PBS following centrifugation (500 g, 5 min, 4 C). Cells were labeled using a live/dead fixable reagent (Miltenyi) following manufacturer recommendations, then incubated with the relevant antibodies (all diluted 1/50 in PBS) for 30 minutes at 4 C. The antibodies used in this study include: anti-CD45 VioBlue, anti-CD11b PE, anti-Ly6C PE-Vio770, anti-Ly6G APC-Vio770, anti-CD3 FITC, anti-CD8a PE, and anti-CD4 PE-Vio770 (all from Miltenyi). For flow cytometry, cells were fixed in 1% PFA, data acquisition was performed on a Celesta SORP cytometer (BD Biosciences), and data were analyzed with FlowJo 10.9. For cell sorting, cells suspensions were prepared as described for flow cytometry but without the final fixation step. Cells were sorted into chilled tubes using a FACSMelody cell sorter (BD Biosciences).

### Bulk mRNA sequencing

CD11b^+^CD45^+^Ly6C^neg^Ly6C^high^ splenic monocytes were sorted, as described above. RNA was extracted and RNA quality was confirmed using a Bioanalyzer RNA Analysis kit (Agilent). RNA sequencing libraries were prepared from 2 ng total of RNA using the Smartseq V4 (Takara) and Nextera XT (Illumina) kits and sequenced at the Institut du Cerveau iGenSeq sequencing facility.

After sequencing, fastq files were aligned to the *Mus musculus* reference genome from Ensembl (Ensemble 106, June 2022) using STAR (version 2.7.9a) with the option “- - quantMode Gene Counts” to extract the raw counts for each gene, and all count files were concatenated into a single file. The count file and descriptive sample file were loaded into our in-house R Shiny application “EyeDVseq” for analysis: the genes with a total count below 10 across samples were filtered out and the differential gene expression analysis was performed using DESeq version 1.40.2 integrated into EyeDVseq, comparing sham as control and vagotomy as the condition. The resulting gene list was then filtered, with |log_2_(FoldChange)| > 0.5 and pAdj values < 0.05 being considered significant.

### Single-cell mRNA sequencing and analysis

Live CD11b^+^CD45^+^ MPs were sorted from eyecups with and without laser, as described above. Single cell mRNA sequencing (scRNAseq) libraries were prepared from sorted cells using the Chromium Next GEM Single Cell 3’ Library v3.1 and dual index TT kits (10X Genomics) according to manufacturer’s recommended protocol. Resulting libraries were sequenced on an Illumina Novaseq 6000. The fastq files were processed using CellRanger v6.1.2 with default options (alignment, count). The resulting expression matrices were merged and analyzed with Seurat v4 (merging, QC filtering, normalization, PCA, clustering, and UMAP). The resulting Seurat object was used to create an H5AD file that was loaded into cellxgene. Further analyses, including differential gene expression analyses, and visualizations were performed in cellxgene using the cellxgene_VIP plugin. Cells other than MPs (including rod photoreceptors and other retinal neurons) were excluded and then the remaining MPs were re-clustered. Differentially expressed genes were identified using the cellxgene Welch’s t test, and the resulting gene lists were filtered with adjusted p-value < 0.05 and |log Fold Change| > 0.2 considered significant. Processing scripts are available on request.

### Statistical analysis

GraphPad Prism 10 was used for data analysis and graphic representation, unless otherwise stated (see above for details of the cohort study and RNA sequencing analyses). Data are presented as mean ± standard error of the mean (SEM). Differences between two groups were analyzed using an unpaired Mann-Whitney test and differences between multiple groups were analyzed using a one-way ANOVA followed by a Tukey’s multiple comparisons test. The n and p-values are indicated in the figure legends.

## Acknowledgements

We would like to thank Daniela Carnevale for initial training in performing cervical vagotomies in mice. We gratefully acknowledge the Cordelier Histology, Imaging, Cytometry and Spatialomics (CHICS) core facility (Centre de Recherche des Cordeliers, U1138, INSERM, Sorbonne Université, Université Paris Cité, 75006 Paris, France) for histological preparation and multiplex immunohistochemistry. This work benefited from equipment and services from the iGenSeq core facility at ICM. We would also like to thank the Imaging Core Facility, the Phénotypage Cellulaire et Tissulaire platform, and the animal facility of the Institut de la Vision.

This work was supported by the Agence Nationale de la Recherche (ANR, France: IHU FOReSIGHT [ANR-18-IAHU-0001], ANR Osaging [ANR-180CE14-0311-02], and ANR KEVINplus [ANR-22-CE14-0015-01]), a Berthe Fouassier Fondation de France post-doctoral fellowship, the MSCA Seal of Excellence program at Sorbonne University (VagalToneAMD), Retina France, Fondation Valentin Hauÿ, the Carnot Institute (France), and the Institut National de la Santé et de la Recherche Médicale (INSERM, France).

