## Supplement for "Vagal Signaling Decline in Age-related Macular Degeneration Drives Spleen-Dependent Retinal Inflammation"

**Supplemental Table 1.** Baseline characteristics 10 years after vagotomy of patients that underwent vagotomy between 1977-1995 and paired control individuals with at least 10 years follow-up time after the index date.

| | Comparison cohort | | Vagotomy | | Truncal vagotomy | | Superselective vagotomy | |
|---|---|---|---|---|---|---|---|---|
| | N | % | N | % | N | % | N | % |
| Total | 85930 | 100.0 | 9465 | 100.0 | 4040 | 100.0 | 5080 | 100.0 |
| **Gender** | | | | | | | | |
| Female | 29765 | 35 | 3275 | 35 | 1570 | 39 | 1580 | 31 |
| Male | 56165 | 65 | 6195 | 65 | 2465 | 61 | 3500 | 69 |
| **Age groups at vagotomy, years** | | | | | | | | |
| 0 - 39 years | 22650 | 26 | 2330 | 25 | 690 | 17 | 1565 | 31 |
| 40 - 64 years | 55120 | 64 | 5990 | 63 | 2600 | 64 | 3150 | 62 |
| 65 - 79 years | 8015 | 9 | 1110 | 12 | 715 | 18 | 355 | 7 |
| 80+ years | 145 | 0 | 40 | 0 | 35 | 1 | 5 | 0 |
| **Calendar period at vagotomy** | | | | | | | | |
| 1977-1981 | 43700 | 51 | 4660 | 49 | 1820 | 45 | 2665 | 52 |
| 1982-1986 | 24730 | 29 | 2780 | 29 | 1130 | 28 | 1560 | 31 |
| 1987-1991 | 12890 | 15 | 1485 | 16 | 785 | 19 | 645 | 13 |
| 1992-1995 | 4610 | 5 | 540 | 6 | 300 | 7 | 205 | 4 |
| **Charlson Comorbidity Index** | | | | | | | | |
| Normal | 75635 | 88 | 7565 | 80 | 3050 | 76 | 4235 | 83 |
| Moderate | 9100 | 11 | 1630 | 17 | 835 | 21 | 735 | 14 |
| Severe | 1195 | 1 | 270 | 3 | 150 | 4 | 110 | 2 |
| **Diabetes mellitus** | | | | | | | | |
| No | 83975 | 98 | 9235 | 98 | 3920 | 97 | 4985 | 98 |
| Yes | 1955 | 2 | 230 | 2 | 120 | 3 | 95 | 2 |
| **Cardiovascular disease** | | | | | | | | |
| No | 80350 | 94 | 8485 | 90 | 3560 | 88 | 4620 | 91 |
| Yes | 5580 | 6 | 980 | 10 | 480 | 12 | 460 | 9 |

| | Comparison cohort | | Vagotomy | | Truncal vagotomy | | Superselective vagotomy | |
|---|---|---|---|---|---|---|---|---|
| | N | % | N | % | N | % | N | % |
| **Rheumatological diseases or arthrosis** | | | | | | | | |
| No | 83860 | 98 | 9150 | 97 | 3860 | 96 | 4960 | 98 |
| Yes | 2070 | 2 | 315 | 3 | 180 | 4 | 120 | 2 |
| **Peptic ulcer** | | | | | | | | |
| No | 84380 | 98 | 1140 | 12 | 430 | 11 | 665 | 13 |
| Yes | 1550 | 2 | 8325 | 88 | 3610 | 89 | 4415 | 87 |
| **Chronic obstructive pulmonary disease (COPD)** | | | | | | | | |
| No | 83785 | 98 | 8990 | 95 | 3815 | 94 | 4960 | 96 |
| Yes | 2145 | 2 | 475 | 5 | 225 | 6 | 220 | 4 |
| **Hypertension** | | | | | | | | |
| No | 83885 | 98 | 9150 | 97 | 3880 | 96 | 4940 | 97 |
| Yes | 2045 | 2 | 315 | 3 | 160 | 4 | 140 | 3 |
| **Obesity related diseases** | | | | | | | | |
| No | 84925 | 99 | 9290 | 98 | 3945 | 98 | 5010 | 99 |
| Yes | 1005 | 1 | 175 | 2 | 95 | 2 | 70 | 1 |

**Supplemental Table 2:** Incidence rates and hazard ratios of AMD for Vagotomy patients surviving 10 years after vagotomy and their comparison cohort. Unadjusted (adjusted for sex and age in the design) and adjusted rates (age, sex, CCI score). Stratified by chronic obstructive pulmonary disease. Hazard ratios were not calculated for COPD cohorts due to the low case numbers.

| Chronic obstructive pulmonary disease (COPD) | Exposure | Vagotomy type | No. of outcome | Incidence Rate (per 1000PYRs) | Unadjusted Hazard Ratio | Adjusted Hazard Ratio |
|---|---|---|---|---|---|---|
| No | Comparison cohort | all | 3005 | 2.38 (2.30-2.47) | | |
| No | Vagotomy | all | 345 | 2.94 (2.64-3.26) | 1.36 (1.21–1.53) | 1.35 (1.20–1.53) |
| No | Comparison cohort | TV | 1360 | 2.82 (2.67-2.97) | | |
| No | Vagotomy | TV | 165 | 3.87 (3.31-4.48) | 1.50 (1.26–1.78) | 1.49 (1.25–1.78) |
| No | Comparison cohort | SSV | 1525 | 2.08 (1.98-2.18) | | |
| No | Vagotomy | SSV | 165 | 2.36 (2.01-2.73) | 1.26 (1.06–1.50) | 1.26 (1.06–1.50) |
| Yes | Comparison cohort | all | 65 | 3.44 (2.67-4.31) | | |
| Yes | Vagotomy | all | 20 | 4.72 (2.80-6.14) | | |
| Yes | Comparison cohort | TV | 35 | 4.31 (3.04-5.81) | | |
| Yes | Vagotomy | TV | 5 | 3.17 (1.02-6.49) | | |
| Yes | Comparison cohort | SSV | 30 | 2.75 (1.83-3.85) | | |
| Yes | Vagotomy | SSV | 15 | 6.36 (3.30-10.26) | | |

**Supplemental Table 3.** International classification of disease codes used in the study.

| | ICD-8 | ICD-10 |
|---|---|---|
| All AMD | 377.10, 377.11 | H35.3; H35.3B; H35.3C; H35.3D; H35.3E; H35.3G; H35.3J; H35.3K; H35.3L; H35.3A+kckd05b* |
| **Charlson Comorbidity:** | | |
| Myocardial infarction | 410 | I21;I22;I23 |
| Congestive heart failure | 427.09; 427.10; 427.11; 427.19; 428.99; 782.49 | I50; I11.0; I13.0; I13.2 |
| Peripheral vascular disease | 440; 441; 442; 443; 444; 445 | I70; I71; I72; I73; I74; I77 |
| Cerebrovascular disease | 430-438 | I60-I69; G45; G46 |
| Dementia | 290.09-290.19; 293.09 | F00-F03; F05.1; G30 |
| Chronic pulmonary disease | 490-493; 515-518 | J40-J47; J60-J67; J68.4; J70.1; J70.3; J84.1; J92.0; J96.1; J98.2; J98.3 |
| Connective tissue disease | 712; 716; 734; 446; 135.99 | M05; M06; M08; M09; M30; M31; M32; M33; M34; M35; M36; D86 |
| Ulcer disease | 530.91; 530.98; 531-534 | K22.1; K25 |
| Mild liver disease | 571; 573.01; 573.04 | B18; K70.0-K70.3; K70.9; K71; K73; K74; K76.0 |
| Diabetes, Type 1 | 249.00; 249.06; 249.07; 249.09; 250.00; 250.06; 250.07; 250.09 | E10.0, E10.1; E10.9 |
| Diabetes, Type 2 | | E11.0; E11.1; E11.9 |

*kckd05b is a procedural code for intravitreal injection with angio-static medicine

**Supplemental Table 4.** Procedural indexing utilized by surgeons in Denmark at the time of the vagotomies.

| | All vagotomy | Truncal vagotomy | Superselective vagotomy |
|---|---|---|---|
| VAGOTOMIA TRUNCALIS ABDOMINALIS | o | o | |
| VAGOTOMIA TRUNCALIS ABDOMINALIS LAPAROSCOPICA | o | | |
| VAGOTOMIA SELECTIVA VENTRICULI | o | o | |
| VAGOTOMIA SELECTIVA VENTRICULI LAPAROSCOPICA | o | | |
| VAGOTOMIA AREAE ACIDOGENIS VENTRICULI | o | | o |
| VAGOTOMIA (FORSKELLIGE TYPER, ATYPISKE) | o | | |
| VAGOTOMIA ENDOSCOPICA | o | | |
| VAGOTOMIA AREAE ACIDOGENIS VENTRICULI ENDOSCOPIC | o | | |
| VAGOTOMIA | o | | |
| VAGOTOMY TRUNCALIS THORACALIS | o | | |

**Supplemental Table 5. Significantly differentially expressed genes in splenic monocyte after vagotomy.** List of genes significantly down- (negative logFC) or up- (positive logFC) regulated in splenic monocytes from vagotomized (VGX) mice compared to sham. Genes are ordered from largest to smallest fold change.

(See excel file “SupplementalTable5.xlsx”)

**Supplemental Table 6. Significantly differentially expressed genes in homeostatic microglia.** List of genes significantly up- (positive logFC) and down- (negative logFC) regulated in homeostatic microglia from vagotomized (VGX) compared to sham, from non-lasered eyes. Genes are ordered from largest to smallest fold change.

| Gene | logFC | p-value | padj |
|---|---|---|---|
| Gm42418 | 1 | 7.92E-132 | 2.56E-127 |
| Ccnd3 | 0.51 | 2.17E-81 | 7.00E-77 |
| Fkbp5 | 0.49 | 2.68E-105 | 8.67E-101 |
| Cmss1 | 0.42 | 1.23E-32 | 3.96E-28 |
| Cadm2 | 0.37 | 3.41E-71 | 1.10E-66 |
| Dmd | 0.32 | 1.18E-65 | 3.80E-61 |
| March1 | 0.32 | 2.82E-50 | 9.10E-46 |
| Abca9 | 0.32 | 1.55E-45 | 4.99E-41 |
| AY036118 | 0.31 | 7.39E-36 | 2.39E-31 |
| Tgfbr1 | 0.29 | 6.27E-37 | 2.02E-32 |
| Ssh2 | 0.26 | 5.52E-39 | 1.78E-34 |
| Mertk | 0.25 | 1.87E-30 | 6.04E-26 |
| Rp1 | 0.23 | 4.03E-36 | 1.30E-31 |
| Mycbp2 | 0.23 | 4.37E-29 | 1.41E-24 |
| Rpl23a | 0.23 | 4.76E-22 | 1.54E-17 |
| Rps23 | 0.23 | 1.68E-19 | 5.42E-15 |
| mt-Nd5 | 0.22 | 1.69E-21 | 5.45E-17 |
| Apoe | 0.22 | 1.42E-12 | 4.60E-08 |
| Hcn1 | 0.21 | 5.52E-36 | 1.78E-31 |
| Entpd1 | 0.21 | 7.11E-20 | 2.30E-15 |
| Gm34455 | -0.21 | 6.62E-27 | 2.14E-22 |
| Socs3 | -0.22 | 9.60E-15 | 3.10E-10 |
| Ier2 | -0.22 | 3.88E-12 | 1.25E-07 |
| Sgk1 | -0.23 | 3.23E-17 | 1.04E-12 |
| H3f3b | -0.23 | 1.91E-14 | 6.17E-10 |
| Cd83 | -0.23 | 2.08E-14 | 6.73E-10 |
| Ubc | -0.24 | 1.21E-16 | 3.89E-12 |
| Ccl3 | -0.24 | 3.04E-16 | 9.82E-12 |
| Atf3 | -0.24 | 9.73E-12 | 3.14E-07 |
| Ier5 | -0.25 | 1.58E-17 | 5.09E-13 |
| Fosb | -0.25 | 1.72E-13 | 5.54E-09 |
| Ccl4 | -0.25 | 1.20E-12 | 3.88E-08 |
| Jun | -0.27 | 5.85E-17 | 1.89E-12 |
| Nfkbiz | -0.27 | 3.51E-15 | 1.13E-10 |
| Plek | -0.3 | 7.21E-26 | 2.33E-21 |
| Dusp1 | -0.3 | 2.89E-16 | 9.33E-12 |
| Jund | -0.32 | 5.97E-30 | 1.93E-25 |
| Klf4 | -0.32 | 4.85E-23 | 1.57E-18 |
| Klf2 | -0.37 | 1.85E-25 | 5.96E-21 |
| Btg2 | -0.37 | 4.10E-23 | 1.32E-18 |
| Zfp36 | -0.39 | 3.64E-25 | 1.18E-20 |
| Junb | -0.44 | 1.89E-32 | 6.10E-28 |
| Egr1 | -0.46 | 1.69E-27 | 5.46E-23 |
| Fos | -0.53 | 1.24E-32 | 4.01E-28 |

**Supplemental Table 7. Significantly differentially expressed genes in activated microglia regulated in vagotomy compared to sham.** List of genes significantly up- (positive logFC) and down- (negative logFC) regulated in the activated microglia 2 cluster 1-day post-laser from vagotomized (VGX) compared to sham. Genes are ordered from largest to smallest fold change.
(See excel file "SupplementalTable7.xlsx")

**Supplemental Table 8. Significantly differentially expressed genes in activated microglia regulated in vagotomy compared to splenectomy.** List of genes significantly up- (positive logFC) and down- (negative logFC) regulated in the activated microglia 2 cluster 1-day post-laser from vagotomized (VGX) compared to vagotomy and splenectomy (VGX/SplX). Genes are ordered from largest to smallest fold change.
(See excel file "SupplementalTable8.xlsx")

**Supplemental Table 9. Significantly differentially expressed genes in infiltrating monocytes regulated in vagotomy compared to sham.** List of genes significantly up- (positive logFC) and down- (negative logFC) regulated in the infiltrating moncoytes cluster 1-day post-laser from vagotomized (VGX) compared to sham. Genes are ordered from largest to smallest fold change.
(See excel file "SupplementalTable9.xlsx")

**Supplemental Table 10. Significantly differentially expressed genes in infiltrating monocytes regulated in vagotomy compared to splenectomy.** List of genes significantly up- (positive logFC) and down- (negative logFC) regulated in the infiltrating monocytes cluster 1-day post-laser from vagotomized (VGX) compared to vagotomy and splenectomy (VGX/SplX). Genes are ordered from largest to smallest fold change.
(See excel file "SupplementalTable10.xlsx")

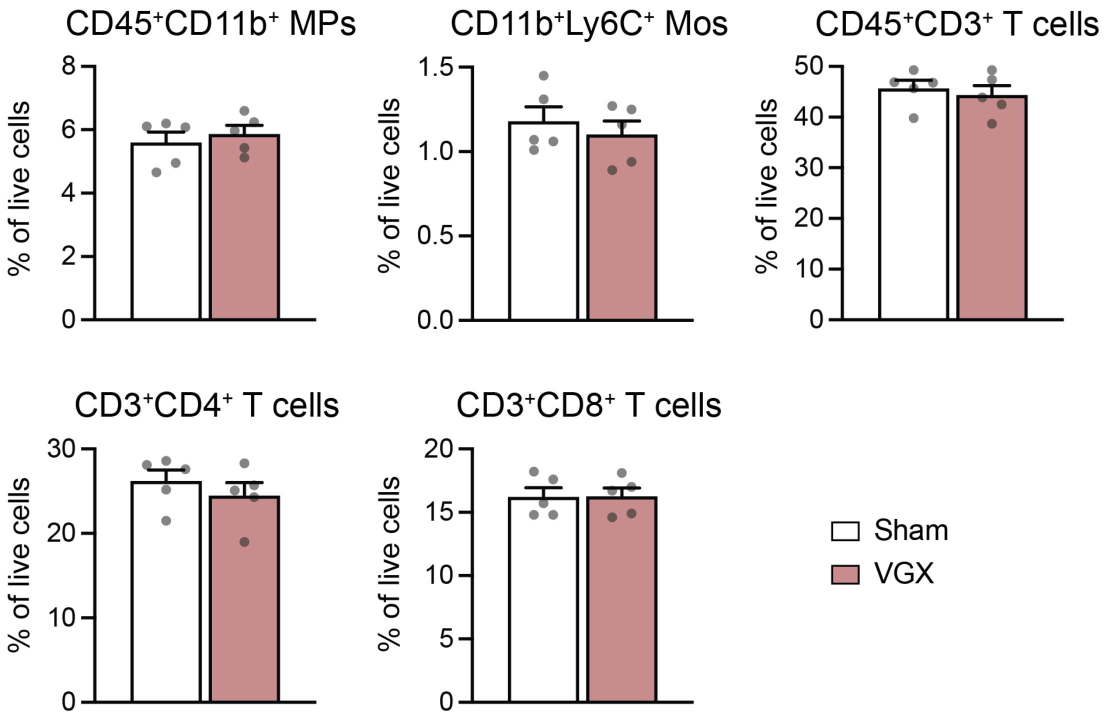

**Supplemental Figure 1. There are no changes in the proportion of immune cell types in the spleen after vagotomy.** Flow cytometric quantification of single, alive splenic $CD45^+CD11b^+$ mononuclear phagocytes (MPs), $CD45^+CD11b^+Ly6C^+$ monocytes (Mos), $CD45^+CD3^+$ T cells, $CD3^+CD4^+$ T cells, and $CD3^+CD8^+$ T cells from 3-month-old male mice 30 days after vagotomy (VGX) or sham surgery (n= 5 per group; Mann-Whitney: all comparisons not statistically significant; replicate represent spleens from different mice, all males).
