## Supplemental Table 5 for "Vagal Signaling Decline in Age-related Macular Degeneration Drives Spleen-Dependent Retinal Inflammation"

| Gene | log2FC | p value | padj | Mean (Sham) | Mean (VGX) |
|---|---|---|---|---|---|
| Gm23935 | -4.6867665 | 3.6847E-63 | 1.8452E-59 | 12195.4293 | 473.422547 |
| Gm24270 | -4.5603769 | 2.1564E-09 | 1.0404E-07 | 304.574686 | 12.9313431 |
| Gm15564 | -4.4365865 | 3.676E-63 | 1.8452E-59 | 2902.31005 | 133.977326 |
| Ighv1-52 | -4.3552095 | 1.797E-06 | 3.9124E-05 | 25.5348687 | 1.23456091 |
| Lars2 | -4.2380941 | 2.7697E-72 | 4.161E-68 | 20204.8923 | 1070.54829 |
| Ighv1-42 | -4.1683867 | 0.00197921 | 0.01311584 | 80.6017164 | 4.48084034 |
| Ssh3 | -3.5407876 | 1.0933E-07 | 3.3725E-06 | 23.2259207 | 1.9971779 |
| Apol7d | -3.1650221 | 7.0938E-09 | 3.0449E-07 | 917.873178 | 102.288753 |
| Plekhg6 | -3.0266257 | 3.2027E-08 | 1.1375E-06 | 26.4378182 | 3.23485764 |
| Abca4 | -2.9495653 | 2.613E-05 | 0.00038001 | 20.991047 | 2.65683417 |
| Cdc42bpb | -2.8940816 | 1.2661E-20 | 7.6085E-18 | 113.159623 | 15.2052459 |
| AC154912.3 | -2.739006 | 1.1568E-08 | 4.7098E-07 | 47.0849497 | 7.11412317 |
| AC154912.1 | -2.7101759 | 3.0994E-05 | 0.00043721 | 18.0963189 | 2.79251802 |
| Ddit4 | -2.5865366 | 1.7503E-18 | 6.4132E-16 | 467.513299 | 77.8038214 |
| Per1 | -2.5840061 | 3.655E-30 | 5.4909E-27 | 209.983439 | 35.0517627 |
| Gm19409 | -2.5579608 | 8.1162E-06 | 0.00014228 | 20.6077414 | 3.50236048 |
| Lgr4 | -2.4240877 | 0.00029289 | 0.00282423 | 23.8264398 | 4.379137 |
| Epn1 | -2.3805652 | 5.8795E-52 | 2.2082E-48 | 768.59269 | 147.447179 |
| Gm42798 | -2.3420559 | 0.00040411 | 0.00363746 | 18.9890278 | 3.74835814 |
| Gm15886 | -2.3120121 | 7.8533E-15 | 1.4045E-12 | 131.906481 | 26.4269345 |
| Prrc2a | -2.2639967 | 1.1599E-11 | 1.0434E-09 | 142.191184 | 29.484952 |
| Ryr3 | -2.25217 | 0.00063592 | 0.00528102 | 16.7866564 | 3.55201618 |
| Kntc1 | -2.2238843 | 4.602E-05 | 0.00061155 | 30.5146605 | 6.53058893 |
| Megf8 | -2.2227129 | 3.4179E-10 | 2.0215E-08 | 66.3200835 | 14.1988193 |
| Cxcr4 | -2.1566294 | 2.674E-20 | 1.4347E-17 | 486.660731 | 109.035008 |
| Tspoap1 | -2.1556329 | 0.00010755 | 0.00125442 | 24.6205916 | 5.5752793 |
| Gm45779 | -2.1436757 | 0.00016005 | 0.00171135 | 24.2015263 | 5.47471053 |
| Actn1 | -2.1303543 | 1.6283E-07 | 4.8155E-06 | 59.2226886 | 13.4992641 |
| Wdr90 | -2.1294991 | 2.7045E-06 | 5.5888E-05 | 58.3382478 | 13.3705639 |
| Mink1 | -2.1227777 | 3.4647E-14 | 5.3113E-12 | 232.368937 | 53.2327509 |
| AC154912.5 | -2.1135148 | 0.00109132 | 0.00819747 | 23.4633116 | 5.36146259 |
| Gm37893 | -2.1027246 | 0.00117505 | 0.00866609 | 25.7624304 | 6.02571804 |
| Msmp | -2.0955415 | 0.00011354 | 0.00130476 | 32.7957178 | 7.62780837 |
| Plxnd1 | -2.0934118 | 7.1938E-16 | 1.7154E-13 | 865.646281 | 202.774155 |
| Prpf19 | -2.093287 | 1.5489E-13 | 2.1347E-11 | 674.322029 | 157.928968 |
| Gm47260 | -2.0592017 | 4.604E-05 | 0.00061155 | 26.1932443 | 6.29873165 |
| Gm20511 | -2.0560496 | 1.4812E-15 | 3.225E-13 | 343.070734 | 82.3203235 |
| Zbtb42 | -2.0539677 | 7.3658E-13 | 8.578E-11 | 186.360741 | 44.7785295 |
| Gm43221 | -2.0518676 | 0.0004582 | 0.00404201 | 20.6711683 | 4.99008744 |
| Vegfa | -2.0410301 | 0.00029688 | 0.00285896 | 36.3185357 | 8.85367779 |
| Tle3 | -2.0209155 | 1.0464E-31 | 2.2457E-28 | 761.179928 | 187.514672 |
| Cebpa | -2.0061395 | 1.5866E-45 | 4.7671E-42 | 959.934021 | 238.815438 |
| Gm6649 | -1.9941349 | 0.00254914 | 0.01602334 | 15.9982229 | 4.05741755 |
| Fscn1 | -1.9887775 | 0.00128236 | 0.00929326 | 17.871636 | 4.52531191 |
| Efnb2 | -1.9875662 | 0.00095273 | 0.00729134 | 26.3973071 | 6.60777194 |

| | | | | | |
|---|---|---|---|---|---|
| Cebpe | -1.9752924 | 2.1608E-08 | 8.1153E-07 | 207.061559 | 52.7458585 |
| Gm12316 | -1.9699628 | 9.2242E-06 | 0.00015855 | 30.2325661 | 7.68973557 |
| Npm3 | -1.9697972 | 0.00012753 | 0.00142944 | 50.4674347 | 12.946014 |
| Ncor2 | -1.9588381 | 2.8395E-26 | 3.0469E-23 | 835.216138 | 214.557467 |
| Eml2 | -1.9546206 | 9.7253E-10 | 5.0555E-08 | 320.284366 | 82.4726019 |
| Plekhg3 | -1.9537467 | 1.7416E-24 | 1.3771E-21 | 901.015671 | 232.427016 |
| Dach1 | -1.9516902 | 0.00202547 | 0.0133576 | 18.5196331 | 4.80310198 |
| C1qtnf6 | -1.9503782 | 0.00077474 | 0.00615164 | 24.8282774 | 6.34929894 |
| Gm44949 | -1.9349259 | 1.5313E-06 | 3.3929E-05 | 50.7498339 | 13.3006604 |
| Adgrg6 | -1.9280321 | 5.3783E-06 | 0.00010151 | 45.364043 | 11.8846793 |
| Ski | -1.9207842 | 3.6749E-12 | 3.8075E-10 | 864.523568 | 228.378735 |
| Gm42418 | -1.9198387 | 9.3915E-05 | 0.00111709 | 10217.9804 | 2700.4353 |
| Phf2 | -1.919613 | 5.4043E-13 | 6.6007E-11 | 272.452739 | 71.9937232 |
| Sbf1 | -1.9190599 | 2.4435E-14 | 3.864E-12 | 990.539387 | 261.782432 |
| Arhgap23 | -1.9182266 | 7.1131E-16 | 1.7154E-13 | 320.884993 | 84.8324813 |
| Fn1 | -1.9105207 | 1.498E-14 | 2.473E-12 | 31075.7647 | 8265.96158 |
| Gm42839 | -1.9049206 | 9.4094E-06 | 0.00016125 | 38.186541 | 10.1564654 |
| Gm20481 | -1.9026929 | 0.00018177 | 0.00189505 | 311.424343 | 83.2922916 |
| Zc3h3 | -1.9006458 | 7.2675E-19 | 2.9508E-16 | 183.470066 | 49.1691456 |
| Ly75 | -1.898766 | 7.718E-08 | 2.4935E-06 | 182.711733 | 49.0173981 |
| Rybp | -1.8938817 | 2.7591E-10 | 1.6514E-08 | 146.716938 | 39.3680227 |
| Ttyh3 | -1.8854553 | 4.4128E-22 | 2.7622E-19 | 647.852359 | 175.111729 |
| Xk | -1.8812522 | 0.00022567 | 0.00227839 | 57.6664393 | 15.6022986 |
| Sik1 | -1.8788605 | 9.9801E-25 | 8.3295E-22 | 854.481462 | 232.077501 |
| Rbm38 | -1.8771601 | 2.4531E-15 | 4.9992E-13 | 252.97743 | 68.6543069 |
| CT010467.1 | -1.8761138 | 1.0936E-06 | 2.5283E-05 | 11427.8308 | 3113.139 |
| Zfp691 | -1.8731232 | 0.00163983 | 0.01135782 | 25.1394769 | 6.80451414 |
| Samd1 | -1.8713949 | 1.0735E-06 | 2.4889E-05 | 97.8329673 | 26.7350013 |
| Nup210l | -1.8698026 | 0.00190237 | 0.01273586 | 19.0507491 | 5.18736013 |
| Caskin2 | -1.8674295 | 0.00041618 | 0.00372606 | 38.507548 | 10.5738735 |
| Notch1 | -1.8672267 | 1.9891E-15 | 4.269E-13 | 754.336698 | 206.629284 |
| Nup188 | -1.8646083 | 1.2693E-16 | 3.4051E-14 | 268.810494 | 73.7268681 |
| BC037034 | -1.8606315 | 1.1161E-25 | 1.1178E-22 | 538.421248 | 148.366038 |
| Hspa1a | -1.8547226 | 0.00104349 | 0.00790933 | 739.957963 | 204.570195 |
| Gm37645 | -1.8443362 | 0.00213236 | 0.01394619 | 25.4401907 | 7.14664247 |
| Igkv4-53 | -1.8441241 | 0.00691549 | 0.0345269 | 26.4307622 | 7.34025414 |
| Rab35 | -1.841015 | 1.2492E-12 | 1.3901E-10 | 360.712173 | 100.727598 |
| Gm14453 | -1.8404536 | 1.0208E-06 | 2.3813E-05 | 43.1372571 | 12.0481134 |
| Hbb-bt | -1.8368812 | 0.0080758 | 0.03867476 | 105.119796 | 29.4267733 |
| Clip2 | -1.8337442 | 0.00024177 | 0.00241763 | 23.3419502 | 6.57652172 |
| Shb | -1.8164508 | 1.0925E-14 | 1.8864E-12 | 112.546509 | 31.8590466 |
| mt-Nd3 | -1.8134322 | 0.00802825 | 0.03854811 | 16.6040447 | 4.72331149 |
| Gm13609 | -1.8092016 | 1.802E-05 | 0.00027681 | 41.4164153 | 11.831197 |
| Rhob | -1.7963081 | 2.7241E-12 | 2.9442E-10 | 482.876971 | 138.982084 |
| Gm12898 | -1.7952397 | 2.6473E-05 | 0.00038462 | 42.1387517 | 12.1020034 |
| Dip2a | -1.792759 | 1.6923E-06 | 3.7006E-05 | 156.017185 | 45.0381091 |

| | | | | | |
|---|---|---|---|---|---|
| E2f8 | -1.7893422 | 9.85E-09 | 4.0991E-07 | 173.215397 | 50.0870785 |
| Hspa1b | -1.782096 | 0.00304346 | 0.01839174 | 1361.84753 | 395.960635 |
| Arap3 | -1.7775159 | 1.0984E-08 | 4.5086E-07 | 256.636293 | 74.809404 |
| Med12 | -1.7740509 | 4.3519E-24 | 3.2689E-21 | 447.491014 | 130.781576 |
| Gm27241 | -1.768195 | 0.00069449 | 0.00566721 | 30.5968466 | 8.98590445 |
| Gm10814 | -1.766263 | 5.9017E-12 | 5.7201E-10 | 112.969587 | 33.1313109 |
| Tst | -1.7636376 | 0.00498434 | 0.02687714 | 16.0249994 | 4.66669124 |
| Gm43071 | -1.757615 | 0.00052748 | 0.00452564 | 26.346764 | 7.83809858 |
| Gm15347 | -1.757367 | 0.00025182 | 0.00249216 | 28.0579217 | 8.3078544 |
| Gm45371 | -1.7570442 | 0.00027018 | 0.00264429 | 21.969375 | 6.48903235 |
| Gm37069 | -1.7552715 | 0.00266782 | 0.01663013 | 20.4140102 | 6.07921946 |
| Gm43359 | -1.7507237 | 0.00071593 | 0.00580747 | 29.5535413 | 8.83715289 |
| Gm42715 | -1.7467422 | 0.00644834 | 0.03274299 | 17.7784758 | 5.35431332 |
| Prkcg | -1.7416332 | 0.00075234 | 0.00602474 | 34.1012626 | 10.1750556 |
| Zc3h12a | -1.7403304 | 3.6812E-19 | 1.5362E-16 | 212.382462 | 63.5710751 |
| Gm43637 | -1.7362676 | 0.01141323 | 0.0497362 | 17.5114363 | 5.28003361 |
| Gm11521 | -1.7355966 | 0.00137129 | 0.00979592 | 23.4023107 | 7.02642997 |
| Gm14471 | -1.7321567 | 0.00033383 | 0.00311834 | 28.2051872 | 8.46357051 |
| Gm10388 | -1.7254855 | 0.00029506 | 0.00284332 | 26.3103296 | 7.90839062 |
| Gm9823 | -1.7235777 | 9.9912E-06 | 0.0001696 | 34.6155614 | 10.424376 |
| Atg2b | -1.7171021 | 1.3121E-12 | 1.4494E-10 | 362.936125 | 110.372172 |
| Rarg | -1.7159996 | 1.4587E-18 | 5.4783E-16 | 563.299126 | 171.256733 |
| Mapkbp1 | -1.7148202 | 1.912E-08 | 7.2354E-07 | 81.3870895 | 24.849814 |
| Gm15445 | -1.7113986 | 0.00032608 | 0.00306361 | 24.5443189 | 7.513445 |
| mt-Ty | -1.710949 | 1.0272E-10 | 7.1063E-09 | 94.2353597 | 28.7892733 |
| Them6 | -1.7092351 | 1.4263E-08 | 5.5656E-07 | 60.6680352 | 18.639576 |
| Abca7 | -1.7087919 | 1.1776E-18 | 4.5363E-16 | 3185.11269 | 974.317674 |
| Gm43813 | -1.7069904 | 1.409E-05 | 0.00022591 | 85.808428 | 26.3283072 |
| Gm20490 | -1.7051475 | 6.6462E-06 | 0.00012073 | 41.5761882 | 12.7475384 |
| Syne1 | -1.7006768 | 1.9847E-09 | 9.6493E-08 | 182.762968 | 56.2320783 |
| Capn1 | -1.699238 | 2.1942E-17 | 7.0133E-15 | 708.302402 | 217.869253 |
| Ulk1 | -1.6983925 | 2.0717E-14 | 3.3109E-12 | 775.582348 | 238.811292 |
| Zfp36l2 | -1.6967965 | 1.0049E-35 | 2.5161E-32 | 5970.88187 | 1841.57399 |
| Frat1 | -1.6939654 | 1.4261E-07 | 4.2762E-06 | 74.6988656 | 23.0087333 |
| Gm37940 | -1.6909129 | 1.1895E-06 | 2.7158E-05 | 89.2909023 | 27.6736558 |
| Depp1 | -1.6874745 | 0.00132771 | 0.0095436 | 44.8292196 | 13.990144 |
| 1700056N10I | -1.685461 | 0.00043049 | 0.00383359 | 28.1536687 | 8.79486943 |
| E530011L22R | -1.6820818 | 5.5273E-05 | 0.00071645 | 88.805051 | 27.7474808 |
| Bahd1 | -1.6760358 | 9.7137E-08 | 3.024E-06 | 184.257201 | 57.6449834 |
| Gpr132 | -1.6751107 | 1.5172E-27 | 1.7532E-24 | 591.559958 | 185.165737 |
| Kif24 | -1.6723084 | 0.00159135 | 0.0110731 | 29.6325222 | 9.35053057 |
| Bag6 | -1.6693086 | 2.9685E-29 | 3.7163E-26 | 608.755381 | 191.252044 |
| Gm10847 | -1.6565721 | 0.00377782 | 0.02167844 | 15.7015781 | 4.97481992 |
| Anxa11 | -1.6530945 | 1.4487E-15 | 3.2006E-13 | 243.553451 | 77.2697125 |
| Cad | -1.6499922 | 1.0131E-06 | 2.3669E-05 | 210.295108 | 66.9846645 |
| Fmnl1 | -1.6412282 | 1.1696E-19 | 5.1678E-17 | 4539.47313 | 1455.05981 |

| | | | | | |
|---|---|---|---|---|---|
| Dnajb5 | -1.6390873 | 2.2669E-08 | 8.409E-07 | 153.562695 | 49.1521557 |
| Man2a2 | -1.6326289 | 1.8349E-17 | 5.9926E-15 | 1290.04583 | 415.867525 |
| Irgq | -1.632157 | 1.2075E-08 | 4.8515E-07 | 203.312851 | 65.3541141 |
| Plekhg2 | -1.628862 | 9.1805E-16 | 2.155E-13 | 259.076971 | 83.6995971 |
| Gm15929 | -1.6283404 | 0.01046016 | 0.04675786 | 16.8083409 | 5.42140551 |
| Fbxl14 | -1.6282471 | 3.4015E-25 | 3.1938E-22 | 632.136626 | 204.508671 |
| Foxp4 | -1.6274667 | 1.1139E-07 | 3.4221E-06 | 87.2715851 | 28.2660235 |
| Lrrc25 | -1.6248012 | 7.0011E-17 | 1.9477E-14 | 582.259129 | 188.759563 |
| Pip4k2b | -1.622487 | 1.7421E-05 | 0.00026815 | 60.5985648 | 19.6670883 |
| Msrb1 | -1.6192039 | 2.151E-18 | 7.6939E-16 | 2974.66193 | 968.121292 |
| CT025731.2 | -1.615925 | 0.0054473 | 0.02879479 | 28.455142 | 9.343485 |
| Eng | -1.612228 | 3.8541E-12 | 3.9232E-10 | 145.160559 | 47.3003104 |
| Olfml2b | -1.6120409 | 1.6886E-05 | 0.00026066 | 202.220166 | 66.1831954 |
| Hyou1 | -1.61161 | 2.452E-17 | 7.3672E-15 | 458.944064 | 150.109166 |
| Kmt2d | -1.6109963 | 3.0546E-14 | 4.7801E-12 | 856.87087 | 280.456274 |
| Nt5dc3 | -1.608664 | 0.00211907 | 0.01387137 | 29.3113425 | 9.68579803 |
| Pelp1 | -1.60541 | 1.4983E-13 | 2.0841E-11 | 148.183994 | 48.6069746 |
| Epg5 | -1.6050953 | 1.3159E-09 | 6.6339E-08 | 166.45323 | 54.8088325 |
| Foxm1 | -1.6043246 | 0.00123412 | 0.00901494 | 72.5323849 | 23.9060606 |
| Kif15 | -1.5973209 | 5.859E-05 | 0.00075361 | 106.014658 | 35.1066752 |
| Lonrf3 | -1.5959749 | 0.0041552 | 0.02323199 | 20.3235387 | 6.68205165 |
| Sipa1l3 | -1.5908237 | 7.868E-12 | 7.2963E-10 | 281.20305 | 93.3684552 |
| Zfp526 | -1.5888941 | 0.00388563 | 0.0221307 | 24.022684 | 7.96919068 |
| Elf4 | -1.5874584 | 2.6994E-16 | 6.6481E-14 | 1887.21566 | 627.803044 |
| Gm48855 | -1.5860341 | 0.00021154 | 0.00216334 | 57.181594 | 19.0798578 |
| Cic | -1.5859223 | 4.6623E-10 | 2.6632E-08 | 428.68855 | 142.64705 |
| Mcf2l | -1.5792114 | 1.1655E-10 | 7.9539E-09 | 147.571889 | 49.2966577 |
| Fam78a | -1.5784731 | 6.1552E-14 | 9.0657E-12 | 263.065587 | 88.0361357 |
| Washc1 | -1.5778514 | 6.6802E-10 | 3.597E-08 | 161.832763 | 54.1179649 |
| Tnks1bp1 | -1.5777726 | 0.00031195 | 0.00296611 | 48.8298476 | 16.2585168 |
| Was | -1.5774439 | 8.6595E-30 | 1.1826E-26 | 808.199131 | 270.661129 |
| Pole | -1.5766279 | 0.00305223 | 0.01842253 | 44.2972583 | 14.808093 |
| D630044L22F | -1.5752196 | 9.7225E-08 | 3.024E-06 | 163.192802 | 54.7179086 |
| Notch2 | -1.5710795 | 2.5211E-13 | 3.2935E-11 | 2306.70351 | 776.159826 |
| Rhou | -1.5705832 | 2.5365E-10 | 1.5243E-08 | 244.913391 | 82.4710708 |
| Vapb | -1.5676162 | 0.00054644 | 0.00465903 | 498.70895 | 168.259194 |
| Gm36992 | -1.5671033 | 0.00039599 | 0.003582 | 32.4077789 | 10.8590167 |
| Adcy9 | -1.5646606 | 0.0001221 | 0.00138541 | 72.5819552 | 24.6103314 |
| Gm4793 | -1.5620951 | 3.5681E-05 | 0.00049542 | 52.3668504 | 17.7337065 |
| Zfp512b | -1.5607287 | 4.6951E-09 | 2.131E-07 | 133.051385 | 45.1499694 |
| 9230112E08F | -1.5606694 | 0.00010103 | 0.00118764 | 41.5137856 | 14.0938935 |
| Ntng2 | -1.5591951 | 5.5924E-06 | 0.00010427 | 43.6353792 | 14.850592 |
| Pcnx3 | -1.5565287 | 1.1123E-19 | 5.0636E-17 | 652.529803 | 221.653698 |
| Abca2 | -1.5558874 | 4.9238E-10 | 2.7808E-08 | 425.500985 | 144.698297 |
| Rnf44 | -1.5554345 | 5.8741E-20 | 3.043E-17 | 767.454083 | 260.902665 |
| Zhx2 | -1.5548006 | 4.1037E-05 | 0.00055515 | 35.7888391 | 12.1393066 |

| | | | | | |
|---|---|---|---|---|---|
| Tns3 | -1.5521616 | 3.1414E-15 | 5.9737E-13 | 432.203618 | 147.26532 |
| Gm15513 | -1.5492368 | 0.00177606 | 0.0121006 | 36.2973244 | 12.384904 |
| Cenpf | -1.5485427 | 0.00233929 | 0.01498004 | 48.1636122 | 16.5224578 |
| Pbx2 | -1.547691 | 4.7458E-10 | 2.6904E-08 | 203.570608 | 69.4909453 |
| Rtn4 | -1.5468789 | 1.2862E-15 | 2.9277E-13 | 883.58264 | 302.369375 |
| Ngfr | -1.5465844 | 0.00920107 | 0.04255742 | 15.9017262 | 5.38151398 |
| Tbc1d24 | -1.5450271 | 1.1E-06 | 2.5345E-05 | 113.852666 | 39.11916 |
| Rtl5 | -1.5389415 | 2.2479E-06 | 4.7496E-05 | 85.6069822 | 29.4921512 |
| Mfsd4a | -1.5386851 | 0.00014011 | 0.00153752 | 110.751265 | 37.994721 |
| AC132455.1 | -1.5377167 | 2.0476E-15 | 4.2723E-13 | 164.784411 | 56.7065422 |
| Gm26904 | -1.537707 | 0.00013124 | 0.00146374 | 62.3144833 | 21.4387276 |
| Tecpr2 | -1.5375855 | 3.6691E-07 | 9.7142E-06 | 119.199127 | 40.9706552 |
| Zbtb34 | -1.5368775 | 1.4535E-14 | 2.4534E-12 | 219.170889 | 75.4469902 |
| Zswim8 | -1.532799 | 1.4749E-11 | 1.2882E-09 | 377.190312 | 130.444004 |
| Stk24 | -1.5300533 | 1.5672E-20 | 8.7202E-18 | 500.319909 | 173.22316 |
| Chdh | -1.5272021 | 0.01073257 | 0.04770278 | 35.3183854 | 12.2172356 |
| Firre | -1.5260769 | 0.00216288 | 0.01407842 | 40.5320092 | 14.1258087 |
| E230001N04I | -1.5250861 | 0.0001663 | 0.00176435 | 91.9677731 | 31.9214625 |
| Msl2 | -1.5240043 | 2.2225E-23 | 1.5899E-20 | 948.402985 | 329.621819 |
| Scrib | -1.5231344 | 6.8616E-13 | 8.0532E-11 | 249.372153 | 86.6226651 |
| Myh9 | -1.52134 | 2.8975E-13 | 3.7204E-11 | 2850.41558 | 992.845966 |
| H2-Q4 | -1.5208069 | 1.7015E-05 | 0.00026218 | 64.169042 | 22.3551919 |
| Capns1 | -1.5154863 | 2.4625E-15 | 4.9992E-13 | 1459.36821 | 510.446197 |
| Nlrp12 | -1.509095 | 0.00260067 | 0.01627232 | 25.2102962 | 8.90081873 |
| Ppp1r37 | -1.5072436 | 1.4164E-12 | 1.5532E-10 | 181.131381 | 63.7404747 |
| Anxa11os | -1.5026256 | 2.7685E-09 | 1.3038E-07 | 93.7068564 | 33.0940269 |
| Fancm | -1.5024976 | 0.00047164 | 0.00412669 | 61.4906006 | 21.6587555 |
| Wdr81 | -1.5023831 | 1.0297E-08 | 4.2614E-07 | 474.643394 | 167.59893 |
| Tnpo2 | -1.502025 | 3.5239E-11 | 2.7717E-09 | 240.586028 | 84.837327 |
| Maz | -1.5014128 | 2.1913E-19 | 9.4055E-17 | 510.246104 | 180.234449 |
| Mcm3ap | -1.4994669 | 6.5368E-09 | 2.83E-07 | 584.635499 | 206.722707 |
| Gm26917 | -1.4967943 | 4.8999E-11 | 3.6806E-09 | 12359.108 | 4379.08546 |
| Gtf3c1 | -1.4967438 | 8.8131E-14 | 1.2609E-11 | 499.117475 | 176.830916 |
| Tmem121b | -1.495371 | 0.00164863 | 0.01139778 | 38.3392166 | 13.5765281 |
| Gm11192 | -1.4949731 | 3.547E-05 | 0.00049339 | 41.6416506 | 14.7757082 |
| Fancd2 | -1.4933475 | 0.00953703 | 0.04368136 | 39.3046185 | 13.9167759 |
| Cpeb4 | -1.4929534 | 5.0366E-07 | 1.276E-05 | 122.282255 | 43.4352235 |
| Ifnlr1 | -1.4896275 | 0.0073713 | 0.03622475 | 35.4590725 | 12.6872144 |
| Gm47586 | -1.4825234 | 0.01109666 | 0.04874025 | 28.6437236 | 10.2585764 |
| Zfp142 | -1.4820305 | 1.4328E-08 | 5.5765E-07 | 213.804193 | 76.4166593 |
| Kcnab2 | -1.4818892 | 1.9122E-16 | 5.0398E-14 | 1429.82488 | 511.740138 |
| Ube2o | -1.4814278 | 1.5823E-10 | 1.0204E-08 | 200.264241 | 71.6050205 |
| Atg2a | -1.4806008 | 1.7441E-08 | 6.6502E-07 | 677.000242 | 242.50776 |
| Gm15956 | -1.4802682 | 0.00460967 | 0.02519137 | 21.9070049 | 7.89180384 |
| Tgfb1 | -1.4795413 | 9.2299E-25 | 8.1565E-22 | 4765.83165 | 1708.72815 |
| Mast2 | -1.4792504 | 8.233E-07 | 1.9853E-05 | 261.336511 | 93.5456466 |

| | | | | | |
|---|---|---|---|---|---|
| Ints1 | -1.4790261 | 2.1187E-10 | 1.3262E-08 | 389.439609 | 139.74534 |
| Arhgap19 | -1.4775423 | 6.357E-05 | 0.00080321 | 203.489598 | 73.0912499 |
| Gm11149 | -1.4762481 | 0.00407716 | 0.02293192 | 21.7510384 | 7.82011248 |
| 4933428G20I | -1.4756542 | 7.1667E-05 | 0.00089127 | 32.5923014 | 11.6890716 |
| Zbtb7b | -1.4745397 | 1.1839E-15 | 2.7363E-13 | 697.709023 | 250.910903 |
| Gm48978 | -1.473661 | 0.0054788 | 0.02892327 | 55.8671791 | 20.1839968 |
| Patl1 | -1.4734473 | 1.4972E-14 | 2.473E-12 | 334.42614 | 120.388692 |
| Gm15978 | -1.4694454 | 3.1356E-08 | 1.1189E-06 | 379.211018 | 136.858038 |
| Ly6g | -1.4671157 | 0.00074165 | 0.00595499 | 55.2992292 | 20.0438013 |
| Zfp598 | -1.4643963 | 5.2501E-15 | 9.7373E-13 | 270.921079 | 98.1437608 |
| Il6ra | -1.4638727 | 2.3786E-30 | 3.9704E-27 | 4374.59813 | 1585.60622 |
| Nav2 | -1.4626635 | 0.00879075 | 0.04121828 | 19.391544 | 7.08617649 |
| Peg13 | -1.4584371 | 6.4237E-12 | 6.1467E-10 | 253.654394 | 92.2773119 |
| Wdr7 | -1.4572309 | 1.5497E-11 | 1.3382E-09 | 463.100511 | 168.732816 |
| 9930024M15 | -1.4561735 | 0.00553993 | 0.02914086 | 20.5432045 | 7.52632057 |
| Angptl2 | -1.4491011 | 7.1911E-05 | 0.00089208 | 79.4862605 | 29.0953441 |
| Nbeal2 | -1.4487073 | 5.3597E-10 | 2.9711E-08 | 842.065379 | 308.41104 |
| Fasn | -1.4467911 | 8.8232E-12 | 8.0823E-10 | 342.882568 | 125.741152 |
| Nxpe3 | -1.4453364 | 6.5906E-06 | 0.00011987 | 71.5762526 | 26.3248543 |
| Aatk | -1.4440807 | 1.6634E-10 | 1.0589E-08 | 270.760727 | 99.5107169 |
| Gatad1 | -1.4429803 | 3.7384E-12 | 3.8467E-10 | 318.88741 | 117.111275 |
| Slc2a4rg-ps | -1.4424259 | 0.00116797 | 0.00862656 | 50.1379706 | 18.4908127 |
| AA467197 | -1.4391774 | 0.00351038 | 0.02044841 | 23.1081866 | 8.51063786 |
| Setd1b | -1.4389324 | 1.2104E-05 | 0.00019973 | 131.922672 | 48.6283945 |
| Fam220a | -1.4370047 | 3.9227E-08 | 1.3641E-06 | 101.746951 | 37.4777248 |
| Ldhd | -1.4370022 | 0.00236629 | 0.01510786 | 29.8086378 | 11.0148748 |
| Nacc1 | -1.4364779 | 1.0632E-12 | 1.2009E-10 | 297.281468 | 109.718054 |
| Pex26 | -1.4356881 | 1.3271E-11 | 1.1727E-09 | 176.699537 | 65.1349668 |
| Hnrnpa0 | -1.4342911 | 3.5273E-06 | 7E-05 | 221.549238 | 81.9733793 |
| Ppp1r9b | -1.4338305 | 6.4678E-12 | 6.1497E-10 | 673.294162 | 249.13609 |
| Ccp110 | -1.4326918 | 0.00010165 | 0.00119309 | 52.7091071 | 19.6123412 |
| Dusp18 | -1.4319228 | 2.3971E-06 | 5.0155E-05 | 126.292074 | 46.6292942 |
| Slc7a5 | -1.431871 | 3.4569E-12 | 3.6064E-10 | 178.656006 | 66.1987788 |
| Fbxo10 | -1.4313318 | 1.7005E-08 | 6.5003E-07 | 116.083832 | 42.9837164 |
| Ncam1 | -1.4312458 | 0.00115157 | 0.00853901 | 84.5229158 | 31.3537253 |
| Tnfsf12 | -1.430688 | 1.4688E-09 | 7.3064E-08 | 131.927653 | 48.9146221 |
| Pkdcc | -1.4306096 | 9.2467E-10 | 4.8234E-08 | 218.658996 | 80.9792061 |
| Gm13571 | -1.4295481 | 0.00639919 | 0.03257708 | 23.4142093 | 8.63070841 |
| Tnrc18 | -1.4280765 | 9.2765E-13 | 1.0638E-10 | 687.379817 | 255.448578 |
| Gm26789 | -1.4269121 | 0.00093147 | 0.00718348 | 134.80471 | 50.2296799 |
| Atrn | -1.4254585 | 1.2819E-13 | 1.8168E-11 | 342.648708 | 127.536919 |
| Celsr1 | -1.4227018 | 0.01098444 | 0.04842114 | 19.7244811 | 7.41248769 |
| Ska3 | -1.4214319 | 0.0006883 | 0.00563196 | 43.3423377 | 16.2211634 |
| Dhcr24 | -1.4186193 | 3.93E-05 | 0.00053576 | 101.322115 | 37.924383 |
| Mta1 | -1.4174568 | 2.2274E-11 | 1.859E-09 | 179.945752 | 67.2436657 |
| Ppp4r2 | -1.4172747 | 5.7802E-12 | 5.6387E-10 | 415.984687 | 155.760593 |

| | | | | | |
|---|---|---|---|---|---|
| Gan | -1.416945 | 4.2694E-06 | 8.2866E-05 | 139.473869 | 52.2471335 |
| Plxnb2 | -1.4149055 | 2.3995E-09 | 1.148E-07 | 6533.55058 | 2450.15729 |
| Gm42659 | -1.4145762 | 0.00460136 | 0.02515952 | 48.1831137 | 18.114784 |
| Acvr1 | -1.4144457 | 2.0167E-05 | 0.00030603 | 150.183963 | 56.1505398 |
| Sympk | -1.4137636 | 3.7362E-06 | 7.3756E-05 | 508.907193 | 190.764435 |
| Gdap10 | -1.4137223 | 2.2158E-07 | 6.2191E-06 | 303.274259 | 113.841043 |
| Khsrp | -1.413113 | 2.3779E-16 | 6.0547E-14 | 860.284503 | 322.73395 |
| Gm26546 | -1.412485 | 0.000402 | 0.00362722 | 33.2778919 | 12.4772434 |
| Mast3 | -1.4122174 | 4.9216E-18 | 1.6777E-15 | 1296.09246 | 486.84575 |
| Hipk3 | -1.4113795 | 5.6163E-13 | 6.8044E-11 | 1011.5513 | 380.179051 |
| Gm37254 | -1.4111689 | 6.2015E-05 | 0.00079062 | 73.4752998 | 27.7324097 |
| Csf2rb2 | -1.4076084 | 2.119E-16 | 5.4885E-14 | 678.051075 | 255.305655 |
| Pelo | -1.4066042 | 2.2192E-05 | 0.0003314 | 44.3247738 | 16.6944266 |
| Pigt | -1.4061712 | 2.956E-08 | 1.0649E-06 | 471.23803 | 177.674385 |
| A630072L19F | -1.4054338 | 1.7243E-06 | 3.7652E-05 | 147.684035 | 55.7027416 |
| Furin | -1.4020134 | 4.7529E-31 | 8.9254E-28 | 2019.23208 | 763.940497 |
| S100a8 | -1.4009686 | 0.0098848 | 0.04495891 | 27446.6991 | 10393.3819 |
| Fam129b | -1.399632 | 9.6784E-19 | 3.8263E-16 | 2011.06249 | 761.977894 |
| Smad3 | -1.3990983 | 2.4341E-13 | 3.2076E-11 | 394.845057 | 149.678513 |
| Gm16214 | -1.3968637 | 1.9399E-06 | 4.1812E-05 | 551.78184 | 209.464082 |
| 4930500M09 | -1.3968152 | 4.1986E-05 | 0.0005657 | 41.5956961 | 15.8272104 |
| Cluh | -1.3961674 | 1.1933E-08 | 4.8192E-07 | 653.598159 | 248.205641 |
| Actr2 | -1.3961456 | 1.4263E-20 | 8.2411E-18 | 4197.82563 | 1594.75304 |
| Gm38319 | -1.3944956 | 0.00738312 | 0.03624724 | 24.2232456 | 9.24603506 |
| Scd2 | -1.3936912 | 7.6402E-09 | 3.2458E-07 | 1230.49484 | 468.305921 |
| 4930404N11I | -1.3928713 | 1.2095E-08 | 4.8515E-07 | 272.356302 | 103.529578 |
| Tmed9 | -1.3919767 | 1.6142E-09 | 7.9768E-08 | 3068.41988 | 1169.01128 |
| Mfsd12 | -1.3879278 | 5.831E-13 | 7.008E-11 | 619.741022 | 236.662337 |
| Pabpn1 | -1.3872487 | 4.8668E-07 | 1.2413E-05 | 136.785407 | 52.3102645 |
| Gm42856 | -1.3867393 | 0.00388609 | 0.0221307 | 67.3137192 | 25.6829696 |
| Heatr5b | -1.3859736 | 1.1705E-10 | 7.9539E-09 | 452.832259 | 173.206009 |
| Prex1 | -1.3822383 | 6.0554E-09 | 2.6599E-07 | 5231.69273 | 2006.88795 |
| Atp11a | -1.3818512 | 9.159E-11 | 6.4949E-09 | 511.203752 | 195.931526 |
| Ncapg2 | -1.3767505 | 0.00036337 | 0.00334701 | 135.401155 | 52.1197474 |
| Rap1gap | -1.3752373 | 0.00959798 | 0.04390697 | 18.2581641 | 7.04129725 |
| Gm28198 | -1.3752283 | 0.00642461 | 0.0326733 | 19.9972952 | 7.65502073 |
| Tbkbp1 | -1.3750345 | 6.17E-20 | 3.0897E-17 | 454.959221 | 175.301929 |
| Rapgef2 | -1.3730518 | 4.1989E-12 | 4.2053E-10 | 290.35348 | 112.04112 |
| Rps6ka3 | -1.3729965 | 1.0641E-06 | 2.4707E-05 | 232.85568 | 89.8269258 |
| Gm44423 | -1.3720272 | 0.0005168 | 0.00445942 | 39.9451084 | 15.4286038 |
| Isoc2a | -1.3712857 | 0.00448835 | 0.02471719 | 18.0948584 | 6.98726534 |
| Gm45014 | -1.3704413 | 0.00300857 | 0.01821756 | 30.4888019 | 11.8510446 |
| Gm29477 | -1.3696087 | 0.00143239 | 0.01015996 | 33.6706795 | 13.0129834 |
| Zfp516 | -1.3685448 | 1.2329E-09 | 6.2786E-08 | 364.299901 | 141.075024 |
| Mknk2 | -1.3659378 | 2.2644E-17 | 7.087E-15 | 3460.56515 | 1342.45763 |
| Wdfy2 | -1.3637434 | 3.7828E-17 | 1.1143E-14 | 1066.67244 | 414.375962 |

| | | | | | |
|---|---|---|---|---|---|
| Mnt | -1.3633076 | 9.9426E-10 | 5.1329E-08 | 195.252668 | 75.721969 |
| Capn15 | -1.3630861 | 7.4974E-15 | 1.357E-12 | 497.672586 | 193.385832 |
| Ubqln2 | -1.3624524 | 4.9747E-09 | 2.2443E-07 | 141.583173 | 54.8999608 |
| Nup210 | -1.3624065 | 1.2331E-06 | 2.7907E-05 | 1244.56291 | 483.961207 |
| Gm44686 | -1.3619455 | 0.00460217 | 0.02515952 | 23.2485914 | 9.10579029 |
| Hsf2 | -1.3614561 | 1.2743E-06 | 2.8745E-05 | 61.9049318 | 24.1732987 |
| Myo9b | -1.3607347 | 5.4222E-14 | 8.0652E-12 | 1601.49347 | 623.588706 |
| Glp2r | -1.3563758 | 5.1982E-06 | 9.8602E-05 | 81.5285045 | 31.7887429 |
| Sort1 | -1.3562772 | 2.4452E-10 | 1.4872E-08 | 1970.57057 | 769.479289 |
| Smcr8 | -1.3561959 | 1.2334E-06 | 2.7907E-05 | 249.890462 | 97.4891842 |
| Alms1 | -1.3549086 | 0.00611407 | 0.0314992 | 28.7957567 | 11.2610688 |
| Fam189a2 | -1.3544326 | 0.00932333 | 0.04303053 | 30.1131361 | 11.7023358 |
| Cnot1 | -1.3537945 | 5.6823E-17 | 1.6107E-14 | 1194.26588 | 467.264749 |
| Bicral | -1.3532711 | 4.2243E-11 | 3.2214E-09 | 387.738244 | 151.704458 |
| Atrnl1 | -1.3530968 | 4.7553E-13 | 5.904E-11 | 769.749564 | 301.317416 |
| Tbc1d22b | -1.3529305 | 1.6928E-07 | 4.9865E-06 | 111.601291 | 43.7296804 |
| 1700126G02I | -1.3515261 | 2.3381E-05 | 0.00034572 | 85.4916206 | 33.3627563 |
| Foxj2 | -1.350421 | 5.5823E-11 | 4.1516E-09 | 820.713088 | 321.774349 |
| Ddx19a | -1.3503637 | 1.274E-08 | 5.0367E-07 | 130.937632 | 51.2161448 |
| Hoxb4 | -1.3502895 | 0.00705599 | 0.03510004 | 19.9054609 | 7.84898988 |
| Znfx1 | -1.3495041 | 3.0638E-11 | 2.488E-09 | 1695.68615 | 665.347463 |
| Adam19 | -1.3490558 | 2.5736E-07 | 7.0401E-06 | 339.446835 | 133.261272 |
| Utp14b | -1.3468822 | 0.00681743 | 0.03412803 | 32.2741826 | 12.6478268 |
| Dennd4b | -1.3467475 | 8.5289E-13 | 9.8562E-11 | 1295.84708 | 509.279833 |
| Dennd4a | -1.3462809 | 4.3642E-13 | 5.4636E-11 | 888.931634 | 349.671838 |
| Zdhhc8 | -1.3458605 | 1.9225E-07 | 5.5224E-06 | 103.219182 | 40.531238 |
| Lyl1 | -1.3451855 | 1.9826E-10 | 1.2515E-08 | 412.485075 | 162.310073 |
| Ice1 | -1.3451828 | 3.1183E-09 | 1.4474E-07 | 422.611892 | 166.246792 |
| Gm28875 | -1.3445444 | 9.1179E-10 | 4.7894E-08 | 271.460799 | 106.942971 |
| Pde8a | -1.343622 | 1.0859E-10 | 7.4493E-09 | 241.409528 | 95.0185092 |
| Gm2788 | -1.3420198 | 0.00390397 | 0.02220727 | 46.5804452 | 18.369281 |
| Pan3 | -1.3417655 | 3.2125E-14 | 4.9753E-12 | 420.839252 | 166.050043 |
| Flna | -1.3406661 | 2.8673E-09 | 1.3461E-07 | 22877.3623 | 9032.6892 |
| Nckap5l | -1.3403162 | 3.4001E-06 | 6.7925E-05 | 77.3832461 | 30.4358235 |
| Oas3 | -1.3392711 | 5.9098E-08 | 1.9513E-06 | 1095.29441 | 432.707731 |
| Ttll4 | -1.3383477 | 2.7777E-12 | 2.9807E-10 | 193.082106 | 76.2684156 |
| Gm45222 | -1.3364084 | 0.00778583 | 0.03774333 | 26.5306725 | 10.5317129 |
| Gm42934 | -1.33445 | 4.2823E-07 | 1.1111E-05 | 89.7512529 | 35.4878455 |
| Prom1 | -1.3340816 | 0.00117054 | 0.00863708 | 104.897247 | 41.698457 |
| Gm42735 | -1.3335643 | 0.00910502 | 0.04225665 | 24.677377 | 9.81782086 |
| Gm44260 | -1.3280521 | 5.4053E-06 | 0.00010189 | 60.9747002 | 24.2994105 |
| Fam217b | -1.3280145 | 4.2254E-06 | 8.2141E-05 | 185.711643 | 73.9273867 |
| Hdac7 | -1.3276333 | 2.6135E-15 | 5.2351E-13 | 321.338959 | 128.133375 |
| Fnip2 | -1.3273057 | 1.8159E-05 | 0.00027865 | 160.123842 | 63.6338147 |
| Urb2 | -1.3258865 | 4.4431E-06 | 8.5905E-05 | 132.670943 | 52.7195013 |
| Slc5a3 | -1.3255442 | 0.00014439 | 0.00157415 | 51.242521 | 20.50107 |

| | | | | | |
|---|---|---|---|---|---|
| Lsm14b | -1.3250647 | 0.0004596 | 0.0040508 | 72.17038 | 28.8307637 |
| Spg11 | -1.3247222 | 7.3123E-09 | 3.1297E-07 | 447.921746 | 178.710892 |
| Gm45413 | -1.3238554 | 0.00535363 | 0.02845984 | 28.8198544 | 11.5732471 |
| Lmtk2 | -1.3233896 | 1.6344E-06 | 3.5845E-05 | 152.253428 | 60.881252 |
| Gm45223 | -1.3216638 | 3.0509E-06 | 6.1735E-05 | 181.060116 | 72.5290102 |
| Trip6 | -1.3214109 | 4.1908E-05 | 0.00056515 | 162.022129 | 64.8846186 |
| Zfp628 | -1.3126405 | 0.0006238 | 0.00519761 | 40.260136 | 16.2156409 |
| Kdm5c | -1.3095617 | 2.3054E-08 | 8.5305E-07 | 753.654959 | 304.038097 |
| Arhgap21 | -1.3045481 | 1.0723E-05 | 0.00017999 | 153.805411 | 62.2108872 |
| 1700007J10R | -1.2996455 | 2.2338E-05 | 0.00033289 | 106.5689 | 43.2234647 |
| Rad54l2 | -1.2994387 | 0.0007031 | 0.00572976 | 68.9966909 | 28.1020854 |
| AC160637.1 | -1.2989248 | 7.0813E-06 | 0.00012695 | 78.029147 | 31.710833 |
| Med13 | -1.2972669 | 8.2925E-09 | 3.4994E-07 | 433.407552 | 176.34193 |
| Anks1 | -1.2968827 | 8.3948E-11 | 6.0342E-09 | 496.56915 | 202.099929 |
| Tiam1 | -1.2954749 | 5.7434E-15 | 1.0522E-12 | 613.310791 | 249.738451 |
| Sec16a | -1.2948001 | 1.0312E-10 | 7.1063E-09 | 670.353857 | 273.105589 |
| Gm48375 | -1.2930219 | 3.8659E-05 | 0.00052894 | 68.0525177 | 27.7096899 |
| Tgfbrap1 | -1.2922355 | 4.4312E-10 | 2.5506E-08 | 427.619259 | 174.47355 |
| Gm15334 | -1.2919312 | 0.00306764 | 0.01849323 | 27.5244209 | 11.1913692 |
| Zfp687 | -1.2913114 | 2.9598E-07 | 8.0116E-06 | 247.911998 | 101.214399 |
| Rab11fip5 | -1.2907618 | 1.2646E-10 | 8.4063E-09 | 306.252155 | 125.060214 |
| Gm37760 | -1.290528 | 0.00081951 | 0.00646956 | 49.5257863 | 20.3067124 |
| Gigyf1 | -1.2900249 | 7.5144E-06 | 0.00013328 | 287.890864 | 117.72933 |
| Vav2 | -1.2896247 | 3.4008E-07 | 9.0746E-06 | 453.546537 | 185.452685 |
| Klf16 | -1.2872426 | 0.00150478 | 0.0105915 | 71.7847577 | 29.3565166 |
| Kpna2-ps | -1.2870111 | 0.00027948 | 0.00271405 | 49.0560095 | 20.0338909 |
| Cep152 | -1.2863876 | 4.6859E-05 | 0.00062023 | 178.610474 | 73.2234553 |
| Dennd4c | -1.285112 | 7.5874E-08 | 2.4566E-06 | 290.458855 | 119.050624 |
| Kidins220 | -1.2839556 | 1.6351E-10 | 1.0453E-08 | 682.563568 | 280.246404 |
| Tmem94 | -1.2815159 | 2.9533E-11 | 2.4113E-09 | 619.377262 | 254.63825 |
| Mmp25 | -1.2810713 | 0.00401166 | 0.02265683 | 82.6362491 | 34.0452294 |
| Trio | -1.2803223 | 9.0438E-09 | 3.7835E-07 | 368.931667 | 151.878167 |
| Camsap1 | -1.2802135 | 5.1096E-07 | 1.2923E-05 | 171.373828 | 70.5403961 |
| Plagl2 | -1.2794005 | 4.5763E-12 | 4.523E-10 | 652.625941 | 268.64283 |
| Dgkd | -1.2787079 | 1.0157E-12 | 1.1559E-10 | 1377.17531 | 567.461187 |
| Agpat1 | -1.2787032 | 5.6483E-07 | 1.419E-05 | 310.852673 | 128.190991 |
| Lclat1 | -1.278522 | 4.428E-08 | 1.5153E-06 | 112.752777 | 46.4188427 |
| C3 | -1.2783595 | 1.1117E-09 | 5.7196E-08 | 10232.4472 | 4218.28753 |
| Zfp703 | -1.2772092 | 3.9049E-11 | 3.0239E-09 | 167.033486 | 68.8826289 |
| Tsc2 | -1.2770618 | 1.5927E-09 | 7.8966E-08 | 1012.37653 | 417.54679 |
| Gas7 | -1.2765895 | 8.7933E-20 | 4.1282E-17 | 826.191578 | 340.815572 |
| Heatr5a | -1.2757068 | 5.4699E-10 | 3.0101E-08 | 741.332995 | 306.214479 |
| Mthfr | -1.2746482 | 4.9527E-11 | 3.7017E-09 | 816.582943 | 337.354743 |
| Cpd | -1.2744465 | 1.4074E-05 | 0.00022589 | 312.971285 | 129.246786 |
| Urb1 | -1.272986 | 2.3798E-08 | 8.7411E-07 | 166.954644 | 68.9538743 |
| S1pr4 | -1.272884 | 9.6734E-08 | 3.0213E-06 | 201.25811 | 83.2221233 |

| | | | | | |
|---|---|---|---|---|---|
| Phactr4 | -1.2722646 | 9.9456E-07 | 2.3309E-05 | 119.664789 | 49.5499392 |
| Chml | -1.2719125 | 3.4305E-08 | 1.207E-06 | 134.342293 | 55.7298768 |
| Anapc1 | -1.2718823 | 1.3544E-08 | 5.3182E-07 | 1057.87039 | 437.953666 |
| Pds5a | -1.2716276 | 3.7322E-13 | 4.7116E-11 | 894.615688 | 370.607831 |
| Dennd2a | -1.268918 | 1.1063E-05 | 0.00018549 | 109.467534 | 45.2542037 |
| Rflnb | -1.2676842 | 1.0241E-05 | 0.00017325 | 443.022999 | 183.977781 |
| Rassf2 | -1.267109 | 2.0372E-15 | 4.2723E-13 | 1442.9845 | 599.506917 |
| 6430511E19F | -1.2667415 | 0.00106928 | 0.00806414 | 69.3791707 | 28.8808271 |
| Gm40309 | -1.2651285 | 0.0024202 | 0.01536066 | 87.7187876 | 36.5894986 |
| Zbtb40 | -1.2645548 | 6.8856E-06 | 0.00012396 | 157.798106 | 65.6710526 |
| Ppme1 | -1.2639123 | 9.103E-11 | 6.4949E-09 | 286.851626 | 119.222816 |
| Stk10 | -1.2618644 | 8.5329E-06 | 0.0001482 | 5936.61556 | 2475.48858 |
| Irf2bp2 | -1.2580595 | 0.0078368 | 0.03791388 | 2058.86931 | 860.876774 |
| Celf3 | -1.2574076 | 0.0078235 | 0.03786484 | 21.6069347 | 9.07598216 |
| Gm26759 | -1.2573901 | 1.2123E-05 | 0.00019973 | 299.996925 | 125.607 |
| A930004J17R | -1.2572429 | 0.00776113 | 0.03768438 | 33.6002696 | 14.127874 |
| Gm48582 | -1.2558739 | 0.00131006 | 0.0094484 | 66.1719725 | 27.818315 |
| Ankfy1 | -1.2557426 | 1.9588E-11 | 1.6532E-09 | 1634.59184 | 684.33827 |
| Nlrp3 | -1.2552077 | 4.1577E-11 | 3.1868E-09 | 372.932891 | 156.229632 |
| Gcnt7 | -1.2547152 | 1.2928E-05 | 0.00021087 | 80.7547245 | 33.9696144 |
| Zfp831 | -1.2540543 | 0.00900624 | 0.04191253 | 29.4983443 | 12.3358523 |
| Tram2 | -1.2538198 | 8.1701E-08 | 2.6282E-06 | 152.316319 | 63.7555075 |
| Gm26916 | -1.2513857 | 2.3046E-05 | 0.00034177 | 53.5651383 | 22.5374126 |
| Bcr | -1.2497975 | 3.6728E-07 | 9.7142E-06 | 225.157333 | 94.5267906 |
| 2610507B11F | -1.2492818 | 5.0255E-18 | 1.6777E-15 | 1525.85585 | 641.745715 |
| Itpripl1 | -1.2477615 | 4.8203E-13 | 5.9357E-11 | 260.265372 | 109.494839 |
| Gm37949 | -1.2472376 | 0.00882331 | 0.04131939 | 26.2510235 | 11.089685 |
| Gm43660 | -1.2441698 | 0.0027954 | 0.01723237 | 36.2357735 | 15.3131058 |
| Slc30a4 | -1.2439314 | 0.00331936 | 0.01958631 | 37.3954198 | 15.8150543 |
| Ttc7 | -1.2429284 | 3.1494E-08 | 1.1212E-06 | 5339.43298 | 2255.84003 |
| Ncoa6 | -1.2428389 | 1.2508E-11 | 1.1185E-09 | 413.520626 | 174.644613 |
| Klc3 | -1.2415432 | 0.00028372 | 0.00275166 | 54.4780039 | 23.1233388 |
| Siglec1 | -1.2373157 | 4.5752E-06 | 8.8145E-05 | 313.045982 | 132.662025 |
| Wipf1 | -1.2373025 | 4.3272E-14 | 6.5664E-12 | 1580.36897 | 670.333823 |
| Amotl2 | -1.2363488 | 2.2449E-05 | 0.00033391 | 108.594196 | 46.0451323 |
| Tab3 | -1.2352593 | 1.2222E-05 | 0.0002007 | 84.9566985 | 35.9966432 |
| Dtx2 | -1.2351315 | 2.5105E-10 | 1.5147E-08 | 526.534903 | 223.375048 |
| Dnajb9 | -1.2344042 | 3.0923E-11 | 2.4976E-09 | 358.303614 | 152.137414 |
| Fer1l5 | -1.2326425 | 3.9422E-07 | 1.0336E-05 | 151.506647 | 64.5463742 |
| Mogat2 | -1.2325398 | 0.00188493 | 0.01265292 | 89.1551107 | 37.9392503 |
| Ccdc88c | -1.2316271 | 7.5754E-09 | 3.2331E-07 | 1624.11207 | 691.541439 |
| Lmnb1 | -1.2302137 | 3.7807E-08 | 1.3178E-06 | 457.907936 | 195.148014 |
| Nufip2 | -1.2266023 | 2.2983E-10 | 1.4209E-08 | 1397.48381 | 597.030397 |
| Inpp4a | -1.2245972 | 8.4787E-07 | 2.0321E-05 | 513.099646 | 219.615636 |
| Gm26772 | -1.2244249 | 0.00407456 | 0.02292586 | 47.5624747 | 20.4354316 |
| Gm37297 | -1.222696 | 0.0011232 | 0.00838241 | 33.1529889 | 14.1635004 |

| | | | | | |
|---|---|---|---|---|---|
| Sgms2 | -1.2224696 | 6.0879E-09 | 2.6664E-07 | 881.246139 | 377.436876 |
| Rfx1 | -1.2218228 | 1.6394E-13 | 2.2389E-11 | 355.426023 | 152.23188 |
| Gfi1b | -1.2217048 | 0.01007007 | 0.04560827 | 43.1538406 | 18.5167677 |
| Edem3 | -1.2190016 | 4.3283E-08 | 1.4846E-06 | 375.461109 | 161.196368 |
| Atp8b2 | -1.2147971 | 2.1135E-09 | 1.0242E-07 | 895.530062 | 385.770314 |
| Ubr3 | -1.2147872 | 2.3226E-07 | 6.4495E-06 | 435.914968 | 187.64611 |
| Pde5a | -1.2134982 | 0.00010058 | 0.00118604 | 45.1700046 | 19.4420728 |
| Gm12000 | -1.2126882 | 0.00793345 | 0.0382491 | 30.0405764 | 13.0069421 |
| Zmiz1os1 | -1.2116735 | 8.8543E-06 | 0.00015307 | 108.669727 | 46.9208971 |
| Trim45 | -1.2114296 | 0.01131838 | 0.04947224 | 22.031947 | 9.54772225 |
| Rnf150 | -1.2109632 | 0.00108674 | 0.00816711 | 48.8994247 | 21.0693102 |
| Bcl6 | -1.2109422 | 1.2409E-10 | 8.2851E-09 | 1100.28842 | 475.112775 |
| Tln1 | -1.2095689 | 4.6908E-11 | 3.5412E-09 | 10357.6939 | 4478.53396 |
| Dop1b | -1.209313 | 3.6237E-11 | 2.8354E-09 | 418.121639 | 180.761013 |
| Arhgap39 | -1.2084553 | 9.1654E-11 | 6.4949E-09 | 794.644187 | 343.942141 |
| Piezo1 | -1.2065947 | 2.5303E-11 | 2.0772E-09 | 1068.5752 | 462.904147 |
| Ctdsp2 | -1.2064464 | 5.0788E-10 | 2.8577E-08 | 1160.04965 | 502.619061 |
| Sema6b | -1.2038887 | 0.0001944 | 0.00201549 | 94.270963 | 40.9078281 |
| Foxk2 | -1.203258 | 1.011E-11 | 9.1498E-10 | 356.225368 | 154.579831 |
| Fam120a | -1.2024924 | 7.1873E-12 | 6.7484E-10 | 2113.46124 | 918.203607 |
| Lrfn1 | -1.2024603 | 0.00041966 | 0.00375275 | 74.4650963 | 32.3855795 |
| Lmbr1l | -1.2021135 | 2.4997E-11 | 2.0645E-09 | 466.743108 | 202.755077 |
| Tmbim6 | -1.2018818 | 7.5291E-20 | 3.6487E-17 | 3337.51322 | 1450.77187 |
| Gm6225 | -1.1990919 | 0.00240751 | 0.01529302 | 28.7002485 | 12.5199345 |
| Adamts7 | -1.1979474 | 0.0039316 | 0.02232106 | 83.1787588 | 36.2063986 |
| Gpbp1l1 | -1.1966729 | 2.5542E-16 | 6.3954E-14 | 444.305244 | 193.842824 |
| Qser1 | -1.1962566 | 3.207E-05 | 0.00044985 | 123.364033 | 53.758704 |
| Zer1 | -1.1961857 | 1.1284E-08 | 4.6189E-07 | 477.804507 | 208.536684 |
| 4930579G18I | -1.1954645 | 0.00550359 | 0.02898016 | 30.2437528 | 13.2264543 |
| Trak1 | -1.1953093 | 8.0603E-08 | 2.5985E-06 | 695.881042 | 303.824737 |
| Itpripl2 | -1.1948571 | 2.4399E-07 | 6.7255E-06 | 418.905522 | 183.022253 |
| Morc2a | -1.1940114 | 1.7442E-11 | 1.4973E-09 | 307.798839 | 134.500511 |
| Usp31 | -1.189402 | 2.3651E-06 | 4.9554E-05 | 125.933746 | 55.2175462 |
| Rassf3 | -1.1888986 | 1.8782E-11 | 1.5941E-09 | 2695.12155 | 1181.95867 |
| Hipk1 | -1.1880915 | 1.828E-11 | 1.5604E-09 | 1415.63316 | 621.206597 |
| Tlnrd1 | -1.1878662 | 2.7534E-15 | 5.4426E-13 | 676.595833 | 296.733022 |
| Smap2 | -1.1873985 | 4.4E-22 | 2.7622E-19 | 1395.50664 | 612.561687 |
| Zfp275 | -1.1869624 | 0.00030706 | 0.00293333 | 73.9780937 | 32.3927022 |
| Clasp1 | -1.1868307 | 9.9662E-09 | 4.136E-07 | 371.998334 | 163.252227 |
| Sp4 | -1.1866921 | 6.3255E-06 | 0.00011561 | 101.102668 | 44.5129529 |
| Tarbp1 | -1.1863154 | 0.00131833 | 0.00949439 | 82.0117865 | 36.0417681 |
| Gm45732 | -1.1845751 | 0.0102029 | 0.04598806 | 24.2391836 | 10.6444112 |
| Dcaf5 | -1.1838016 | 1.6019E-07 | 4.7559E-06 | 686.391338 | 301.903135 |
| Zfp608 | -1.1831579 | 5.8194E-08 | 1.9257E-06 | 175.810796 | 77.441726 |
| Atp10a | -1.1826623 | 1.0432E-06 | 2.4259E-05 | 377.639129 | 166.183463 |
| Man2b2 | -1.1821061 | 3.4895E-11 | 2.7591E-09 | 400.705296 | 176.548151 |

| | | | | | |
|---|---|---|---|---|---|
| Gm11651 | -1.1820356 | 0.00610777 | 0.03147756 | 767.576581 | 338.219497 |
| Crocc | -1.1799172 | 0.00308406 | 0.01857729 | 72.2517825 | 31.8403903 |
| Hira | -1.1789318 | 0.01044267 | 0.04672172 | 29.4749607 | 13.0677422 |
| Ern1 | -1.1782236 | 6.6694E-23 | 4.5543E-20 | 623.424635 | 275.560781 |
| Utp20 | -1.1761269 | 1.6532E-05 | 0.00025669 | 144.796735 | 64.1671038 |
| Dnajb13 | -1.1751704 | 0.00067746 | 0.00555839 | 152.378985 | 67.5143897 |
| Gm10734 | -1.1751554 | 0.00023268 | 0.00234282 | 47.8166585 | 21.0895654 |
| Clcn6 | -1.1746881 | 0.00025566 | 0.00252348 | 98.2660869 | 43.5046341 |
| Zmiz2 | -1.1737998 | 7.8861E-10 | 4.1863E-08 | 1710.34422 | 758.009331 |
| Tkfc | -1.1737332 | 0.00251152 | 0.0158133 | 43.8845091 | 19.4524829 |
| Clasp2 | -1.1725098 | 0.00015477 | 0.00166552 | 321.704515 | 142.63751 |
| Uhmk1 | -1.1707983 | 2.1625E-10 | 1.348E-08 | 645.359939 | 286.681375 |
| Gm12523 | -1.1701907 | 0.00014235 | 0.00155601 | 76.1143972 | 33.6780534 |
| Gm37419 | -1.1693546 | 0.00500451 | 0.02696656 | 33.5395312 | 14.8744774 |
| Cnksr3 | -1.1692745 | 1.5371E-08 | 5.9211E-07 | 198.046803 | 87.8666068 |
| Dock11 | -1.1691286 | 2.8119E-06 | 5.7841E-05 | 1465.77994 | 651.709024 |
| Setd1a | -1.1689962 | 0.00317114 | 0.01895374 | 61.3496459 | 27.3132286 |
| Purg | -1.1688325 | 0.00068076 | 0.00558223 | 51.7087463 | 22.967073 |
| Gnaq | -1.1661866 | 2.11E-07 | 5.9696E-06 | 314.372619 | 140.141109 |
| Sfxn5 | -1.1654069 | 0.00049123 | 0.00427318 | 63.2813134 | 28.3173559 |
| Ace | -1.1646612 | 0.00251143 | 0.0158133 | 5107.54978 | 2278.24718 |
| Itga6 | -1.1646225 | 0.0022314 | 0.01440581 | 97.832062 | 43.5696652 |
| Gm44199 | -1.1626093 | 0.00038616 | 0.00351165 | 60.9259366 | 27.2825633 |
| Flnb | -1.162465 | 5.3754E-09 | 2.3892E-07 | 181.360856 | 80.9815818 |
| Tmem131 | -1.1622859 | 1.578E-10 | 1.0204E-08 | 1420.59587 | 634.57843 |
| Gm44164 | -1.1616003 | 0.00072545 | 0.00586254 | 59.2480506 | 26.5422594 |
| Abca3 | -1.1609107 | 1.3386E-11 | 1.176E-09 | 1433.16081 | 640.694166 |
| Rxra | -1.1608818 | 1.1554E-12 | 1.2953E-10 | 1002.67145 | 448.263213 |
| Birc6 | -1.1606695 | 8.1966E-10 | 4.3359E-08 | 705.309923 | 315.529633 |
| Gm42809 | -1.1602342 | 4.0461E-11 | 3.1172E-09 | 302.736791 | 135.325317 |
| Luc7l2 | -1.1595144 | 3.1864E-07 | 8.5633E-06 | 281.464443 | 126.105527 |
| Synj1 | -1.1591391 | 2.3905E-13 | 3.2043E-11 | 1649.02961 | 738.190121 |
| Zfhx2 | -1.1579745 | 0.00072259 | 0.00584886 | 51.9686799 | 23.2788436 |
| Gm37676 | -1.157878 | 0.01082967 | 0.04794994 | 32.254868 | 14.3913081 |
| Ywhag | -1.1575639 | 1.001E-16 | 2.7342E-14 | 2501.94259 | 1121.45168 |
| Mark4 | -1.1574522 | 9.5734E-06 | 0.00016362 | 217.010611 | 97.208017 |
| Cdv3 | -1.1563276 | 1.9582E-07 | 5.614E-06 | 687.171265 | 308.399981 |
| Arhgef11 | -1.1555249 | 3.63E-10 | 2.1386E-08 | 391.492739 | 175.689724 |
| Plxna1 | -1.1549597 | 5.1941E-05 | 0.00068089 | 113.514315 | 51.0312555 |
| Inppl1 | -1.1545127 | 1.8731E-08 | 7.1061E-07 | 261.937892 | 117.509273 |
| Gm16282 | -1.1533448 | 1.5826E-10 | 1.0204E-08 | 222.450884 | 99.9578255 |
| Itgb3 | -1.1530526 | 6.0272E-07 | 1.5042E-05 | 225.480728 | 101.326924 |
| Creb3l2 | -1.1530212 | 2.7791E-07 | 7.5497E-06 | 278.134095 | 125.020119 |
| 4921516A02F | -1.1524697 | 3.7189E-10 | 2.1824E-08 | 393.249768 | 176.870445 |
| Gm38115 | -1.1509751 | 1.4837E-05 | 0.00023562 | 74.7737723 | 33.6881559 |
| Zkscan8 | -1.1509414 | 0.00021981 | 0.00223273 | 75.9359359 | 34.2207845 |

| | | | | | |
|---|---|---|---|---|---|
| Rubcn | -1.1505123 | 7.3813E-07 | 1.7972E-05 | 540.766153 | 243.471768 |
| Atp13a3 | -1.1502317 | 9.2443E-10 | 4.8234E-08 | 688.234763 | 309.974194 |
| Gm26751 | -1.1496518 | 0.0038207 | 0.0218245 | 43.0575958 | 19.338508 |
| Gm43788 | -1.1494109 | 0.00316066 | 0.01893247 | 39.2278729 | 17.7824432 |
| Mprip | -1.1491304 | 5.1137E-10 | 2.8666E-08 | 1224.23393 | 551.910224 |
| Itga5 | -1.1459865 | 6.8416E-13 | 8.0532E-11 | 1773.41086 | 801.066709 |
| Lfng | -1.1451256 | 1.5717E-08 | 6.0388E-07 | 1401.46179 | 633.771887 |
| Agrn | -1.1443339 | 5.5118E-06 | 0.00010338 | 144.67856 | 65.3461216 |
| Fosl2 | -1.1442938 | 1.4157E-09 | 7.0658E-08 | 612.520293 | 277.091062 |
| Rab5b | -1.1442357 | 4.2969E-18 | 1.5012E-15 | 903.402963 | 408.605438 |
| Pi4ka | -1.143199 | 1.2833E-08 | 5.0602E-07 | 749.894866 | 339.462075 |
| Gtf2a1 | -1.1430069 | 2.4029E-10 | 1.4781E-08 | 356.283404 | 161.385106 |
| Met | -1.1422256 | 4.8852E-08 | 1.6604E-06 | 567.209422 | 256.952608 |
| Plekhg1 | -1.1418135 | 3.8223E-10 | 2.2257E-08 | 365.809012 | 165.655894 |
| Arid3a | -1.1415485 | 2.028E-08 | 7.6356E-07 | 849.726253 | 385.031464 |
| Mlec | -1.1409673 | 9.051E-15 | 1.5997E-12 | 2726.94772 | 1236.4027 |
| Zc3h4 | -1.1407617 | 3.6072E-09 | 1.6522E-07 | 271.971679 | 123.278329 |
| Hsd17b11 | -1.1405927 | 0.00032018 | 0.00301764 | 427.196614 | 193.634195 |
| Pym1 | -1.1384162 | 8.2088E-08 | 2.6351E-06 | 113.379237 | 51.5041184 |
| Dvl3 | -1.1379726 | 1.9998E-10 | 1.257E-08 | 433.678201 | 197.011204 |
| Ica1l | -1.137949 | 6.5167E-07 | 1.6155E-05 | 194.43808 | 88.1638325 |
| Trak2 | -1.1372732 | 1.5224E-06 | 3.3782E-05 | 454.769873 | 206.577502 |
| Scaf1 | -1.1364549 | 6.2076E-09 | 2.711E-07 | 433.966712 | 197.279879 |
| Nek9 | -1.1358503 | 9.8481E-11 | 6.8813E-09 | 839.7385 | 381.969643 |
| Kctd17 | -1.1340786 | 5.2665E-09 | 2.3477E-07 | 202.820891 | 92.3025818 |
| Snrnp200 | -1.13334 | 2.5746E-07 | 7.0401E-06 | 1323.1033 | 603.024474 |
| Sptan1 | -1.1332678 | 1.6191E-07 | 4.7975E-06 | 1630.27565 | 743.157695 |
| Gm43331 | -1.1331497 | 8.8375E-05 | 0.00106127 | 69.9739165 | 31.9013301 |
| Arhgef18 | -1.133125 | 3.0156E-07 | 8.1334E-06 | 1099.58181 | 501.243437 |
| Cramp1l | -1.1331168 | 4.3883E-06 | 8.4955E-05 | 104.479636 | 47.6070044 |
| Nhsl2 | -1.1328579 | 2.5011E-11 | 2.0645E-09 | 2232.8831 | 1017.95084 |
| Dcaf6 | -1.1310413 | 0.00125427 | 0.00913634 | 61.6906827 | 28.1772842 |
| Frs2 | -1.1306686 | 2.1517E-07 | 6.0762E-06 | 317.912743 | 145.208771 |
| Gm17106 | -1.1293264 | 0.01076442 | 0.04780192 | 28.8495571 | 13.143545 |
| Gm26699 | -1.1290636 | 0.00946699 | 0.04347928 | 28.5719831 | 13.1333997 |
| Prdm15 | -1.1284123 | 1.3242E-06 | 2.9647E-05 | 211.211729 | 96.5390277 |
| Cltc | -1.1276954 | 2.243E-07 | 6.2749E-06 | 3845.06879 | 1759.51952 |
| 5830417I10R | -1.1263819 | 4.9618E-06 | 9.4357E-05 | 122.633437 | 56.0639105 |
| Ptprs | -1.1260848 | 1.4048E-07 | 4.2207E-06 | 405.013198 | 185.645368 |
| 9430034N14I | -1.1253788 | 1.4047E-05 | 0.0002257 | 107.921396 | 49.4194394 |
| Zfp592 | -1.1248304 | 2.9567E-07 | 8.0116E-06 | 1114.05027 | 510.653584 |
| Gm42727 | -1.1245711 | 0.00019466 | 0.00201683 | 69.9669515 | 32.1108198 |
| Iqsec1 | -1.1243441 | 3.5154E-09 | 1.625E-07 | 177.895278 | 81.7026539 |
| Crebl2 | -1.1225812 | 8.48E-05 | 0.0010249 | 158.209641 | 72.6289571 |
| Zbtb26 | -1.1219373 | 8.9564E-05 | 0.00107213 | 81.896754 | 37.5662677 |
| Gm43628 | -1.1217346 | 0.01046405 | 0.04675832 | 25.4696233 | 11.7225291 |

| | | | | | |
|---|---|---|---|---|---|
| Gm44860 | -1.1214705 | 0.00065023 | 0.00537909 | 82.8780792 | 38.1679692 |
| Galc | -1.1200631 | 1.2603E-11 | 1.1203E-09 | 336.562921 | 154.684585 |
| Arrb1 | -1.1199111 | 1.5394E-14 | 2.5137E-12 | 1167.77251 | 537.242316 |
| Fam117a | -1.1175457 | 4.3194E-11 | 3.2773E-09 | 438.804789 | 202.31815 |
| Tbc1d2b | -1.1169922 | 2.6868E-05 | 0.00038924 | 1854.23612 | 854.856659 |
| Prrc2b | -1.1157864 | 1.3209E-06 | 2.9619E-05 | 539.438458 | 248.951427 |
| Naip6 | -1.1154147 | 6.2401E-08 | 2.0513E-06 | 347.017263 | 160.130928 |
| Sorl1 | -1.1153694 | 1.1932E-08 | 4.8192E-07 | 7974.23296 | 3680.59012 |
| Gm20404 | -1.1153586 | 0.00231601 | 0.01486507 | 73.8439217 | 34.17644 |
| Gm28033 | -1.1152407 | 0.00043245 | 0.00384422 | 66.9859988 | 30.7833015 |
| Cap1 | -1.1150297 | 7.5544E-11 | 5.4562E-09 | 1487.61765 | 686.629304 |
| Itpkb | -1.1146863 | 4.0526E-07 | 1.0588E-05 | 339.916685 | 157.003965 |
| Phf12 | -1.114256 | 3.4614E-08 | 1.215E-06 | 418.220617 | 193.275364 |
| Gm27232 | -1.1141861 | 0.00247795 | 0.01565416 | 41.1766732 | 19.0098166 |
| Thbd | -1.1138554 | 6.796E-07 | 1.6764E-05 | 1483.01878 | 685.093345 |
| Mief1 | -1.1121125 | 1.3248E-15 | 2.9705E-13 | 457.072013 | 211.283012 |
| Cgnl1 | -1.1117762 | 0.00199884 | 0.01321682 | 41.3466367 | 19.05435 |
| Bicra | -1.109352 | 4.0068E-05 | 0.00054474 | 236.597959 | 109.626987 |
| Fnbp4 | -1.1092615 | 2.0761E-07 | 5.8847E-06 | 1102.54616 | 511.130438 |
| Gm42867 | -1.1074656 | 0.00872931 | 0.04107853 | 35.0320844 | 16.3419683 |
| C230096K16F | -1.1073345 | 1.8942E-06 | 4.1063E-05 | 183.954856 | 85.4318644 |
| Pip4k2c | -1.1046684 | 9.1844E-12 | 8.3623E-10 | 397.140989 | 184.515673 |
| Nynrin | -1.1045669 | 0.01110793 | 0.04875091 | 60.457043 | 28.013212 |
| Abl1 | -1.1045231 | 6.2309E-10 | 3.4039E-08 | 943.122963 | 438.460421 |
| Sprtn | -1.1042961 | 3.691E-05 | 0.00050871 | 101.321813 | 47.1809314 |
| Vcl | -1.1039582 | 3.2542E-06 | 6.5271E-05 | 1703.28053 | 792.223008 |
| Zfp646 | -1.1036157 | 3.8685E-10 | 2.2439E-08 | 624.189908 | 290.268528 |
| Nat8l | -1.1034315 | 1.445E-08 | 5.6092E-07 | 621.905882 | 289.302696 |
| F5 | -1.102336 | 3.7005E-06 | 7.3245E-05 | 532.774671 | 247.970283 |
| Nup153 | -1.101278 | 4.0162E-12 | 4.0493E-10 | 672.190922 | 313.191313 |
| Myof | -1.1001017 | 2.5381E-05 | 0.00037164 | 503.348325 | 234.767417 |
| Nav1 | -1.0997881 | 1.8077E-07 | 5.2529E-06 | 910.725987 | 425.013512 |
| Pdpr | -1.0996468 | 1.0266E-06 | 2.3912E-05 | 245.733119 | 114.676488 |
| Zfyve26 | -1.0973837 | 8.7848E-08 | 2.7847E-06 | 319.274375 | 149.23054 |
| BC048403 | -1.0973364 | 7.1649E-05 | 0.00089127 | 65.3759513 | 30.6097604 |
| mt-Tp | -1.0972722 | 0.00277268 | 0.01710635 | 39.0528814 | 18.3257401 |
| Gm37010 | -1.0972703 | 0.00680244 | 0.03410414 | 48.9808708 | 23.0001494 |
| Plxnc1 | -1.0969447 | 6.342E-09 | 2.7537E-07 | 553.562373 | 258.670661 |
| Ube2z | -1.0968492 | 3.1345E-12 | 3.3162E-10 | 815.835282 | 381.187063 |
| Rnf126 | -1.0966289 | 2.4204E-10 | 1.4781E-08 | 399.981424 | 186.913013 |
| Cpsf7 | -1.0951479 | 1.1182E-05 | 0.00018665 | 329.441969 | 154.287894 |
| Wdfy3 | -1.0943844 | 9.3578E-11 | 6.6001E-09 | 1539.73805 | 721.000862 |
| Adpgk | -1.0919386 | 3.014E-08 | 1.0832E-06 | 1429.66242 | 670.746252 |
| Rab11fip4 | -1.0918035 | 0.00033531 | 0.00312493 | 81.6037582 | 38.2641845 |
| 4932442E05F | -1.0913598 | 0.00032062 | 0.0030199 | 67.9657873 | 31.8871235 |
| Smad7 | -1.0905237 | 1.6883E-05 | 0.00026066 | 80.6357251 | 37.765198 |

| | | | | | |
|---|---|---|---|---|---|
| Trim25 | -1.0904716 | 3.0918E-10 | 1.8359E-08 | 1601.96702 | 752.22804 |
| Kdm2b | -1.0890213 | 1.855E-09 | 9.1072E-08 | 416.757725 | 195.82695 |
| Sptbn1 | -1.0882331 | 7.1404E-10 | 3.8174E-08 | 962.318408 | 452.53926 |
| Jarid2 | -1.0878297 | 3.9725E-08 | 1.3764E-06 | 2364.91623 | 1112.40569 |
| Gm43361 | -1.0877963 | 4.8179E-08 | 1.6413E-06 | 246.627132 | 116.043848 |
| Zfp26 | -1.0877933 | 4.4433E-07 | 1.143E-05 | 223.727339 | 105.21009 |
| Nectin1 | -1.0873174 | 0.00217994 | 0.01415265 | 137.72816 | 64.8237332 |
| Dpysl2 | -1.086599 | 4.3066E-15 | 8.0872E-13 | 796.315849 | 374.961526 |
| Thra | -1.0864152 | 8.8038E-07 | 2.0927E-05 | 138.469215 | 65.2080766 |
| Map4k4 | -1.0861577 | 5.6872E-10 | 3.1182E-08 | 826.382968 | 389.218049 |
| Midn | -1.0856475 | 0.00520234 | 0.02781307 | 64.2976959 | 30.1721472 |
| Atp10d | -1.0838975 | 6.248E-13 | 7.4494E-11 | 692.322167 | 326.579397 |
| Usp9x | -1.0822495 | 5.5495E-08 | 1.8404E-06 | 388.467853 | 183.339518 |
| Gm26826 | -1.0817108 | 0.00251897 | 0.01585358 | 47.0784881 | 22.2132895 |
| Lrrc8a | -1.0809396 | 0.00857514 | 0.04043452 | 44.0218499 | 20.8624227 |
| Bri3 | -1.0805723 | 3.0407E-06 | 6.1735E-05 | 119.017343 | 56.1751667 |
| Slc13a3 | -1.07792 | 0.003157 | 0.0189181 | 31.6233937 | 15.0191911 |
| Zdhhc23 | -1.0763522 | 8.9747E-07 | 2.1183E-05 | 237.8378 | 112.762262 |
| Fndc3b | -1.0756016 | 1.2301E-08 | 4.889E-07 | 544.750092 | 258.319562 |
| Maml1 | -1.07506 | 0.00012702 | 0.00142775 | 151.417105 | 71.8525059 |
| A430088P11I | -1.074154 | 0.00084398 | 0.00661751 | 54.6266279 | 26.0296278 |
| Gm43460 | -1.0734748 | 0.00626187 | 0.0320518 | 31.6987409 | 15.1305776 |
| Fnip1 | -1.0734088 | 5.2906E-06 | 0.0001001 | 253.059198 | 120.268367 |
| Mex3d | -1.0727211 | 4.448E-05 | 0.00059345 | 149.25475 | 70.8566667 |
| Rnf111 | -1.0724013 | 6.407E-10 | 3.4748E-08 | 834.756844 | 396.940561 |
| Aff1 | -1.0707877 | 3.9349E-07 | 1.0334E-05 | 431.646155 | 205.319578 |
| Atxn7l3 | -1.0696305 | 8.8101E-10 | 4.644E-08 | 326.687588 | 155.731678 |
| Gm26749 | -1.0695829 | 0.00158767 | 0.01105776 | 50.50526 | 24.0859396 |
| Nup205 | -1.068751 | 0.00026154 | 0.00257139 | 285.174443 | 135.834783 |
| Sos2 | -1.0685699 | 2.8962E-13 | 3.7204E-11 | 707.757964 | 337.430217 |
| Lrp5 | -1.0684561 | 0.00050468 | 0.00436241 | 66.6285248 | 31.8597116 |
| Zbtb37 | -1.0670105 | 2.524E-05 | 0.00036993 | 97.9318239 | 46.7586902 |
| Adamts6 | -1.0644885 | 0.01097949 | 0.04842114 | 43.135597 | 20.7095062 |
| Rap1gap2 | -1.0628355 | 1.0957E-06 | 2.5286E-05 | 1281.83297 | 613.473865 |
| Madd | -1.062561 | 0.00010892 | 0.00126719 | 1882.32889 | 901.175527 |
| Gnb2 | -1.0621053 | 3.0567E-15 | 5.8872E-13 | 4611.46795 | 2208.39749 |
| Bmf | -1.0610381 | 0.00011789 | 0.00134476 | 264.444329 | 126.84457 |
| Camsap2 | -1.058492 | 4.422E-07 | 1.1414E-05 | 253.082237 | 121.398376 |
| Rmnd5a | -1.0582217 | 4.9717E-09 | 2.2443E-07 | 625.308161 | 300.186343 |
| Slx4 | -1.0578846 | 0.00216272 | 0.01407842 | 145.938272 | 70.056297 |
| Ankmy1 | -1.0576726 | 0.00077338 | 0.00614733 | 71.5531444 | 34.3011525 |
| Polr2a | -1.0573375 | 1.7391E-10 | 1.1024E-08 | 880.343616 | 422.861825 |
| Mbd6 | -1.0573225 | 0.00057036 | 0.0048355 | 116.29626 | 55.8960574 |
| Nbas | -1.0570149 | 1.537E-06 | 3.3957E-05 | 174.812662 | 83.8810718 |
| Gm13556 | -1.056144 | 0.00394025 | 0.02234595 | 39.1261897 | 18.9155046 |
| Prdm16 | -1.0557552 | 0.00408454 | 0.02294766 | 73.3517259 | 35.3160122 |

| | | | | | |
|---|---|---|---|---|---|
| Nod2 | -1.0557353 | 6.4145E-05 | 0.00080843 | 74.7885199 | 35.8945587 |
| Atf7 | -1.0556146 | 1.2933E-07 | 3.9093E-06 | 390.357843 | 187.682433 |
| Ube2q1 | -1.0547815 | 5.4435E-17 | 1.5726E-14 | 1324.92105 | 637.806824 |
| Ripor1 | -1.0544709 | 2.1638E-06 | 4.6174E-05 | 307.246834 | 147.842091 |
| Fry | -1.0543973 | 3.093E-08 | 1.109E-06 | 518.37033 | 249.659527 |
| Foxred2 | -1.0541482 | 1.2573E-05 | 0.00020554 | 617.667947 | 297.346475 |
| Pom121 | -1.0530976 | 5.0143E-07 | 1.2746E-05 | 611.638342 | 294.592814 |
| Kdm4a | -1.0521524 | 1.1123E-05 | 0.00018608 | 280.132008 | 134.919717 |
| Pgap1 | -1.0517768 | 5.2109E-08 | 1.7592E-06 | 326.73987 | 157.532738 |
| Zcchc2 | -1.0515925 | 1.296E-09 | 6.5552E-08 | 571.436091 | 275.548871 |
| Suz12 | -1.0510915 | 1.0544E-08 | 4.34E-07 | 389.685314 | 188.002421 |
| Hcfc1 | -1.0503509 | 2.1942E-10 | 1.3621E-08 | 1007.17326 | 486.10469 |
| Gm44154 | -1.0499369 | 0.00223807 | 0.01444267 | 76.6414303 | 36.916887 |
| Rab11fip1 | -1.0498185 | 1.8886E-13 | 2.556E-11 | 651.827547 | 314.737049 |
| Myo18a | -1.0493505 | 2.9447E-05 | 0.00041853 | 1206.05338 | 582.709191 |
| Atg9b | -1.0491133 | 0.01096209 | 0.04837942 | 27.3983613 | 13.208591 |
| Ep300 | -1.0486906 | 1.9684E-14 | 3.1796E-12 | 1408.25926 | 680.701689 |
| Gm10602 | -1.0477679 | 0.00033638 | 0.00313104 | 65.374965 | 31.6123221 |
| E2f2 | -1.0468344 | 3.2536E-11 | 2.6138E-09 | 1981.14077 | 958.765285 |
| Dele1 | -1.0463249 | 1.1667E-05 | 0.00019367 | 105.578298 | 50.9780499 |
| Trappc10 | -1.0458637 | 3.9123E-07 | 1.0293E-05 | 413.957363 | 200.517879 |
| Tshz1 | -1.0453619 | 3.8169E-05 | 0.00052319 | 317.390273 | 153.630322 |
| Tet3 | -1.0445164 | 1.6658E-07 | 4.9166E-06 | 793.114984 | 384.546623 |
| Picalm | -1.0428385 | 1.5187E-05 | 0.00023991 | 1247.39038 | 605.44275 |
| Megf9 | -1.0413899 | 8.3172E-08 | 2.6642E-06 | 554.729736 | 269.33256 |
| Ddb1 | -1.0411585 | 6.842E-11 | 4.9656E-09 | 2366.74637 | 1149.79638 |
| Asna1 | -1.0407237 | 6.2771E-14 | 9.1554E-12 | 569.730591 | 276.73743 |
| Arid1b | -1.0398393 | 4.1655E-07 | 1.0864E-05 | 852.914328 | 414.732193 |
| Rc3h2 | -1.03971 | 2.5774E-07 | 7.0401E-06 | 412.244072 | 200.680288 |
| Nbeal1 | -1.0388834 | 1.6712E-05 | 0.0002591 | 214.296017 | 104.41771 |
| 5830462I19R | -1.0379671 | 0.00065981 | 0.00543902 | 147.417937 | 71.7210489 |
| Plcxd2 | -1.0377581 | 0.000504 | 0.00435898 | 67.3558511 | 32.9258847 |
| Kdm7a | -1.0375178 | 1.3552E-13 | 1.9027E-11 | 2649.2773 | 1290.39512 |
| Nle1 | -1.0370555 | 2.5021E-05 | 0.00036708 | 73.4969711 | 35.8958278 |
| Asxl1 | -1.036489 | 2.3347E-07 | 6.4713E-06 | 647.216618 | 315.293171 |
| Lemd3 | -1.0354966 | 9.424E-06 | 0.00016125 | 325.925207 | 158.887381 |
| Gm26778 | -1.0354458 | 6.8013E-08 | 2.2261E-06 | 648.072139 | 316.184868 |
| Borcs6 | -1.0352613 | 4.1738E-10 | 2.4117E-08 | 461.182928 | 224.944617 |
| Dnajc13 | -1.0344007 | 1.2977E-10 | 8.5884E-09 | 653.472807 | 319.091072 |
| Spef1 | -1.0343577 | 0.00255092 | 0.01602777 | 52.8248556 | 25.7970245 |
| Extl3 | -1.0342645 | 4.5765E-06 | 8.8145E-05 | 485.426475 | 236.932811 |
| C130089K02F | -1.0331812 | 0.00161774 | 0.01121 | 198.888279 | 97.3490468 |
| AC154492.2 | -1.0326804 | 6.9676E-05 | 0.00087228 | 101.632291 | 49.8052506 |
| Nphp3 | -1.031959 | 0.00068499 | 0.00560792 | 54.1337747 | 26.3877816 |
| Mtmr3 | -1.0306173 | 4.6118E-07 | 1.1803E-05 | 2461.70009 | 1204.82737 |
| Hs6st1 | -1.0304007 | 1.067E-07 | 3.2983E-06 | 798.522386 | 390.761886 |

| | | | | | |
|---|---|---|---|---|---|
| Wiz | -1.0298277 | 1.1961E-06 | 2.7266E-05 | 302.423627 | 148.051843 |
| Gm42547 | -1.0298086 | 0.00130467 | 0.00941404 | 77.0509437 | 37.6514536 |
| Gsk3a | -1.0295818 | 1.3318E-14 | 2.2736E-12 | 965.792055 | 472.902805 |
| Gabpb2 | -1.0286273 | 1.4739E-08 | 5.7069E-07 | 403.163657 | 197.707591 |
| Tbrg4 | -1.0271404 | 0.00011221 | 0.00129676 | 113.068333 | 55.4992503 |
| Agap1 | -1.0265107 | 9.9042E-10 | 5.1307E-08 | 221.745966 | 108.854661 |
| Arid2 | -1.0262068 | 1.8514E-06 | 4.0193E-05 | 589.910353 | 289.516881 |
| Psme4 | -1.0257655 | 1.8832E-07 | 5.4594E-06 | 524.183033 | 257.429261 |
| Sos1 | -1.0252301 | 1.1719E-06 | 2.6837E-05 | 218.205379 | 107.10954 |
| Ldlr | -1.0240403 | 2.7365E-05 | 0.00039481 | 2992.10552 | 1471.20363 |
| Fam196b | -1.0237811 | 0.0044717 | 0.02466165 | 76.0003677 | 37.4236994 |
| Nin | -1.0233921 | 5.4238E-09 | 2.4036E-07 | 1001.97326 | 492.80163 |
| Ranbp2 | -1.0228496 | 4.6853E-10 | 2.6662E-08 | 588.900257 | 289.926071 |
| Hist1h4d | -1.0197311 | 5.3707E-05 | 0.00069916 | 283.560011 | 139.81967 |
| Fam160b1 | -1.0189721 | 1.5293E-10 | 9.9889E-09 | 630.990441 | 311.287536 |
| Leng8 | -1.0165499 | 4.7704E-06 | 9.1179E-05 | 1197.46637 | 591.910365 |
| Tmem143 | -1.0162925 | 0.00331653 | 0.01957731 | 55.3404305 | 27.3958038 |
| Lyst | -1.0160724 | 1.153E-09 | 5.8917E-08 | 812.342521 | 401.731844 |
| Zfand3 | -1.0155415 | 1.2146E-05 | 0.00019985 | 260.548271 | 128.78624 |
| Zmiz1 | -1.0150906 | 3.5934E-09 | 1.6509E-07 | 2205.39852 | 1091.17643 |
| Lrp1 | -1.0150007 | 2.1972E-08 | 8.2112E-07 | 10298.2221 | 5095.80595 |
| Apaf1 | -1.0139539 | 7.3713E-08 | 2.397E-06 | 1190.91947 | 589.550632 |
| Gm15859 | -1.0122172 | 0.01101825 | 0.04852748 | 36.6607815 | 18.1810264 |
| Hnrnpl | -1.0119034 | 2.4423E-17 | 7.3672E-15 | 1620.76023 | 803.454351 |
| Gm7694 | -1.0108745 | 0.00011564 | 0.00132379 | 161.824523 | 80.23562 |
| Nrp2 | -1.0104613 | 8.4959E-05 | 0.001026 | 146.899248 | 72.895162 |
| Rps6ka2 | -1.0102407 | 0.00126327 | 0.00918595 | 173.018488 | 85.7598487 |
| 9530052E02F | -1.010181 | 4.0168E-07 | 1.0513E-05 | 171.569054 | 85.1085305 |
| Tnfrsf14 | -1.0100925 | 0.0003055 | 0.00292641 | 115.65915 | 57.4835527 |
| Klhl18 | -1.0092565 | 4.2403E-12 | 4.2187E-10 | 467.524141 | 232.320599 |
| Cnnm4 | -1.0087003 | 2.2189E-07 | 6.2191E-06 | 296.860733 | 147.605565 |
| Sept9 | -1.005765 | 3.7798E-11 | 2.9422E-09 | 4808.5102 | 2394.45106 |
| Numa1 | -1.0046261 | 4.1637E-05 | 0.00056201 | 923.786408 | 460.343817 |
| Gm42893 | -1.0042985 | 0.00690496 | 0.03450872 | 46.6233718 | 23.2709483 |
| Gm26787 | -1.0042971 | 5.7383E-05 | 0.00074124 | 113.209947 | 56.401589 |
| Ppp2cb | -1.0041908 | 1.1794E-10 | 7.9539E-09 | 1583.94763 | 789.555901 |
| Slco4c1 | -1.0041548 | 0.00497089 | 0.02681422 | 88.3833976 | 44.1212114 |
| Gm12264 | -1.0040529 | 2.8777E-06 | 5.8979E-05 | 124.875743 | 62.1422728 |
| Faddos | -1.0033455 | 0.00596095 | 0.03085851 | 34.4498405 | 17.1319823 |
| Arl4c | -1.003201 | 6.5907E-06 | 0.00011987 | 361.449488 | 180.394099 |
| Pi16 | -1.0031076 | 9.7512E-08 | 3.0267E-06 | 2044.32794 | 1019.88021 |
| Wdtc1 | -1.0018578 | 0.00016462 | 0.00174784 | 68.4532619 | 34.1063945 |
| Zfp770 | -1.0005304 | 8.7558E-05 | 0.00105463 | 102.644368 | 51.28929 |
| Tbcel | -1.0003483 | 1.1612E-08 | 4.7149E-07 | 276.841206 | 138.429874 |
| Gm26780 | -0.999525 | 0.0004872 | 0.00424304 | 87.132196 | 43.7530588 |
| Atxn2l | -0.9975559 | 7.2743E-10 | 3.8752E-08 | 951.73414 | 476.599686 |

| | | | | | |
|---|---|---|---|---|---|
| Htt | -0.9973258 | 4.9054E-06 | 9.3401E-05 | 178.655005 | 89.7159711 |
| Sowahc | -0.9967532 | 9.838E-11 | 6.8813E-09 | 1415.49268 | 709.145974 |
| Stard9 | -0.9966106 | 1.1424E-05 | 0.00019006 | 665.914591 | 333.884483 |
| Nfya | -0.9959584 | 3.8383E-05 | 0.00052564 | 103.782835 | 51.9981599 |
| Slf2 | -0.9934759 | 6.3169E-07 | 1.5738E-05 | 641.289696 | 321.950859 |
| Gmeb2 | -0.9932457 | 5.2929E-05 | 0.00069204 | 293.558557 | 147.254287 |
| Serinc5 | -0.993012 | 3.612E-08 | 1.2649E-06 | 689.967515 | 346.408251 |
| Ube4bos3 | -0.9920448 | 0.00983907 | 0.04479161 | 42.4900411 | 21.2440242 |
| Hectd4 | -0.9901872 | 6.7123E-06 | 0.00012164 | 189.704157 | 95.521634 |
| Nlrc5 | -0.9883535 | 3.7905E-05 | 0.00052004 | 743.541724 | 374.843207 |
| Sec14l1 | -0.9866176 | 2.5302E-08 | 9.2486E-07 | 2226.32957 | 1123.39842 |
| Ube4b | -0.985765 | 3.0068E-07 | 8.1243E-06 | 681.002699 | 343.760836 |
| Tial1 | -0.984735 | 2.8612E-05 | 0.00040937 | 288.053511 | 145.686756 |
| Plekhm1 | -0.9841694 | 3.7914E-07 | 1.001E-05 | 1098.81199 | 555.289935 |
| Ipcef1 | -0.9838164 | 6.5195E-05 | 0.00082029 | 263.456154 | 133.271705 |
| Gm17477 | -0.9837981 | 0.00916253 | 0.04244485 | 43.0152544 | 21.789459 |
| Gm20300 | -0.9832476 | 0.00021723 | 0.00220955 | 83.3281665 | 42.2918792 |
| Gm43323 | -0.9830751 | 0.00500813 | 0.02696955 | 54.0907835 | 27.4608573 |
| Nfatc2 | -0.9823143 | 0.00022998 | 0.00231878 | 414.844702 | 209.947469 |
| Pkd1 | -0.982133 | 3.4769E-06 | 6.9275E-05 | 1026.39684 | 519.39152 |
| Zfp629 | -0.9815921 | 0.00946701 | 0.04347928 | 115.519328 | 58.5175448 |
| Gm26852 | -0.9793934 | 0.00584247 | 0.03040229 | 33.525863 | 17.0330977 |
| Insr | -0.9792615 | 1.0094E-05 | 0.00017095 | 704.781722 | 357.515885 |
| Kansl1 | -0.979113 | 5.9407E-11 | 4.3964E-09 | 1141.68972 | 579.116498 |
| Gm26517 | -0.9790475 | 0.0034018 | 0.01996298 | 54.7407574 | 27.7841116 |
| Adamts10 | -0.9788836 | 7.4793E-05 | 0.000921 | 873.745878 | 443.286713 |
| E330009J07R | -0.9788656 | 5.365E-05 | 0.00069903 | 126.418376 | 64.0235033 |
| Tmed8 | -0.9788103 | 0.00137038 | 0.00979414 | 121.154304 | 61.3117249 |
| Mir99ahg | -0.9779224 | 0.01102537 | 0.04854457 | 33.8897346 | 17.2281749 |
| Prkce | -0.9774314 | 6.2048E-05 | 0.00079062 | 121.524889 | 61.7717121 |
| Marf1 | -0.9748308 | 4.795E-06 | 9.1415E-05 | 170.078069 | 86.6330759 |
| Upf1 | -0.9738623 | 4.4833E-07 | 1.1494E-05 | 1401.58477 | 713.430513 |
| Sipa1l2 | -0.9735496 | 0.00119358 | 0.00876403 | 248.624181 | 126.59535 |
| Paip2b | -0.9716362 | 0.00075623 | 0.00605262 | 55.4544526 | 28.3167253 |
| Dip2b | -0.9710145 | 5.0546E-09 | 2.2667E-07 | 477.497308 | 243.590385 |
| Gm26538 | -0.9708391 | 0.00022467 | 0.00227135 | 81.1484291 | 41.45529 |
| Per2 | -0.9698682 | 0.00128952 | 0.00933613 | 84.9482992 | 43.3197065 |
| Nsd2 | -0.9694382 | 1.21E-07 | 3.6871E-06 | 335.732943 | 171.43873 |
| Gm16599 | -0.9690672 | 0.0002012 | 0.0020746 | 76.6181875 | 39.1833372 |
| Ppp3cb | -0.9674866 | 3.7765E-06 | 7.4358E-05 | 267.692888 | 137.006758 |
| Ubr4 | -0.9670496 | 1.8501E-05 | 0.00028332 | 988.541947 | 505.71304 |
| Acaca | -0.9667851 | 0.00011103 | 0.00128606 | 116.447545 | 59.6570402 |
| Ppm1a | -0.9661887 | 1.0472E-14 | 1.8293E-12 | 697.007568 | 356.667175 |
| Sik2 | -0.9659527 | 1.9954E-08 | 7.5319E-07 | 557.148484 | 285.395838 |
| Zfp568 | -0.9658561 | 0.00035257 | 0.00325604 | 119.559595 | 61.2763406 |
| Kif16b | -0.9652238 | 0.0003129 | 0.00297136 | 267.271608 | 136.744798 |

| | | | | | |
|---|---|---|---|---|---|
| Isg20l2 | -0.9650841 | 6.6125E-09 | 2.8546E-07 | 451.940726 | 231.372493 |
| Bcorl1 | -0.9647843 | 0.00270604 | 0.01679868 | 43.9694361 | 22.513319 |
| Pdzd8 | -0.9642671 | 2.1367E-12 | 2.326E-10 | 600.828434 | 307.826611 |
| Gm16726 | -0.964117 | 0.00030714 | 0.00293333 | 84.5475603 | 43.3466113 |
| Hmbox1 | -0.9632795 | 1.1941E-07 | 3.6462E-06 | 226.64008 | 116.406686 |
| Zfp609 | -0.9627075 | 0.00054063 | 0.0046173 | 82.887484 | 42.4633624 |
| Smg5 | -0.9617966 | 2.2534E-08 | 8.3796E-07 | 614.3932 | 315.641734 |
| Itprid2 | -0.9615562 | 5.717E-09 | 2.5261E-07 | 719.61104 | 369.290959 |
| Gcn1l1 | -0.9613279 | 8.4392E-06 | 0.00014742 | 496.96353 | 255.361581 |
| Hip1 | -0.9609616 | 1.3645E-05 | 0.00022018 | 1650.56784 | 847.851911 |
| Arel1 | -0.960412 | 5.5873E-06 | 0.00010427 | 813.029873 | 417.597868 |
| Abcb7 | -0.9587599 | 2.1887E-05 | 0.00032751 | 460.937195 | 237.012289 |
| Irs2 | -0.9586797 | 1.4966E-06 | 3.3308E-05 | 867.40066 | 446.166013 |
| Pik3c2a | -0.9582738 | 1.252E-07 | 3.7996E-06 | 364.624109 | 187.651545 |
| Arhgap26 | -0.9568156 | 7.4137E-14 | 1.0709E-11 | 2680.6224 | 1380.93344 |
| Mtmr4 | -0.9561948 | 1.29E-06 | 2.9055E-05 | 927.691819 | 478.018869 |
| Slc22a15 | -0.9560967 | 4.1684E-06 | 8.1222E-05 | 371.202469 | 191.16307 |
| Ssh1 | -0.9541275 | 2.531E-06 | 5.2663E-05 | 607.54832 | 313.48747 |
| Kremen1 | -0.9536789 | 0.00026546 | 0.0026058 | 456.746631 | 235.595133 |
| Fus | -0.9534051 | 1.5989E-05 | 0.00025021 | 486.625786 | 251.432837 |
| Zfp619 | -0.9531821 | 0.00014688 | 0.00159465 | 76.8528767 | 39.6784322 |
| Plcg1 | -0.953052 | 0.00179955 | 0.01221625 | 157.78503 | 81.4783801 |
| Xpo7 | -0.9530159 | 2.4461E-06 | 5.1039E-05 | 846.763438 | 437.217847 |
| Tmem86b | -0.952227 | 0.0003159 | 0.00299417 | 87.1971484 | 45.1889302 |
| Igf1r | -0.9518524 | 1.5707E-05 | 0.00024605 | 129.346659 | 66.8595077 |
| Fam84b | -0.9517448 | 2.1307E-05 | 0.00031977 | 339.187651 | 175.169715 |
| Fadd | -0.9506218 | 6.2248E-05 | 0.00079181 | 107.020955 | 55.2504657 |
| Vwa8 | -0.9505489 | 0.00027166 | 0.00265522 | 154.695571 | 80.0179051 |
| Trim65 | -0.9497037 | 2.4596E-07 | 6.7676E-06 | 504.114245 | 260.944898 |
| Tnks2 | -0.9494513 | 8.532E-08 | 2.7272E-06 | 1692.70325 | 876.354478 |
| Atp2a3 | -0.9490713 | 2.9806E-05 | 0.00042255 | 4529.35537 | 2345.86026 |
| Gm42729 | -0.948579 | 0.00393037 | 0.02232106 | 50.9489338 | 26.5113359 |
| Mst1r | -0.9473984 | 0.01148263 | 0.04997204 | 195.038687 | 101.097951 |
| Ulbp1 | -0.9473383 | 0.00447642 | 0.02466957 | 48.1078392 | 25.0140294 |
| Stim1 | -0.9450758 | 7.7867E-12 | 7.2659E-10 | 810.737412 | 420.961644 |
| Nxn | -0.9443192 | 0.00387645 | 0.02209255 | 46.844117 | 24.4036212 |
| A230103J11R | -0.944232 | 0.00090469 | 0.00702652 | 65.9494487 | 34.2495316 |
| Prr33 | -0.9430485 | 0.00486201 | 0.02633095 | 62.8531826 | 32.7350418 |
| Dcaf7 | -0.9418482 | 8.9701E-08 | 2.8251E-06 | 2605.45221 | 1356.09183 |
| Pdp2 | -0.9411766 | 6.3902E-05 | 0.00080672 | 167.978693 | 87.5205272 |
| Wdr26 | -0.9404996 | 1.4299E-07 | 4.2793E-06 | 1783.90466 | 929.346264 |
| Fam151a | -0.9404568 | 1.5231E-05 | 0.00024035 | 139.652393 | 72.6741085 |
| Ahr | -0.9382532 | 1.9023E-09 | 9.2786E-08 | 1940.16325 | 1012.34066 |
| Rere | -0.9353263 | 8.9609E-07 | 2.1183E-05 | 822.939053 | 430.279756 |
| 2010008C14F | -0.9351597 | 0.00018252 | 0.00190152 | 89.4047397 | 46.6868761 |
| Klhl2 | -0.9350328 | 5.2942E-10 | 2.9457E-08 | 793.590038 | 414.875812 |

| | | | | | |
|---|---|---|---|---|---|
| Acads | -0.9348593 | 2.7248E-09 | 1.2872E-07 | 365.109072 | 190.878977 |
| Kdm2a | -0.9347705 | 9.9762E-11 | 6.9385E-09 | 1053.3812 | 551.184532 |
| Plec | -0.9342783 | 1.4014E-09 | 7.0414E-08 | 2549.40057 | 1333.92312 |
| Rab27a | -0.9338464 | 1.5634E-05 | 0.00024516 | 769.84425 | 402.782424 |
| Pgam2 | -0.9333904 | 0.00092737 | 0.00715553 | 76.5933835 | 40.0223911 |
| Zzef1 | -0.9332007 | 2.5258E-07 | 6.9369E-06 | 622.436594 | 326.006756 |
| Avl9 | -0.9329593 | 6.8856E-08 | 2.2488E-06 | 468.19172 | 245.087343 |
| Mrc1 | -0.9324854 | 1.2031E-05 | 0.00019884 | 178.617291 | 93.4449923 |
| Uggt1 | -0.9323904 | 1.7072E-09 | 8.409E-08 | 2988.90175 | 1565.91201 |
| Atg4c | -0.93227 | 2.8564E-10 | 1.7028E-08 | 494.024937 | 258.829998 |
| Tmcc2 | -0.9322436 | 0.00068921 | 0.00563332 | 212.9138 | 111.499508 |
| Dennd5a | -0.9314727 | 8.7771E-08 | 2.7847E-06 | 2177.90921 | 1141.80227 |
| 9130221H12I | -0.930457 | 3.7993E-10 | 2.2209E-08 | 419.883808 | 220.163003 |
| Zfhx3 | -0.9290468 | 0.00033156 | 0.00310147 | 301.212892 | 158.186559 |
| Spag9 | -0.9286437 | 5.4244E-10 | 2.996E-08 | 792.31678 | 416.4228 |
| Ranbp10 | -0.9286212 | 3.2122E-08 | 1.1381E-06 | 621.314936 | 326.415221 |
| Pura | -0.9275258 | 3.0352E-09 | 1.4205E-07 | 383.16947 | 201.443005 |
| 0610009E02F | -0.9272874 | 0.00547931 | 0.02892327 | 49.1322064 | 25.8832565 |
| Scap | -0.9264562 | 0.0001272 | 0.00142775 | 596.068858 | 313.499851 |
| Snx33 | -0.926263 | 0.00286575 | 0.01759384 | 112.570493 | 59.2807049 |
| Depdc5 | -0.9262004 | 2.2575E-05 | 0.00033512 | 303.744502 | 159.834447 |
| Rfx7 | -0.9261937 | 2.3018E-07 | 6.4038E-06 | 297.296877 | 156.494838 |
| Phlpp1 | -0.925461 | 0.00063048 | 0.00524456 | 147.487986 | 77.5506832 |
| Svil | -0.9248679 | 7.53E-08 | 2.4433E-06 | 2021.13795 | 1064.48259 |
| Ubqln4 | -0.9246259 | 2.3671E-07 | 6.5489E-06 | 479.13539 | 252.319724 |
| Prmt1 | -0.9245468 | 3.2968E-13 | 4.1972E-11 | 634.433342 | 334.409875 |
| Nf1 | -0.9229326 | 0.00318057 | 0.01897823 | 104.574267 | 55.1273181 |
| Arih1 | -0.9226893 | 5.2453E-08 | 1.7668E-06 | 972.532518 | 512.87165 |
| Setd5 | -0.9220755 | 2.7474E-05 | 0.00039573 | 427.897406 | 226.001975 |
| Ppp2r5e | -0.9215501 | 0.0040204 | 0.02268058 | 455.872812 | 240.403303 |
| Ambra1 | -0.9214525 | 2.6778E-05 | 0.0003883 | 579.650663 | 306.006485 |
| Gm20659 | -0.9210604 | 2.5005E-05 | 0.00036708 | 304.843351 | 161.003948 |
| Elmsan1 | -0.920866 | 3.8419E-06 | 7.5446E-05 | 588.261493 | 310.57624 |
| Yeats2 | -0.9207785 | 0.00074857 | 0.00599772 | 265.759072 | 140.402293 |
| Rreb1 | -0.9207229 | 1.4756E-05 | 0.00023482 | 343.652433 | 181.596379 |
| Sdc3 | -0.9199349 | 0.00024461 | 0.00243524 | 3804.2039 | 2010.4586 |
| Prdm2 | -0.9197611 | 2.379E-08 | 8.7411E-07 | 321.810919 | 170.068032 |
| Xpr1 | -0.9177418 | 3.8543E-07 | 1.0158E-05 | 1033.89922 | 547.282088 |
| Hif1an | -0.9174932 | 7.6485E-09 | 3.2458E-07 | 565.720671 | 299.466768 |
| Tet2 | -0.9169374 | 1.4506E-07 | 4.3324E-06 | 805.003543 | 426.499906 |
| Ikzf1 | -0.9169233 | 5.4825E-08 | 1.8263E-06 | 2109.67494 | 1117.38934 |
| Chd8 | -0.9168775 | 5.9973E-05 | 0.00076809 | 291.672168 | 154.680784 |
| Lmo4 | -0.9159792 | 7.4777E-06 | 0.00013279 | 1263.5257 | 669.598067 |
| Cnnm3 | -0.9156115 | 1.2645E-09 | 6.4177E-08 | 730.926125 | 387.289521 |
| Slc27a4 | -0.9154598 | 4.4991E-10 | 2.5798E-08 | 482.390051 | 255.531594 |
| Rbm33 | -0.9152537 | 5.24E-09 | 2.3429E-07 | 429.934031 | 227.907498 |

| | | | | | |
|---|---|---|---|---|---|
| Ppard | -0.9144208 | 0.00076522 | 0.00610833 | 72.4668206 | 38.4874196 |
| Slc39a1 | -0.9139739 | 0.00150592 | 0.0105915 | 71.4758162 | 37.9223945 |
| Rnpepl1 | -0.9136509 | 5.4952E-08 | 1.8264E-06 | 1964.8346 | 1042.79115 |
| Etnk1 | -0.9131613 | 1.1974E-10 | 8.0309E-09 | 1019.39457 | 541.187264 |
| Micall1 | -0.9128819 | 3.9259E-05 | 0.00053569 | 234.186095 | 124.24657 |
| Rapgef1 | -0.9128818 | 1.0093E-07 | 3.1262E-06 | 753.117712 | 400.078927 |
| Virma | -0.910926 | 2.9385E-06 | 6.0061E-05 | 631.548624 | 335.718555 |
| Smarcc2 | -0.9093192 | 9.6925E-05 | 0.00114969 | 1743.72038 | 928.32018 |
| E2f3 | -0.9086288 | 0.00813986 | 0.0389319 | 55.2831612 | 29.4610121 |
| C1galt1 | -0.908553 | 1.1049E-07 | 3.4013E-06 | 573.980348 | 305.640043 |
| Pbx1 | -0.9079667 | 0.00111491 | 0.00833496 | 164.322287 | 87.6963521 |
| Timmdc1 | -0.906789 | 0.00027896 | 0.00271077 | 125.709583 | 66.9554539 |
| Kctd12 | -0.906731 | 8.5106E-06 | 0.00014798 | 1117.07388 | 595.846438 |
| Ralgapb | -0.9062315 | 3.739E-05 | 0.00051392 | 441.373398 | 235.602793 |
| Cep170 | -0.9060902 | 1.0466E-05 | 0.00017626 | 874.946039 | 466.601956 |
| Lrrk2 | -0.9053601 | 6.2029E-06 | 0.00011392 | 623.787614 | 333.086119 |
| Hnrnpul1 | -0.9046708 | 4.9493E-14 | 7.4354E-12 | 1324.28727 | 707.146892 |
| Ptprj | -0.9041613 | 6.0275E-07 | 1.5042E-05 | 902.403682 | 482.048128 |
| 2310022B05F | -0.9034905 | 0.00045974 | 0.0040508 | 197.933563 | 105.691833 |
| Gm9776 | -0.9026604 | 0.01021302 | 0.04601984 | 30.348868 | 16.2112647 |
| Map7d1 | -0.9023657 | 2.1608E-09 | 1.0404E-07 | 2759.21677 | 1476.02627 |
| Apc | -0.9020373 | 1.3785E-07 | 4.1503E-06 | 424.812319 | 227.360499 |
| Arap1 | -0.9019286 | 5.1531E-06 | 9.787E-05 | 5556.4785 | 2973.45038 |
| Cers6 | -0.8997275 | 5.5106E-12 | 5.4108E-10 | 2893.52676 | 1551.06838 |
| H6pd | -0.8996975 | 0.00016826 | 0.00178134 | 1093.01801 | 585.603601 |
| Eif4g1 | -0.8995328 | 1.2142E-08 | 4.8515E-07 | 2244.79387 | 1203.12172 |
| Rtel1 | -0.8989704 | 0.00234521 | 0.01501153 | 170.542813 | 91.3405351 |
| Pacs2 | -0.8989315 | 3.0551E-09 | 1.4254E-07 | 1641.04467 | 879.886439 |
| Ppp1r15b | -0.8987349 | 5.2883E-08 | 1.7733E-06 | 2942.02449 | 1577.68827 |
| F420014N23I | -0.8984501 | 0.00580652 | 0.03028865 | 55.0722297 | 29.4513098 |
| Esrp2 | -0.8982078 | 0.00236506 | 0.01510644 | 53.458962 | 28.6531892 |
| Dstyk | -0.8978702 | 3.401E-05 | 0.00047573 | 899.61384 | 482.656273 |
| Usp2 | -0.897805 | 0.00281186 | 0.01731963 | 66.8157099 | 35.8647549 |
| Rab21 | -0.8974993 | 3.865E-12 | 3.9232E-10 | 1095.03042 | 587.680234 |
| Mtor | -0.8974127 | 4.7199E-06 | 9.0559E-05 | 287.306259 | 154.41835 |
| Bcor | -0.8972957 | 8.9292E-05 | 0.00107058 | 381.533806 | 204.789529 |
| Aida | -0.8968845 | 5.1664E-08 | 1.7481E-06 | 391.924888 | 210.388714 |
| Rad21 | -0.8963266 | 1.4568E-06 | 3.2519E-05 | 490.680109 | 263.742215 |
| Slc36a1 | -0.8959838 | 0.00094723 | 0.00726031 | 183.604923 | 98.724045 |
| Arfgef2 | -0.8946224 | 3.9809E-05 | 0.00054171 | 242.435445 | 130.45236 |
| Tmem250-ps | -0.8931867 | 2.0093E-07 | 5.717E-06 | 269.831919 | 145.172635 |
| Klhl42 | -0.8930367 | 0.00013275 | 0.0014784 | 255.320315 | 137.341347 |
| Abcc5 | -0.8928309 | 2.9618E-06 | 6.0455E-05 | 959.980203 | 516.948918 |
| Lrba | -0.8925878 | 0.00641782 | 0.03264982 | 54.4645408 | 29.2756233 |
| Smg1 | -0.8923953 | 2.8622E-06 | 5.8741E-05 | 566.445547 | 305.268788 |
| Cpeb2 | -0.8919898 | 0.00092304 | 0.00712951 | 171.001353 | 92.0554902 |

| | | | | | |
|---|---|---|---|---|---|
| Prkag2 | -0.8917897 | 5.9904E-11 | 4.4115E-09 | 760.581482 | 409.592002 |
| Map4 | -0.8891539 | 3.2445E-05 | 0.00045425 | 1481.3099 | 799.708521 |
| Apbb1ip | -0.8890842 | 2.9982E-12 | 3.1945E-10 | 1583.6644 | 854.973909 |
| Nsd3 | -0.8889923 | 3.4396E-12 | 3.6064E-10 | 1258.19506 | 679.121597 |
| Eps15l1 | -0.8887704 | 1.1442E-09 | 5.8668E-08 | 680.419327 | 367.335233 |
| Gm15265 | -0.8885772 | 0.00296614 | 0.01801874 | 51.8213862 | 28.0183608 |
| Iqgap1 | -0.8878381 | 3.5016E-06 | 6.9582E-05 | 11794.2457 | 6373.64115 |
| Apba1 | -0.8875028 | 9.1371E-08 | 2.8657E-06 | 470.573977 | 254.314517 |
| Tmem150b | -0.8872117 | 2.0531E-05 | 0.0003103 | 232.728674 | 125.928908 |
| Zcchc24 | -0.8865495 | 2.3234E-08 | 8.576E-07 | 1513.66161 | 818.486782 |
| Trrap | -0.8863713 | 2.9318E-06 | 6.0006E-05 | 627.245338 | 339.520749 |
| 2700016F22F | -0.8863244 | 9.6083E-06 | 0.00016403 | 310.512272 | 167.909364 |
| Otud4 | -0.8856684 | 6.8696E-06 | 0.00012389 | 332.620768 | 179.872484 |
| Fbrsl1 | -0.8851773 | 1.8861E-07 | 5.4594E-06 | 841.095684 | 455.406818 |
| Ercc2 | -0.8850761 | 0.0001197 | 0.00136442 | 121.522731 | 65.917808 |
| Rfx2 | -0.8839716 | 2.6515E-08 | 9.645E-07 | 720.988765 | 390.492606 |
| Esyt2 | -0.883965 | 8.1421E-07 | 1.9697E-05 | 679.870646 | 368.328574 |
| Ankib1 | -0.8835377 | 3.4071E-05 | 0.00047614 | 391.086482 | 211.950906 |
| Arhgap31 | -0.8835065 | 1.3289E-05 | 0.00021587 | 500.259397 | 271.159358 |
| 4933416M07 | -0.8833332 | 0.00024099 | 0.00241364 | 73.8485338 | 40.0056741 |
| Slc12a6 | -0.8832033 | 3.4152E-11 | 2.7146E-09 | 1069.26558 | 579.554117 |
| Lpar5 | -0.8830925 | 2.3347E-06 | 4.9124E-05 | 478.247466 | 259.178542 |
| Cbl | -0.8822657 | 6.2802E-10 | 3.4184E-08 | 1869.7638 | 1014.27033 |
| Taok2 | -0.882258 | 2.1157E-06 | 4.5407E-05 | 788.935916 | 427.920213 |
| Trp53inp1 | -0.8811777 | 5.8899E-06 | 0.00010924 | 691.952642 | 375.546898 |
| Kat6b | -0.8803742 | 0.00217235 | 0.01411561 | 101.647254 | 55.232136 |
| Ahctf1 | -0.8799315 | 3.2635E-06 | 6.537E-05 | 254.431189 | 138.362027 |
| Dock5 | -0.8796603 | 1.5114E-06 | 3.3588E-05 | 761.401167 | 413.736449 |
| Il6st | -0.8796178 | 1.9719E-07 | 5.6318E-06 | 2116.70719 | 1150.32152 |
| Dhcr7 | -0.8795124 | 7.3984E-06 | 0.00013185 | 329.747162 | 179.353082 |
| Gm28437 | -0.8794544 | 5.2045E-05 | 0.00068167 | 226.24977 | 122.910143 |
| 9930014A18F | -0.8792894 | 0.00042964 | 0.00383103 | 178.356772 | 96.8708976 |
| Atp6v0a1 | -0.8786825 | 3.4708E-06 | 6.9246E-05 | 630.172744 | 342.497302 |
| Gm26616 | -0.8780685 | 1.0124E-06 | 2.3669E-05 | 370.930212 | 201.584628 |
| Smarcd1 | -0.8776164 | 2.091E-05 | 0.00031508 | 202.160957 | 109.847054 |
| Hivep1 | -0.87754 | 0.0011301 | 0.00842137 | 158.089538 | 86.1708909 |
| Sema4b | -0.8772206 | 0.00087484 | 0.00683452 | 119.040417 | 64.9557302 |
| Kdm3b | -0.8762644 | 1.411E-10 | 9.2569E-09 | 973.79902 | 530.315061 |
| Wdfy4 | -0.8755708 | 9.4229E-06 | 0.00016125 | 773.809841 | 421.630631 |
| Lats2 | -0.8754365 | 4.3954E-05 | 0.00058852 | 811.928478 | 442.315388 |
| Zbtb45 | -0.8754294 | 2.4082E-06 | 5.0318E-05 | 373.566818 | 203.52223 |
| Fbrs | -0.8746587 | 0.00059984 | 0.00503426 | 267.857228 | 146.145712 |
| Cttnbp2nl | -0.8746083 | 0.00199061 | 0.01316821 | 142.68043 | 77.8282423 |
| Adcy7 | -0.8745167 | 1.7675E-07 | 5.166E-06 | 2852.21489 | 1555.62849 |
| Hipk2 | -0.8739086 | 5.4363E-06 | 0.00010221 | 412.26234 | 224.989908 |
| 4931406P16F | -0.8735747 | 7.6406E-06 | 0.0001352 | 428.363387 | 233.60102 |

| | | | | | |
|---|---|---|---|---|---|
| 1700084C06F | -0.872758 | 5.3E-06 | 0.00010015 | 158.297646 | 86.5248156 |
| Man2a1 | -0.8726828 | 6.3009E-06 | 0.00011544 | 450.557278 | 246.181488 |
| Fam126a | -0.8725727 | 0.0001803 | 0.00188227 | 1002.77945 | 547.537154 |
| Tra2a | -0.8724206 | 8.0143E-05 | 0.00098125 | 639.815685 | 349.449984 |
| Myo5a | -0.8720225 | 3.0612E-06 | 6.1812E-05 | 1115.03495 | 609.24885 |
| Iqcg | -0.8716063 | 2.4257E-05 | 0.00035692 | 402.276216 | 219.713762 |
| Snapc4 | -0.8714786 | 0.00031922 | 0.00301163 | 222.897542 | 121.83101 |
| Gm26720 | -0.8712714 | 0.00423855 | 0.02363615 | 58.6957502 | 32.109366 |
| 9330175E14F | -0.8708647 | 0.00011558 | 0.00132379 | 169.646986 | 92.8413914 |
| Mib1 | -0.8707221 | 9.947E-06 | 0.00016904 | 502.232674 | 274.503354 |
| Kif13b | -0.8706057 | 9.6119E-05 | 0.00114149 | 170.102629 | 93.095276 |
| Dmxl2 | -0.8705962 | 0.00289866 | 0.01775237 | 175.559515 | 96.084343 |
| Hcls1 | -0.86991 | 2.8926E-15 | 5.6436E-13 | 3250.75933 | 1778.62094 |
| Man1b1 | -0.8688395 | 2.4441E-09 | 1.1656E-07 | 484.809658 | 265.469569 |
| Shprh | -0.8687964 | 0.00020183 | 0.00207816 | 343.93321 | 188.328217 |
| Ass1 | -0.8684884 | 1.9846E-05 | 0.00030207 | 155.691047 | 85.1818568 |
| Dapk1 | -0.8684196 | 0.00060452 | 0.00506792 | 329.316184 | 180.342167 |
| Ssh2 | -0.8681469 | 6.2987E-09 | 2.7428E-07 | 5771.48294 | 3161.6861 |
| Pign | -0.8680362 | 3.2107E-06 | 6.4485E-05 | 225.937346 | 123.830311 |
| Dyrk2 | -0.8679596 | 1.69E-05 | 0.00026066 | 598.059467 | 327.671158 |
| Xpo1 | -0.8676391 | 1.6268E-10 | 1.0444E-08 | 1083.63947 | 593.662519 |
| Herc1 | -0.8674447 | 5.8917E-07 | 1.4752E-05 | 812.869602 | 445.660427 |
| Elovl6 | -0.8669232 | 1.985E-06 | 4.2723E-05 | 208.45578 | 114.423255 |
| Gm37472 | -0.866773 | 0.00351889 | 0.02049003 | 128.120568 | 70.3566296 |
| Helz | -0.8658775 | 3.6715E-07 | 9.7142E-06 | 433.423307 | 237.793098 |
| Pik3cg | -0.8656879 | 2.9436E-05 | 0.00041853 | 2299.00489 | 1261.4447 |
| Ascc3 | -0.864142 | 2.4712E-08 | 9.0548E-07 | 587.488889 | 322.904067 |
| Kifc3 | -0.8640227 | 9.2246E-07 | 2.1653E-05 | 594.163299 | 326.222124 |
| Abca9 | -0.8636339 | 8.6543E-09 | 3.6317E-07 | 2122.27149 | 1166.31591 |
| March8 | -0.8636119 | 1.7888E-07 | 5.2079E-06 | 342.468179 | 188.217547 |
| Abr | -0.8629347 | 5.7824E-06 | 0.00010751 | 3133.14175 | 1722.51453 |
| Kcnk6 | -0.8617882 | 7.0359E-10 | 3.775E-08 | 455.099042 | 250.176735 |
| Clock | -0.8614187 | 3.9459E-06 | 7.7389E-05 | 326.205761 | 179.629197 |
| Pikfyve | -0.8612643 | 2.4831E-06 | 5.1739E-05 | 402.542346 | 221.574376 |
| Ccnk | -0.861013 | 0.00532683 | 0.02837698 | 51.2429529 | 28.2390001 |
| Ncapd3 | -0.8606835 | 0.00030942 | 0.00294759 | 384.56862 | 211.721355 |
| Gm26798 | -0.8604733 | 0.00736467 | 0.03620401 | 51.1160751 | 28.1261598 |
| Atp2a2 | -0.8603446 | 5.5753E-07 | 1.403E-05 | 2513.97167 | 1384.49743 |
| Phc1 | -0.8594946 | 0.00130131 | 0.00940788 | 97.5240733 | 53.8914344 |
| Eif4g2 | -0.858286 | 3.7472E-08 | 1.3092E-06 | 9828.59843 | 5421.34082 |
| Neurl4 | -0.8576354 | 0.00257865 | 0.01615471 | 162.292985 | 89.7158732 |
| Urm1 | -0.8572049 | 1.2327E-05 | 0.00020217 | 210.615507 | 116.120863 |
| 9930021J03R | -0.8566077 | 8.8517E-05 | 0.00106214 | 212.079478 | 117.196614 |
| Ahnak | -0.8562651 | 5.8357E-07 | 1.4636E-05 | 9595.67916 | 5300.44921 |
| Gm26590 | -0.8562092 | 0.00596698 | 0.0308736 | 45.9758085 | 25.4622918 |
| Ralgapa1 | -0.8559622 | 8.7507E-08 | 2.7847E-06 | 709.169364 | 391.794902 |

| | | | | | |
|---|---|---|---|---|---|
| Kmt2b | -0.8554209 | 0.00091002 | 0.00705432 | 234.80232 | 129.880624 |
| Csrnp1 | -0.855123 | 3.7753E-06 | 7.4358E-05 | 229.109311 | 126.786216 |
| Slc35e1 | -0.8546563 | 2.1964E-06 | 4.6737E-05 | 2249.35356 | 1243.6621 |
| Polr3gl | -0.8545028 | 6.2385E-05 | 0.00079196 | 124.282458 | 68.7374774 |
| AC162182.1 | -0.8543012 | 5.1528E-07 | 1.301E-05 | 535.896239 | 296.261926 |
| Gm5960 | -0.853955 | 0.00405577 | 0.02283729 | 85.6882589 | 47.3681869 |
| Gm11508 | -0.8535264 | 3.6956E-05 | 0.00050888 | 112.248171 | 61.9712892 |
| Emc1 | -0.8528811 | 2.9018E-05 | 0.00041399 | 1304.91028 | 722.379404 |
| Nfatc3 | -0.8523857 | 6.3446E-06 | 0.00011581 | 945.949586 | 523.768925 |
| Dock4 | -0.8516289 | 2.5998E-05 | 0.00037874 | 576.533068 | 319.751943 |
| Abcc1 | -0.8513658 | 2.159E-07 | 6.0853E-06 | 748.763018 | 414.787752 |
| Cdc7 | -0.8510566 | 0.0001987 | 0.00205156 | 96.9796668 | 53.9117933 |
| Kif3c | -0.8504122 | 0.00011336 | 0.00130401 | 273.279363 | 151.444859 |
| Traf3 | -0.8503961 | 2.0236E-07 | 5.7468E-06 | 579.032653 | 321.077215 |
| Xpo5 | -0.8503275 | 3.135E-05 | 0.00044181 | 471.481597 | 261.333304 |
| Ube4bos1 | -0.8501704 | 0.00138202 | 0.00985852 | 94.6583351 | 52.374842 |
| Klhdc10 | -0.8500325 | 6.9313E-07 | 1.7014E-05 | 402.368974 | 223.22518 |
| Daam1 | -0.8498977 | 0.00114068 | 0.00847919 | 116.802949 | 64.8307979 |
| Dnmt1 | -0.8493704 | 0.00286167 | 0.0175832 | 902.720989 | 500.96751 |
| Dhx37 | -0.8492728 | 0.00056013 | 0.00477033 | 273.237048 | 151.488703 |
| Ykt6 | -0.8464992 | 6.4212E-12 | 6.1467E-10 | 1014.3262 | 563.890829 |
| Plk2 | -0.8463792 | 0.00219769 | 0.01423099 | 52.2847653 | 29.1220334 |
| Slc12a7 | -0.8462409 | 3.1858E-05 | 0.00044729 | 374.099819 | 207.969731 |
| Smurf1 | -0.8449086 | 1.797E-05 | 0.00027631 | 453.078778 | 252.264984 |
| Abi2 | -0.8446882 | 2.4099E-05 | 0.00035528 | 213.325036 | 118.803478 |
| Fam193a | -0.844399 | 0.00174654 | 0.01193194 | 258.861785 | 144.085923 |
| Osbpl7 | -0.8442057 | 4.0527E-06 | 7.9275E-05 | 296.997174 | 165.437732 |
| Retreg3 | -0.8438727 | 8.8289E-05 | 0.00106109 | 569.309193 | 317.083617 |
| Pou2f2 | -0.8436165 | 7.2971E-07 | 1.7825E-05 | 1379.89616 | 768.694658 |
| Tmem216 | -0.8435373 | 0.0001983 | 0.00204888 | 211.350727 | 117.92084 |
| 4931440J10R | -0.8433879 | 0.00990621 | 0.04501543 | 54.8703196 | 30.6359266 |
| Foxj3 | -0.842485 | 2.4683E-05 | 0.00036282 | 380.490932 | 212.079877 |
| Fam193b | -0.8414748 | 1.3873E-06 | 3.1015E-05 | 402.795596 | 224.980502 |
| Gtf3c4 | -0.8411375 | 0.00069415 | 0.00566721 | 279.080528 | 155.846671 |
| Crebbp | -0.8397164 | 2.6105E-08 | 9.5187E-07 | 1091.50673 | 609.81448 |
| Tnks | -0.8391177 | 1.2087E-06 | 2.7513E-05 | 646.948907 | 361.486362 |
| Vps13c | -0.8390079 | 1.4879E-05 | 0.00023604 | 470.707826 | 263.154078 |
| Gm20457 | -0.8364352 | 0.00275001 | 0.01700822 | 78.039611 | 43.6383188 |
| Dock7 | -0.8360519 | 2.0321E-05 | 0.00030743 | 296.399872 | 165.958197 |
| Capn12 | -0.8358655 | 0.0102649 | 0.04612911 | 60.4012757 | 33.8458563 |
| Khnyn | -0.8353111 | 4.0043E-06 | 7.843E-05 | 1356.46613 | 760.199234 |
| Asb1 | -0.8350886 | 0.00157156 | 0.01099511 | 139.667488 | 78.257821 |
| Prtn3 | -0.8345303 | 0.00559943 | 0.0293921 | 392.55286 | 220.134859 |
| Zfp91 | -0.8340215 | 2.9457E-08 | 1.0638E-06 | 532.04162 | 298.631148 |
| Ecpas | -0.8333978 | 2.4169E-10 | 1.4781E-08 | 735.075392 | 412.445003 |
| Hddc3 | -0.8310603 | 0.00010765 | 0.00125461 | 138.513919 | 77.751032 |

| | | | | | |
|---|---|---|---|---|---|
| Kbtbd11 | -0.8309335 | 8.0226E-06 | 0.0001413 | 242.947961 | 136.634249 |
| Gm44751 | -0.8308142 | 0.00767643 | 0.03736972 | 39.2431339 | 22.1092133 |
| Rnf169 | -0.8305432 | 1.1101E-06 | 2.5494E-05 | 502.896052 | 282.762011 |
| Zswim6 | -0.8288725 | 0.00098861 | 0.00753904 | 316.343005 | 178.161472 |
| D330041H03I | -0.828791 | 2.7246E-06 | 5.6225E-05 | 433.934415 | 244.19977 |
| C4b | -0.8284754 | 0.00507795 | 0.0273035 | 132.905566 | 74.7984975 |
| Zbed4 | -0.8278081 | 0.00150521 | 0.0105915 | 104.108902 | 58.4960484 |
| Rin3 | -0.8278023 | 6.6792E-10 | 3.597E-08 | 4333.49155 | 2441.14452 |
| Vasp | -0.827488 | 3.2963E-11 | 2.634E-09 | 1824.86159 | 1028.10766 |
| Nup214 | -0.8274426 | 3.6276E-07 | 9.6456E-06 | 741.907517 | 418.060713 |
| N4bp2 | -0.8273122 | 0.00013029 | 0.00145425 | 159.483037 | 89.7821828 |
| Trim24 | -0.8272102 | 0.00125396 | 0.00913634 | 111.205208 | 62.5084638 |
| Gm44777 | -0.8268302 | 0.00490766 | 0.02653989 | 68.1732421 | 38.3845742 |
| Trerf1 | -0.8235767 | 0.00015647 | 0.00168024 | 299.844184 | 169.506 |
| Tmem259 | -0.8230384 | 5.7794E-05 | 0.00074592 | 558.742495 | 315.832148 |
| Gm26716 | -0.823017 | 9.3386E-05 | 0.00111167 | 467.978016 | 264.355039 |
| Dcp1a | -0.8226317 | 1.1319E-05 | 0.00018852 | 522.794218 | 295.497016 |
| Cbfa2t3 | -0.8213796 | 1.3484E-07 | 4.0678E-06 | 1742.53142 | 986.011412 |
| Map3k20 | -0.8197447 | 0.00016081 | 0.00171701 | 308.378735 | 174.609172 |
| Arhgef37 | -0.818885 | 0.00013617 | 0.001505 | 2821.94115 | 1599.49395 |
| Ralgapa2 | -0.8186023 | 0.00035263 | 0.00325604 | 243.667151 | 138.228404 |
| Slmap | -0.8181759 | 9.1074E-08 | 2.8624E-06 | 865.723582 | 490.753923 |
| Cemip2 | -0.8176908 | 4.5166E-05 | 0.00060153 | 292.408395 | 165.961315 |
| mt-Rnr2 | -0.8174727 | 0.0003143 | 0.00298275 | 14514.6358 | 8236.05036 |
| Ulk2 | -0.8174085 | 1.8307E-08 | 6.9625E-07 | 803.440201 | 455.807741 |
| Slc26a2 | -0.8170218 | 8.1575E-12 | 7.5184E-10 | 803.166215 | 455.860184 |
| Atxn7 | -0.8161631 | 4.28E-07 | 1.1111E-05 | 453.462192 | 257.646034 |
| Zbtb18 | -0.8161275 | 3.0612E-05 | 0.00043263 | 202.600454 | 115.095145 |
| Rad23b | -0.8161207 | 5.1314E-08 | 1.7402E-06 | 390.602715 | 221.93468 |
| Parp14 | -0.8157239 | 5.5246E-05 | 0.00071645 | 1248.02012 | 708.980287 |
| Slc35c1 | -0.8151097 | 1.8875E-09 | 9.2363E-08 | 2691.28133 | 1529.27031 |
| Ptpn12 | -0.8149451 | 8.1354E-05 | 0.00099283 | 368.170746 | 209.35572 |
| U2af2 | -0.8139993 | 5.1728E-10 | 2.8889E-08 | 606.394087 | 344.71284 |
| Gm15601 | -0.8139115 | 4.5705E-06 | 8.8145E-05 | 231.256191 | 131.504471 |
| Mboat7 | -0.8128299 | 3.8588E-09 | 1.7567E-07 | 1505.79485 | 856.921933 |
| Lilrb4a | -0.8126896 | 0.00011586 | 0.00132467 | 1319.91074 | 751.652611 |
| Helz2 | -0.8117948 | 0.00017303 | 0.00182338 | 447.716778 | 255.088985 |
| Asb7 | -0.8103098 | 0.00147534 | 0.01043508 | 98.7042097 | 56.2309447 |
| Topbp1 | -0.8100534 | 1.8913E-05 | 0.00028875 | 519.601605 | 296.130001 |
| Ankrd17 | -0.8093336 | 2.8145E-06 | 5.7841E-05 | 591.750841 | 337.781836 |
| Ocrl | -0.8089473 | 3.1478E-06 | 6.3306E-05 | 232.395915 | 132.626088 |
| Ddhd1 | -0.8087073 | 2.2248E-06 | 4.7208E-05 | 1286.2626 | 734.305857 |
| Sfpq | -0.8083726 | 5.3392E-08 | 1.7864E-06 | 1195.85726 | 682.734493 |
| Btaf1 | -0.8081395 | 0.00143725 | 0.01018769 | 654.210085 | 373.770981 |
| Rbm41 | -0.8076819 | 0.0002014 | 0.00207521 | 108.786637 | 62.1043847 |
| Dcun1d3 | -0.8074545 | 0.00119335 | 0.00876403 | 158.22682 | 90.3448651 |

| | | | | | |
|---|---|---|---|---|---|
| Fryl | -0.8074114 | 1.5293E-05 | 0.00024108 | 488.803214 | 279.415497 |
| Pum2 | -0.8073387 | 0.00046192 | 0.00405861 | 401.194043 | 229.264547 |
| Szrd1 | -0.8071801 | 2.3118E-09 | 1.1096E-07 | 979.547799 | 559.947147 |
| Jup | -0.806719 | 0.00211893 | 0.01387137 | 348.478392 | 199.05428 |
| Mapk3 | -0.8047564 | 1.2201E-08 | 4.8619E-07 | 2203.22624 | 1261.02968 |
| Adamtsl4 | -0.8042074 | 0.00047588 | 0.00415977 | 134.071001 | 76.6869861 |
| Gm15491 | -0.8040659 | 5.2977E-05 | 0.00069206 | 321.409478 | 183.87676 |
| Amigo1 | -0.8039884 | 0.00086083 | 0.00673946 | 75.9608396 | 43.4455033 |
| Ncoa2 | -0.8036538 | 3.049E-06 | 6.1735E-05 | 516.491189 | 295.793981 |
| Gm16235 | -0.8034918 | 0.0041112 | 0.02306294 | 66.7328618 | 38.1169294 |
| R3hdm2 | -0.803352 | 9.2153E-06 | 0.00015855 | 436.227854 | 249.859354 |
| Cacna1d | -0.8031772 | 0.00072517 | 0.00586254 | 109.175285 | 62.6083932 |
| Ppp1r12b | -0.8028105 | 0.00859925 | 0.04053548 | 99.7827441 | 57.3863565 |
| Rbm15b | -0.802286 | 4.1084E-05 | 0.00055515 | 527.735235 | 302.612681 |
| Crtc3 | -0.8022072 | 2.5747E-05 | 0.00037627 | 469.497397 | 269.321902 |
| Dock1 | -0.8015419 | 3.0499E-06 | 6.1735E-05 | 590.71274 | 338.95856 |
| Pias3 | -0.8008728 | 8.0909E-06 | 0.00014202 | 283.767002 | 162.971792 |
| Mbtd1 | -0.7994316 | 1.1807E-10 | 7.9539E-09 | 474.376469 | 272.646509 |
| Ttf2 | -0.7988594 | 0.00082746 | 0.00651515 | 170.535735 | 97.924857 |
| Gemin5 | -0.7988404 | 0.00093826 | 0.00721735 | 289.669447 | 166.496514 |
| Ppip5k1 | -0.7984873 | 0.00014113 | 0.00154534 | 170.983015 | 98.2892597 |
| Acot11 | -0.7978377 | 0.00020463 | 0.00210267 | 403.351971 | 231.817308 |
| Lipe | -0.7977786 | 6.3687E-11 | 4.6445E-09 | 779.516112 | 448.306595 |
| Gse1 | -0.7973702 | 0.00016221 | 0.0017283 | 306.633681 | 176.574844 |
| Kif21b | -0.7973292 | 0.00013834 | 0.00152255 | 327.345195 | 188.342495 |
| Med14 | -0.7970012 | 6.1439E-06 | 0.0001131 | 509.900155 | 293.345205 |
| Prcc | -0.7960311 | 0.00017529 | 0.00184279 | 152.868772 | 87.8361136 |
| Copg2 | -0.7959713 | 0.00059781 | 0.0050229 | 831.688065 | 479.027879 |
| Huwe1 | -0.7958108 | 0.00053026 | 0.00454424 | 481.102482 | 277.167234 |
| Usp32 | -0.7955687 | 3.0998E-06 | 6.2425E-05 | 302.188596 | 174.095844 |
| Mllt1 | -0.7955679 | 9.7213E-06 | 0.00016558 | 500.54109 | 288.180946 |
| Clec4a1 | -0.794556 | 2.2237E-08 | 8.2894E-07 | 2931.26486 | 1689.72564 |
| Skil | -0.7941446 | 0.00027482 | 0.00268096 | 478.51505 | 275.917216 |
| Mosmo | -0.7940988 | 0.00118038 | 0.00869256 | 107.842407 | 62.2683344 |
| Gm15243 | -0.7935213 | 0.00308994 | 0.01860528 | 77.0769509 | 44.3897988 |
| Abcc4 | -0.7933553 | 0.00035092 | 0.00325019 | 849.541358 | 490.093999 |
| Pank3 | -0.7926485 | 8.0923E-06 | 0.00014202 | 428.177919 | 247.061449 |
| 6430550D23I | -0.7925211 | 0.00331642 | 0.01957731 | 83.4123577 | 48.1039726 |
| Tug1 | -0.7923637 | 1.4953E-07 | 4.4571E-06 | 852.00771 | 491.873968 |
| Kif13a | -0.7909104 | 0.00016001 | 0.00171135 | 465.030077 | 268.594233 |
| Med13l | -0.790537 | 2.7468E-06 | 5.6605E-05 | 505.210756 | 292.184647 |
| Akap13 | -0.790531 | 8.0741E-09 | 3.4168E-07 | 7664.46672 | 4430.70019 |
| Phrf1 | -0.7901644 | 0.00195113 | 0.0129412 | 1127.12209 | 651.764423 |
| Dag1 | -0.7897503 | 0.000374 | 0.00343045 | 448.896299 | 259.453068 |
| Fam53b | -0.7895051 | 1.5499E-11 | 1.3382E-09 | 1127.03794 | 651.918928 |
| Cep250 | -0.7891718 | 5.5888E-06 | 0.00010427 | 539.228888 | 312.065235 |

| | | | | | |
|---|---|---|---|---|---|
| Synrg | -0.7881759 | 2.252E-05 | 0.00033463 | 566.229497 | 327.884781 |
| Golga3 | -0.7881225 | 0.00014626 | 0.00159126 | 535.227618 | 309.789377 |
| Phlda1 | -0.7872815 | 0.00542828 | 0.02873467 | 80.9891563 | 46.9970007 |
| A930037H05I | -0.7870296 | 0.0021668 | 0.01409782 | 263.100855 | 152.475375 |
| Map3k14 | -0.7865637 | 0.00031768 | 0.0030016 | 1149.06174 | 666.035133 |
| Gm17344 | -0.7854608 | 4.2265E-06 | 8.2141E-05 | 255.072194 | 147.829569 |
| Uso1 | -0.7849265 | 1.0521E-05 | 0.000177 | 1290.64609 | 749.00602 |
| Lbp | -0.7845582 | 0.00158137 | 0.01102922 | 268.097097 | 155.603618 |
| Pi4kb | -0.7834162 | 0.0001132 | 0.00130317 | 303.961806 | 176.460004 |
| Mdm4 | -0.783182 | 1.3292E-05 | 0.00021587 | 591.903692 | 344.152653 |
| Card10 | -0.7830073 | 0.0094412 | 0.04342777 | 247.086675 | 143.455154 |
| Gm22107 | -0.7829707 | 0.00401793 | 0.0226752 | 89.0715673 | 51.7362906 |
| Ccdc6 | -0.7826315 | 1.4496E-05 | 0.00023118 | 216.595771 | 126.076143 |
| Clcn5 | -0.7824622 | 0.00076681 | 0.00611781 | 103.273329 | 59.9981504 |
| Fam8a1 | -0.7824062 | 0.0051544 | 0.02762559 | 93.1355987 | 54.1037701 |
| Gm12758 | -0.7819552 | 0.00064404 | 0.00534255 | 165.65656 | 96.2741825 |
| Abca1 | -0.7807426 | 0.00378766 | 0.02171002 | 211.976207 | 123.277277 |
| Smurf2 | -0.7807257 | 0.00108101 | 0.00814039 | 402.839437 | 234.311978 |
| Srsf1 | -0.7804518 | 6.7749E-12 | 6.4012E-10 | 1604.61627 | 933.941623 |
| Mroh1 | -0.7797817 | 2.8968E-05 | 0.00041367 | 841.754468 | 490.08215 |
| Tsc1 | -0.779681 | 0.0001289 | 0.0014398 | 305.687893 | 177.893659 |
| Prkag2os2 | -0.7790476 | 0.00784103 | 0.03791388 | 55.4105063 | 32.287788 |
| Gpc1 | -0.778973 | 0.00094019 | 0.00722483 | 498.662015 | 290.642255 |
| Rara | -0.7786947 | 8.9821E-07 | 2.1183E-05 | 3851.35515 | 2244.56625 |
| Dop1a | -0.7782371 | 1.2289E-06 | 2.7889E-05 | 391.475681 | 228.336279 |
| Cep192 | -0.7778375 | 0.00010495 | 0.00122794 | 276.246417 | 161.101997 |
| Rgcc | -0.7769819 | 0.00272556 | 0.01691285 | 133.459089 | 77.853916 |
| Gbf1 | -0.7768946 | 4.7697E-06 | 9.1179E-05 | 823.669015 | 480.666745 |
| Zfyve28 | -0.7762012 | 0.00089906 | 0.00699461 | 317.731301 | 185.479459 |
| Mfhas1 | -0.7757888 | 5.079E-05 | 0.00066756 | 277.886584 | 162.330521 |
| Gm45091 | -0.775022 | 0.01101637 | 0.04852748 | 46.8702837 | 27.2666234 |
| Zfp605 | -0.7749289 | 0.00363276 | 0.0210714 | 63.4840481 | 37.0870517 |
| Mgea5 | -0.7747655 | 1.4667E-05 | 0.00023366 | 1666.42066 | 973.688966 |
| Irf4 | -0.7742592 | 1.1299E-07 | 3.4641E-06 | 573.54251 | 335.070898 |
| Zfat | -0.7735768 | 0.01083619 | 0.04796467 | 149.727018 | 87.5626343 |
| Tmx4 | -0.7733926 | 3.021E-06 | 6.1496E-05 | 621.599729 | 363.548575 |
| Tbc1d9 | -0.7732257 | 0.00178015 | 0.0121134 | 241.824345 | 141.445753 |
| Ube3b | -0.772127 | 0.0023001 | 0.01477946 | 1653.81954 | 968.275286 |
| Trmt61a | -0.7718536 | 5.1276E-05 | 0.00067335 | 156.243804 | 91.422426 |
| Sema4d | -0.7715665 | 0.00024358 | 0.00242825 | 7658.09136 | 4485.76969 |
| Kmt2c | -0.7712855 | 4.0822E-06 | 7.9748E-05 | 842.639522 | 493.86073 |
| Cox6b2 | -0.770861 | 0.00509945 | 0.02738061 | 163.466886 | 95.7881443 |
| Gm38365 | -0.7700779 | 0.01022079 | 0.04602708 | 69.1168271 | 40.5149095 |
| Ctbp2 | -0.7700647 | 1.115E-05 | 0.00018633 | 651.859755 | 382.10998 |
| Phf8 | -0.7699244 | 3.1418E-05 | 0.00044236 | 921.275425 | 540.078099 |
| Cplane1 | -0.7692956 | 0.01024054 | 0.04608855 | 115.163482 | 67.6161988 |

| | | | | | |
|---|---|---|---|---|---|
| 4732487G21I | -0.768484 | 0.00062107 | 0.00518064 | 141.440117 | 82.9912159 |
| Dock10 | -0.7684006 | 1.7114E-07 | 5.0216E-06 | 3938.07071 | 2311.84365 |
| Smad5 | -0.7683229 | 2.7781E-05 | 0.000399 | 242.826076 | 142.403264 |
| Dot1l | -0.7673758 | 0.0011774 | 0.00867913 | 192.62374 | 113.198266 |
| Fam117b | -0.7668755 | 4.6627E-06 | 8.969E-05 | 1609.14659 | 945.45869 |
| Prpf8 | -0.765865 | 1.5002E-05 | 0.00023773 | 1011.78217 | 595.055359 |
| Ammecr1 | -0.765255 | 3.899E-05 | 0.00053249 | 141.21488 | 83.0590314 |
| Ndst1 | -0.7642982 | 0.00101813 | 0.00774059 | 492.059286 | 289.599442 |
| Hmgxb3 | -0.7631597 | 0.00077407 | 0.00614961 | 525.426497 | 309.514901 |
| Stard10 | -0.7621072 | 0.00224909 | 0.01449508 | 70.3385377 | 41.517421 |
| Setx | -0.7608998 | 8.3745E-05 | 0.00101453 | 539.520244 | 318.29467 |
| Scnn1a | -0.760881 | 7.793E-05 | 0.00095649 | 227.87505 | 134.418782 |
| Sh3pxd2b | -0.7579591 | 0.00690357 | 0.03450872 | 86.3700236 | 51.0642936 |
| Limk1 | -0.7575696 | 0.00797561 | 0.03836621 | 282.125272 | 166.976013 |
| Gatad2b | -0.7572403 | 0.00472588 | 0.02564919 | 292.210265 | 172.869582 |
| Stard8 | -0.7571882 | 0.00136109 | 0.00974162 | 1248.92088 | 738.794717 |
| Gm13562 | -0.7568007 | 0.00259721 | 0.01625744 | 76.4916984 | 45.3684206 |
| Dusp3 | -0.7558386 | 1.1841E-06 | 2.7075E-05 | 905.833693 | 536.392043 |
| Gm26563 | -0.7554211 | 0.00804885 | 0.03858259 | 46.7640793 | 27.7030637 |
| Dyrk1a | -0.7553738 | 6.0038E-06 | 0.00011067 | 482.012263 | 285.412716 |
| Cdan1 | -0.7539568 | 0.00021846 | 0.00222054 | 342.312509 | 202.858114 |
| Taf4 | -0.7536877 | 0.00057798 | 0.00488904 | 240.877119 | 143.008991 |
| Ogt | -0.753478 | 7.4532E-07 | 1.8118E-05 | 2595.30935 | 1539.6364 |
| Camk1d | -0.7533823 | 2.4027E-07 | 6.6354E-06 | 2223.8453 | 1319.05703 |
| Naip5 | -0.7513992 | 8.6852E-07 | 2.07E-05 | 880.56499 | 522.920207 |
| Cep85l | -0.7513059 | 0.00040942 | 0.00367429 | 246.985296 | 146.755978 |
| Heg1 | -0.7509474 | 0.00963312 | 0.04402748 | 335.248503 | 199.144975 |
| Ppp4r1 | -0.7508323 | 1.2224E-05 | 0.0002007 | 1439.17246 | 855.266119 |
| Gm15651 | -0.7504563 | 0.00292849 | 0.01784776 | 104.726776 | 62.1858402 |
| Trim56 | -0.7504109 | 0.00076002 | 0.00607654 | 287.983135 | 171.241272 |
| Dgkq | -0.7503262 | 8.9413E-05 | 0.00107118 | 429.591777 | 255.38353 |
| Oas2 | -0.7500763 | 0.00013307 | 0.00148079 | 961.168213 | 571.525888 |
| Srgap2 | -0.7497837 | 1.2499E-08 | 4.9546E-07 | 1066.59179 | 634.130621 |
| Hmg20b | -0.7488997 | 8.8467E-08 | 2.7921E-06 | 363.525469 | 216.38261 |
| Stx4a | -0.7478318 | 0.00056625 | 0.00480454 | 850.700068 | 506.533946 |
| Gm44649 | -0.7473379 | 0.0013495 | 0.00967248 | 1128.76282 | 672.164482 |
| Kat6a | -0.7471115 | 3.6156E-05 | 0.00050108 | 791.926698 | 471.577694 |
| Tmcc1 | -0.7469106 | 2.3642E-06 | 4.9554E-05 | 3138.75982 | 1870.36541 |
| Slfn5os | -0.7451192 | 9.5326E-05 | 0.00113298 | 628.686833 | 374.788972 |
| Trps1 | -0.7446632 | 2.8379E-08 | 1.0298E-06 | 2210.17522 | 1319.0625 |
| Ino80 | -0.7434368 | 0.0005261 | 0.00451889 | 164.897944 | 98.5600269 |
| Gab1 | -0.7422869 | 0.00224499 | 0.01448113 | 227.040262 | 135.635176 |
| Brpf3 | -0.7414297 | 0.00081188 | 0.00641268 | 129.39466 | 77.3166988 |
| 5031439G07I | -0.7413532 | 1.0453E-08 | 4.3142E-07 | 6439.46314 | 3851.58206 |
| Ireb2 | -0.7411165 | 0.00017602 | 0.00184921 | 401.918981 | 240.536697 |
| Rhobtb2 | -0.7395723 | 0.008225 | 0.03920182 | 173.97681 | 104.09472 |

| | | | | | |
|---|---|---|---|---|---|
| Rbm3 | -0.7390165 | 1.6876E-05 | 0.00026066 | 1448.82587 | 868.021033 |
| Ptk2b | -0.738701 | 0.00015561 | 0.00167216 | 3329.23804 | 1994.83742 |
| Ccdc88b | -0.7385407 | 0.0012014 | 0.00881281 | 871.917995 | 522.617957 |
| Ylpm1 | -0.7385182 | 1.036E-05 | 0.0001747 | 257.804286 | 154.698959 |
| Zc3h12c | -0.7382447 | 0.00566561 | 0.0297062 | 82.6787719 | 49.5566925 |
| Dazap1 | -0.7380486 | 1.133E-08 | 4.6254E-07 | 434.396419 | 260.40919 |
| Gm19582 | -0.7378335 | 0.0039231 | 0.02229917 | 99.1700865 | 59.6301906 |
| Spire1 | -0.7375849 | 0.00753429 | 0.03680451 | 98.7723575 | 59.1387721 |
| 1500004A13I | -0.7372624 | 0.00265523 | 0.01657227 | 94.4311014 | 56.6621199 |
| Tbl1xr1 | -0.736388 | 1.4142E-09 | 7.0658E-08 | 1263.31928 | 758.2229 |
| Dido1 | -0.7362479 | 2.7093E-07 | 7.387E-06 | 551.187425 | 331.039594 |
| 9630028H03I | -0.7356193 | 0.00797295 | 0.03836566 | 94.3220257 | 56.5395037 |
| Mtmr10 | -0.7354366 | 1.9596E-05 | 0.00029857 | 693.710878 | 416.440933 |
| Chd4 | -0.7344308 | 2.3213E-06 | 4.891E-05 | 749.558464 | 450.47763 |
| Purb | -0.7342429 | 1.8954E-07 | 5.476E-06 | 1189.86288 | 715.154341 |
| Itpr1 | -0.7338858 | 6.9064E-07 | 1.6981E-05 | 987.824514 | 594.037527 |
| Atp8b4 | -0.7338224 | 8.368E-07 | 2.0146E-05 | 2048.07448 | 1231.28476 |
| Il3ra | -0.733492 | 0.00346459 | 0.02024447 | 225.459177 | 135.598891 |
| Prkdc | -0.732623 | 0.00021429 | 0.00218703 | 319.049552 | 191.972027 |
| Hlx | -0.7325521 | 3.4989E-07 | 9.3198E-06 | 601.461484 | 361.965573 |
| Abcc10 | -0.732069 | 0.00067381 | 0.0055317 | 98.0604864 | 58.9471294 |
| Zfyve1 | -0.7319916 | 0.00189143 | 0.01267958 | 270.666272 | 163.049813 |
| Map2k7 | -0.7317036 | 9.8927E-06 | 0.00016831 | 393.756402 | 236.942572 |
| Gm37335 | -0.7311212 | 0.0019201 | 0.0128089 | 147.152786 | 88.4737682 |
| Rbm3os | -0.7306117 | 0.00135378 | 0.00969854 | 158.237854 | 95.2482202 |
| R3hdm1 | -0.7301919 | 1.2334E-07 | 3.751E-06 | 499.550471 | 301.020664 |
| Rel | -0.7299123 | 4.3266E-08 | 1.4846E-06 | 881.252135 | 531.336474 |
| Kmt2a | -0.72868 | 0.00030006 | 0.00288773 | 427.354191 | 257.944213 |
| Git1 | -0.7283614 | 2.3005E-07 | 6.4038E-06 | 634.188324 | 382.588757 |
| Dmxl1 | -0.7279707 | 0.00037572 | 0.0034396 | 199.099165 | 120.224743 |
| Cdk12 | -0.727597 | 8.1531E-05 | 0.00099419 | 275.971534 | 166.65276 |
| Snx19 | -0.7273494 | 0.00024316 | 0.00242567 | 591.639666 | 357.249658 |
| Pfkfb3 | -0.7271361 | 1.12E-05 | 0.00018675 | 614.200946 | 371.099926 |
| Gnptab | -0.7261851 | 7.4411E-06 | 0.00013229 | 1024.98555 | 619.53525 |
| Mkl1 | -0.7257934 | 0.00066375 | 0.00546084 | 2568.91789 | 1553.18888 |
| Ttll3 | -0.7257117 | 0.01105197 | 0.04861895 | 108.298446 | 65.5841019 |
| Tgm4 | -0.7255575 | 0.00488692 | 0.02643725 | 115.563866 | 69.8532008 |
| Acsf3 | -0.7248061 | 0.00461302 | 0.02519723 | 94.0016567 | 56.7264517 |
| Vkorc1l1 | -0.7239593 | 0.00010587 | 0.00123687 | 268.989488 | 162.657082 |
| Fbxw7 | -0.7238322 | 0.00012864 | 0.00143894 | 275.608138 | 166.741061 |
| Braf | -0.7236727 | 3.5721E-09 | 1.6461E-07 | 504.827425 | 305.630456 |
| Abcg1 | -0.7231425 | 1.654E-05 | 0.00025669 | 997.696291 | 604.282712 |
| Peli2 | -0.7230923 | 0.00335754 | 0.01977276 | 122.858172 | 74.3844237 |
| 9330160F10I | -0.7227004 | 0.0002417 | 0.00241763 | 311.820366 | 189.137856 |
| Usp42 | -0.7222679 | 2.3533E-05 | 0.00034763 | 288.334696 | 174.699365 |
| mt-Nd5 | -0.721883 | 0.00015142 | 0.00163653 | 4099.52419 | 2485.38068 |

| | | | | | |
|---|---|---|---|---|---|
| Trip12 | -0.7216526 | 7.2096E-07 | 1.764E-05 | 1019.08231 | 618.00775 |
| Gpatch2l | -0.7210363 | 0.00013977 | 0.00153492 | 707.479128 | 429.069813 |
| Dtx3 | -0.7209251 | 0.00374723 | 0.02155208 | 111.504432 | 67.6649513 |
| Togaram1 | -0.7208561 | 5.4192E-06 | 0.00010202 | 516.028627 | 313.140334 |
| Aknaos | -0.7207977 | 0.00209752 | 0.01375428 | 313.625101 | 190.166108 |
| Itga1 | -0.7204466 | 0.00112407 | 0.00838474 | 458.447897 | 278.091148 |
| Nup155 | -0.7198981 | 0.00461884 | 0.02521395 | 287.271654 | 174.242642 |
| Ttll12 | -0.7193595 | 1.1828E-05 | 0.00019592 | 184.131569 | 111.941305 |
| Zfp335 | -0.7187801 | 0.00290594 | 0.01777524 | 837.808816 | 508.980817 |
| Usp12 | -0.7182888 | 5.9088E-06 | 0.00010945 | 1214.67875 | 738.089118 |
| Larp1 | -0.7181685 | 4.6545E-07 | 1.1892E-05 | 2441.9364 | 1484.21963 |
| Drosha | -0.717273 | 0.00273922 | 0.0169574 | 228.296237 | 138.81473 |
| Mga | -0.7172656 | 0.00012262 | 0.00139016 | 415.046721 | 252.475114 |
| Nup98 | -0.7171027 | 9.0212E-05 | 0.00107817 | 598.070383 | 363.630477 |
| Chd1 | -0.7169656 | 2.1677E-06 | 4.6193E-05 | 771.310831 | 469.268827 |
| Thada | -0.7168646 | 2.3165E-05 | 0.00034321 | 202.734058 | 123.315799 |
| Trappc8 | -0.7166937 | 0.000202 | 0.00207855 | 1212.26396 | 737.461905 |
| Sipa1l1 | -0.7165835 | 0.0006811 | 0.00558223 | 207.80288 | 126.259956 |
| Ppp1r10 | -0.7162475 | 9.7128E-06 | 0.00016558 | 924.997975 | 562.83091 |
| Samd4b | -0.7152416 | 7.0431E-07 | 1.7261E-05 | 538.652293 | 327.91844 |
| Tada2b | -0.7151876 | 7.3506E-05 | 0.000909 | 320.456862 | 195.037689 |
| Zbtb22 | -0.7141736 | 6.1509E-06 | 0.0001131 | 421.573766 | 256.775449 |
| Git2 | -0.7135917 | 1.6881E-08 | 6.4693E-07 | 1783.6747 | 1087.7899 |
| Klf4 | -0.7133759 | 8.6946E-07 | 2.07E-05 | 956.868846 | 583.607874 |
| Lpl | -0.713206 | 0.00322879 | 0.01917994 | 849.717892 | 518.183337 |
| Cherp | -0.7128424 | 0.00016202 | 0.00172743 | 1265.4749 | 771.924469 |
| Agps | -0.7122659 | 6.9667E-06 | 0.00012504 | 1667.94486 | 1018.04348 |
| Susd6 | -0.712061 | 8.6491E-07 | 2.069E-05 | 2196.7043 | 1340.65608 |
| Pik3cd | -0.7118771 | 2.0973E-05 | 0.00031571 | 8092.27822 | 4940.31734 |
| Taf1 | -0.711598 | 0.0003266 | 0.00306658 | 285.917184 | 174.541069 |
| Ltn1 | -0.7114637 | 0.00021088 | 0.00215805 | 657.596587 | 401.523394 |
| 6230400D17I | -0.7106368 | 0.00012287 | 0.00139016 | 337.352894 | 206.046774 |
| Zswim4 | -0.7105785 | 0.0003239 | 0.00304689 | 1055.07166 | 644.582401 |
| Mef2d | -0.7103888 | 1.0939E-06 | 2.5283E-05 | 1350.32186 | 825.020771 |
| Atp6v0d1 | -0.710388 | 6.6761E-08 | 2.1898E-06 | 1898.2683 | 1159.79781 |
| Tmem164 | -0.7097436 | 1.2404E-05 | 0.00020321 | 3438.59593 | 2102.30581 |
| Ahcyl2 | -0.7089778 | 8.5248E-09 | 3.5873E-07 | 2674.29343 | 1635.77629 |
| Fbxo42 | -0.708949 | 1.2126E-08 | 4.8515E-07 | 703.176299 | 430.007335 |
| Map3k2 | -0.7086497 | 6.2581E-05 | 0.00079337 | 359.116319 | 219.753689 |
| Wfs1 | -0.7077886 | 0.00193576 | 0.01288475 | 326.529165 | 199.769724 |
| Slc25a28 | -0.7076329 | 1.4862E-06 | 3.3126E-05 | 583.037767 | 356.949545 |
| Mon2 | -0.7073503 | 0.00945855 | 0.04347928 | 256.560282 | 157.080609 |
| Fzd7 | -0.707219 | 0.00034085 | 0.0031687 | 237.80464 | 145.638002 |
| Usp45 | -0.7071902 | 8.0398E-05 | 0.00098276 | 458.993799 | 281.093482 |
| Rin2 | -0.7070248 | 0.00012881 | 0.0014398 | 1046.65995 | 640.921613 |
| Rptor | -0.7062549 | 1.5496E-05 | 0.00024377 | 433.02549 | 265.43453 |

| | | | | | |
|---|---|---|---|---|---|
| Stk35 | -0.7057445 | 3.1648E-05 | 0.00044489 | 222.386318 | 136.27425 |
| Rbpj | -0.7056903 | 0.00014501 | 0.00157973 | 680.12616 | 416.873068 |
| Cdk8 | -0.7051607 | 0.00131749 | 0.00949285 | 198.6225 | 121.82258 |
| Zfp874b | -0.7043298 | 0.00080198 | 0.00634445 | 159.604664 | 97.9478927 |
| Abcb10 | -0.704155 | 0.00014388 | 0.00156977 | 460.870076 | 282.716492 |
| Arhgef2 | -0.7040637 | 5.025E-07 | 1.2752E-05 | 1564.99547 | 960.704221 |
| Mrs2 | -0.7038754 | 2.1676E-07 | 6.0982E-06 | 603.644274 | 370.559312 |
| Eif2ak3 | -0.703776 | 0.00088079 | 0.00687381 | 1243.43232 | 763.239976 |
| Zfp217 | -0.7021238 | 0.00540943 | 0.02865512 | 286.82043 | 176.31724 |
| Fhod1 | -0.7020474 | 0.0099291 | 0.04510577 | 97.846896 | 60.3033832 |
| Pcnx | -0.7010217 | 0.00022179 | 0.00224773 | 981.50964 | 603.514615 |
| Rnf216 | -0.700524 | 6.3254E-06 | 0.00011561 | 1125.85293 | 692.627626 |
| 1700088E04F | -0.6997525 | 0.01123898 | 0.04918239 | 103.654469 | 63.7121631 |
| Ino80d | -0.6985354 | 0.00031707 | 0.00300151 | 301.546212 | 185.940024 |
| 4930599N23I | -0.6974225 | 8.025E-07 | 1.9445E-05 | 375.920054 | 231.8541 |
| Tsc22d3 | -0.6973438 | 9.3997E-08 | 2.9419E-06 | 14868.0835 | 9169.11941 |
| Ift172 | -0.6965064 | 0.00540738 | 0.02865433 | 313.140662 | 193.12198 |
| Utrn | -0.6963494 | 8.2335E-05 | 0.00100318 | 1129.8686 | 697.366066 |
| Dock2 | -0.6962324 | 0.0012982 | 0.00938993 | 7437.46639 | 4590.20859 |
| mt-Rnr1 | -0.695219 | 0.00642981 | 0.03268866 | 998.704819 | 616.782438 |
| Wasf2 | -0.6948607 | 1.7084E-07 | 5.0216E-06 | 1823.85531 | 1126.90312 |
| Ap2a1 | -0.6948065 | 0.00040837 | 0.00366921 | 789.01453 | 487.193735 |
| Snap29 | -0.6947652 | 1.3558E-08 | 5.3182E-07 | 646.59477 | 399.244586 |
| Slc4a2 | -0.6945849 | 0.00011276 | 0.00130076 | 703.86099 | 434.758006 |
| Fem1b | -0.6944593 | 7.9409E-06 | 0.00014018 | 335.143413 | 206.990554 |
| Dicer1 | -0.6943095 | 0.00013852 | 0.00152341 | 410.24499 | 253.688595 |
| Lrrc20 | -0.6936845 | 0.00198672 | 0.01314826 | 198.869767 | 123.025978 |
| Supt6 | -0.6936253 | 0.00016463 | 0.00174784 | 718.405172 | 444.108396 |
| Clvs1 | -0.6935191 | 0.00263418 | 0.01644776 | 123.589999 | 76.3566393 |
| Afap1 | -0.6934978 | 0.00519641 | 0.02779125 | 94.5092608 | 58.3804049 |
| Pwwp2b | -0.693236 | 0.00150509 | 0.0105915 | 203.208414 | 125.653181 |
| Nucks1 | -0.6930351 | 0.00070885 | 0.0057656 | 356.176806 | 220.260265 |
| Ddi2 | -0.6928703 | 5.0214E-05 | 0.00066114 | 1414.72037 | 874.919618 |
| Syk | -0.6926602 | 2.7193E-05 | 0.00039281 | 7878.07293 | 4874.06051 |
| Kansl3 | -0.6923827 | 4.238E-05 | 0.00056999 | 816.974711 | 505.628891 |
| Rpl37a | -0.692116 | 0.00040824 | 0.00366921 | 7728.95193 | 4783.6458 |
| Gm15336 | -0.6920219 | 0.00305043 | 0.01841908 | 69.5165874 | 43.0631066 |
| Crybg1 | -0.6917815 | 0.00047617 | 0.00415977 | 1445.65625 | 894.862755 |
| Csnk1g1 | -0.6915829 | 0.00022409 | 0.00226698 | 297.162315 | 184.143768 |
| B4galt5 | -0.6910542 | 0.00017695 | 0.00185641 | 1885.7407 | 1167.85004 |
| Icmt | -0.6910541 | 0.00017663 | 0.00185434 | 593.642383 | 367.545755 |
| Dcaf15 | -0.6908631 | 0.00159374 | 0.01108462 | 212.462037 | 131.678926 |
| Slc41a1 | -0.6907078 | 0.00037043 | 0.0034026 | 492.265126 | 304.795528 |
| Ankrd13c | -0.6904739 | 3.451E-05 | 0.00048137 | 735.259457 | 455.415699 |
| Ap2b1 | -0.6881683 | 1.1582E-07 | 3.5438E-06 | 814.680365 | 505.727604 |
| Nsd1 | -0.6881091 | 1.1978E-05 | 0.00019817 | 873.129402 | 542.076795 |

| | | | | | |
|---|---|---|---|---|---|
| Osbp | -0.6878428 | 4.6593E-05 | 0.00061726 | 1583.61502 | 982.877811 |
| Tsc22d2 | -0.6877816 | 6.9377E-05 | 0.00086927 | 928.927483 | 576.568268 |
| Dpp8 | -0.6870302 | 9.2153E-05 | 0.00109874 | 1131.06255 | 702.516917 |
| Ralgps1 | -0.6860247 | 0.00884422 | 0.04139256 | 108.777965 | 67.5911593 |
| Fam168b | -0.6846908 | 4.3837E-05 | 0.00058748 | 1745.57853 | 1085.90697 |
| Slc45a4 | -0.6838379 | 0.00045736 | 0.00403936 | 463.64205 | 288.452932 |
| Tjp2 | -0.683496 | 0.00109842 | 0.00824218 | 267.109605 | 166.225882 |
| Elk1 | -0.682116 | 0.00630075 | 0.03217434 | 135.332748 | 84.3244475 |
| Chpf2 | -0.6804705 | 3.6441E-05 | 0.00050372 | 1046.46543 | 652.812503 |
| 4932438A13I | -0.6803283 | 2.0113E-05 | 0.00030552 | 673.47017 | 420.314374 |
| Tbc1d14 | -0.6801762 | 8.8278E-07 | 2.0951E-05 | 1724.50271 | 1076.03328 |
| Top2b | -0.6799691 | 0.00021469 | 0.00218961 | 1937.10675 | 1208.90704 |
| Mepce | -0.6795321 | 4.6393E-05 | 0.00061514 | 274.157195 | 171.000668 |
| Trpm7 | -0.6791099 | 3.3102E-08 | 1.1701E-06 | 805.023692 | 502.923095 |
| Nol6 | -0.6786569 | 0.00550223 | 0.02898016 | 685.410903 | 427.990854 |
| P2ry6 | -0.6786019 | 1.5148E-08 | 5.8502E-07 | 1652.66614 | 1032.39832 |
| Exoc8 | -0.6775098 | 2.0263E-05 | 0.00030686 | 737.940792 | 461.357345 |
| Edc4 | -0.6774136 | 0.00204104 | 0.01343521 | 1197.13008 | 748.40367 |
| Sp1 | -0.6772894 | 0.00045122 | 0.00399447 | 475.550003 | 297.405458 |
| Syne3 | -0.6763629 | 0.00275111 | 0.01700822 | 290.778145 | 181.838015 |
| Eif4g3 | -0.6762265 | 0.00016331 | 0.00173757 | 2062.69597 | 1290.69407 |
| Usf3 | -0.6758798 | 0.00211333 | 0.01384839 | 208.67828 | 130.513123 |
| Ip6k1 | -0.6754462 | 1.5494E-05 | 0.00024377 | 2204.62703 | 1380.34072 |
| Casc3 | -0.6753355 | 4.1092E-05 | 0.00055515 | 1179.78915 | 738.731601 |
| Pld1 | -0.6751116 | 0.00065632 | 0.00541755 | 319.588463 | 200.105997 |
| Raph1 | -0.6748016 | 0.00275585 | 0.01701653 | 249.220686 | 156.180386 |
| Ptbp3 | -0.6746315 | 3.0174E-06 | 6.1496E-05 | 2845.12123 | 1782.18049 |
| Wdr37 | -0.6742902 | 8.4759E-07 | 2.0321E-05 | 687.524268 | 430.569778 |
| Rassf4 | -0.6740096 | 3.7499E-09 | 1.7123E-07 | 14450.9509 | 9057.10701 |
| Atr | -0.6739761 | 0.00279497 | 0.01723237 | 140.403876 | 88.0992506 |
| Dhx33 | -0.673826 | 7.5905E-05 | 0.00093393 | 331.394255 | 207.608161 |
| Ncoa1 | -0.6738213 | 0.00106834 | 0.00806109 | 1820.86058 | 1141.26587 |
| Osbpl8 | -0.6732636 | 9.1834E-07 | 2.159E-05 | 891.635339 | 559.050354 |
| Brd2 | -0.6731191 | 7.5578E-07 | 1.8343E-05 | 2993.34309 | 1877.10326 |
| Gm14455 | -0.6722174 | 0.0031998 | 0.01906808 | 85.916606 | 53.9825967 |
| Arrb2 | -0.6718045 | 0.0003783 | 0.00345693 | 1642.85518 | 1031.22715 |
| Nemp1 | -0.6709871 | 0.01121713 | 0.04910108 | 116.306626 | 72.9390003 |
| Cdk19 | -0.6690481 | 1.6522E-06 | 3.6181E-05 | 738.661997 | 464.43719 |
| Mafg | -0.6680609 | 7.0228E-09 | 3.023E-07 | 1194.53675 | 751.928928 |
| Slc9a1 | -0.6675617 | 0.00250016 | 0.01576458 | 520.821741 | 327.735149 |
| Cep350 | -0.6670825 | 2.7168E-07 | 7.394E-06 | 739.301924 | 465.668583 |
| Map3k1 | -0.6665067 | 1.2743E-06 | 2.8745E-05 | 1063.60395 | 670.25662 |
| Rlim | -0.66626 | 8.2733E-06 | 0.00014486 | 603.781123 | 380.354348 |
| Fxr2 | -0.6650766 | 7.1755E-05 | 0.00089162 | 768.856768 | 484.936572 |
| Taok1 | -0.6647363 | 0.00026078 | 0.00256561 | 523.476266 | 330.110578 |
| Ipo8 | -0.6646677 | 0.00045815 | 0.00404201 | 413.124987 | 260.519496 |

| | | | | | |
|---|---|---|---|---|---|
| Cul9 | -0.6645612 | 0.00238443 | 0.01519139 | 151.953228 | 95.7513651 |
| Ap3d1 | -0.6639399 | 5.9547E-05 | 0.00076459 | 2316.38625 | 1461.92006 |
| Akna | -0.6636944 | 0.00071294 | 0.00579258 | 7968.21275 | 5029.86197 |
| Nup160 | -0.6635514 | 0.00017162 | 0.00181062 | 527.10141 | 332.493982 |
| Camta2 | -0.6630186 | 0.00180225 | 0.01222912 | 1817.1453 | 1147.56945 |
| Atxn2 | -0.6629903 | 1.0696E-05 | 0.00017974 | 856.098095 | 540.712327 |
| Ankrd44 | -0.6625956 | 0.0005068 | 0.00437822 | 2454.99977 | 1550.84821 |
| Prkd3 | -0.6623138 | 2.9374E-05 | 0.00041829 | 951.858279 | 601.122328 |
| Gm9843 | -0.6621046 | 0.00393289 | 0.02232106 | 439.229001 | 277.478001 |
| Ids | -0.6614993 | 1.0361E-05 | 0.0001747 | 2097.99261 | 1326.10105 |
| Zfp292 | -0.6614751 | 0.00037887 | 0.00345789 | 489.781022 | 309.718241 |
| Tef | -0.6605838 | 2.7774E-05 | 0.000399 | 1144.71237 | 724.011789 |
| Ypel2 | -0.6603982 | 9.215E-05 | 0.00109874 | 175.762267 | 111.303031 |
| Kdm3a | -0.6598928 | 0.00065669 | 0.00541764 | 814.716616 | 515.680283 |
| Xpo6 | -0.6590955 | 0.00012769 | 0.00142944 | 1837.83433 | 1163.60788 |
| Tyk2 | -0.658704 | 0.00020977 | 0.00214964 | 3548.93094 | 2247.90132 |
| Dvl1 | -0.6583173 | 9.7817E-05 | 0.001158 | 699.856002 | 443.571034 |
| Usp20 | -0.6583165 | 0.00657485 | 0.03324529 | 171.127456 | 108.534682 |
| Ncoa3 | -0.6577205 | 0.00184179 | 0.01243427 | 1630.84362 | 1033.72173 |
| Creb1 | -0.6576873 | 1.4468E-05 | 0.00023098 | 1107.81482 | 702.147283 |
| Zbtb44 | -0.6575927 | 0.00017325 | 0.00182391 | 630.866247 | 399.777556 |
| Gm38399 | -0.6573784 | 0.0066984 | 0.03371946 | 102.247011 | 64.9231299 |
| Acap3 | -0.6573115 | 0.00010898 | 0.00126719 | 778.128071 | 493.320116 |
| Tmtc3 | -0.6565203 | 0.00317431 | 0.01895374 | 121.656439 | 77.1301946 |
| Xpo4 | -0.6562001 | 0.00172261 | 0.01182759 | 217.708932 | 138.366904 |
| Scaf8 | -0.6557178 | 0.00039062 | 0.00354367 | 602.025155 | 382.007313 |
| Psd | -0.6547772 | 0.00222009 | 0.01434512 | 142.727383 | 90.6928001 |
| Gak | -0.654225 | 0.0007301 | 0.00588744 | 2214.46035 | 1406.84347 |
| Zfp12 | -0.6537873 | 0.00427756 | 0.02382092 | 379.298343 | 240.969402 |
| Ccdc9 | -0.6537479 | 0.00090503 | 0.00702652 | 437.972566 | 278.33633 |
| AW549877 | -0.6527055 | 5.3366E-05 | 0.00069654 | 649.644507 | 413.325057 |
| A430078G23I | -0.6524888 | 0.00799316 | 0.03843829 | 533.386573 | 339.486599 |
| Tbc1d2 | -0.6515054 | 0.00491079 | 0.02654726 | 250.068414 | 159.184987 |
| Slc39a9 | -0.6511797 | 0.0001463 | 0.00159126 | 783.115517 | 498.350011 |
| Inpp5d | -0.6509445 | 0.00102171 | 0.00775993 | 4871.90752 | 3102.55794 |
| Slc25a27 | -0.6506966 | 0.00750619 | 0.03670751 | 179.220929 | 114.289202 |
| Iqce | -0.650655 | 0.00011191 | 0.00129523 | 296.845774 | 188.944453 |
| Ppm1k | -0.6498416 | 0.0012656 | 0.00919393 | 149.844447 | 95.5452502 |
| Pdpk1 | -0.649405 | 4.8898E-05 | 0.00064487 | 646.052318 | 411.815733 |
| Trim33 | -0.6493767 | 0.00847574 | 0.0400598 | 213.267938 | 135.962855 |
| Lrig2 | -0.6491318 | 0.00024204 | 0.00241763 | 252.477329 | 160.907445 |
| Smg7 | -0.6481278 | 0.00029255 | 0.00282423 | 476.79197 | 304.315455 |
| Epm2aip1 | -0.6480556 | 0.00301336 | 0.01822452 | 208.535658 | 132.986792 |
| Slc35e2 | -0.6479749 | 0.00105543 | 0.00799176 | 235.570365 | 150.548899 |
| Strn | -0.6478138 | 4.5027E-05 | 0.00060022 | 439.799333 | 280.41688 |
| Osbpl11 | -0.6477147 | 0.00030078 | 0.00289285 | 964.441014 | 615.38881 |

| | | | | | |
|---|---|---|---|---|---|
| Trim28 | -0.6473868 | 1.9791E-07 | 5.6418E-06 | 640.192838 | 408.790396 |
| Rttn | -0.6471253 | 0.003781 | 0.02168843 | 90.1924723 | 57.5776307 |
| Spen | -0.6468046 | 0.00064091 | 0.00531956 | 654.506397 | 417.956928 |
| Ubr2 | -0.6465359 | 0.00165533 | 0.01142835 | 894.924764 | 571.744246 |
| Tlr8 | -0.646384 | 0.00015301 | 0.00165131 | 1155.3483 | 737.797423 |
| Pip5k1c | -0.6459623 | 0.00110645 | 0.00828626 | 2415.79585 | 1543.67784 |
| Pitpnm1 | -0.6455887 | 0.00125463 | 0.00913634 | 3479.20075 | 2223.89989 |
| Setd2 | -0.6454464 | 4.7631E-06 | 9.1179E-05 | 712.702639 | 455.603063 |
| Nfrkb | -0.645354 | 0.00983389 | 0.0447816 | 291.206069 | 186.172557 |
| Lrch3 | -0.6442986 | 0.00087904 | 0.00686374 | 744.429442 | 476.253348 |
| Senp5 | -0.6442914 | 0.00011745 | 0.00134175 | 297.07177 | 190.104045 |
| Traf6 | -0.6433747 | 0.00206505 | 0.01356506 | 770.192046 | 492.95243 |
| I830127L07Ri | -0.6432252 | 0.00497 | 0.02681422 | 3966.46427 | 2539.5254 |
| Ctdsp1 | -0.6424145 | 0.00710789 | 0.03529976 | 349.270528 | 223.816828 |
| Tmem65 | -0.6415312 | 8.1752E-07 | 1.9745E-05 | 694.193306 | 444.771297 |
| Gm11973 | -0.6412234 | 4.5266E-05 | 0.00060232 | 380.587569 | 244.074216 |
| Gm9725 | -0.6407388 | 0.00082224 | 0.00648428 | 219.809839 | 140.822207 |
| Tmsb10 | -0.6404124 | 4.2255E-05 | 0.00056881 | 10357.0179 | 6644.13319 |
| Ptbp2 | -0.6401965 | 0.00070448 | 0.00573626 | 145.328066 | 93.3699747 |
| Sp2 | -0.63951 | 0.00038066 | 0.00346791 | 704.127042 | 451.845851 |
| Olfm1 | -0.6394801 | 0.00033603 | 0.00312967 | 3448.47087 | 2213.67297 |
| Slfn5 | -0.6394738 | 0.00030829 | 0.00294244 | 3840.69461 | 2465.22982 |
| Pcm1 | -0.6391299 | 0.00019274 | 0.00200241 | 170.961421 | 109.843774 |
| Tob1 | -0.6386969 | 2.6527E-05 | 0.00038504 | 249.78553 | 160.314466 |
| Zfp318 | -0.6385914 | 0.0076483 | 0.03726904 | 93.2663565 | 59.8685548 |
| Alpk1 | -0.6383082 | 7.3205E-06 | 0.00013077 | 377.303746 | 242.573333 |
| Lig3 | -0.6374974 | 0.00518594 | 0.02775503 | 443.203918 | 284.872577 |
| Arpc3 | -0.6372091 | 0.01080778 | 0.04794359 | 5142.02312 | 3305.95674 |
| Rprd2 | -0.6362445 | 0.00069012 | 0.00563763 | 279.844917 | 180.217413 |
| Kdm6a | -0.6358646 | 0.00046035 | 0.00405387 | 487.325668 | 313.448618 |
| Slc7a1 | -0.6349006 | 0.00964534 | 0.04405323 | 176.955047 | 114.022006 |
| Gna13 | -0.6348907 | 8.6674E-07 | 2.07E-05 | 2268.44052 | 1460.62259 |
| Map3k5 | -0.6348083 | 0.00013928 | 0.00153065 | 1678.78728 | 1081.09137 |
| Clk2 | -0.6343513 | 2.1284E-06 | 4.5566E-05 | 695.665907 | 448.256677 |
| Gm9844 | -0.6341035 | 0.00027543 | 0.00268501 | 720.063483 | 463.808802 |
| Frmd4a | -0.6336283 | 0.00016762 | 0.00177588 | 904.787972 | 582.956829 |
| Txnl4b | -0.6336176 | 0.00687789 | 0.03440182 | 198.900744 | 128.171513 |
| Itgav | -0.6324678 | 0.0002735 | 0.00267149 | 911.68494 | 588.119527 |
| Mtss1 | -0.6322792 | 0.001795 | 0.01219092 | 176.685203 | 113.933249 |
| Herc2 | -0.6320788 | 0.00016919 | 0.00178868 | 620.131705 | 400.153739 |
| N4bp1 | -0.6316124 | 1.5369E-06 | 3.3957E-05 | 1319.01594 | 851.214303 |
| Cblb | -0.631502 | 0.00462826 | 0.02525624 | 913.606183 | 589.721578 |
| Pprc1 | -0.6290874 | 0.00580559 | 0.03028865 | 537.213332 | 347.184574 |
| Agl | -0.6290799 | 0.00051314 | 0.00443044 | 324.663387 | 209.880441 |
| Arid1a | -0.6280735 | 0.0016152 | 0.01120274 | 950.27762 | 614.902713 |
| Arvcf | -0.6277061 | 0.00609546 | 0.03142486 | 96.6097061 | 62.4280884 |

| | | | | | |
|---|---|---|---|---|---|
| Gm12359 | -0.6273334 | 0.00032702 | 0.00306856 | 198.446642 | 128.538819 |
| Dctn1 | -0.6266464 | 0.00250063 | 0.01576458 | 1841.59778 | 1192.63293 |
| Rnf144a | -0.6256835 | 0.00180774 | 0.0122553 | 573.31143 | 371.547056 |
| Hk2 | -0.6254314 | 0.00125744 | 0.00915237 | 798.48776 | 517.727602 |
| Dync1i2 | -0.624565 | 1.9513E-05 | 0.0002976 | 608.668667 | 394.659377 |
| Ybx1 | -0.6234124 | 6.0365E-09 | 2.6594E-07 | 4461.93297 | 2896.38864 |
| Xiap | -0.6228755 | 0.00549722 | 0.02897681 | 453.203633 | 294.322573 |
| St8sia4 | -0.6228199 | 7.3128E-06 | 0.00013077 | 2890.465 | 1876.76805 |
| Exoc6b | -0.6213207 | 0.00028463 | 0.00275874 | 410.184956 | 266.528082 |
| Fam102b | -0.6210821 | 1.2782E-07 | 3.8715E-06 | 2171.4017 | 1411.46148 |
| Cdh23 | -0.6204464 | 0.0080875 | 0.03870612 | 577.181617 | 375.533647 |
| Mark2 | -0.6201372 | 6.3153E-05 | 0.00079928 | 1700.20847 | 1106.00012 |
| Snx25 | -0.6198447 | 0.01028235 | 0.04618459 | 265.929777 | 172.786461 |
| Tusc1 | -0.619635 | 1.3195E-05 | 0.00021477 | 345.872173 | 224.918749 |
| Fam208b | -0.6191315 | 0.00573974 | 0.03003417 | 255.954828 | 166.83873 |
| Mbp | -0.6191066 | 0.00122842 | 0.0089803 | 4813.03362 | 3133.3842 |
| Ints6 | -0.6187678 | 3.0911E-06 | 6.2332E-05 | 784.126161 | 510.531885 |
| Ago2 | -0.6182559 | 7.025E-05 | 0.00087801 | 1029.29277 | 670.444474 |
| Zfp282 | -0.6173497 | 0.00077175 | 0.00614411 | 373.040657 | 243.08189 |
| Prpf38b | -0.6172269 | 8.4602E-06 | 0.00014745 | 1108.56618 | 722.955704 |
| Dennd1a | -0.6170742 | 0.0002303 | 0.00232046 | 928.045808 | 604.872365 |
| Rasgrp4 | -0.6168794 | 1.929E-06 | 4.1637E-05 | 2135.12209 | 1392.38919 |
| Akap11 | -0.6156126 | 0.003163 | 0.01893288 | 127.305929 | 83.226323 |
| Gxylt1 | -0.6156116 | 1.1105E-05 | 0.00018599 | 498.234344 | 325.10712 |
| Syvn1 | -0.6146354 | 2.9815E-05 | 0.00042255 | 1104.55421 | 721.241712 |
| Epb41 | -0.6135791 | 0.00352503 | 0.02051785 | 994.684474 | 649.929323 |
| Usp30 | -0.6132997 | 0.00058447 | 0.00493566 | 159.514824 | 104.231806 |
| Pogz | -0.6130122 | 0.0036976 | 0.02134037 | 154.553555 | 100.944589 |
| Lrrk1 | -0.6128757 | 0.00114205 | 0.00848518 | 1857.62946 | 1214.84442 |
| Nfkbiz | -0.6127502 | 0.00954953 | 0.04372525 | 584.538942 | 382.415317 |
| Csrp1 | -0.6127459 | 0.00027407 | 0.00267536 | 367.654599 | 240.260897 |
| Itch | -0.6121676 | 0.00034911 | 0.00323741 | 3036.00095 | 1986.06568 |
| Gsk3b | -0.6121408 | 0.00037508 | 0.00343585 | 669.654754 | 438.130606 |
| Hlcs | -0.6119306 | 8.424E-05 | 0.00101896 | 713.630187 | 466.785728 |
| Mkl2 | -0.6115374 | 0.00722176 | 0.03573517 | 114.336081 | 74.7456588 |
| Polg | -0.6113966 | 0.00040268 | 0.00363112 | 1494.49114 | 978.048384 |
| Rnf38 | -0.6109219 | 0.00026041 | 0.00256364 | 886.852138 | 580.395899 |
| Hmox1 | -0.6093358 | 0.00115553 | 0.00854727 | 3352.5167 | 2197.48241 |
| Tnrc6c | -0.609334 | 0.00030602 | 0.00292641 | 444.580966 | 291.607805 |
| Cd24a | -0.608441 | 0.01050275 | 0.04690332 | 537.478863 | 352.411167 |
| Grk2 | -0.6078464 | 3.6447E-05 | 0.00050372 | 6770.81651 | 4442.62394 |
| Rnf213 | -0.6068088 | 0.0077674 | 0.03769046 | 1555.51136 | 1021.47712 |
| Gm11769 | -0.6067574 | 0.00366849 | 0.02122133 | 238.322052 | 156.425043 |
| Hps5 | -0.6062548 | 0.0008976 | 0.0069869 | 1296.7585 | 851.647697 |
| Gm10076 | -0.6045937 | 0.00570174 | 0.02986655 | 254.693749 | 167.471622 |
| Atp8a1 | -0.6044825 | 3.1657E-05 | 0.00044489 | 1092.91006 | 718.701842 |

| | | | | | |
|---|---|---|---|---|---|
| Ei24 | -0.6044737 | 1.574E-06 | 3.4673E-05 | 1049.71911 | 690.168094 |
| Naip2 | -0.604065 | 8.0877E-05 | 0.00098782 | 1098.59234 | 722.562749 |
| Mxd1 | -0.6037782 | 0.00015373 | 0.00165793 | 783.9019 | 515.854171 |
| Stk11ip | -0.6036152 | 0.00058698 | 0.00495126 | 940.641565 | 618.805617 |
| Ogfod1 | -0.6024889 | 0.00706066 | 0.03511165 | 138.094012 | 91.0684176 |
| Reep5 | -0.6023904 | 1.8568E-05 | 0.00028378 | 2960.53421 | 1949.74505 |
| Ehmt2 | -0.6015396 | 1.5585E-05 | 0.00024466 | 1045.88279 | 689.343798 |
| Gins2 | -0.6014674 | 0.00624983 | 0.03200109 | 180.781466 | 119.219108 |
| Gm37844 | -0.6003238 | 0.00790445 | 0.03816128 | 103.233735 | 68.1239805 |
| Castor2 | -0.5999326 | 0.00032084 | 0.00302006 | 275.905674 | 182.052896 |
| Phf14 | -0.5999299 | 0.01011134 | 0.04572625 | 364.908073 | 240.801033 |
| Tbc1d10b | -0.5998008 | 0.00322508 | 0.01916552 | 2517.35871 | 1660.87652 |
| Pds5b | -0.5993255 | 0.00412725 | 0.02313569 | 360.157922 | 237.745985 |
| Pxn | -0.5980471 | 0.00206027 | 0.01353959 | 3254.70994 | 2149.98622 |
| Rcan3 | -0.5963649 | 0.00383038 | 0.02187148 | 292.245809 | 193.131012 |
| Slc35a3 | -0.5961109 | 0.00010371 | 0.00121436 | 502.719797 | 332.400671 |
| Trib1 | -0.5957179 | 0.00427332 | 0.02381236 | 385.038449 | 254.744714 |
| Arfgef1 | -0.5956131 | 7.4785E-05 | 0.000921 | 1036.45496 | 685.557797 |
| C2cd3 | -0.5953699 | 0.00041108 | 0.00368478 | 540.304175 | 357.481543 |
| Yod1 | -0.5950989 | 0.00440584 | 0.02437648 | 178.345561 | 118.110045 |
| AI837181 | -0.5950708 | 0.00378575 | 0.02170737 | 488.086673 | 323.058989 |
| Tnpo1 | -0.5944943 | 2.2436E-06 | 4.7473E-05 | 584.806442 | 387.374581 |
| Pde4dip | -0.5942942 | 0.00782323 | 0.03786484 | 716.399622 | 474.566732 |
| Myo9a | -0.5940622 | 0.00174312 | 0.01192472 | 232.99883 | 154.413309 |
| Erbin | -0.5932382 | 0.00218728 | 0.01418192 | 1080.6838 | 716.273111 |
| Phka2 | -0.59283 | 0.00030331 | 0.0029123 | 733.868587 | 486.45204 |
| Mllt3 | -0.5925548 | 0.00753582 | 0.03680451 | 134.005815 | 88.9754818 |
| Pabpc1 | -0.5919438 | 7.0731E-05 | 0.00088181 | 3874.55532 | 2570.49705 |
| Pik3r5 | -0.5915903 | 4.0784E-05 | 0.00055345 | 3570.00766 | 2368.76235 |
| Zdhhc5 | -0.5908305 | 0.00012008 | 0.00136479 | 1324.58654 | 879.162677 |
| 4931414P19F | -0.5905762 | 0.01084779 | 0.04799565 | 84.261969 | 55.89705 |
| Rai1 | -0.5904555 | 0.00514566 | 0.02760832 | 199.167893 | 132.263872 |
| Gm16192 | -0.5903408 | 0.00166214 | 0.0114648 | 280.99447 | 186.530279 |
| Sec24c | -0.5899975 | 0.00041736 | 0.0037344 | 3174.33475 | 2108.67943 |
| Dennd1b | -0.5894467 | 0.00015531 | 0.0016702 | 463.625724 | 307.944919 |
| Msl1 | -0.589155 | 0.00026939 | 0.00263825 | 577.497267 | 383.965997 |
| Trpm2 | -0.5887616 | 0.00884445 | 0.04139256 | 228.294404 | 151.692925 |
| Ap1g1 | -0.5886568 | 0.00037636 | 0.00344132 | 1145.23907 | 761.310315 |
| Nktr | -0.5883252 | 0.00429023 | 0.02384464 | 909.417011 | 605.130114 |
| Arrdc3 | -0.5878207 | 3.0533E-06 | 6.1735E-05 | 1450.27524 | 964.820366 |
| Zfp653 | -0.5867597 | 0.00911508 | 0.04229023 | 144.086471 | 95.9664849 |
| Ctdnep1 | -0.5855562 | 0.00033969 | 0.00315989 | 340.171136 | 226.752623 |
| Cux1 | -0.5854558 | 1.3029E-06 | 2.9258E-05 | 1823.21336 | 1214.81796 |
| Ubn2 | -0.5849351 | 0.00057232 | 0.00484665 | 749.664187 | 499.935865 |
| Phf3 | -0.5848595 | 0.00144757 | 0.01025308 | 669.288627 | 446.192284 |
| Srrt | -0.5847259 | 0.00014175 | 0.00155096 | 722.633371 | 481.93392 |

| | | | | | |
|---|---|---|---|---|---|
| Zc3h18 | -0.5843236 | 0.00114507 | 0.00849501 | 727.248124 | 484.961837 |
| Sh2b1 | -0.5840141 | 3.0971E-05 | 0.00043721 | 484.158589 | 322.775906 |
| Pcgf3 | -0.5836927 | 9.1967E-05 | 0.00109827 | 285.681868 | 190.53873 |
| Ttpal | -0.5836238 | 0.00024594 | 0.00244526 | 632.732073 | 422.092438 |
| Uhrf1bp1l | -0.5829129 | 6.8896E-06 | 0.00012396 | 1154.43048 | 770.610445 |
| Pdxk | -0.5824078 | 8.4087E-06 | 0.00014706 | 1420.03681 | 948.319115 |
| Ccdc50 | -0.5815563 | 8.3729E-05 | 0.00101453 | 2025.99349 | 1353.69643 |
| mt-Co1 | -0.581441 | 0.00290953 | 0.0177832 | 15997.9959 | 10691.2095 |
| Itga4 | -0.5813787 | 1.771E-07 | 5.1662E-06 | 4459.76143 | 2980.42546 |
| Ifih1 | -0.5807469 | 0.00148384 | 0.01046562 | 584.109875 | 390.433236 |
| Bace1 | -0.5796725 | 0.00146642 | 0.01037684 | 266.492446 | 178.079414 |
| Kdm5b | -0.578857 | 0.00606007 | 0.03127463 | 718.425862 | 480.988959 |
| Cd2ap | -0.5786002 | 1.3327E-05 | 0.00021622 | 676.619016 | 453.040899 |
| Fyco1 | -0.5779786 | 8.2563E-05 | 0.00100433 | 454.23885 | 304.438543 |
| Dusp1 | -0.5779284 | 0.00569039 | 0.02981749 | 1794.3556 | 1201.9452 |
| Esrra | -0.5778369 | 0.00308015 | 0.01856121 | 465.901153 | 312.188792 |
| Ago4 | -0.577293 | 0.00376213 | 0.02160796 | 141.193026 | 94.4976909 |
| Sdr39u1 | -0.5769311 | 0.00073169 | 0.00589711 | 317.637021 | 212.936572 |
| C530005A16F | -0.5761191 | 0.00919589 | 0.04254694 | 95.2250057 | 63.8303368 |
| Klf2 | -0.5760832 | 2.5616E-05 | 0.00037471 | 9230.48615 | 6191.32771 |
| Zeb2os | -0.5756085 | 2.1211E-05 | 0.00031866 | 527.767151 | 354.214641 |
| Acsl4 | -0.5745107 | 0.00046557 | 0.00408068 | 403.564111 | 270.830829 |
| Actr1a | -0.573667 | 9.154E-06 | 0.00015789 | 2263.28756 | 1520.34803 |
| Vps13b | -0.5725373 | 0.00024195 | 0.00241763 | 655.161393 | 440.726292 |
| Cdk13 | -0.5723661 | 2.6017E-05 | 0.00037874 | 1059.68089 | 712.531461 |
| Itsn1 | -0.5722818 | 0.00175907 | 0.0120066 | 215.79819 | 144.884508 |
| Heatr1 | -0.572254 | 3.881E-05 | 0.00053053 | 863.665148 | 580.777702 |
| Lrsam1 | -0.5717855 | 0.01067416 | 0.04755065 | 192.832913 | 129.556087 |
| Wrn | -0.5717267 | 1.6507E-05 | 0.00025669 | 441.998646 | 297.304345 |
| Larp4b | -0.5713161 | 0.00018332 | 0.00190717 | 1484.14661 | 998.653949 |
| Patz1 | -0.5710881 | 0.00304823 | 0.01841315 | 410.787482 | 276.524107 |
| Sfswap | -0.5705901 | 0.00020297 | 0.00208709 | 608.322466 | 409.623405 |
| Slc38a2 | -0.5695193 | 5.5944E-06 | 0.00010427 | 2586.21377 | 1742.44498 |
| Kmt5b | -0.5682082 | 8.3159E-05 | 0.00100995 | 749.577611 | 505.547889 |
| Scyl2 | -0.5681203 | 0.0064633 | 0.03279233 | 228.470476 | 153.850573 |
| Dgkz | -0.5679542 | 0.00187533 | 0.01260711 | 2074.35429 | 1399.13155 |
| Ticam1 | -0.5678698 | 0.00020724 | 0.00212667 | 331.559023 | 223.621921 |
| Fam13b | -0.5678264 | 0.00342635 | 0.02006785 | 588.692225 | 397.119609 |
| Slc16a3 | -0.567537 | 0.00158608 | 0.01105692 | 1442.62071 | 973.195547 |
| Plekho1 | -0.5674519 | 4.1267E-08 | 1.4252E-06 | 1688.4344 | 1139.33317 |
| Hsp90aa1 | -0.5670903 | 0.01092957 | 0.04828761 | 2883.69434 | 1946.4228 |
| Pgd | -0.5667631 | 5.3559E-05 | 0.00069846 | 5451.04923 | 3679.8218 |
| Zfp106 | -0.5666212 | 0.00054113 | 0.004619 | 1633.62904 | 1103.05749 |
| Csf2rb | -0.5663678 | 6.4002E-05 | 0.0008073 | 5718.69473 | 3861.53949 |
| Mob3b | -0.5659142 | 0.0001941 | 0.00201384 | 902.885308 | 609.659851 |
| Steap3 | -0.5650792 | 0.01132436 | 0.04948394 | 158.699597 | 107.049345 |

| | | | | | |
|---|---|---|---|---|---|
| Rbbp6 | -0.5650743 | 0.00024851 | 0.00246268 | 558.687601 | 377.446123 |
| Hmgcr | -0.5644245 | 0.00455464 | 0.02496329 | 1039.03099 | 702.514659 |
| Whamm | -0.5639363 | 0.00133451 | 0.00957878 | 429.310539 | 290.259168 |
| Zfp710 | -0.5636903 | 5.0752E-05 | 0.00066756 | 3096.54195 | 2094.75179 |
| Golga2 | -0.563404 | 8.7611E-05 | 0.00105463 | 577.251972 | 390.509058 |
| Iqgap2 | -0.5632789 | 0.00011282 | 0.00130076 | 1788.50863 | 1210.43315 |
| Zyg11b | -0.563096 | 0.00031272 | 0.00297136 | 275.883675 | 186.871595 |
| Cyb561d1 | -0.5627131 | 0.00082335 | 0.00648961 | 528.568287 | 357.520186 |
| Insig1 | -0.5614472 | 0.00222186 | 0.01435036 | 347.76959 | 235.684857 |
| Polr3a | -0.561338 | 0.00655073 | 0.03315754 | 443.837638 | 300.600581 |
| Dync1h1 | -0.5612255 | 0.00330529 | 0.01953396 | 1157.30402 | 784.343376 |
| Dnajc5 | -0.5609663 | 1.2125E-05 | 0.00019973 | 2111.29536 | 1430.94017 |
| Zbtb6 | -0.5604862 | 0.00493951 | 0.02668328 | 161.839588 | 109.771244 |
| Tbc1d1 | -0.5600615 | 0.00247792 | 0.01565416 | 1150.29351 | 780.12881 |
| Ece1 | -0.5596971 | 0.00105839 | 0.00800484 | 1154.24148 | 782.945562 |
| Nisch | -0.5595183 | 0.00115812 | 0.00856224 | 1774.25909 | 1204.11492 |
| Dyrk3 | -0.5591978 | 0.0009797 | 0.00747489 | 390.498532 | 264.974085 |
| Zfp385a | -0.5590827 | 2.5939E-05 | 0.00037833 | 899.405716 | 610.333075 |
| D430042O09 | -0.5589997 | 0.01021875 | 0.04602708 | 207.278775 | 140.849875 |
| Gm48236 | -0.5588487 | 0.00594365 | 0.0308008 | 274.709049 | 186.548691 |
| Gpsm3 | -0.5584016 | 3.7123E-06 | 7.3382E-05 | 671.760751 | 455.894393 |
| Ehmt1 | -0.558382 | 8.2899E-05 | 0.00100759 | 1039.68457 | 705.884394 |
| Zfp219 | -0.5579694 | 0.00964755 | 0.04405323 | 126.170553 | 85.713019 |
| Trim44 | -0.5573826 | 0.0031506 | 0.01889156 | 588.942417 | 400.348566 |
| mt-Nd6 | -0.5571725 | 0.00739394 | 0.03628853 | 1305.83904 | 887.342795 |
| Naa30 | -0.5557525 | 8.7874E-05 | 0.00105695 | 393.99324 | 268.066723 |
| Alg2 | -0.5551901 | 0.00339077 | 0.01992159 | 212.864752 | 144.799711 |
| Usp24 | -0.553767 | 0.00057162 | 0.00484345 | 380.69166 | 259.509109 |
| Gmip | -0.553414 | 0.00030529 | 0.00292641 | 2844.67646 | 1938.17644 |
| Slc35d1 | -0.5532492 | 0.00274705 | 0.01699707 | 215.17315 | 146.441472 |
| Sufu | -0.5529073 | 0.00034531 | 0.00320614 | 385.960527 | 263.039324 |
| Sbk1 | -0.5527624 | 0.00876063 | 0.04114126 | 1201.28407 | 818.731995 |
| D5Ertd579e | -0.552257 | 4.7168E-05 | 0.00062377 | 416.781897 | 284.251828 |
| Ppp1cb | -0.5519269 | 0.00010259 | 0.00120309 | 2358.89298 | 1608.72712 |
| Smchd1 | -0.5515367 | 0.00037621 | 0.00344132 | 1145.68245 | 781.791893 |
| Smarcd3 | -0.5513559 | 6.6489E-05 | 0.00083587 | 579.569389 | 395.354305 |
| Zfp658 | -0.551292 | 0.01093165 | 0.04828761 | 200.035625 | 136.462605 |
| Ubr5 | -0.550772 | 0.00035562 | 0.00328165 | 426.939652 | 291.429971 |
| Spred1 | -0.5503586 | 0.00037508 | 0.00343585 | 528.123181 | 360.491157 |
| Gga1 | -0.5503458 | 0.00077089 | 0.00614056 | 1653.60555 | 1128.93565 |
| Gm10353 | -0.5500212 | 0.00067383 | 0.0055317 | 248.720634 | 169.75848 |
| Lrrc58 | -0.5493304 | 3.4901E-06 | 6.9445E-05 | 804.952679 | 549.820871 |
| Foxk1 | -0.5488046 | 0.00418879 | 0.02341077 | 265.190434 | 181.358115 |
| Ube4a | -0.5480422 | 0.00074273 | 0.00596049 | 588.690825 | 402.454824 |
| Fndc3a | -0.5480095 | 2.3439E-06 | 4.9248E-05 | 1299.67598 | 889.019774 |
| Armh3 | -0.5477494 | 0.00040312 | 0.00363288 | 268.768076 | 184.003017 |

| | | | | | |
|---|---|---|---|---|---|
| Tmem104 | -0.5477464 | 0.0001519 | 0.00164054 | 1372.57507 | 938.767842 |
| Wnk1 | -0.5474343 | 5.3841E-05 | 0.0006997 | 1018.8281 | 697.183844 |
| Nomo1 | -0.5473246 | 0.00151298 | 0.01063119 | 558.89609 | 382.462026 |
| Sbno2 | -0.5471918 | 8.3236E-05 | 0.00101006 | 5129.99221 | 3510.4265 |
| B230354K17F | -0.5471165 | 0.00207853 | 0.01364761 | 152.504202 | 104.285919 |
| Atad2b | -0.547044 | 6.1036E-05 | 0.00077971 | 798.352442 | 546.609859 |
| Senp7 | -0.5468912 | 0.00014875 | 0.00161346 | 371.118467 | 254.020742 |
| Stag1 | -0.5450585 | 0.00076896 | 0.0061307 | 792.736678 | 543.370559 |
| Zfp715 | -0.5449622 | 0.00072854 | 0.00588431 | 348.765976 | 238.914744 |
| Arid3b | -0.5448155 | 0.00061934 | 0.00517198 | 298.059136 | 204.421208 |
| Spin1 | -0.544787 | 0.00014638 | 0.00159126 | 622.11504 | 426.3533 |
| Hdac4 | -0.5445514 | 0.0023493 | 0.01502489 | 496.412195 | 340.232315 |
| Srebf2 | -0.5444219 | 0.00771348 | 0.03748937 | 2227.55189 | 1527.14372 |
| Mex3c | -0.5442895 | 0.00186497 | 0.01255265 | 547.392139 | 375.171294 |
| Tent4b | -0.5442579 | 0.00239368 | 0.01524383 | 379.019087 | 259.764671 |
| P2rx7 | -0.5434363 | 0.0001805 | 0.00188305 | 1028.10035 | 705.211822 |
| Fmn1 | -0.5432129 | 0.01113178 | 0.04884132 | 105.613662 | 72.6052476 |
| Sin3a | -0.5427098 | 0.00238268 | 0.01518669 | 592.16506 | 406.462594 |
| Chrnb2 | -0.5425675 | 0.00059876 | 0.00502804 | 498.670386 | 342.14084 |
| B4galt1 | -0.5424896 | 7.0486E-05 | 0.00087949 | 1031.87021 | 708.273378 |
| Acap2 | -0.5424328 | 0.00115264 | 0.0085394 | 1370.99093 | 941.109303 |
| Elk4 | -0.5417511 | 0.00427875 | 0.02382092 | 271.186586 | 186.330727 |
| Gm37494 | -0.5411223 | 0.00742943 | 0.03641371 | 369.156828 | 253.771669 |
| Det1 | -0.5409116 | 0.00442019 | 0.02442247 | 235.593905 | 161.782106 |
| Npc1 | -0.5405591 | 0.00381551 | 0.02180387 | 743.200602 | 510.901563 |
| Entpd7 | -0.5399472 | 0.00282351 | 0.01736288 | 323.448201 | 222.199203 |
| Cep295 | -0.5396108 | 0.00275531 | 0.01701653 | 173.547618 | 119.567079 |
| Amd1 | -0.5395843 | 7.0924E-05 | 0.00088349 | 477.239914 | 328.483776 |
| Xylt1 | -0.5394858 | 0.00024636 | 0.00244784 | 734.691439 | 505.628227 |
| Brd1 | -0.5394143 | 0.00035124 | 0.00325121 | 1022.64688 | 703.581236 |
| Llgl1 | -0.5392877 | 0.0042028 | 0.02345416 | 869.457958 | 598.127858 |
| C2cd5 | -0.5392683 | 0.00084094 | 0.00660745 | 846.179529 | 582.163873 |
| Asap1 | -0.5387925 | 3.4426E-05 | 0.00048065 | 3730.02854 | 2567.36056 |
| Slc20a1 | -0.5387577 | 0.0034558 | 0.02020094 | 252.104687 | 173.450633 |
| AU040320 | -0.5386673 | 0.00846105 | 0.04003475 | 853.640179 | 587.471303 |
| Gpld1 | -0.5381189 | 0.0046141 | 0.02519723 | 237.497411 | 163.380766 |
| Hinfp | -0.5379561 | 0.00161274 | 0.01119083 | 310.209046 | 213.749928 |
| Snx27 | -0.5378526 | 0.00293264 | 0.01786227 | 1041.95612 | 717.477508 |
| Fam91a1 | -0.5378029 | 0.0053677 | 0.0285044 | 1271.80126 | 875.919669 |
| Tbc1d5 | -0.5376437 | 0.0002444 | 0.00243474 | 872.041876 | 600.608499 |
| Itgam | -0.536184 | 0.0002422 | 0.00241763 | 2362.79539 | 1629.04812 |
| Ctbp1 | -0.5360182 | 8.4938E-06 | 0.00014786 | 3146.11125 | 2169.58538 |
| Ppp6r3 | -0.5359104 | 0.00043137 | 0.00383917 | 1012.81586 | 698.559249 |
| Zadh2 | -0.5351142 | 0.00051846 | 0.0044686 | 313.015473 | 215.907997 |
| Slc16a10 | -0.5345669 | 0.0071654 | 0.03552415 | 353.092644 | 243.73943 |
| Usp7 | -0.5327636 | 8.9987E-05 | 0.00107633 | 1203.68688 | 832.168589 |

| | | | | | |
|---|---|---|---|---|---|
| Crkl | -0.532714 | 0.00216783 | 0.01409841 | 993.225828 | 686.23507 |
| C2cd2l | -0.5322643 | 0.00173854 | 0.01190975 | 494.095675 | 341.36002 |
| Mfsd13a | -0.5321257 | 0.00158694 | 0.01105776 | 441.63249 | 305.233089 |
| Macf1 | -0.5317359 | 0.00267618 | 0.01667537 | 1234.9645 | 854.460525 |
| Pcnt | -0.5316235 | 0.00548727 | 0.02895445 | 510.340982 | 353.210235 |
| Fnbp1 | -0.531445 | 1.1711E-05 | 0.00019419 | 1796.57311 | 1242.80953 |
| 2410006H16I | -0.5301453 | 0.00731426 | 0.03602695 | 550.8238 | 381.355702 |
| Smg6 | -0.5300534 | 0.00016839 | 0.00178151 | 357.282366 | 247.627294 |
| Pag1 | -0.5299999 | 0.00165287 | 0.01142185 | 349.14469 | 241.860752 |
| Impad1 | -0.5292899 | 0.00329131 | 0.019482 | 459.021097 | 317.841094 |
| Pafah1b1 | -0.5292332 | 0.00053775 | 0.00459798 | 667.027262 | 461.981509 |
| Trp53inp2 | -0.5291791 | 0.0101296 | 0.04578128 | 252.628835 | 174.8752 |
| March6 | -0.5288059 | 0.00433813 | 0.02405749 | 302.870577 | 209.865468 |
| Sft2d2 | -0.5279191 | 0.00341844 | 0.02002935 | 2416.48425 | 1675.69828 |
| Asxl2 | -0.5272638 | 0.00046442 | 0.0040753 | 565.477003 | 392.469836 |
| AU022252 | -0.5258808 | 0.00582651 | 0.03036129 | 278.190716 | 193.322937 |
| Esyt1 | -0.5250816 | 0.0012026 | 0.00881303 | 6877.88556 | 4779.33338 |
| Rab43 | -0.5247191 | 0.00239747 | 0.01525509 | 330.894002 | 230.030602 |
| Atp13a2 | -0.5245041 | 0.00072165 | 0.00584742 | 4602.98623 | 3199.72215 |
| Lpin2 | -0.523116 | 0.00329525 | 0.01949765 | 798.80678 | 555.726749 |
| Arhgap11a | -0.5220076 | 0.00584165 | 0.03040229 | 839.087195 | 584.140738 |
| Map3k10 | -0.5215728 | 0.00319185 | 0.01903578 | 209.641674 | 145.952354 |
| Plekhm3 | -0.5207792 | 5.859E-06 | 0.0001088 | 1577.19716 | 1099.35918 |
| Atl1 | -0.5198352 | 0.00790962 | 0.03817099 | 109.625917 | 76.3545099 |
| Hspa13 | -0.5197893 | 0.00341617 | 0.02002935 | 374.848888 | 261.296564 |
| Ifrd2 | -0.5194957 | 0.0087586 | 0.04114126 | 131.831548 | 92.0388798 |
| Gm9754 | -0.5194679 | 0.00283143 | 0.01740448 | 214.637271 | 149.641061 |
| Zfyve16 | -0.5194418 | 0.00214739 | 0.01401402 | 298.0443 | 207.864945 |
| Foxn2 | -0.5193082 | 0.00018293 | 0.00190447 | 858.021608 | 598.545671 |
| Pus1 | -0.5185664 | 0.00192452 | 0.01281563 | 318.279893 | 222.12057 |
| Plcxd1 | -0.5180346 | 0.00176362 | 0.01203218 | 358.231634 | 250.179633 |
| Trafd1 | -0.5179114 | 4.8935E-05 | 0.00064487 | 3260.5426 | 2276.95104 |
| Sel1l | -0.5177157 | 0.0011094 | 0.00830423 | 2522.9043 | 1762.0875 |
| Slc43a2 | -0.5168202 | 8.4572E-06 | 0.00014745 | 3221.7828 | 2251.48591 |
| Kpnb1 | -0.5166926 | 0.00449333 | 0.02473553 | 446.671757 | 312.287495 |
| Pum1 | -0.5156116 | 0.00275477 | 0.01701653 | 1510.82008 | 1056.71685 |
| Slc9a8 | -0.5150546 | 0.00191706 | 0.01279996 | 544.428156 | 380.908608 |
| Atxn3 | -0.5143373 | 0.00345522 | 0.02020094 | 688.777466 | 481.997991 |
| Fto | -0.5138246 | 0.00021661 | 0.00220473 | 841.18754 | 589.027037 |
| Ube2d2a | -0.5131629 | 2.6086E-06 | 5.4128E-05 | 1214.30085 | 850.576534 |
| Fbxw8 | -0.513068 | 9.8331E-05 | 0.00116226 | 695.309195 | 486.936641 |
| Plcb1 | -0.5118152 | 0.00011302 | 0.00130211 | 1139.04141 | 798.82639 |
| Zbtb20 | -0.5116185 | 0.00676387 | 0.03396176 | 674.622198 | 473.363169 |
| Gon4l | -0.5114194 | 0.00802883 | 0.03854811 | 249.358249 | 174.995736 |
| Arhgap30 | -0.5103337 | 0.00017815 | 0.00186503 | 7571.36723 | 5315.40008 |
| Mier3 | -0.5098645 | 0.00273951 | 0.0169574 | 336.744284 | 236.406197 |

| | | | | | |
|---|---|---|---|---|---|
| Kif1b | -0.509799 | 0.00078872 | 0.00624618 | 470.684701 | 330.565832 |
| Stt3b | -0.5087782 | 0.00066001 | 0.00543902 | 2296.21332 | 1613.59443 |
| 4732471J01R | -0.5086024 | 0.00111531 | 0.00833496 | 328.954691 | 231.264429 |
| Tpcn1 | -0.5082452 | 0.00616339 | 0.03169893 | 259.015168 | 182.16229 |
| Ckap4 | -0.5080541 | 0.00397616 | 0.02251559 | 1867.02416 | 1312.59977 |
| Cmtr1 | -0.507884 | 0.00941441 | 0.04331781 | 210.253038 | 148.022387 |
| Pik3ca | -0.5075442 | 0.00481708 | 0.02610641 | 986.979368 | 694.219147 |
| Rgl2 | -0.5074073 | 0.00519301 | 0.02778298 | 1533.46749 | 1078.96902 |
| Zbtb43 | -0.5068754 | 0.00159084 | 0.0110731 | 337.572523 | 237.38332 |
| Acin1 | -0.5066044 | 0.0007844 | 0.00621852 | 751.037293 | 528.935317 |
| Spty2d1 | -0.5058995 | 0.00558724 | 0.0293589 | 892.815645 | 628.7315 |
| Fem1a | -0.5058948 | 0.00115276 | 0.0085394 | 970.26549 | 683.385092 |
| Hivep3 | -0.5058901 | 0.00673289 | 0.03385149 | 434.139139 | 305.885145 |
| Chd6 | -0.5058045 | 0.00430184 | 0.0238827 | 341.434521 | 240.682431 |
| 2900097C17F | -0.5052685 | 7.1881E-05 | 0.00089208 | 1173.37321 | 826.487102 |
| Rap1gds1 | -0.5033739 | 0.00106611 | 0.00804834 | 1518.43265 | 1071.04701 |
| Fam76a | -0.503276 | 0.00051879 | 0.00446889 | 588.805354 | 415.330195 |
| Btbd7 | -0.5032642 | 0.00049741 | 0.00431443 | 367.785958 | 259.601008 |
| Vps54 | -0.5028802 | 0.00321476 | 0.01913661 | 957.282492 | 675.440428 |
| Lats1 | -0.5025333 | 0.00835035 | 0.03959826 | 168.489852 | 118.904969 |
| Ccdc71 | -0.5024153 | 0.00341737 | 0.02002935 | 751.191993 | 529.970516 |
| Coa3 | 0.5002936 | 0.00017831 | 0.00186542 | 1220.56247 | 1726.57584 |
| Prpsap2 | 0.50159401 | 3.2297E-05 | 0.00045261 | 404.622715 | 572.743647 |
| Acap1 | 0.50213347 | 0.01070202 | 0.04759514 | 691.037859 | 978.911376 |
| Msantd4 | 0.50268315 | 0.0023648 | 0.01510644 | 284.992786 | 403.556018 |
| Evi2a | 0.50276743 | 2.2302E-05 | 0.00033272 | 3540.9255 | 5017.11914 |
| Ptpn6 | 0.50279134 | 1.6781E-05 | 0.0002599 | 18972.1534 | 26882.4618 |
| Zfp54 | 0.50279687 | 0.00776707 | 0.03769046 | 88.1089191 | 124.9918 |
| Klf10 | 0.5029375 | 0.00028746 | 0.00277893 | 4293.81875 | 6084.4047 |
| Gm26768 | 0.503637 | 0.00088257 | 0.00688413 | 188.754642 | 267.46784 |
| Iqcc | 0.50400467 | 0.01105973 | 0.04863887 | 147.731342 | 209.704506 |
| Pea15a | 0.504813 | 0.00072954 | 0.00588607 | 1297.03031 | 1840.76796 |
| Tor2a | 0.50552501 | 0.00138799 | 0.00989641 | 938.679505 | 1332.4145 |
| Prrg2 | 0.50588081 | 0.00039371 | 0.00356546 | 273.948021 | 388.866064 |
| Gm21370 | 0.50619061 | 2.01E-05 | 0.00030552 | 2376.17127 | 3374.67506 |
| Cabp7 | 0.5076912 | 0.00108424 | 0.00816055 | 362.103898 | 514.754929 |
| Dusp2 | 0.50805122 | 0.00596798 | 0.0308736 | 794.730379 | 1130.57126 |
| Gtf2h3 | 0.50835337 | 0.00030598 | 0.00292641 | 344.730527 | 490.236203 |
| Rpf2 | 0.50896238 | 3.7681E-05 | 0.00051744 | 418.426912 | 595.332512 |
| Otulin | 0.50910057 | 3.1277E-08 | 1.1187E-06 | 1406.18248 | 2001.14886 |
| Zc3hc1 | 0.51015106 | 0.00094921 | 0.00727179 | 450.135886 | 640.970384 |
| Pex16 | 0.51034326 | 4.0819E-05 | 0.00055345 | 434.869888 | 619.573712 |
| Nprl2 | 0.51041394 | 0.00046197 | 0.00405861 | 397.167908 | 565.918397 |
| Hmces | 0.51084267 | 0.00473272 | 0.02567701 | 151.511975 | 215.950009 |
| C130050O18I | 0.51087558 | 0.00056615 | 0.00480454 | 1722.15923 | 2453.7177 |
| Tmem59 | 0.51101073 | 0.00026757 | 0.00262382 | 7437.38577 | 10598.5403 |

| | | | | | |
|---|---|---|---|---|---|
| Ercc1 | 0.51185502 | 0.00052882 | 0.00453448 | 454.712726 | 648.450568 |
| Lcp2 | 0.51224186 | 0.00052146 | 0.0044893 | 5272.63769 | 7519.91858 |
| Aldh3b1 | 0.51227506 | 0.00108544 | 0.00816148 | 3473.65175 | 4954.13907 |
| Chmp2b | 0.51270491 | 0.00025892 | 0.00255069 | 1564.93878 | 2232.38254 |
| Gm11626 | 0.51273364 | 0.00016293 | 0.00173471 | 828.421896 | 1182.11473 |
| Ptcd3 | 0.51279805 | 0.00691293 | 0.03452557 | 341.873837 | 487.833897 |
| Gm15829 | 0.51342321 | 0.00370948 | 0.02138848 | 97.9522789 | 139.941868 |
| Arpc5l | 0.51347651 | 4.3519E-07 | 1.1272E-05 | 1147.63209 | 1638.31091 |
| Cers4 | 0.51354358 | 0.00456537 | 0.02501298 | 106.585959 | 151.899658 |
| Gm5547 | 0.51462131 | 0.00266434 | 0.01661534 | 233.757484 | 334.144608 |
| Ppcs | 0.51478301 | 0.00890353 | 0.0415784 | 189.990003 | 271.479879 |
| Pithd1 | 0.51499918 | 0.0008421 | 0.00661008 | 763.162975 | 1090.53824 |
| Cfp | 0.51503468 | 0.00928982 | 0.04288903 | 3729.89138 | 5330.05866 |
| Rab8a | 0.51535098 | 2.0949E-06 | 4.5024E-05 | 5207.30597 | 7442.73895 |
| Cd300a | 0.51693459 | 6.0039E-06 | 0.00011067 | 12934.7004 | 18508.0851 |
| Ubb | 0.51698541 | 1.9135E-07 | 5.507E-06 | 17971.1925 | 25716.1195 |
| E130208F15R | 0.51841181 | 0.00049674 | 0.00431131 | 279.224533 | 400.168988 |
| Pilrb1 | 0.51877181 | 0.00013472 | 0.0014918 | 1576.26651 | 2258.35391 |
| Fez2 | 0.51890148 | 4.2678E-05 | 0.00057348 | 649.583189 | 930.60747 |
| Aaas | 0.51942831 | 0.00041327 | 0.00370217 | 303.857564 | 435.346445 |
| Cyb561a3 | 0.52021696 | 0.00073313 | 0.00590549 | 1293.76721 | 1855.41413 |
| Ndufaf1 | 0.52022131 | 0.00768202 | 0.03737795 | 205.769854 | 294.830229 |
| 2310031A07R | 0.52027299 | 0.00087046 | 0.00680382 | 426.477313 | 611.570966 |
| Kpna2 | 0.520354 | 0.00095264 | 0.00729134 | 1711.65173 | 2454.84978 |
| Slamf8 | 0.52083567 | 0.00039983 | 0.00361191 | 2226.49969 | 3194.81879 |
| Dph2 | 0.52301976 | 0.00215571 | 0.01405612 | 220.890155 | 317.163705 |
| Trf | 0.52415969 | 0.00056101 | 0.0047751 | 264.345026 | 380.162465 |
| Abi3 | 0.52487076 | 3.5828E-06 | 7.1009E-05 | 5783.41578 | 8321.25485 |
| Faap24 | 0.52657103 | 0.00017389 | 0.00182933 | 258.358339 | 372.250946 |
| Gas8 | 0.52690949 | 0.00534342 | 0.02843676 | 104.023864 | 149.825836 |
| Cd40 | 0.52721853 | 0.00015113 | 0.00163456 | 928.072295 | 1337.56633 |
| Cbr1 | 0.52786714 | 0.00593708 | 0.03078096 | 233.18041 | 335.942971 |
| Fra10ac1 | 0.52940051 | 0.00115369 | 0.00854209 | 143.093568 | 206.838164 |
| Cyp4f16 | 0.52961622 | 1.2766E-05 | 0.00020846 | 3323.08033 | 4797.30012 |
| Clpx | 0.52984887 | 0.0001015 | 0.00119215 | 600.644435 | 867.148803 |
| Clec7a | 0.53027708 | 1.5103E-05 | 0.00023909 | 6299.27531 | 9097.07096 |
| Rars2 | 0.5304554 | 0.00118933 | 0.00874564 | 345.959173 | 499.653771 |
| Nsdhl | 0.530829 | 0.00996513 | 0.0452421 | 224.882584 | 324.626771 |
| Tmem176a | 0.53127262 | 0.00187558 | 0.01260711 | 2904.89605 | 4198.56547 |
| Crcp | 0.53187424 | 2.9869E-05 | 0.00042292 | 621.874885 | 899.182881 |
| Pilrb2 | 0.53287281 | 4.3584E-05 | 0.00058461 | 1414.26671 | 2046.30637 |
| BC035044 | 0.53294864 | 0.00056639 | 0.00480454 | 1272.33654 | 1841.00977 |
| Fam32a | 0.53463264 | 2.238E-06 | 4.7422E-05 | 4153.5737 | 6016.61343 |
| Rnase6 | 0.53564163 | 5.326E-07 | 1.3425E-05 | 3429.65596 | 4971.41204 |
| Crem | 0.53736286 | 0.00824578 | 0.03927503 | 92.7707766 | 134.794815 |
| Hnrnph2 | 0.53793918 | 0.00034258 | 0.00318282 | 2170.76518 | 3151.49506 |

| | | | | | |
|---|---|---|---|---|---|
| Gm20531 | 0.53825024 | 2.0217E-05 | 0.00030648 | 3974.75929 | 5771.96848 |
| Mtrr | 0.53962819 | 0.00948984 | 0.04353157 | 109.736203 | 159.264613 |
| A230005M16 | 0.53993874 | 0.00012179 | 0.00138292 | 741.745119 | 1078.59494 |
| AW011738 | 0.5401427 | 0.00242823 | 0.01540512 | 148.441421 | 215.970597 |
| Exoc7 | 0.5406259 | 3.6096E-05 | 0.00050071 | 936.37062 | 1361.52251 |
| Samsn1 | 0.54185749 | 0.00042995 | 0.00383103 | 3919.13646 | 5705.46503 |
| Cd27 | 0.54203525 | 0.00333841 | 0.01967304 | 133.859299 | 195.29041 |
| Tfpt | 0.54213465 | 4.097E-05 | 0.00055499 | 258.447223 | 376.380547 |
| Bcl2a1d | 0.54242605 | 0.0016003 | 0.01111993 | 253.239019 | 369.078435 |
| Ech1 | 0.54263303 | 6.2417E-05 | 0.00079196 | 1912.65548 | 2785.96304 |
| Coq7 | 0.54476221 | 0.0051644 | 0.02766931 | 109.247465 | 159.366804 |
| Slc2a8 | 0.54493163 | 0.00538148 | 0.02856748 | 190.111987 | 277.658268 |
| Abhd12 | 0.54493448 | 8.0795E-06 | 0.00014202 | 2547.04534 | 3715.66175 |
| Tdp1 | 0.54518628 | 0.00174428 | 0.01192472 | 283.954456 | 414.190762 |
| Gab3 | 0.54667107 | 0.00204171 | 0.01343521 | 343.449224 | 501.912171 |
| Mrps27 | 0.54681962 | 0.00269617 | 0.01675124 | 290.871904 | 424.99108 |
| Pdrg1 | 0.54687163 | 4.418E-05 | 0.00059102 | 871.549089 | 1273.618 |
| Pim2 | 0.54788033 | 0.00074849 | 0.00599772 | 250.043988 | 365.983982 |
| Usp39 | 0.54825028 | 0.00039374 | 0.00356546 | 727.183932 | 1063.44791 |
| Gm10434 | 0.54880897 | 0.00298429 | 0.01809875 | 166.435398 | 243.459579 |
| BC147527 | 0.54926089 | 1.312E-05 | 0.00021378 | 1993.81657 | 2917.68371 |
| Coq8a | 0.5496937 | 0.00282165 | 0.01736288 | 326.26645 | 477.313318 |
| B9d2 | 0.54971255 | 5.9545E-06 | 0.00011003 | 937.111353 | 1371.61924 |
| Acadvl | 0.54980583 | 7.2305E-06 | 0.00012947 | 1303.27208 | 1907.82052 |
| Isoc2b | 0.54984725 | 0.00073995 | 0.0059477 | 177.308715 | 259.558083 |
| Zfp961 | 0.55097488 | 0.00083658 | 0.00657659 | 1926.00448 | 2821.73089 |
| Ttc33 | 0.55171592 | 0.00014691 | 0.00159465 | 507.357927 | 743.733398 |
| Gm45507 | 0.55226047 | 6.2196E-05 | 0.00079181 | 278.40665 | 408.123816 |
| Zc2hc1a | 0.55567699 | 0.00290525 | 0.01777524 | 128.948028 | 189.543155 |
| Otulinl | 0.55621463 | 1.2434E-05 | 0.00020349 | 10162.861 | 14943.5975 |
| Gm42477 | 0.55621716 | 0.00097549 | 0.00744653 | 284.593256 | 418.564698 |
| Sumo1 | 0.55641897 | 3.9762E-08 | 1.3764E-06 | 2087.28183 | 3069.61293 |
| Bnip3 | 0.55670624 | 0.00026299 | 0.00258402 | 469.310919 | 690.250144 |
| Snrnp27 | 0.55703056 | 8.6704E-05 | 0.00104602 | 509.845698 | 750.188637 |
| Galm | 0.55709158 | 0.00301134 | 0.01821961 | 255.308858 | 375.225137 |
| Sat1 | 0.55710937 | 4.4558E-07 | 1.1443E-05 | 19497.8423 | 28687.7583 |
| Cdc42ep3 | 0.55718448 | 3.3849E-07 | 9.0482E-06 | 2571.29918 | 3783.15208 |
| Gm15931 | 0.55768144 | 6.7988E-06 | 0.00012306 | 1042.3118 | 1533.97964 |
| Bmyc | 0.55953247 | 7.42E-06 | 0.00013207 | 996.839024 | 1469.326 |
| Rogdi | 0.55968849 | 6.1024E-05 | 0.00077971 | 1460.68589 | 2153.02586 |
| Cyp39a1 | 0.56161254 | 0.00022245 | 0.00225191 | 431.167245 | 636.041047 |
| Cd200r1 | 0.56188658 | 0.00141058 | 0.01001946 | 288.850419 | 426.667495 |
| Ifngr1 | 0.56235505 | 2.3361E-05 | 0.00034572 | 22808.3957 | 33680.5131 |
| 4933412E12F | 0.5637779 | 0.00940563 | 0.04330396 | 86.768728 | 128.430644 |
| Tbcc | 0.56378043 | 8.2539E-05 | 0.00100433 | 552.501989 | 816.673064 |
| Ccdc130 | 0.56587975 | 0.00296268 | 0.01800995 | 314.308713 | 465.336386 |

| | | | | | |
|---|---|---|---|---|---|
| Nosip | 0.56593614 | 0.00010071 | 0.00118666 | 1116.09748 | 1652.3218 |
| Lin37 | 0.56610442 | 2.2028E-06 | 4.6808E-05 | 510.163233 | 755.190128 |
| Mettl18 | 0.56622764 | 0.00325693 | 0.01930131 | 73.204202 | 108.3956 |
| Oplah | 0.56694073 | 0.00013584 | 0.00150277 | 207.50499 | 307.693548 |
| Tax1bp3 | 0.56748047 | 0.00128827 | 0.00933155 | 706.761851 | 1047.14062 |
| Cnn3 | 0.56792819 | 1.3721E-05 | 0.00022117 | 513.285328 | 760.828655 |
| Sucla2 | 0.56817077 | 1.6384E-05 | 0.00025534 | 1144.32324 | 1696.55436 |
| Exosc4 | 0.56867863 | 8.7256E-06 | 0.00015102 | 710.086296 | 1053.51228 |
| Stat4 | 0.56889741 | 0.00560148 | 0.02939262 | 187.288918 | 277.739612 |
| Rrad | 0.56988828 | 5.7046E-05 | 0.00073752 | 333.309464 | 494.969744 |
| Metrnl | 0.57223246 | 1.1085E-06 | 2.5494E-05 | 6552.43312 | 9742.24899 |
| Sdhaf1 | 0.57397007 | 0.00015402 | 0.0016599 | 348.085058 | 518.328547 |
| Cenpl | 0.57441027 | 0.0047081 | 0.02559889 | 150.440646 | 223.844694 |
| Cwc22 | 0.57532696 | 0.00231657 | 0.01486507 | 308.173482 | 458.863062 |
| Chn2 | 0.57533212 | 0.01039561 | 0.04659105 | 206.723499 | 308.111476 |
| Pars2 | 0.576835 | 0.00384914 | 0.02196186 | 106.883082 | 159.510577 |
| Chmp6 | 0.57739437 | 7.0083E-05 | 0.00087665 | 1048.4514 | 1564.08432 |
| 4833407H14I | 0.57764206 | 0.00152865 | 0.01073129 | 574.878834 | 858.154375 |
| Trem3 | 0.57868818 | 0.00015088 | 0.00163308 | 965.203159 | 1441.19796 |
| Ptcd2 | 0.5790971 | 2.142E-06 | 4.5775E-05 | 963.667474 | 1439.68028 |
| Ube2l6 | 0.57940804 | 0.00107548 | 0.00810685 | 1712.65106 | 2558.81367 |
| Pilra | 0.57973873 | 7.5991E-05 | 0.00093421 | 2642.64535 | 3949.50373 |
| Cops4 | 0.57976273 | 0.00011559 | 0.00132379 | 1105.73175 | 1652.62104 |
| Gm49500 | 0.58105951 | 0.00044176 | 0.00391768 | 572.318727 | 855.978387 |
| H2-Q7 | 0.58122949 | 0.00709342 | 0.03523956 | 1000.31872 | 1497.02995 |
| Cd83 | 0.58314196 | 0.0061932 | 0.0317978 | 380.151905 | 569.164614 |
| Scimp | 0.5836918 | 0.01028647 | 0.04618459 | 117.929593 | 177.011583 |
| Dnajc8 | 0.58654517 | 1.9253E-06 | 4.1618E-05 | 2062.77892 | 3097.4418 |
| Slc46a3 | 0.58688341 | 0.00086088 | 0.00673946 | 2365.76248 | 3553.37707 |
| Slc19a1 | 0.58702454 | 0.00553248 | 0.02911188 | 145.927477 | 219.039659 |
| Tmem206 | 0.58962021 | 0.00252174 | 0.01586437 | 358.988022 | 540.155475 |
| Gngt2 | 0.58967614 | 6.7064E-06 | 0.00012164 | 7529.18624 | 11330.966 |
| Bphl | 0.5899547 | 0.00181001 | 0.01226516 | 328.240875 | 494.037898 |
| Ginm1 | 0.59035918 | 4.3514E-06 | 8.4349E-05 | 602.854344 | 907.381607 |
| Sugp1 | 0.5910719 | 0.00366318 | 0.02119873 | 725.240597 | 1092.3741 |
| Tnni2 | 0.59149855 | 0.00224762 | 0.01449181 | 135.819058 | 204.572078 |
| Kctd13 | 0.59150343 | 1.3635E-05 | 0.00022018 | 297.142522 | 447.792868 |
| 0610040J01R | 0.59184701 | 0.00313045 | 0.01881147 | 783.03052 | 1180.02792 |
| Brix1 | 0.59331675 | 2.1017E-05 | 0.00031606 | 691.124686 | 1042.65955 |
| Gm36551 | 0.59348761 | 0.00033398 | 0.00311834 | 419.438859 | 632.722682 |
| Susd3 | 0.59372022 | 8.8047E-08 | 2.7847E-06 | 3542.43429 | 5345.85758 |
| Ccr5 | 0.59380201 | 0.00844983 | 0.04000687 | 222.520256 | 335.915644 |
| Hilpda | 0.59438987 | 0.00846898 | 0.04005296 | 75.5155013 | 113.739703 |
| Acad12 | 0.59499451 | 0.00888163 | 0.04151485 | 57.3386021 | 86.5520545 |
| Timm17a | 0.59703838 | 2.3121E-06 | 4.8785E-05 | 634.262339 | 959.55547 |
| Cutc | 0.59809195 | 0.00059333 | 0.00499083 | 352.138477 | 532.591786 |

| | | | | | |
|---|---|---|---|---|---|
| Ighm | 0.60002449 | 0.00040864 | 0.00366948 | 13151.7813 | 19934.7259 |
| Zcwpw1 | 0.60100069 | 0.00148968 | 0.01050188 | 216.080486 | 327.70856 |
| Ecm1 | 0.60121525 | 0.00077535 | 0.00615323 | 1354.72147 | 2055.18945 |
| Srfbp1 | 0.60178813 | 1.4406E-05 | 0.00023024 | 199.087191 | 301.952493 |
| Zfp677 | 0.6048315 | 0.00660725 | 0.03335372 | 111.400299 | 169.63988 |
| Aamdc | 0.60520291 | 0.0001201 | 0.00136479 | 174.745035 | 265.936038 |
| Gm26613 | 0.60683307 | 0.00247895 | 0.01565416 | 122.778484 | 186.836635 |
| Cst7 | 0.60728878 | 0.00323728 | 0.01921517 | 155.190904 | 236.597098 |
| Mrm2 | 0.60797004 | 0.00260287 | 0.01627929 | 111.643739 | 169.841313 |
| Acaa2 | 0.60831965 | 5.6963E-05 | 0.00073752 | 2486.25832 | 3789.86879 |
| Rpp30 | 0.60858369 | 0.00015423 | 0.0016609 | 238.992788 | 364.338392 |
| Gm1821 | 0.60875256 | 5.9676E-05 | 0.00076559 | 608.060805 | 927.209226 |
| M1ap | 0.61505767 | 0.00013355 | 0.00148397 | 129.802566 | 198.841869 |
| Rtp4 | 0.61585818 | 0.00035954 | 0.0033158 | 1875.49098 | 2873.98168 |
| Kmo | 0.61638597 | 0.00012289 | 0.00139016 | 872.274082 | 1337.282 |
| Tmx2 | 0.61696806 | 6.5071E-05 | 0.00081942 | 477.680249 | 732.515859 |
| Igtp | 0.62228571 | 8.5512E-06 | 0.00014834 | 1110.39519 | 1709.31128 |
| Manbal | 0.62300802 | 0.00017308 | 0.00182338 | 485.689765 | 747.853365 |
| Nudt1 | 0.62347422 | 0.0002756 | 0.00268501 | 111.23879 | 171.477784 |
| Gm21958 | 0.62609317 | 0.00886211 | 0.04144939 | 71.0493845 | 109.771295 |
| Tmem37 | 0.62651621 | 0.01109432 | 0.04874025 | 102.7309 | 158.927933 |
| Nmd3 | 0.62813955 | 0.0001965 | 0.0020317 | 346.562329 | 535.587313 |
| Rras2 | 0.62877271 | 5.2324E-05 | 0.00068472 | 192.67036 | 297.728785 |
| Mettl22 | 0.63045046 | 0.00620803 | 0.03186305 | 74.1793357 | 115.08742 |
| Ehd3 | 0.63151966 | 0.01066524 | 0.04754417 | 73.3636688 | 113.309001 |
| Sertad3 | 0.63295405 | 6.5107E-07 | 1.6155E-05 | 452.140384 | 700.837747 |
| Tmem51os1 | 0.63437855 | 0.00010058 | 0.00118604 | 411.920013 | 639.349733 |
| Pla2g4a | 0.63646684 | 1.0262E-05 | 0.00017342 | 472.357033 | 734.457245 |
| Commd9 | 0.63792635 | 0.00046535 | 0.00408068 | 274.853062 | 427.640777 |
| Grk5 | 0.64076009 | 6.6913E-05 | 0.0008391 | 754.195975 | 1175.70665 |
| Bco2 | 0.64116478 | 0.00045426 | 0.0040186 | 156.305998 | 243.635389 |
| Trmt10b | 0.64337548 | 0.00073428 | 0.00591166 | 141.940073 | 221.903603 |
| H2-Ob | 0.64364535 | 0.00495068 | 0.026734 | 302.513785 | 472.374069 |
| Pigh | 0.64454145 | 0.00012408 | 0.00140192 | 304.003302 | 475.285153 |
| Dtd1 | 0.64480047 | 0.00459851 | 0.02515782 | 136.287486 | 212.854076 |
| 3110082I17R | 0.64822257 | 6.2811E-06 | 0.00011521 | 1737.67378 | 2723.11751 |
| Acot9 | 0.64909842 | 5.2844E-08 | 1.7733E-06 | 1440.36092 | 2258.49205 |
| Gm5086 | 0.65078645 | 0.00074152 | 0.00595499 | 123.251648 | 193.324867 |
| Lax1 | 0.65113809 | 0.00818888 | 0.03907926 | 144.715019 | 227.504625 |
| Cd81 | 0.6515369 | 0.0047038 | 0.02558478 | 349.322604 | 548.701571 |
| C5ar1 | 0.65210077 | 0.00708814 | 0.03522499 | 899.059 | 1412.70469 |
| Gimap1os | 0.65441511 | 0.0072241 | 0.03573517 | 97.5886558 | 153.67948 |
| Ttc16 | 0.65583056 | 0.00013334 | 0.00148275 | 135.20824 | 212.99276 |
| Mccc2 | 0.65586459 | 0.00056166 | 0.00477791 | 155.959107 | 245.671683 |
| Snap47 | 0.65716639 | 0.00122447 | 0.00895577 | 120.235649 | 189.675262 |
| Cyb5r3 | 0.65725292 | 7.0325E-05 | 0.00087822 | 1527.40919 | 2408.72292 |

| | | | | | |
|---|---|---|---|---|---|
| Gm43495 | 0.65852599 | 4.6982E-06 | 9.0258E-05 | 376.122639 | 593.629692 |
| Gm2a | 0.66670192 | 3.1297E-07 | 8.426E-06 | 26756.8087 | 42474.676 |
| Adgre4 | 0.66715537 | 7.9217E-05 | 0.00097069 | 7011.08269 | 11133.0281 |
| Zc3h8 | 0.66741797 | 0.00402716 | 0.02269922 | 58.3446627 | 92.6471148 |
| Gpr18 | 0.66783934 | 2.7144E-09 | 1.2864E-07 | 1636.33453 | 2600.05502 |
| Cep57l1 | 0.66848372 | 0.01104124 | 0.0486002 | 48.4245983 | 77.1019447 |
| Cd19 | 0.6687573 | 0.00419348 | 0.02342352 | 360.097256 | 572.408586 |
| Il1rn | 0.67227724 | 0.0006128 | 0.00512875 | 487.653301 | 777.251417 |
| Zfp72 | 0.67309722 | 0.00847023 | 0.04005296 | 38.9510137 | 62.1798181 |
| Hpgd | 0.67383097 | 0.00026913 | 0.0026374 | 4993.19119 | 7965.51801 |
| Slc14a1 | 0.6750727 | 0.01113631 | 0.04884695 | 87.6444036 | 139.826376 |
| Gbp2 | 0.67531906 | 0.00060396 | 0.00506601 | 989.510212 | 1580.21453 |
| Slamf9 | 0.67710842 | 0.00826902 | 0.03933675 | 79.0921545 | 126.324985 |
| Samd10 | 0.67804717 | 0.00129082 | 0.00934103 | 148.901884 | 238.262457 |
| Cd55 | 0.67968446 | 0.01131484 | 0.04947224 | 127.394631 | 203.658477 |
| AW112010 | 0.67970677 | 0.00737434 | 0.03622781 | 1433.49131 | 2296.62384 |
| Gm38287 | 0.68122681 | 0.00717386 | 0.03553343 | 113.033147 | 181.349006 |
| Plgrkt | 0.68342151 | 1.5119E-07 | 4.4976E-06 | 508.343157 | 816.6123 |
| Gdf3 | 0.6839447 | 0.00350868 | 0.02044642 | 119.337711 | 191.409765 |
| Upb1 | 0.68641305 | 0.00320901 | 0.01911536 | 205.271585 | 330.608553 |
| Nipsnap1 | 0.68706086 | 0.00048923 | 0.00425825 | 88.3221147 | 141.986445 |
| Ociad1 | 0.68868049 | 0.00398031 | 0.02252212 | 1413.34382 | 2277.96318 |
| Cd1d1 | 0.68983235 | 0.00188577 | 0.01265292 | 217.356968 | 350.697624 |
| Tmem51 | 0.69072582 | 8.9968E-06 | 0.00015535 | 3376.38594 | 5449.74697 |
| Lpar6 | 0.69123319 | 8.6178E-06 | 0.00014933 | 1718.3247 | 2774.90052 |
| Pgap3 | 0.69258827 | 0.01139618 | 0.04971105 | 53.4206134 | 86.33295 |
| Tubd1 | 0.69270024 | 0.00193732 | 0.01288947 | 117.462284 | 189.714242 |
| Fcrl1 | 0.69361805 | 0.00014312 | 0.00156255 | 188.05624 | 303.824214 |
| Nup37 | 0.69429552 | 0.00016174 | 0.00172572 | 93.0220776 | 150.496167 |
| Cyp4f18 | 0.6946081 | 2.7091E-05 | 0.00039209 | 8180.11605 | 13239.099 |
| Nmnat3 | 0.6949975 | 0.00288208 | 0.01767966 | 62.3283458 | 100.85149 |
| Serpina3g | 0.69546727 | 0.00056406 | 0.00479565 | 352.327167 | 570.923869 |
| Plxna4os1 | 0.69551188 | 0.00214915 | 0.01401943 | 242.447382 | 392.557458 |
| Itm2a | 0.69556456 | 0.00734087 | 0.03613432 | 77.5573089 | 125.691582 |
| Engase | 0.69801873 | 0.0001366 | 0.00150675 | 190.103142 | 308.56074 |
| Cd74 | 0.70029517 | 0.00010299 | 0.00120684 | 64804.0385 | 105296.478 |
| Gm17315 | 0.70195421 | 8.7982E-08 | 2.7847E-06 | 480.272652 | 780.973538 |
| Rpap2 | 0.70270254 | 0.00173809 | 0.01190975 | 166.829017 | 271.38386 |
| Phf11b | 0.70352655 | 1.3348E-05 | 0.00021633 | 1369.92969 | 2230.67602 |
| Gss | 0.70472032 | 0.00043444 | 0.00385964 | 677.691985 | 1104.44049 |
| Cxcr3 | 0.70568811 | 0.00324207 | 0.01923605 | 185.479595 | 302.749452 |
| Gm13068 | 0.70625866 | 0.01096871 | 0.0483944 | 38.4338655 | 62.7972026 |
| Zfp775 | 0.70749227 | 0.00306101 | 0.0184607 | 94.4616124 | 154.11317 |
| Mthfsl | 0.70787263 | 3.3258E-08 | 1.1728E-06 | 476.410872 | 778.048187 |
| Tbc1d10c | 0.70805716 | 0.00032483 | 0.00305375 | 752.860845 | 1230.21688 |
| Il21r | 0.70935234 | 2.1292E-06 | 4.5566E-05 | 292.91591 | 479.261321 |

| | | | | | |
|---|---|---|---|---|---|
| Tgtp2 | 0.70952475 | 3.6614E-05 | 0.00050557 | 309.423531 | 506.125725 |
| AC126457.1 | 0.71124243 | 0.00118813 | 0.00874106 | 80.582069 | 132.26868 |
| Calhm6 | 0.71181552 | 9.132E-07 | 2.1503E-05 | 1262.70672 | 2068.60958 |
| Rab3il1 | 0.71283685 | 0.000111 | 0.00128606 | 288.285334 | 472.260212 |
| Spns3 | 0.71363659 | 0.0001694 | 0.00178971 | 187.922338 | 308.351275 |
| Cd7 | 0.71600396 | 0.00643232 | 0.03269035 | 1305.39868 | 2144.56831 |
| Stra6l | 0.71650234 | 0.00880627 | 0.04125593 | 35.0012812 | 57.4509767 |
| Rcan1 | 0.71757082 | 0.00010093 | 0.00118731 | 648.858375 | 1066.92779 |
| Ppp1r15a | 0.72129459 | 0.00049538 | 0.00430431 | 333.050884 | 549.164451 |
| Slc29a1 | 0.7251307 | 8.3807E-05 | 0.00101453 | 1013.19091 | 1674.90412 |
| Lef1 | 0.72517203 | 0.00415525 | 0.02323199 | 129.410892 | 214.221048 |
| P2ry13 | 0.72766702 | 0.00520829 | 0.02782509 | 242.494866 | 401.547891 |
| Pkp3 | 0.72795559 | 2.6186E-06 | 5.4261E-05 | 460.31532 | 762.623768 |
| Nt5c3b | 0.72809364 | 0.00214663 | 0.01401402 | 109.810059 | 181.829952 |
| Irgc1 | 0.72821707 | 0.00011775 | 0.00134421 | 107.711201 | 178.129247 |
| Slc41a2 | 0.72821846 | 0.00808727 | 0.03870612 | 68.0353469 | 112.771542 |
| Tefm | 0.72924802 | 1.2994E-06 | 2.9224E-05 | 168.299167 | 278.953475 |
| Gm11775 | 0.73158808 | 0.00293325 | 0.01786227 | 544.703079 | 904.690927 |
| Vnn3 | 0.73375177 | 4.1078E-06 | 8.0146E-05 | 263.963557 | 438.740145 |
| Cep72 | 0.73553819 | 0.01081495 | 0.04794359 | 56.6986265 | 94.2298806 |
| Blnk | 0.73567391 | 0.00644921 | 0.03274299 | 234.884792 | 391.155259 |
| Gm6377 | 0.73670206 | 0.0066999 | 0.03371946 | 118.847702 | 197.746928 |
| Cd79b | 0.73764995 | 0.00023692 | 0.00237754 | 867.932891 | 1447.01406 |
| Tmem35b | 0.74180506 | 0.0003798 | 0.00346248 | 207.089476 | 346.398291 |
| Ctsw | 0.74313366 | 0.00665979 | 0.03355134 | 392.89916 | 658.018061 |
| Gm43545 | 0.74315576 | 0.00013625 | 0.001505 | 67.3920729 | 112.724558 |
| Ctsf | 0.74337019 | 0.00120225 | 0.00881303 | 63.255624 | 105.831467 |
| H2-DMb1 | 0.74442045 | 6.7065E-07 | 1.6571E-05 | 4573.65488 | 7662.72482 |
| Casp8 | 0.74566694 | 0.00010082 | 0.00118697 | 2091.91989 | 3507.62495 |
| Mir6381 | 0.74676825 | 0.00225604 | 0.01453363 | 75.4463119 | 126.649243 |
| Slc25a53 | 0.75069088 | 0.00106225 | 0.00802725 | 102.406692 | 172.692155 |
| AC140186.1 | 0.75174174 | 0.00613985 | 0.03161031 | 584.97655 | 985.241243 |
| Hmgn3 | 0.75325322 | 0.00325328 | 0.01928729 | 79.7400552 | 134.520659 |
| Il1b | 0.75623371 | 0.00011982 | 0.00136474 | 860.092787 | 1453.1173 |
| Gm32633 | 0.75664404 | 1.9028E-07 | 5.4868E-06 | 308.086864 | 520.710445 |
| Tspan13 | 0.75810108 | 9.0666E-09 | 3.7835E-07 | 3981.55884 | 6734.11367 |
| Rab19 | 0.75875097 | 0.00046716 | 0.00409171 | 307.599463 | 520.581724 |
| Mthfs | 0.75965073 | 0.00046737 | 0.00409171 | 76.2184339 | 128.883661 |
| Pls3 | 0.76136585 | 0.00829026 | 0.03938789 | 115.408069 | 195.645388 |
| Coq3 | 0.76683909 | 0.00164228 | 0.01136954 | 139.578957 | 237.350192 |
| Paqr5 | 0.76747881 | 0.01050217 | 0.04690332 | 50.3392296 | 85.628435 |
| Tmem221 | 0.76843575 | 0.00315132 | 0.01889156 | 185.173135 | 315.634369 |
| Gm17080 | 0.77808108 | 2.1551E-05 | 0.00032311 | 346.468066 | 594.13524 |
| Hes1 | 0.77869356 | 0.00065522 | 0.00541141 | 440.160395 | 755.151031 |
| Dok2 | 0.78205548 | 0.00066094 | 0.00544366 | 585.80994 | 1007.87126 |
| Dnajc17 | 0.78410903 | 4.7815E-06 | 9.1274E-05 | 115.996794 | 199.473517 |

| | | | | | |
|---|---|---|---|---|---|
| 6430562O15I | 0.7852034 | 0.0003888 | 0.00352925 | 101.260564 | 174.612181 |
| Dhrs3 | 0.78544334 | 0.01117089 | 0.04892721 | 290.207428 | 500.228512 |
| H2-Oa | 0.78765396 | 0.00012518 | 0.00140968 | 134.209331 | 231.517457 |
| Trim5 | 0.79252473 | 3.8107E-06 | 7.4933E-05 | 160.791823 | 278.368964 |
| Phf11a | 0.79824127 | 0.00017949 | 0.00187511 | 204.141485 | 355.024955 |
| Tnfrsf13c | 0.80149332 | 0.00337356 | 0.01985155 | 166.498441 | 290.02707 |
| Siglecg | 0.80237189 | 0.00021996 | 0.00223273 | 211.165936 | 368.259614 |
| Ltb | 0.80294324 | 2.5691E-06 | 5.3383E-05 | 721.428248 | 1258.51741 |
| Grap | 0.8032798 | 1.7864E-06 | 3.8951E-05 | 398.116182 | 694.772684 |
| Mmp11 | 0.80336392 | 0.00918423 | 0.04252014 | 36.6749227 | 63.9067246 |
| H2-DMb2 | 0.80500969 | 1.8088E-06 | 3.9326E-05 | 501.831066 | 876.558473 |
| Gm21596 | 0.80620323 | 0.00448813 | 0.02471719 | 31.0008787 | 54.1306331 |
| 1700071M16 | 0.80697151 | 3.2083E-07 | 8.6068E-06 | 1911.65567 | 3344.45653 |
| Gm3608 | 0.81069827 | 0.00804695 | 0.03858259 | 37.503559 | 65.6669272 |
| Cd209a | 0.81593865 | 0.00035177 | 0.00325243 | 4198.62533 | 7391.65991 |
| Lair1 | 0.8161365 | 6.3982E-07 | 1.5914E-05 | 1243.57795 | 2189.63679 |
| Slain1 | 0.81743013 | 0.00544254 | 0.02877974 | 64.506395 | 113.713333 |
| Gm33023 | 0.81889464 | 0.00012726 | 0.00142775 | 78.9283411 | 139.374936 |
| Cd274 | 0.82352058 | 0.00109743 | 0.00823921 | 273.780938 | 484.797546 |
| Slamf6 | 0.82547274 | 1.3815E-05 | 0.00022245 | 121.701873 | 215.467323 |
| Chil5 | 0.82610127 | 0.00048662 | 0.00424043 | 46.1152954 | 81.6876991 |
| Zfp3 | 0.82690573 | 0.00202328 | 0.01334901 | 64.196593 | 113.982366 |
| S1pr1 | 0.82704361 | 7.4222E-05 | 0.00091622 | 239.685704 | 425.399382 |
| Ms4a1 | 0.83539546 | 0.00212661 | 0.01391465 | 460.738539 | 821.780617 |
| Hsh2d | 0.83649789 | 0.0001962 | 0.00202998 | 191.414862 | 341.625304 |
| Zcchc18 | 0.84152426 | 0.00269214 | 0.01673455 | 57.0510121 | 102.354892 |
| Gprc5c | 0.84248876 | 0.00131643 | 0.00948983 | 113.836875 | 204.176413 |
| Gimap3 | 0.84256951 | 0.00037403 | 0.00343045 | 632.753077 | 1135.08356 |
| Skap1 | 0.84264514 | 0.00246053 | 0.01557055 | 148.240346 | 266.040562 |
| Gm10634 | 0.84482935 | 5.757E-06 | 0.00010717 | 98.2160482 | 176.376894 |
| Gimap5 | 0.84605124 | 0.00030279 | 0.00291029 | 112.873287 | 203.27229 |
| Pla2g7 | 0.84917424 | 2.5579E-09 | 1.2161E-07 | 9147.64336 | 16479.1245 |
| Gpr34 | 0.85165661 | 0.00178036 | 0.0121134 | 92.330999 | 166.822504 |
| Slc25a33 | 0.85222153 | 1.5572E-05 | 0.00024466 | 84.886615 | 153.398572 |
| Gm10384 | 0.85451953 | 1.6174E-05 | 0.00025258 | 143.890222 | 260.329998 |
| Ifi205 | 0.86005611 | 0.000125 | 0.00140879 | 694.620627 | 1260.86074 |
| C1qb | 0.86327052 | 3.5012E-05 | 0.00048747 | 186.453821 | 339.407397 |
| Cd200r4 | 0.8645945 | 0.00589073 | 0.03061102 | 42.8217609 | 78.2170073 |
| H2-Eb1 | 0.86829218 | 2.2878E-07 | 6.3885E-06 | 7446.73992 | 13594.679 |
| Cd6 | 0.87139695 | 0.00381107 | 0.02179433 | 115.551972 | 211.293461 |
| Cxcr5 | 0.87202665 | 0.00603779 | 0.03118109 | 127.951209 | 233.800383 |
| 2610035D17I | 0.87523934 | 0.00060735 | 0.0050888 | 143.861208 | 264.055676 |
| Tbxa2r | 0.87593485 | 0.00161663 | 0.0112075 | 55.2945451 | 101.346577 |
| Klra17 | 0.87746603 | 0.00338869 | 0.01991717 | 117.211436 | 215.522529 |
| Gbp4 | 0.8775686 | 0.00245871 | 0.0155656 | 126.521903 | 232.736724 |
| Ryr1 | 0.88095668 | 0.00170144 | 0.01170137 | 93.3891986 | 171.894215 |

| | | | | | |
|---|---|---|---|---|---|
| Cd38 | 0.88171006 | 0.00032841 | 0.0030759 | 216.52881 | 398.762853 |
| Id3 | 0.88329788 | 4.5717E-08 | 1.5609E-06 | 198.1629 | 365.366443 |
| Fcrla | 0.88973664 | 1.8424E-05 | 0.00028243 | 114.975314 | 212.766669 |
| Ms4a4b | 0.88977869 | 5.2783E-06 | 9.9995E-05 | 1103.65773 | 2045.29133 |
| Rcn3 | 0.89214298 | 0.00211372 | 0.01384839 | 76.5989348 | 142.381178 |
| Gm16008 | 0.89561824 | 0.00053126 | 0.00455022 | 61.3808167 | 114.473466 |
| Ccnd1 | 0.89736647 | 1.4208E-05 | 0.00022756 | 110.7667 | 206.425722 |
| Pparg | 0.89762567 | 7.5866E-06 | 0.0001344 | 208.722888 | 388.647544 |
| Rab37 | 0.90069059 | 0.00036427 | 0.00335319 | 60.5841857 | 113.300212 |
| H2-Aa | 0.90127065 | 4.2464E-07 | 1.1056E-05 | 13588.579 | 25380.1633 |
| Il2rg | 0.90330898 | 2.1558E-11 | 1.8093E-09 | 2095.3213 | 3919.40746 |
| Slc25a20 | 0.90947069 | 6.1246E-11 | 4.4883E-09 | 604.343839 | 1134.92229 |
| Xlr4b | 0.91838326 | 0.0032596 | 0.01930954 | 23.8975738 | 45.2859233 |
| Klf11 | 0.91848761 | 2.4769E-10 | 1.5004E-08 | 536.18435 | 1013.77306 |
| Rpp38 | 0.9200524 | 0.00026556 | 0.0026058 | 45.6348833 | 86.5647999 |
| Cd247 | 0.9212351 | 0.00028712 | 0.00277746 | 111.156065 | 210.486501 |
| Top1mt | 0.92514063 | 0.00140899 | 0.01001763 | 49.2363166 | 93.5526533 |
| Gimap9 | 0.92534377 | 2.7976E-05 | 0.00040141 | 133.60592 | 254.018325 |
| Bcl2a1a | 0.92720719 | 2.1828E-05 | 0.00032694 | 149.136503 | 283.993792 |
| Ly6i | 0.93041054 | 7.3511E-07 | 1.7928E-05 | 760.926307 | 1450.51826 |
| Fgf13 | 0.93364526 | 0.00186886 | 0.01257316 | 31.0360588 | 59.1839125 |
| Gimap1 | 0.93485324 | 7.3615E-06 | 0.00013134 | 313.407044 | 599.539353 |
| St14 | 0.93671119 | 0.0074724 | 0.03656606 | 38.0412437 | 72.6035644 |
| Ptprcap | 0.93810016 | 4.4126E-07 | 1.141E-05 | 610.739952 | 1170.50608 |
| Gm47077 | 0.94766344 | 0.00364936 | 0.021144 | 23.926527 | 45.9841284 |
| Tnfrsf18 | 0.95047148 | 8.8449E-07 | 2.0959E-05 | 107.559227 | 208.149946 |
| Zfp566 | 0.95318792 | 0.00170305 | 0.01170398 | 29.9588969 | 58.1055982 |
| Cd2 | 0.95372625 | 0.00033484 | 0.00312244 | 366.169115 | 709.532034 |
| Sh2d2a | 0.96370144 | 0.00177546 | 0.0121006 | 147.621782 | 288.096453 |
| Smyd4 | 0.96681756 | 0.01020099 | 0.04598806 | 47.8168798 | 93.5789927 |
| Plin2 | 0.97336073 | 1.9178E-06 | 4.1514E-05 | 4802.65905 | 9429.38905 |
| Apod | 0.97390613 | 0.00322082 | 0.01915533 | 62.8354975 | 123.475028 |
| Siglech | 0.98206289 | 2.5788E-05 | 0.0003765 | 509.682099 | 1006.7849 |
| Cpm | 0.98305887 | 0.01074108 | 0.04772649 | 30.3738509 | 59.820707 |
| Ephx1 | 0.98685665 | 1.3523E-05 | 0.00021868 | 88.5917713 | 175.478729 |
| St3gal6 | 0.98692743 | 0.00041046 | 0.00368141 | 77.7844127 | 154.613373 |
| Tdrkh | 0.98810133 | 0.00317227 | 0.01895374 | 20.9979815 | 41.5952347 |
| Il4i1 | 0.9898715 | 0.0001136 | 0.00130476 | 296.656864 | 589.328483 |
| Hexim2 | 0.99404878 | 0.00409829 | 0.0230077 | 19.7869825 | 39.2913744 |
| Gimap4 | 0.99419765 | 9.7958E-05 | 0.00115875 | 579.957304 | 1155.64918 |
| Iglv3 | 0.99432407 | 0.0008926 | 0.00695518 | 27.4940073 | 54.7302045 |
| Gm31718 | 0.99493714 | 0.00873823 | 0.04108647 | 26.7777928 | 53.5677926 |
| Gnb1l | 0.99749749 | 0.000747 | 0.00599154 | 27.3218228 | 54.5139605 |
| Cd79a | 1.00135368 | 2.7384E-05 | 0.00039481 | 684.011696 | 1368.94903 |
| Ly6a | 1.00592783 | 1.7199E-07 | 5.0365E-06 | 362.312458 | 728.222303 |
| Sox4 | 1.00825455 | 0.00028531 | 0.00276349 | 59.7839284 | 120.459916 |

| | | | | | |
|---|---|---|---|---|---|
| Gpr83 | 1.0097738 | 0.00019989 | 0.00206242 | 42.6462484 | 85.9710895 |
| Gimap7 | 1.01430355 | 0.00127937 | 0.00928056 | 135.1971 | 273.418532 |
| Cox6a2 | 1.01775163 | 0.00379565 | 0.02174752 | 29.192482 | 59.0428333 |
| Ly6d | 1.01850642 | 7.2455E-08 | 2.3612E-06 | 762.39709 | 1544.09353 |
| Tle6 | 1.02405509 | 0.00671003 | 0.03375916 | 19.9488276 | 40.3088034 |
| Chchd10 | 1.02545755 | 5.5244E-06 | 0.00010348 | 62.0028532 | 126.178974 |
| Ccl4 | 1.02835959 | 0.00187907 | 0.01262489 | 48.4349333 | 99.1669725 |
| Prss30 | 1.03789136 | 0.00053274 | 0.00456034 | 32.3080672 | 66.4015739 |
| Fcer2a | 1.04554342 | 0.00119802 | 0.00879229 | 201.75519 | 416.040581 |
| Hs3st1 | 1.04724857 | 0.00164335 | 0.01137172 | 36.5620561 | 75.7505357 |
| Il27ra | 1.04772122 | 5.9615E-08 | 1.964E-06 | 123.937524 | 256.226114 |
| Serpinb6b | 1.05345637 | 0.00374243 | 0.02153861 | 51.064831 | 106.289248 |
| Acp5 | 1.05412333 | 2.4102E-13 | 3.2043E-11 | 733.193513 | 1522.67156 |
| Pmaip1 | 1.05680999 | 6.8085E-06 | 0.00012309 | 1427.20496 | 2969.25226 |
| 9130019P16F | 1.06271221 | 0.00071405 | 0.00579537 | 24.7207828 | 51.8670869 |
| Fcgr4 | 1.0663025 | 2.1912E-08 | 8.2092E-07 | 352.419966 | 737.996089 |
| Gpat3 | 1.06701828 | 0.00409768 | 0.0230077 | 18.7644055 | 39.2429534 |
| Gm14161 | 1.06948125 | 0.00712616 | 0.03537881 | 28.8411644 | 60.7700053 |
| Spic | 1.07023002 | 3.4595E-05 | 0.00048212 | 371.036393 | 779.605663 |
| Hrh1 | 1.07129062 | 0.00103666 | 0.00786152 | 72.9260884 | 153.414987 |
| Klrc1 | 1.07149466 | 0.00010803 | 0.00125811 | 114.315532 | 240.662829 |
| Cd3g | 1.07224418 | 0.00065468 | 0.00540995 | 129.908694 | 273.13247 |
| 6530413G14I | 1.07385947 | 1.3214E-10 | 8.7067E-09 | 168.908422 | 355.514949 |
| Gm33104 | 1.07519455 | 0.00202656 | 0.01335895 | 18.9962009 | 40.1677698 |
| Gbp8 | 1.07582458 | 0.00012276 | 0.00139016 | 63.257092 | 133.678359 |
| Maged1 | 1.07589075 | 0.00046147 | 0.00405861 | 29.7616393 | 62.8398863 |
| Treml4 | 1.08105252 | 4.9236E-07 | 1.2537E-05 | 3816.24824 | 8073.50123 |
| H2-Ab1 | 1.0855902 | 2.9051E-08 | 1.0516E-06 | 26838.511 | 56958.4589 |
| Gm37126 | 1.08783087 | 0.0038945 | 0.02217019 | 16.7988218 | 35.7920291 |
| Gm19585 | 1.09358853 | 0.00454111 | 0.02491642 | 66.4135637 | 142.04392 |
| Klra1 | 1.09662695 | 0.00050351 | 0.00435727 | 23.48317 | 50.3741641 |
| Gimap6 | 1.09988068 | 3.1216E-09 | 1.4474E-07 | 496.43448 | 1064.3912 |
| Jchain | 1.10027575 | 0.0087833 | 0.04120909 | 714.604223 | 1532.01412 |
| Cd300c | 1.10983006 | 2.7674E-05 | 0.00039822 | 54.9529701 | 118.840889 |
| Xkrx | 1.11109178 | 0.00362988 | 0.02106283 | 47.9871068 | 103.65883 |
| Cma1 | 1.11769909 | 0.00111573 | 0.00833496 | 136.84442 | 297.269395 |
| Fcmr | 1.12358715 | 0.00014068 | 0.0015415 | 383.239193 | 834.608891 |
| Lat | 1.12764876 | 0.00114356 | 0.00848803 | 108.01725 | 236.244335 |
| Cd8b1 | 1.1292576 | 8.4813E-07 | 2.0321E-05 | 330.34307 | 722.831523 |
| Ighd | 1.1296785 | 0.0001406 | 0.0015415 | 478.37189 | 1046.29953 |
| Kcnj8 | 1.1330298 | 0.00347186 | 0.02027118 | 69.7500583 | 153.354779 |
| Trbc2 | 1.14014372 | 2.6435E-06 | 5.4701E-05 | 190.284431 | 419.551961 |
| Cd8a | 1.14763312 | 5.4439E-08 | 1.8174E-06 | 290.898183 | 644.78132 |
| Znrd1as | 1.15486341 | 0.00273152 | 0.01692889 | 16.9553656 | 37.6784738 |
| 2010300F17F | 1.15512213 | 0.00080598 | 0.00637273 | 28.4294406 | 63.3615536 |
| Car2 | 1.15619772 | 4.4276E-05 | 0.00059178 | 44.1347599 | 98.6016452 |

| | | | | | |
|---|---|---|---|---|---|
| Phactr3 | 1.1642857 | 0.00897312 | 0.04181239 | 17.9952488 | 40.54664 |
| Cd160 | 1.16676915 | 0.00051814 | 0.00446842 | 37.0785934 | 83.4976546 |
| Ocstamp | 1.16815131 | 1.5844E-06 | 3.4814E-05 | 190.569204 | 428.497401 |
| Lck | 1.16859281 | 1.004E-05 | 0.00017024 | 603.902884 | 1357.96325 |
| Adgrl3 | 1.17803227 | 0.00501964 | 0.02700935 | 14.1986607 | 32.2335863 |
| 7530414M10 | 1.18102162 | 0.00825331 | 0.03927503 | 11.0157163 | 24.9005777 |
| Gm4956 | 1.18293887 | 0.00339835 | 0.01995055 | 36.0230564 | 82.1942157 |
| Cd209d | 1.19816959 | 5.489E-05 | 0.00071271 | 429.293146 | 985.44354 |
| Gm15157 | 1.20552592 | 0.00364595 | 0.02113975 | 14.2368996 | 33.1039139 |
| Cd28 | 1.21843714 | 4.6081E-05 | 0.00061156 | 48.8025329 | 113.551535 |
| Bcl11b | 1.2385463 | 0.00054537 | 0.00465253 | 55.3950511 | 130.665368 |
| Cd3d | 1.23948726 | 1.4232E-08 | 5.5656E-07 | 116.421438 | 274.823884 |
| Pxdc1 | 1.24109188 | 1.5487E-06 | 3.4165E-05 | 65.6587293 | 154.974781 |
| Sit1 | 1.24151178 | 0.01101813 | 0.04852748 | 30.4891685 | 71.932589 |
| Gm36329 | 1.24957284 | 0.00750985 | 0.03671345 | 11.7771368 | 28.2136683 |
| Iglv1 | 1.25662213 | 0.00513568 | 0.0275646 | 312.541583 | 746.653847 |
| Dapl1 | 1.25869854 | 5.384E-05 | 0.0006997 | 19.6849195 | 47.0435259 |
| Gm49463 | 1.26324692 | 0.00123436 | 0.00901494 | 23.8408199 | 57.3027333 |
| Rpusd2 | 1.26967725 | 0.00799871 | 0.03845267 | 10.2896811 | 24.7723043 |
| Gm6637 | 1.29071794 | 0.00025521 | 0.0025207 | 68.2415333 | 167.407589 |
| Trbv16 | 1.29098961 | 0.01146062 | 0.04989072 | 11.1290333 | 27.3069761 |
| Rnf180 | 1.29607931 | 0.00347173 | 0.02027118 | 9.45301665 | 23.1593654 |
| Kcng2 | 1.3015323 | 0.00140182 | 0.00997608 | 14.7916053 | 36.7259638 |
| Tsc22d1 | 1.31166596 | 2.4224E-05 | 0.00035677 | 146.335931 | 363.437883 |
| Trac | 1.33124922 | 3.2941E-07 | 8.8214E-06 | 167.170399 | 420.638839 |
| Pdcd1lg2 | 1.33341728 | 0.00202918 | 0.01337034 | 17.7023452 | 44.6572751 |
| Cd209b | 1.33534915 | 0.00118549 | 0.00872592 | 19.4118319 | 49.1255782 |
| Cd200 | 1.35219702 | 1.9644E-07 | 5.6213E-06 | 34.1361388 | 87.2050163 |
| Tcrg-C2 | 1.36440243 | 0.00822111 | 0.03920182 | 8.26159272 | 21.1299875 |
| Lefty1 | 1.3657779 | 9.1752E-06 | 0.00015807 | 36.4747746 | 94.2501016 |
| Cxcr6 | 1.37563323 | 0.00013763 | 0.0015158 | 48.0836947 | 124.63623 |
| St6galnac3 | 1.38482421 | 0.00434531 | 0.02407955 | 8.98164615 | 23.4248147 |
| Lilra5 | 1.38494523 | 4.9965E-09 | 2.2474E-07 | 125.0726 | 326.972313 |
| 5730409E04F | 1.38957743 | 0.00687899 | 0.03440182 | 12.6180806 | 33.216643 |
| Nbl1 | 1.39636762 | 0.00925326 | 0.04275968 | 12.8005611 | 33.778792 |
| Xlr4c | 1.40770999 | 0.00905932 | 0.04209656 | 6.15448329 | 16.2903209 |
| Ifitm10 | 1.42489971 | 6.834E-06 | 0.0001234 | 62.7814097 | 168.89972 |
| Crtam | 1.42657153 | 0.00909316 | 0.04221464 | 19.1897943 | 51.6065594 |
| Ndrg2 | 1.43596346 | 0.00584183 | 0.03040229 | 9.29664221 | 25.239964 |
| Spint1 | 1.4366838 | 0.00218346 | 0.01416325 | 17.5673796 | 47.5704986 |
| Pacsin1 | 1.47141247 | 0.00036811 | 0.0033844 | 53.5845667 | 148.423638 |
| Rho | 1.47496809 | 0.0095186 | 0.04363685 | 8.83199783 | 24.3841696 |
| Bhlhb9 | 1.47737284 | 3.632E-05 | 0.00050288 | 19.8500654 | 55.3529491 |
| Ifngas1 | 1.49547181 | 0.00581689 | 0.03033222 | 6.54792208 | 18.5796331 |
| Cd209e | 1.49728474 | 6.5902E-07 | 1.631E-05 | 132.206825 | 373.278453 |
| Ube2cbp | 1.52012914 | 0.00768309 | 0.03737795 | 7.48685133 | 21.3453124 |

| | | | | | |
|---|---|---|---|---|---|
| Iglc1 | 1.5275203 | 0.00059651 | 0.00501478 | 311.74709 | 898.678519 |
| Chrnb1 | 1.54965992 | 0.0072606 | 0.03583311 | 5.80988854 | 17.0950249 |
| Clec4g | 1.58278122 | 0.00013207 | 0.00147191 | 16.7539913 | 50.3355577 |
| Il20rb | 1.67962277 | 0.00415057 | 0.02323177 | 5.04442635 | 16.2386883 |
| Gm6969 | 1.69026899 | 0.00061438 | 0.00513913 | 7.52165497 | 24.0876782 |
| Gm45669 | 1.69929204 | 0.00029288 | 0.00282423 | 10.6488273 | 34.6741351 |
| Il5ra | 1.71736547 | 0.00183162 | 0.01238919 | 5.95783117 | 19.4253441 |
| Faah | 1.72875007 | 8.023E-06 | 0.0001413 | 19.207947 | 63.7343834 |
| Cym | 1.7464002 | 0.00093892 | 0.00721869 | 15.469889 | 52.2263358 |
| Igkv4-61 | 1.89544181 | 0.00920382 | 0.04255742 | 4.64558408 | 17.2333538 |
| Fbxo17 | 1.91248299 | 0.00058839 | 0.00495633 | 6.77023318 | 25.6862686 |
| Pomgnt2 | 1.99117047 | 0.00134005 | 0.00961396 | 4.32202496 | 17.1961787 |
| Pvrig | 2.02503505 | 0.00043751 | 0.00388226 | 4.87730894 | 19.7871706 |
| AC091309.5 | 2.18804591 | 1.4781E-05 | 0.00023498 | 9.78480144 | 44.8811865 |
| Tnfrsf25 | 2.24475761 | 0.00103106 | 0.00782305 | 4.53271419 | 21.5988809 |
| Icos | 2.2683947 | 0.00024771 | 0.00245793 | 7.27527991 | 34.8675568 |
| Igkv10-96 | 2.72882601 | 0.00047626 | 0.00415977 | 19.3967065 | 128.405487 |
| Ighv2-2 | 3.03321295 | 0.00025406 | 0.002511 | 2.78410206 | 22.8249947 |
| Igkv12-44 | 3.23385232 | 0.00059199 | 0.00498237 | 46.9587858 | 441.513564 |
| Igkv10-94 | 3.74095788 | 0.00020959 | 0.00214922 | 6.85961741 | 91.7155086 |
| Slc15a2 | 4.00326809 | 0.00142208 | 0.0100964 | 2.62891571 | 41.8741505 |
| Igkv19-93 | 4.26101595 | 1.2204E-06 | 2.7737E-05 | 15.7466071 | 301.636311 |
| Ighv1-75 | 4.48824878 | 4.3203E-08 | 1.4846E-06 | 4.98199223 | 112.461671 |
