## Supplemental Table 7 for "Vagal Signaling Decline in Age-related Macular Degeneration Drives Spleen-Dependent Retinal Inflammation"

| Gene | Log Fold Change | p-value | padj |
|---|---|---|---|
| Cst3 | 1.8 | 9.67E-25 | 3.12E-20 |
| Fkbp5 | 1.4 | 5.00E-53 | 1.61E-48 |
| Rnf169 | 1.25 | 1.81E-38 | 5.83E-34 |
| Ctss | 1.04 | 7.40E-14 | 2.39E-09 |
| Srgn | 1.02 | 7.78E-15 | 2.51E-10 |
| Sla | 0.95 | 6.47E-33 | 2.09E-28 |
| Slc24a3 | 0.94 | 5.66E-25 | 1.83E-20 |
| C1qb | 0.94 | 5.52E-11 | 1.78E-06 |
| Serinc3 | 0.93 | 5.84E-23 | 1.89E-18 |
| Sparc | 0.91 | 1.71E-16 | 5.53E-12 |
| Cd300lf | 0.86 | 2.63E-24 | 8.50E-20 |
| Cacnb2 | 0.83 | 1.84E-17 | 5.95E-13 |
| Rnf180 | 0.77 | 9.15E-19 | 2.95E-14 |
| Ier3 | 0.72 | 1.39E-10 | 4.48E-06 |
| B2m | 0.72 | 4.59E-08 | 1.48E-03 |
| Rps29 | 0.72 | 7.02E-07 | 2.27E-02 |
| Klf13 | 0.7 | 5.55E-20 | 1.79E-15 |
| Siglech | 0.7 | 1.42E-19 | 4.57E-15 |
| C1qc | 0.7 | 4.20E-07 | 1.36E-02 |
| B4galt1 | 0.69 | 9.99E-18 | 3.23E-13 |
| Laptm5 | 0.69 | 4.65E-11 | 1.50E-06 |
| Sort1 | 0.68 | 2.67E-26 | 8.62E-22 |
| Trf | 0.67 | 4.16E-14 | 1.34E-09 |
| Jun | 0.67 | 1.07E-07 | 3.46E-03 |
| Slc38a2 | 0.66 | 1.71E-19 | 5.51E-15 |
| Wnk1 | 0.66 | 6.62E-15 | 2.14E-10 |
| Fcgr3 | 0.66 | 2.97E-10 | 9.60E-06 |
| Cd81 | 0.66 | 3.56E-09 | 1.15E-04 |
| Gpr65 | 0.65 | 4.36E-21 | 1.41E-16 |
| Rbm26 | 0.65 | 2.14E-16 | 6.92E-12 |
| Rps27 | 0.65 | 4.72E-08 | 1.52E-03 |
| Ccnd3 | 0.64 | 9.59E-16 | 3.10E-11 |
| H2-D1 | 0.64 | 1.09E-09 | 3.51E-05 |
| Ogfrl1 | 0.63 | 9.43E-23 | 3.05E-18 |
| Gm46224 | 0.63 | 2.65E-16 | 8.55E-12 |
| Dst | 0.62 | 1.17E-13 | 3.78E-09 |
| Slfn2 | 0.61 | 3.34E-12 | 1.08E-07 |
| Snx24 | 0.61 | 1.28E-11 | 4.14E-07 |
| Cited2 | 0.6 | 8.36E-13 | 2.70E-08 |
| 01-Mar | 0.6 | 7.21E-11 | 2.33E-06 |
| Prkcb | 0.59 | 4.69E-11 | 1.52E-06 |
| Chka | 0.58 | 4.32E-10 | 1.40E-05 |
| Itm2b | 0.58 | 8.30E-07 | 2.68E-02 |
| Tlr7 | 0.57 | 3.77E-16 | 1.22E-11 |
| Il6ra | 0.57 | 1.91E-15 | 6.18E-11 |

| | | | |
|---|---|---|---|
| Pik3r1 | 0.57 | 3.46E-12 | 1.12E-07 |
| Mir142hg | 0.57 | 2.27E-09 | 7.32E-05 |
| Cd33 | 0.56 | 9.37E-17 | 3.03E-12 |
| Peli1 | 0.56 | 5.75E-10 | 1.86E-05 |
| Lyn | 0.55 | 1.07E-10 | 3.46E-06 |
| Gtdc1 | 0.55 | 1.23E-10 | 3.98E-06 |
| Ier2 | 0.55 | 8.93E-07 | 2.88E-02 |
| P2ry12 | 0.54 | 1.95E-09 | 6.29E-05 |
| Cd53 | 0.54 | 3.08E-09 | 9.93E-05 |
| Ssh2 | 0.54 | 3.93E-08 | 1.27E-03 |
| Med12l | 0.53 | 5.06E-15 | 1.63E-10 |
| Enah | 0.52 | 6.30E-21 | 2.04E-16 |
| Tcf7l2 | 0.52 | 3.62E-13 | 1.17E-08 |
| Chd2 | 0.52 | 3.06E-12 | 9.89E-08 |
| Adap2 | 0.52 | 8.32E-10 | 2.69E-05 |
| Glul | 0.52 | 9.28E-09 | 3.00E-04 |
| Mbnl1 | 0.52 | 1.20E-08 | 3.87E-04 |
| Rbbp6 | 0.51 | 3.73E-12 | 1.20E-07 |
| Selenop | 0.51 | 8.66E-12 | 2.80E-07 |
| Mtdh | 0.51 | 1.82E-08 | 5.87E-04 |
| Ccdc86 | 0.5 | 3.71E-12 | 1.20E-07 |
| Irs2 | 0.49 | 2.09E-18 | 6.74E-14 |
| Gm10790 | 0.49 | 1.21E-16 | 3.89E-12 |
| Slc12a9 | 0.49 | 1.21E-15 | 3.90E-11 |
| Ptpn1 | 0.49 | 4.56E-09 | 1.47E-04 |
| Fcho2 | 0.47 | 5.22E-10 | 1.69E-05 |
| St3gal6 | 0.47 | 3.94E-09 | 1.27E-04 |
| Runx1 | 0.47 | 1.57E-07 | 5.08E-03 |
| Cd86 | 0.47 | 2.64E-07 | 8.51E-03 |
| Coro2a | 0.46 | 3.83E-15 | 1.24E-10 |
| Tm6sf1 | 0.46 | 2.23E-10 | 7.18E-06 |
| Gnl3 | 0.46 | 1.84E-09 | 5.94E-05 |
| Irf2bp2 | 0.46 | 1.31E-08 | 4.24E-04 |
| Klf9 | 0.45 | 2.51E-21 | 8.10E-17 |
| Eps8 | 0.45 | 3.68E-11 | 1.19E-06 |
| Kdm7a | 0.45 | 2.81E-09 | 9.06E-05 |
| Sdc4 | 0.45 | 3.47E-09 | 1.12E-04 |
| Txnip | 0.45 | 5.97E-09 | 1.93E-04 |
| Atp8a1 | 0.45 | 2.68E-07 | 8.67E-03 |
| Tg | 0.43 | 4.33E-18 | 1.40E-13 |
| Gm43305 | 0.43 | 7.02E-18 | 2.27E-13 |
| Il15ra | 0.43 | 4.65E-15 | 1.50E-10 |
| Abtb2 | 0.43 | 1.18E-14 | 3.80E-10 |
| Tasor2 | 0.43 | 1.07E-11 | 3.44E-07 |
| Csf1r | 0.43 | 9.27E-07 | 2.99E-02 |
| 0610040J01R | 0.42 | 2.69E-09 | 8.69E-05 |

| | | | |
|---|---|---|---|
| Larp1 | 0.42 | 6.86E-09 | 2.21E-04 |
| Ube2h | 0.42 | 7.72E-09 | 2.49E-04 |
| Epsti1 | 0.42 | 1.19E-08 | 3.83E-04 |
| Ifngr1 | 0.42 | 2.13E-08 | 6.89E-04 |
| Kdm6b | 0.41 | 8.23E-09 | 2.66E-04 |
| Lrrfip1 | 0.41 | 3.15E-07 | 1.02E-02 |
| Gns | 0.41 | 3.93E-07 | 1.27E-02 |
| Chn2 | 0.41 | 7.86E-07 | 2.54E-02 |
| Pten | 0.4 | 2.61E-08 | 8.43E-04 |
| Pik3cg | 0.39 | 2.89E-11 | 9.35E-07 |
| Il4ra | 0.39 | 8.79E-09 | 2.84E-04 |
| Nrip1 | 0.39 | 1.37E-07 | 4.43E-03 |
| Stat3 | 0.39 | 1.71E-07 | 5.52E-03 |
| Rbm25 | 0.39 | 6.29E-07 | 2.03E-02 |
| Zeb1 | 0.39 | 6.77E-07 | 2.19E-02 |
| Man2b1 | 0.39 | 8.74E-07 | 2.82E-02 |
| Cep152 | 0.38 | 1.09E-11 | 3.52E-07 |
| Il17ra | 0.38 | 2.97E-08 | 9.59E-04 |
| Pacsin2 | 0.38 | 5.54E-08 | 1.79E-03 |
| Adora3 | 0.37 | 4.09E-14 | 1.32E-09 |
| Zbtb16 | 0.37 | 1.43E-13 | 4.61E-09 |
| Gmip | 0.37 | 4.80E-10 | 1.55E-05 |
| Mt2 | 0.37 | 7.60E-10 | 2.46E-05 |
| Peli2 | 0.37 | 2.04E-09 | 6.58E-05 |
| Ifih1 | 0.37 | 6.33E-09 | 2.04E-04 |
| Ripor2 | 0.37 | 1.30E-06 | 4.20E-02 |
| Il18rap | 0.36 | 2.85E-15 | 9.21E-11 |
| Mlxip | 0.36 | 9.67E-09 | 3.12E-04 |
| Dram2 | 0.36 | 1.81E-08 | 5.84E-04 |
| Adap2os | 0.36 | 6.72E-08 | 2.17E-03 |
| Lrrk1 | 0.36 | 3.26E-07 | 1.05E-02 |
| Bmf | 0.35 | 7.79E-18 | 2.52E-13 |
| Abhd15 | 0.35 | 2.16E-17 | 6.99E-13 |
| Ltc4s | 0.34 | 2.78E-10 | 8.96E-06 |
| Gbp7 | 0.34 | 6.06E-10 | 1.96E-05 |
| Hs3st3b1 | 0.34 | 3.39E-09 | 1.10E-04 |
| Ptbp2 | 0.34 | 1.13E-07 | 3.64E-03 |
| Wdr43 | 0.34 | 5.74E-07 | 1.85E-02 |
| Gm42031 | 0.33 | 1.48E-14 | 4.77E-10 |
| Lag3 | 0.32 | 6.71E-13 | 2.17E-08 |
| Cntrl | 0.32 | 4.89E-08 | 1.58E-03 |
| Ttc39b | 0.32 | 8.66E-08 | 2.80E-03 |
| Scamp1 | 0.32 | 9.79E-07 | 3.16E-02 |
| Spata13 | 0.32 | 1.18E-06 | 3.82E-02 |
| Il6 | 0.31 | 2.46E-09 | 7.95E-05 |
| Nfam1 | 0.31 | 7.74E-09 | 2.50E-04 |

| | | | |
|---|---|---|---|
| Tns1 | 0.31 | 4.06E-08 | 1.31E-03 |
| Map3k8 | 0.31 | 4.60E-07 | 1.49E-02 |
| Slc7a5 | 0.31 | 4.76E-07 | 1.54E-02 |
| Tagap | 0.3 | 1.29E-10 | 4.16E-06 |
| Il1r2 | 0.3 | 4.21E-10 | 1.36E-05 |
| Rap1gap2 | 0.3 | 1.11E-08 | 3.59E-04 |
| Afap1l1 | 0.3 | 3.15E-08 | 1.02E-03 |
| Pstpip2 | 0.3 | 2.50E-07 | 8.07E-03 |
| Heatr1 | 0.3 | 6.05E-07 | 1.95E-02 |
| Yars | 0.27 | 1.12E-06 | 3.63E-02 |
| Tspan13 | 0.26 | 3.71E-07 | 1.20E-02 |
| Timm9 | 0.26 | 6.57E-07 | 2.12E-02 |
| Gm22146 | 0.25 | 3.80E-11 | 1.23E-06 |
| 1600020E01F | 0.25 | 1.86E-07 | 6.01E-03 |
| Utp4 | 0.25 | 4.29E-07 | 1.38E-02 |
| Nfil3 | 0.25 | 1.32E-06 | 4.27E-02 |
| Rab37 | 0.23 | 3.59E-11 | 1.16E-06 |
| Pde1b | 0.23 | 1.51E-08 | 4.87E-04 |
| Cd82 | 0.23 | 1.63E-08 | 5.27E-04 |
| Ctnnal1 | 0.23 | 7.82E-08 | 2.52E-03 |
| Nuak2 | 0.23 | 3.13E-07 | 1.01E-02 |
| E230001N04F | 0.22 | 1.90E-11 | 6.13E-07 |
| Sgip1 | 0.22 | 1.41E-06 | 4.56E-02 |
| Gm49662 | 0.21 | 7.13E-09 | 2.30E-04 |
| Gm31718 | 0.21 | 2.57E-07 | 8.31E-03 |
| Atad2 | -0.22 | 3.22E-07 | 1.04E-02 |
| Rho | -0.23 | 5.93E-09 | 1.92E-04 |
| Jam2 | -0.23 | 5.09E-08 | 1.64E-03 |
| Nagk | -0.23 | 1.27E-07 | 4.12E-03 |
| Prr5l | -0.23 | 2.12E-07 | 6.83E-03 |
| Fblim1 | -0.24 | 3.70E-07 | 1.19E-02 |
| Vav2 | -0.24 | 7.82E-07 | 2.53E-02 |
| Speg | -0.25 | 2.57E-08 | 8.29E-04 |
| Rasa3 | -0.26 | 4.84E-07 | 1.56E-02 |
| Nes | -0.27 | 1.06E-08 | 3.41E-04 |
| Hk3 | -0.27 | 1.89E-08 | 6.10E-04 |
| Cd36 | -0.28 | 4.16E-08 | 1.34E-03 |
| Gas2l3 | -0.28 | 5.68E-08 | 1.83E-03 |
| Hyou1 | -0.3 | 1.74E-07 | 5.60E-03 |
| S100a6 | -0.31 | 4.35E-08 | 1.40E-03 |
| Kcnma1 | -0.32 | 5.05E-07 | 1.63E-02 |
| Npnt | -0.33 | 9.40E-09 | 3.04E-04 |
| Bcl2 | -0.33 | 1.32E-07 | 4.25E-03 |
| Taco1 | -0.33 | 5.72E-07 | 1.85E-02 |
| Arhgap10 | -0.34 | 1.74E-10 | 5.61E-06 |
| Rab7b | -0.35 | 1.77E-10 | 5.70E-06 |

| | | | |
|---|---|---|---|
| Zmynd8 | -0.35 | 1.61E-07 | 5.20E-03 |
| Gm2245 | -0.35 | 2.33E-07 | 7.53E-03 |
| Plxna4 | -0.36 | 1.35E-06 | 4.37E-02 |
| Bcar3 | -0.37 | 5.49E-10 | 1.77E-05 |
| Creld2 | -0.37 | 2.65E-09 | 8.56E-05 |
| Lgals1 | -0.37 | 4.45E-09 | 1.44E-04 |
| Galnt7 | -0.37 | 5.62E-07 | 1.81E-02 |
| Etl4 | -0.38 | 1.04E-10 | 3.37E-06 |
| Atxn1 | -0.38 | 3.47E-09 | 1.12E-04 |
| Anxa5 | -0.38 | 9.89E-08 | 3.19E-03 |
| Map4 | -0.38 | 2.06E-07 | 6.64E-03 |
| Fmn1 | -0.38 | 5.77E-07 | 1.86E-02 |
| Phkb | -0.39 | 4.69E-09 | 1.51E-04 |
| Sgms1 | -0.39 | 5.98E-08 | 1.93E-03 |
| Fabp5 | -0.39 | 2.70E-07 | 8.73E-03 |
| Xylt1 | -0.39 | 4.51E-07 | 1.46E-02 |
| Ppm1h | -0.39 | 4.88E-07 | 1.58E-02 |
| Macf1 | -0.39 | 7.54E-07 | 2.44E-02 |
| Tpi1 | -0.4 | 3.21E-07 | 1.04E-02 |
| Nuak1 | -0.41 | 7.32E-11 | 2.36E-06 |
| Tent5c | -0.41 | 1.36E-08 | 4.39E-04 |
| Iqgap1 | -0.41 | 4.20E-08 | 1.36E-03 |
| Capg | -0.41 | 5.14E-08 | 1.66E-03 |
| Vps13c | -0.42 | 2.84E-08 | 9.17E-04 |
| Olfr111 | -0.43 | 1.15E-12 | 3.70E-08 |
| Rai14 | -0.43 | 1.12E-07 | 3.61E-03 |
| Supt3 | -0.44 | 6.47E-12 | 2.09E-07 |
| Vim | -0.44 | 1.05E-07 | 3.37E-03 |
| Specc1 | -0.45 | 1.27E-07 | 4.09E-03 |
| Lair1 | -0.45 | 2.11E-07 | 6.82E-03 |
| Slc9a9 | -0.45 | 2.87E-07 | 9.26E-03 |
| Tmcc3 | -0.45 | 6.56E-07 | 2.12E-02 |
| Arhgap22 | -0.46 | 1.45E-09 | 4.69E-05 |
| Cadm1 | -0.46 | 2.35E-07 | 7.57E-03 |
| Itga6 | -0.47 | 8.15E-09 | 2.63E-04 |
| Dock2 | -0.47 | 4.91E-08 | 1.59E-03 |
| Pdgfb | -0.48 | 2.00E-10 | 6.45E-06 |
| Dynll1 | -0.48 | 8.99E-09 | 2.90E-04 |
| Rasgef1b | -0.5 | 1.40E-11 | 4.53E-07 |
| Cd72 | -0.5 | 3.74E-09 | 1.21E-04 |
| Cybb | -0.51 | 5.37E-08 | 1.74E-03 |
| Nrp1 | -0.52 | 9.72E-13 | 3.14E-08 |
| Mitf | -0.52 | 2.76E-07 | 8.92E-03 |
| Slc7a8 | -0.55 | 2.03E-11 | 6.55E-07 |
| Fam20c | -0.55 | 2.33E-11 | 7.52E-07 |
| Wwox | -0.55 | 2.06E-08 | 6.65E-04 |

| | | | |
|---|---|---|---|
| Gpr34 | -0.56 | 1.34E-12 | 4.34E-08 |
| Lhfpl2 | -0.57 | 2.70E-14 | 8.72E-10 |
| Apbb2 | -0.58 | 4.40E-10 | 1.42E-05 |
| Hivep2 | -0.59 | 2.46E-12 | 7.94E-08 |
| Hsp90b1 | -0.59 | 9.27E-07 | 2.99E-02 |
| Asph | -0.6 | 2.15E-14 | 6.94E-10 |
| Cd180 | -0.6 | 4.31E-14 | 1.39E-09 |
| Tbxas1 | -0.6 | 4.65E-11 | 1.50E-06 |
| Pdia4 | -0.61 | 3.72E-14 | 1.20E-09 |
| Zeb2 | -0.61 | 1.42E-09 | 4.60E-05 |
| Pdia6 | -0.61 | 1.21E-08 | 3.90E-04 |
| Apoe | -0.61 | 7.19E-07 | 2.32E-02 |
| Sdf2l1 | -0.62 | 6.35E-12 | 2.05E-07 |
| Pitpnc1 | -0.64 | 1.06E-11 | 3.41E-07 |
| Calr | -0.65 | 1.34E-07 | 4.33E-03 |
| Immp2l | -0.66 | 1.38E-12 | 4.47E-08 |
| Abhd12 | -0.66 | 3.63E-12 | 1.17E-07 |
| Dnmt3a | -0.67 | 6.33E-20 | 2.04E-15 |
| Mir99ahg | -0.68 | 6.86E-11 | 2.21E-06 |
| Myo1f | -0.7 | 1.43E-17 | 4.63E-13 |
| Lgals3 | -0.75 | 9.99E-12 | 3.23E-07 |
| Bin1 | -0.79 | 2.95E-19 | 9.51E-15 |
| Lrmda | -0.82 | 5.25E-17 | 1.69E-12 |
| Nav3 | -0.82 | 1.83E-14 | 5.92E-10 |
| Lpl | -0.82 | 6.93E-12 | 2.24E-07 |
| Hspa5 | -0.82 | 6.17E-11 | 1.99E-06 |
| Fn1 | -0.86 | 1.80E-12 | 5.80E-08 |
| Spp1 | -1.14 | 1.03E-14 | 3.33E-10 |
