## Supplemental Table 8 for "Vagal Signaling Decline in Age-related Macular Degeneration Drives Spleen-Dependent Retinal Inflammation"

| Gene | Log Fold Change | p-value | padj |
|---|---|---|---|
| Cst3 | 1.81 | 7.22E-11 | 2.33E-06 |
| Ctss | 1.15 | 3.45E-08 | 1.11E-03 |
| C1qb | 1.14 | 3.38E-07 | 1.09E-02 |
| Rnf169 | 1.1 | 4.24E-17 | 1.37E-12 |
| Fkbp5 | 1.09 | 7.16E-17 | 2.31E-12 |
| Srgn | 1.01 | 3.42E-08 | 1.11E-03 |
| Jun | 0.93 | 2.25E-07 | 7.26E-03 |
| Sparc | 0.91 | 6.96E-09 | 2.25E-04 |
| Itm2b | 0.91 | 1.09E-07 | 3.52E-03 |
| Slc24a3 | 0.88 | 1.25E-12 | 4.05E-08 |
| H2-D1 | 0.87 | 1.08E-08 | 3.50E-04 |
| Rnf180 | 0.86 | 1.82E-12 | 5.89E-08 |
| Ier3 | 0.86 | 9.04E-08 | 2.92E-03 |
| Sla | 0.85 | 1.37E-14 | 4.43E-10 |
| Cd300lf | 0.84 | 3.61E-12 | 1.16E-07 |
| Klf13 | 0.8 | 1.40E-14 | 4.53E-10 |
| B4galt1 | 0.76 | 1.06E-11 | 3.41E-07 |
| Ier2 | 0.74 | 1.28E-06 | 4.13E-02 |
| Cmss1 | 0.73 | 3.17E-09 | 1.02E-04 |
| Slfn2 | 0.71 | 1.43E-09 | 4.63E-05 |
| H2-K1 | 0.71 | 4.41E-07 | 1.42E-02 |
| Ccnd3 | 0.69 | 9.50E-10 | 3.07E-05 |
| Trf | 0.69 | 1.90E-08 | 6.14E-04 |
| Slc15a3 | 0.69 | 3.16E-07 | 1.02E-02 |
| Snx24 | 0.68 | 1.97E-08 | 6.36E-04 |
| Serinc3 | 0.67 | 1.87E-07 | 6.03E-03 |
| Sort1 | 0.65 | 1.30E-12 | 4.21E-08 |
| Rbm47 | 0.65 | 4.25E-08 | 1.37E-03 |
| Gpr65 | 0.64 | 1.60E-11 | 5.18E-07 |
| Gm46224 | 0.64 | 2.58E-10 | 8.33E-06 |
| Cited2 | 0.64 | 3.93E-08 | 1.27E-03 |
| Il6ra | 0.62 | 3.47E-10 | 1.12E-05 |
| Rbbp6 | 0.62 | 1.25E-09 | 4.03E-05 |
| Irf2bp2 | 0.62 | 3.09E-08 | 9.97E-04 |
| Runx1 | 0.62 | 5.94E-07 | 1.92E-02 |
| Chka | 0.62 | 1.00E-06 | 3.23E-02 |
| Cd33 | 0.61 | 7.65E-12 | 2.47E-07 |
| Gtdc1 | 0.61 | 7.37E-07 | 2.38E-02 |
| Lrrfip1 | 0.6 | 5.23E-08 | 1.69E-03 |
| Siglech | 0.59 | 2.00E-07 | 6.44E-03 |
| Ogfrl1 | 0.58 | 1.99E-10 | 6.43E-06 |
| Rbm26 | 0.58 | 1.53E-08 | 4.93E-04 |
| Tlr7 | 0.57 | 9.93E-09 | 3.21E-04 |
| Epsti1 | 0.57 | 2.13E-08 | 6.88E-04 |
| Fcho2 | 0.57 | 4.81E-08 | 1.55E-03 |

| | | | |
|---|---|---|---|
| Dst | 0.57 | 4.42E-07 | 1.43E-02 |
| Irs2 | 0.56 | 1.50E-13 | 4.83E-09 |
| Adap2 | 0.56 | 2.51E-07 | 8.10E-03 |
| Filip1l | 0.55 | 8.09E-10 | 2.61E-05 |
| Gns | 0.55 | 1.44E-06 | 4.64E-02 |
| Man2b1 | 0.53 | 1.29E-06 | 4.16E-02 |
| Gnl3 | 0.52 | 1.36E-06 | 4.40E-02 |
| Med12l | 0.51 | 7.25E-08 | 2.34E-03 |
| Enah | 0.49 | 1.79E-09 | 5.76E-05 |
| Klf9 | 0.47 | 9.70E-13 | 3.13E-08 |
| Igsf6 | 0.46 | 2.68E-07 | 8.66E-03 |
| Tg | 0.45 | 4.29E-11 | 1.38E-06 |
| Peli2 | 0.45 | 6.19E-08 | 2.00E-03 |
| Coro2a | 0.43 | 1.37E-07 | 4.42E-03 |
| Gm10790 | 0.42 | 9.32E-07 | 3.01E-02 |
| Cdk11b | 0.41 | 7.26E-07 | 2.34E-02 |
| Mt2 | 0.41 | 1.23E-06 | 3.96E-02 |
| Ltc4s | 0.4 | 3.32E-08 | 1.07E-03 |
| Cep152 | 0.39 | 1.91E-07 | 6.16E-03 |
| Zbtb16 | 0.38 | 3.45E-08 | 1.11E-03 |
| Bmf | 0.37 | 7.74E-11 | 2.50E-06 |
| Gm43305 | 0.37 | 6.88E-07 | 2.22E-02 |
| Il18rap | 0.34 | 1.37E-07 | 4.43E-03 |
| Tspan13 | 0.34 | 6.28E-07 | 2.03E-02 |
| Abhd15 | 0.29 | 1.24E-06 | 4.01E-02 |
| Cry1 | 0.28 | 9.10E-07 | 2.94E-02 |
| Cd82 | 0.28 | 1.31E-06 | 4.23E-02 |
| Def6 | 0.27 | 1.07E-06 | 3.46E-02 |
| E230001N04I | 0.24 | 7.22E-08 | 2.33E-03 |
| Pf4 | -0.33 | 8.27E-08 | 2.67E-03 |
| Supt3 | -0.43 | 7.09E-07 | 2.29E-02 |
| Cd36 | -0.46 | 1.63E-07 | 5.26E-03 |
| Tanc2 | -0.48 | 4.33E-07 | 1.40E-02 |
| Dnmt3a | -0.49 | 3.62E-07 | 1.17E-02 |
| Ubn2 | -0.49 | 1.14E-06 | 3.68E-02 |
| Tnfaip8 | -0.54 | 6.73E-07 | 2.17E-02 |
| Nrp1 | -0.55 | 2.64E-08 | 8.52E-04 |
| Dennd1a | -0.57 | 2.40E-08 | 7.76E-04 |
| Tbc1d5 | -0.57 | 3.36E-07 | 1.09E-02 |
| Dock2 | -0.58 | 4.02E-07 | 1.30E-02 |
| Asph | -0.59 | 4.76E-08 | 1.54E-03 |
| Pdia4 | -0.6 | 7.55E-08 | 2.44E-03 |
| Ophn1 | -0.62 | 1.41E-07 | 4.54E-03 |
| Itga6 | -0.62 | 2.15E-07 | 6.94E-03 |
| Lrmda | -0.66 | 1.01E-06 | 3.27E-02 |
| Pla2g4a | -0.68 | 4.62E-08 | 1.49E-03 |

| | | | |
|---|---|---|---|
| Tbxas1 | -0.7 | 1.88E-08 | 6.07E-04 |
| Bin1 | -0.74 | 9.59E-10 | 3.10E-05 |
| Immp2l | -0.87 | 1.73E-10 | 5.59E-06 |
| Nav3 | -0.9 | 8.49E-09 | 2.74E-04 |
| Spp1 | -0.97 | 2.69E-07 | 8.68E-03 |
| Hspa1b | -1.08 | 2.31E-12 | 7.46E-08 |
| Fn1 | -1.24 | 1.01E-11 | 3.26E-07 |
