## Supplemental Table 9 for "Vagal Signaling Decline in Age-related Macular Degeneration Drives Spleen-Dependent Retinal Inflammation"

| Gene | Log Fold Change | p-value | padj |
|---|---|---|---|
| Cxcl2 | 0.75 | 1.32E-18 | 4.27E-14 |
| Id2 | 0.74 | 1.84E-49 | 5.94E-45 |
| Cxcl3 | 0.74 | 4.91E-24 | 1.58E-19 |
| Ccl2 | 0.62 | 1.44E-17 | 4.64E-13 |
| Thbs1 | 0.6 | 2.62E-19 | 8.45E-15 |
| Nfkbia | 0.58 | 3.60E-37 | 1.16E-32 |
| Clec4e | 0.55 | 1.78E-35 | 5.75E-31 |
| Slc7a11 | 0.55 | 2.16E-25 | 6.99E-21 |
| Clec4d | 0.54 | 4.29E-33 | 1.39E-28 |
| Srgn | 0.53 | 4.66E-49 | 1.50E-44 |
| Vcan | 0.53 | 1.94E-21 | 6.28E-17 |
| Uba52 | 0.52 | 9.58E-63 | 3.09E-58 |
| Plaur | 0.51 | 3.58E-36 | 1.15E-31 |
| Mt1 | 0.51 | 4.01E-20 | 1.30E-15 |
| Eno1 | 0.49 | 6.90E-44 | 2.23E-39 |
| Pim1 | 0.47 | 3.22E-33 | 1.04E-28 |
| Rps29 | 0.46 | 8.53E-51 | 2.75E-46 |
| Furin | 0.45 | 2.53E-50 | 8.17E-46 |
| Ninj1 | 0.45 | 1.25E-29 | 4.02E-25 |
| Sdc4 | 0.45 | 4.54E-27 | 1.47E-22 |
| Ccl7 | 0.45 | 4.07E-15 | 1.31E-10 |
| Rpl38 | 0.44 | 8.30E-46 | 2.68E-41 |
| Plin2 | 0.44 | 4.35E-29 | 1.40E-24 |
| Cdkn1a | 0.43 | 5.30E-27 | 1.71E-22 |
| Arrdc4 | 0.42 | 5.91E-48 | 1.91E-43 |
| Rps27 | 0.42 | 1.39E-38 | 4.50E-34 |
| Wnk1 | 0.42 | 1.77E-31 | 5.71E-27 |
| Pkm | 0.41 | 7.43E-41 | 2.40E-36 |
| Fkbp5 | 0.41 | 6.87E-29 | 2.22E-24 |
| Chmp4b | 0.4 | 1.09E-51 | 3.51E-47 |
| Mkrn1 | 0.4 | 4.31E-34 | 1.39E-29 |
| Hif1a | 0.4 | 3.21E-28 | 1.04E-23 |
| Lgals3 | 0.39 | 1.37E-27 | 4.42E-23 |
| Cd80 | 0.39 | 7.67E-25 | 2.48E-20 |
| Noct | 0.38 | 3.51E-43 | 1.13E-38 |
| Emilin2 | 0.38 | 6.11E-38 | 1.97E-33 |
| Jak2 | 0.38 | 2.89E-32 | 9.34E-28 |
| Rnh1 | 0.38 | 3.95E-30 | 1.27E-25 |
| Adam8 | 0.38 | 8.34E-27 | 2.69E-22 |
| Hilpda | 0.38 | 2.77E-20 | 8.95E-16 |
| Il1r2 | 0.38 | 3.05E-19 | 9.85E-15 |
| Ier3 | 0.38 | 7.37E-12 | 2.38E-07 |
| Rbms1 | 0.37 | 4.13E-38 | 1.33E-33 |
| Rbbp6 | 0.37 | 6.30E-34 | 2.03E-29 |
| Tes | 0.37 | 3.02E-27 | 9.74E-23 |

| | | | |
|---|---|---|---|
| Btg1 | 0.37 | 5.90E-27 | 1.90E-22 |
| Crtc3 | 0.37 | 9.61E-27 | 3.10E-22 |
| Cd44 | 0.37 | 1.82E-20 | 5.88E-16 |
| Chst11 | 0.37 | 3.18E-17 | 1.03E-12 |
| Plcb1 | 0.37 | 2.82E-11 | 9.11E-07 |
| Eif3j1 | 0.36 | 2.04E-37 | 6.58E-33 |
| Rara | 0.36 | 6.21E-32 | 2.01E-27 |
| Nfil3 | 0.36 | 3.54E-30 | 1.14E-25 |
| Rpl35 | 0.36 | 8.33E-29 | 2.69E-24 |
| Lilrb4a | 0.36 | 4.12E-28 | 1.33E-23 |
| Card19 | 0.36 | 1.03E-27 | 3.31E-23 |
| Klf9 | 0.36 | 1.12E-27 | 3.61E-23 |
| Lilr4b | 0.36 | 6.00E-26 | 1.94E-21 |
| Tgfb1 | 0.36 | 1.12E-25 | 3.62E-21 |
| Klf6 | 0.36 | 2.46E-23 | 7.94E-19 |
| Rgcc | 0.36 | 1.14E-17 | 3.70E-13 |
| Gm11808 | 0.35 | 4.07E-51 | 1.31E-46 |
| Riok3 | 0.35 | 4.19E-36 | 1.35E-31 |
| Atp5md | 0.35 | 1.62E-33 | 5.23E-29 |
| Fam129b | 0.35 | 8.63E-31 | 2.78E-26 |
| Ap2a2 | 0.35 | 1.77E-26 | 5.72E-22 |
| Abtb2 | 0.35 | 1.59E-23 | 5.13E-19 |
| Uck2 | 0.35 | 6.51E-23 | 2.10E-18 |
| Ehd1 | 0.35 | 1.59E-19 | 5.13E-15 |
| Gsr | 0.35 | 2.46E-14 | 7.96E-10 |
| Cd14 | 0.35 | 9.39E-14 | 3.03E-09 |
| Emp1 | 0.35 | 1.11E-13 | 3.60E-09 |
| Picalm | 0.34 | 3.79E-26 | 1.22E-21 |
| Ube2d3 | 0.33 | 5.39E-40 | 1.74E-35 |
| Cd53 | 0.33 | 5.83E-38 | 1.88E-33 |
| Tet2 | 0.33 | 3.76E-26 | 1.21E-21 |
| Rps28 | 0.33 | 1.83E-24 | 5.90E-20 |
| Kdm6b | 0.33 | 9.22E-22 | 2.98E-17 |
| Klhl6 | 0.33 | 1.80E-21 | 5.80E-17 |
| Rnf149 | 0.33 | 2.12E-21 | 6.85E-17 |
| Prdx6 | 0.33 | 2.27E-19 | 7.31E-15 |
| Eps8 | 0.33 | 2.78E-16 | 8.96E-12 |
| Dusp1 | 0.33 | 4.32E-10 | 1.39E-05 |
| Mir22hg | 0.32 | 3.80E-37 | 1.23E-32 |
| Hnrnpa3 | 0.32 | 1.41E-29 | 4.56E-25 |
| Il4ra | 0.32 | 3.75E-24 | 1.21E-19 |
| Por | 0.32 | 5.84E-24 | 1.88E-19 |
| Cmip | 0.32 | 7.25E-24 | 2.34E-19 |
| Ets2 | 0.32 | 1.15E-22 | 3.70E-18 |
| Jmjd1c | 0.32 | 1.11E-21 | 3.60E-17 |
| Ltb4r1 | 0.32 | 2.12E-21 | 6.83E-17 |

| | | | |
|---|---|---|---|
| Fndc3a | 0.32 | 4.45E-21 | 1.44E-16 |
| Mindy3 | 0.32 | 4.49E-21 | 1.45E-16 |
| Ccr1 | 0.32 | 3.18E-18 | 1.03E-13 |
| Gm17268 | 0.32 | 4.86E-17 | 1.57E-12 |
| Plek | 0.32 | 2.92E-16 | 9.43E-12 |
| Morrbid | 0.32 | 5.74E-13 | 1.85E-08 |
| Fmnl2 | 0.32 | 7.47E-13 | 2.41E-08 |
| Osm | 0.32 | 3.35E-11 | 1.08E-06 |
| Hmox1 | 0.32 | 1.11E-10 | 3.60E-06 |
| Fxyd5 | 0.31 | 1.02E-27 | 3.29E-23 |
| Vav3 | 0.31 | 1.11E-16 | 3.58E-12 |
| Gab1 | 0.31 | 1.09E-15 | 3.51E-11 |
| Ell2 | 0.31 | 1.42E-14 | 4.59E-10 |
| Dennd4a | 0.31 | 7.28E-12 | 2.35E-07 |
| Vim | 0.31 | 1.21E-11 | 3.90E-07 |
| F13a1 | 0.31 | 8.47E-08 | 2.73E-03 |
| Gm19951 | 0.3 | 5.40E-30 | 1.74E-25 |
| Psma6 | 0.3 | 5.66E-26 | 1.83E-21 |
| Fosl2 | 0.3 | 2.17E-21 | 7.02E-17 |
| Anxa2 | 0.3 | 1.55E-16 | 5.01E-12 |
| Aff1 | 0.3 | 2.45E-16 | 7.92E-12 |
| Vapa | 0.29 | 1.32E-26 | 4.27E-22 |
| Ddit4 | 0.29 | 1.15E-25 | 3.73E-21 |
| Cdk8 | 0.29 | 1.16E-25 | 3.75E-21 |
| Atp5k | 0.29 | 1.18E-25 | 3.82E-21 |
| Map3k20 | 0.29 | 1.88E-25 | 6.06E-21 |
| Zfp91 | 0.29 | 7.98E-23 | 2.58E-18 |
| Cndp2 | 0.29 | 6.67E-21 | 2.15E-16 |
| Tuba1c | 0.29 | 5.29E-20 | 1.71E-15 |
| Jdp2 | 0.29 | 5.90E-20 | 1.90E-15 |
| Syk | 0.29 | 2.48E-17 | 8.02E-13 |
| Slc15a3 | 0.29 | 5.39E-17 | 1.74E-12 |
| S100a11 | 0.29 | 7.16E-17 | 2.31E-12 |
| Tnfaip3 | 0.29 | 1.24E-16 | 4.01E-12 |
| Bcl2a1d | 0.29 | 5.96E-16 | 1.92E-11 |
| Fbxo11 | 0.29 | 3.89E-15 | 1.26E-10 |
| Gapdh | 0.29 | 7.27E-15 | 2.35E-10 |
| Ahnak | 0.29 | 4.15E-14 | 1.34E-09 |
| Rbpj | 0.29 | 1.01E-13 | 3.26E-09 |
| Sgms1 | 0.29 | 2.05E-13 | 6.63E-09 |
| Msr1 | 0.29 | 4.76E-13 | 1.54E-08 |
| Pde4b | 0.29 | 3.51E-09 | 1.13E-04 |
| Rgs1 | 0.29 | 2.65E-07 | 8.56E-03 |
| Zfp52 | 0.28 | 6.77E-30 | 2.18E-25 |
| Erh | 0.28 | 2.16E-25 | 6.99E-21 |
| Ndufb1-ps | 0.28 | 5.36E-23 | 1.73E-18 |

| | | | |
|---|---|---|---|
| Diaph1 | 0.28 | 1.12E-22 | 3.60E-18 |
| Eif5 | 0.28 | 4.40E-22 | 1.42E-17 |
| H2afj | 0.28 | 1.10E-21 | 3.56E-17 |
| Mrpl52 | 0.28 | 1.85E-21 | 5.97E-17 |
| Errfi1 | 0.28 | 8.84E-21 | 2.85E-16 |
| Tmem189 | 0.28 | 1.64E-19 | 5.29E-15 |
| Arih1 | 0.28 | 5.63E-19 | 1.82E-14 |
| Gm26532 | 0.28 | 5.31E-18 | 1.72E-13 |
| Cebpb | 0.28 | 3.32E-17 | 1.07E-12 |
| Cyth1 | 0.28 | 3.91E-16 | 1.26E-11 |
| Inpp5d | 0.28 | 1.40E-15 | 4.51E-11 |
| Phlda1 | 0.28 | 3.25E-14 | 1.05E-09 |
| Mmp19 | 0.28 | 2.39E-13 | 7.73E-09 |
| Esd | 0.28 | 6.47E-13 | 2.09E-08 |
| Alcam | 0.28 | 4.15E-09 | 1.34E-04 |
| Ccl9 | 0.28 | 9.94E-08 | 3.21E-03 |
| Ralgps2 | 0.27 | 4.52E-30 | 1.46E-25 |
| Ppp2ca | 0.27 | 1.38E-23 | 4.46E-19 |
| Tpm3 | 0.27 | 5.88E-23 | 1.90E-18 |
| Map3k8 | 0.27 | 1.11E-21 | 3.59E-17 |
| Rab7 | 0.27 | 2.33E-21 | 7.52E-17 |
| Mdm2 | 0.27 | 1.57E-18 | 5.08E-14 |
| Slc23a2 | 0.27 | 5.37E-18 | 1.73E-13 |
| Ahr | 0.27 | 2.22E-16 | 7.17E-12 |
| Sec61g | 0.27 | 6.87E-16 | 2.22E-11 |
| Ptafr | 0.27 | 9.48E-15 | 3.06E-10 |
| Lmna | 0.27 | 5.69E-10 | 1.84E-05 |
| Foxn3 | 0.27 | 1.13E-09 | 3.64E-05 |
| Cacna1d | 0.27 | 1.22E-07 | 3.94E-03 |
| Son | 0.26 | 1.59E-26 | 5.12E-22 |
| Cers2 | 0.26 | 1.45E-23 | 4.69E-19 |
| Dhx15 | 0.26 | 9.45E-23 | 3.05E-18 |
| Cul3 | 0.26 | 6.10E-20 | 1.97E-15 |
| Hmgb1 | 0.26 | 7.03E-20 | 2.27E-15 |
| Rpl41 | 0.26 | 2.67E-18 | 8.64E-14 |
| Ucp2 | 0.26 | 7.75E-18 | 2.50E-13 |
| Gch1 | 0.26 | 1.01E-17 | 3.27E-13 |
| Pfkp | 0.26 | 2.87E-17 | 9.26E-13 |
| Braf | 0.26 | 1.27E-16 | 4.10E-12 |
| Zyx | 0.26 | 2.05E-16 | 6.60E-12 |
| Rpl36 | 0.26 | 3.38E-16 | 1.09E-11 |
| Slc11a1 | 0.26 | 2.11E-15 | 6.80E-11 |
| Tlr4 | 0.26 | 1.05E-13 | 3.39E-09 |
| Txnrd1 | 0.26 | 5.23E-12 | 1.69E-07 |
| Havcr2 | 0.26 | 1.76E-11 | 5.68E-07 |
| Gab2 | 0.26 | 8.01E-11 | 2.59E-06 |

| | | | |
|---|---|---|---|
| Cytip | 0.26 | 8.09E-09 | 2.61E-04 |
| Gm2000 | 0.25 | 9.80E-27 | 3.16E-22 |
| Elob | 0.25 | 1.50E-18 | 4.83E-14 |
| Stk24 | 0.25 | 7.69E-18 | 2.48E-13 |
| Fkbp1a | 0.25 | 3.52E-17 | 1.14E-12 |
| Sqstm1 | 0.25 | 1.74E-14 | 5.61E-10 |
| Cd83 | 0.25 | 1.23E-13 | 3.98E-09 |
| Ezr | 0.25 | 3.08E-13 | 9.96E-09 |
| Ftl1 | 0.25 | 2.35E-12 | 7.60E-08 |
| Smox | 0.25 | 2.49E-11 | 8.05E-07 |
| Fbxl5 | 0.25 | 5.43E-11 | 1.75E-06 |
| Polr2l | 0.24 | 2.12E-25 | 6.85E-21 |
| Pofut2 | 0.24 | 1.74E-23 | 5.61E-19 |
| Ube2d2a | 0.24 | 5.07E-22 | 1.64E-17 |
| Rbm39 | 0.24 | 1.57E-21 | 5.08E-17 |
| Pitpna | 0.24 | 1.64E-21 | 5.28E-17 |
| Ppp2cb | 0.24 | 1.24E-20 | 4.01E-16 |
| Sap18 | 0.24 | 3.69E-20 | 1.19E-15 |
| Stk40 | 0.24 | 9.95E-20 | 3.21E-15 |
| Brd4 | 0.24 | 1.93E-19 | 6.23E-15 |
| Arhgdia | 0.24 | 2.29E-19 | 7.40E-15 |
| Met | 0.24 | 6.69E-19 | 2.16E-14 |
| U2af1 | 0.24 | 1.24E-18 | 3.99E-14 |
| Ndufa3 | 0.24 | 1.77E-18 | 5.71E-14 |
| Atp6v0d1 | 0.24 | 5.38E-18 | 1.74E-13 |
| Fmnl1 | 0.24 | 2.37E-17 | 7.65E-13 |
| Eif2s2 | 0.24 | 4.13E-17 | 1.33E-12 |
| St3gal5 | 0.24 | 4.93E-17 | 1.59E-12 |
| Ric1 | 0.24 | 8.64E-17 | 2.79E-12 |
| Hnrnpdl | 0.24 | 8.94E-16 | 2.88E-11 |
| Smim3 | 0.24 | 4.17E-15 | 1.35E-10 |
| Xbp1 | 0.24 | 4.75E-15 | 1.53E-10 |
| Atp6v0c | 0.24 | 2.30E-14 | 7.42E-10 |
| Slfn2 | 0.24 | 1.81E-13 | 5.84E-09 |
| Jarid2 | 0.24 | 2.41E-13 | 7.77E-09 |
| Tcf7l2 | 0.24 | 4.67E-11 | 1.51E-06 |
| Pdpn | 0.24 | 8.46E-11 | 2.73E-06 |
| S100a4 | 0.24 | 9.20E-11 | 2.97E-06 |
| Gda | 0.24 | 2.67E-10 | 8.61E-06 |
| Trf | 0.24 | 1.46E-09 | 4.71E-05 |
| Osbpl8 | 0.24 | 1.66E-09 | 5.35E-05 |
| F10 | 0.24 | 3.06E-07 | 9.88E-03 |
| Tatdn2 | 0.23 | 4.82E-25 | 1.56E-20 |
| Cirbp | 0.23 | 1.51E-23 | 4.88E-19 |
| Tmem248 | 0.23 | 2.04E-22 | 6.60E-18 |
| Psmb7 | 0.23 | 9.31E-20 | 3.00E-15 |

| | | | |
|---|---|---|---|
| Atf4 | 0.23 | 1.32E-18 | 4.27E-14 |
| Ccdc71l | 0.23 | 1.47E-18 | 4.76E-14 |
| Gm31718 | 0.23 | 1.60E-18 | 5.16E-14 |
| Ptbp2 | 0.23 | 3.01E-18 | 9.72E-14 |
| Eif1 | 0.23 | 3.64E-18 | 1.18E-13 |
| Ufm1 | 0.23 | 3.67E-18 | 1.18E-13 |
| Atp1a1 | 0.23 | 5.21E-18 | 1.68E-13 |
| Fus | 0.23 | 6.81E-17 | 2.20E-12 |
| Ube2k | 0.23 | 1.61E-16 | 5.19E-12 |
| Map2k4 | 0.23 | 2.26E-16 | 7.29E-12 |
| Atp1b3 | 0.23 | 2.32E-15 | 7.50E-11 |
| Lrrc8c | 0.23 | 2.67E-15 | 8.63E-11 |
| Arih2 | 0.23 | 1.77E-14 | 5.72E-10 |
| Cflar | 0.23 | 2.43E-14 | 7.84E-10 |
| Cycs | 0.23 | 4.14E-14 | 1.34E-09 |
| Ndel1 | 0.23 | 4.28E-14 | 1.38E-09 |
| Cux1 | 0.23 | 7.61E-14 | 2.46E-09 |
| Ubb | 0.23 | 1.85E-13 | 5.97E-09 |
| Rab8b | 0.23 | 4.35E-13 | 1.40E-08 |
| Dennd4c | 0.23 | 5.31E-13 | 1.71E-08 |
| Plec | 0.23 | 1.60E-12 | 5.16E-08 |
| Cblb | 0.23 | 1.77E-12 | 5.70E-08 |
| Eea1 | 0.23 | 7.38E-12 | 2.38E-07 |
| Myo5a | 0.23 | 9.39E-12 | 3.03E-07 |
| Ptgs2 | 0.23 | 2.74E-09 | 8.84E-05 |
| Junb | 0.23 | 2.57E-08 | 8.30E-04 |
| Malt1 | 0.23 | 1.32E-07 | 4.25E-03 |
| B930036N10I | 0.22 | 8.89E-25 | 2.87E-20 |
| Cdk2ap1 | 0.22 | 1.90E-21 | 6.15E-17 |
| Pgs1 | 0.22 | 2.53E-18 | 8.17E-14 |
| Otulin | 0.22 | 1.47E-17 | 4.76E-13 |
| Tle3 | 0.22 | 4.17E-17 | 1.34E-12 |
| Gm6377 | 0.22 | 5.13E-17 | 1.66E-12 |
| Scamp1 | 0.22 | 5.58E-17 | 1.80E-12 |
| Rbm26 | 0.22 | 2.12E-16 | 6.84E-12 |
| Mcl1 | 0.22 | 6.36E-16 | 2.05E-11 |
| Rffl | 0.22 | 1.06E-15 | 3.43E-11 |
| Ssr3 | 0.22 | 1.75E-15 | 5.66E-11 |
| Bzw1 | 0.22 | 4.96E-15 | 1.60E-10 |
| Srsf5 | 0.22 | 5.35E-15 | 1.73E-10 |
| Psmd14 | 0.22 | 6.05E-15 | 1.95E-10 |
| Luc7l2 | 0.22 | 9.05E-15 | 2.92E-10 |
| Abcc1 | 0.22 | 4.82E-14 | 1.56E-09 |
| Tomm20 | 0.22 | 9.87E-14 | 3.19E-09 |
| Atp11b | 0.22 | 1.89E-13 | 6.10E-09 |
| Fem1c | 0.22 | 3.12E-13 | 1.01E-08 |

| | | | |
|---|---|---|---|
| Klhl2 | 0.22 | 8.23E-13 | 2.66E-08 |
| Zfand3 | 0.22 | 1.90E-12 | 6.13E-08 |
| Eif5a | 0.22 | 2.73E-12 | 8.81E-08 |
| Hspa8 | 0.22 | 1.71E-11 | 5.53E-07 |
| Ptpn12 | 0.22 | 2.06E-11 | 6.65E-07 |
| Cdk6 | 0.22 | 2.53E-11 | 8.18E-07 |
| Mdfic | 0.22 | 6.00E-10 | 1.94E-05 |
| Neat1 | 0.22 | 1.57E-09 | 5.06E-05 |
| S100a10 | 0.22 | 9.04E-08 | 2.92E-03 |
| Socs3 | 0.22 | 6.60E-07 | 2.13E-02 |
| Rnaset2a | 0.21 | 1.14E-19 | 3.68E-15 |
| Eif4h | 0.21 | 8.09E-17 | 2.61E-12 |
| Vamp8 | 0.21 | 1.63E-16 | 5.27E-12 |
| Psmc1 | 0.21 | 2.83E-16 | 9.12E-12 |
| Pet100 | 0.21 | 3.00E-16 | 9.68E-12 |
| Ube2l3 | 0.21 | 5.20E-16 | 1.68E-11 |
| Emc7 | 0.21 | 8.43E-16 | 2.72E-11 |
| Gnb1 | 0.21 | 9.34E-16 | 3.02E-11 |
| Pabpn1 | 0.21 | 1.23E-15 | 3.96E-11 |
| Ppp1r2 | 0.21 | 4.13E-15 | 1.33E-10 |
| Epc2 | 0.21 | 6.61E-15 | 2.14E-10 |
| Med13 | 0.21 | 7.18E-15 | 2.32E-10 |
| Nisch | 0.21 | 9.14E-15 | 2.95E-10 |
| Map2k3 | 0.21 | 1.07E-14 | 3.45E-10 |
| Crebbp | 0.21 | 1.12E-14 | 3.61E-10 |
| Arpp19 | 0.21 | 1.38E-14 | 4.47E-10 |
| Sh2b2 | 0.21 | 1.74E-14 | 5.62E-10 |
| Foxn2 | 0.21 | 3.08E-14 | 9.95E-10 |
| Etf1 | 0.21 | 9.51E-14 | 3.07E-09 |
| Atp6v1e1 | 0.21 | 9.59E-14 | 3.10E-09 |
| Srsf3 | 0.21 | 1.06E-13 | 3.42E-09 |
| Abhd17b | 0.21 | 2.08E-13 | 6.71E-09 |
| Ankrd11 | 0.21 | 3.37E-13 | 1.09E-08 |
| Mkln1 | 0.21 | 3.40E-13 | 1.10E-08 |
| Hnrnpll | 0.21 | 5.16E-13 | 1.67E-08 |
| Tfec | 0.21 | 7.40E-13 | 2.39E-08 |
| H3f3b | 0.21 | 7.93E-13 | 2.56E-08 |
| Lrrfip2 | 0.21 | 1.03E-12 | 3.33E-08 |
| Flna | 0.21 | 1.70E-12 | 5.48E-08 |
| Gm13986 | 0.21 | 1.97E-12 | 6.34E-08 |
| Prkce | 0.21 | 2.45E-12 | 7.92E-08 |
| Slk | 0.21 | 6.36E-12 | 2.05E-07 |
| Slc38a2 | 0.21 | 6.96E-12 | 2.25E-07 |
| Glipr2 | 0.21 | 8.67E-12 | 2.80E-07 |
| Tiam1 | 0.21 | 1.16E-11 | 3.76E-07 |
| Camk1d | 0.21 | 3.04E-11 | 9.81E-07 |

| | | | |
|---|---|---|---|
| Trem1 | 0.21 | 3.98E-11 | 1.28E-06 |
| Lrrfip1 | 0.21 | 5.35E-11 | 1.73E-06 |
| Lyn | 0.21 | 5.71E-11 | 1.84E-06 |
| Baz2b | 0.21 | 9.19E-11 | 2.97E-06 |
| Fam107b | 0.21 | 2.03E-10 | 6.56E-06 |
| Kdm7a | 0.21 | 3.03E-10 | 9.79E-06 |
| Etv6 | 0.21 | 1.29E-09 | 4.17E-05 |
| Chd7 | 0.21 | 5.38E-09 | 1.74E-04 |
| Gas7 | 0.21 | 1.35E-07 | 4.37E-03 |
| Tnfaip8l2 | -0.21 | 1.48E-14 | 4.77E-10 |
| Shtn1 | -0.21 | 1.38E-13 | 4.46E-09 |
| St6gal1 | -0.21 | 2.59E-12 | 8.38E-08 |
| Rpl23a | -0.21 | 1.42E-10 | 4.59E-06 |
| Nrp1 | -0.21 | 8.58E-10 | 2.77E-05 |
| Cxcl16 | -0.21 | 1.19E-09 | 3.83E-05 |
| Ms4a6c | -0.21 | 9.74E-08 | 3.15E-03 |
| Aif1 | -0.21 | 3.55E-07 | 1.15E-02 |
| Spint1 | -0.21 | 3.91E-07 | 1.26E-02 |
| Parp1 | -0.22 | 1.50E-19 | 4.84E-15 |
| Gm34084 | -0.22 | 1.13E-15 | 3.66E-11 |
| Abi3 | -0.22 | 1.14E-15 | 3.67E-11 |
| Grcc10 | -0.22 | 1.62E-15 | 5.24E-11 |
| Fcgr4 | -0.22 | 2.51E-15 | 8.11E-11 |
| Pik3cd | -0.22 | 8.98E-15 | 2.90E-10 |
| Unc93b1 | -0.22 | 7.66E-14 | 2.47E-09 |
| Vps13c | -0.22 | 2.16E-13 | 6.98E-09 |
| Ptprc | -0.22 | 1.09E-10 | 3.51E-06 |
| Selenop | -0.22 | 1.11E-07 | 3.59E-03 |
| Tmcc3 | -0.23 | 1.82E-18 | 5.88E-14 |
| Fam174a | -0.23 | 4.06E-18 | 1.31E-13 |
| Ypel3 | -0.23 | 2.76E-16 | 8.93E-12 |
| Epsti1 | -0.23 | 5.20E-11 | 1.68E-06 |
| Gm5150 | -0.23 | 2.44E-10 | 7.87E-06 |
| Tbc1d5 | -0.23 | 6.51E-10 | 2.10E-05 |
| Pde7b | -0.23 | 7.21E-10 | 2.33E-05 |
| Gpr141 | -0.23 | 2.38E-08 | 7.70E-04 |
| Ms4a4c | -0.23 | 8.95E-08 | 2.89E-03 |
| Arl6ip1 | -0.24 | 1.19E-16 | 3.85E-12 |
| Timm10b | -0.24 | 1.49E-15 | 4.83E-11 |
| Hacd4 | -0.24 | 7.82E-15 | 2.53E-10 |
| Ehd4 | -0.24 | 2.68E-13 | 8.64E-09 |
| Mndal | -0.24 | 1.11E-12 | 3.59E-08 |
| Clec4a3 | -0.24 | 9.12E-11 | 2.94E-06 |
| Itgb5 | -0.24 | 2.97E-10 | 9.58E-06 |
| Arhgap22 | -0.25 | 3.26E-33 | 1.05E-28 |
| Itga6 | -0.25 | 1.65E-23 | 5.32E-19 |

| | | | |
|---|---|---|---|
| Aoah | -0.25 | 1.92E-14 | 6.21E-10 |
| Cd38 | -0.25 | 2.01E-13 | 6.50E-09 |
| Pbx1 | -0.25 | 2.17E-13 | 7.01E-09 |
| Trem2 | -0.25 | 1.49E-11 | 4.82E-07 |
| Myo1e | -0.25 | 1.75E-09 | 5.66E-05 |
| Slc8a1 | -0.25 | 9.42E-07 | 3.04E-02 |
| H2-K1 | -0.26 | 1.03E-21 | 3.33E-17 |
| Otulinl | -0.26 | 3.75E-15 | 1.21E-10 |
| Pnp | -0.26 | 4.72E-15 | 1.52E-10 |
| Cx3cr1 | -0.26 | 2.27E-13 | 7.33E-09 |
| Psmb9 | -0.27 | 1.47E-25 | 4.75E-21 |
| Lamtor2 | -0.27 | 6.50E-22 | 2.10E-17 |
| Cd48 | -0.27 | 2.60E-18 | 8.39E-14 |
| Itm2b | -0.27 | 1.65E-17 | 5.34E-13 |
| Ifi30 | -0.27 | 2.27E-14 | 7.31E-10 |
| Ncf1 | -0.27 | 3.58E-14 | 1.16E-09 |
| Prkcb | -0.27 | 4.55E-14 | 1.47E-09 |
| Gngt2 | -0.27 | 1.08E-13 | 3.47E-09 |
| Arhgap24 | -0.27 | 5.24E-10 | 1.69E-05 |
| Rnase4 | -0.28 | 1.50E-22 | 4.83E-18 |
| Rsrp1 | -0.29 | 7.03E-24 | 2.27E-19 |
| Myo1f | -0.29 | 1.89E-17 | 6.11E-13 |
| Anxa3 | -0.29 | 2.47E-12 | 7.96E-08 |
| Arhgap15 | -0.29 | 4.04E-09 | 1.31E-04 |
| Prdx5 | -0.3 | 6.04E-19 | 1.95E-14 |
| Fgd4 | -0.3 | 9.30E-17 | 3.00E-12 |
| Dynll1 | -0.3 | 4.31E-15 | 1.39E-10 |
| F11r | -0.31 | 3.07E-31 | 9.92E-27 |
| Pou2f2 | -0.31 | 1.49E-20 | 4.82E-16 |
| Cd300c2 | -0.32 | 5.95E-21 | 1.92E-16 |
| Hexb | -0.32 | 9.67E-21 | 3.12E-16 |
| Mef2c | -0.32 | 7.95E-19 | 2.57E-14 |
| Lpcat2 | -0.33 | 2.43E-25 | 7.85E-21 |
| Frmd4b | -0.34 | 6.61E-22 | 2.13E-17 |
| Ly86 | -0.36 | 8.35E-21 | 2.70E-16 |
| Tmsb4x | -0.37 | 3.43E-27 | 1.11E-22 |
| Lair1 | -0.38 | 1.10E-31 | 3.56E-27 |
| C1qc | -0.38 | 6.99E-13 | 2.26E-08 |
| C1qb | -0.39 | 1.16E-09 | 3.76E-05 |
| Cybb | -0.4 | 9.54E-18 | 3.08E-13 |
| Ms4a6b | -0.41 | 5.56E-26 | 1.80E-21 |
| C1qa | -0.45 | 3.26E-15 | 1.05E-10 |
| Ctsc | -0.49 | 1.76E-33 | 5.69E-29 |
| Spp1 | -0.49 | 9.16E-11 | 2.96E-06 |
| Lrmda | -0.51 | 2.25E-27 | 7.26E-23 |
