## Supplemental Table 10 for "Vagal Signaling Decline in Age-related Macular Degeneration Drives Spleen-Dependent Retinal Inflammation"

| Gene | Log Fold Change | p-value | padj |
|---|---|---|---|
| Gsr | 0.82 | 2.33E-57 | 7.54E-53 |
| Cytip | 0.72 | 9.54E-46 | 3.08E-41 |
| Hp | 0.66 | 4.14E-31 | 1.34E-26 |
| Plac8 | 0.65 | 6.39E-20 | 2.06E-15 |
| Plaur | 0.6 | 1.37E-42 | 4.43E-38 |
| Il1r2 | 0.57 | 2.31E-38 | 7.46E-34 |
| Filip1l | 0.55 | 6.01E-51 | 1.94E-46 |
| Crtc3 | 0.51 | 1.24E-42 | 4.01E-38 |
| Stk17b | 0.51 | 1.15E-34 | 3.71E-30 |
| Klf13 | 0.5 | 3.17E-37 | 1.02E-32 |
| Lyn | 0.49 | 3.99E-39 | 1.29E-34 |
| Havcr2 | 0.49 | 1.03E-34 | 3.34E-30 |
| Napsa | 0.49 | 1.36E-32 | 4.38E-28 |
| Ptpn1 | 0.48 | 1.49E-37 | 4.82E-33 |
| Slk | 0.47 | 8.85E-44 | 2.86E-39 |
| Cox7a2l | 0.47 | 2.05E-38 | 6.63E-34 |
| Btg1 | 0.47 | 1.89E-33 | 6.10E-29 |
| Cmss1 | 0.46 | 2.96E-49 | 9.55E-45 |
| Klhl6 | 0.46 | 4.06E-35 | 1.31E-30 |
| Ly6c2 | 0.46 | 2.99E-21 | 9.64E-17 |
| Rps27 | 0.45 | 7.99E-30 | 2.58E-25 |
| Cdk8 | 0.44 | 2.08E-46 | 6.72E-42 |
| Rps29 | 0.44 | 1.99E-32 | 6.42E-28 |
| Ifitm6 | 0.44 | 2.95E-21 | 9.52E-17 |
| Fkbp5 | 0.43 | 2.94E-26 | 9.48E-22 |
| Inpp5d | 0.42 | 4.25E-30 | 1.37E-25 |
| Emb | 0.42 | 1.18E-26 | 3.81E-22 |
| Txnip | 0.42 | 3.41E-23 | 1.10E-18 |
| Il17ra | 0.41 | 3.41E-30 | 1.10E-25 |
| Runx3 | 0.41 | 6.62E-23 | 2.14E-18 |
| Plbd1 | 0.41 | 7.25E-23 | 2.34E-18 |
| Sorl1 | 0.4 | 4.75E-28 | 1.53E-23 |
| Mgst1 | 0.4 | 1.35E-25 | 4.36E-21 |
| Smox | 0.4 | 4.70E-23 | 1.52E-18 |
| Pla2g7 | 0.4 | 1.51E-16 | 4.89E-12 |
| Sla | 0.39 | 1.28E-26 | 4.15E-22 |
| Syk | 0.39 | 2.83E-26 | 9.12E-22 |
| Foxn3 | 0.39 | 3.54E-15 | 1.14E-10 |
| Itch | 0.38 | 7.62E-26 | 2.46E-21 |
| Rps28 | 0.38 | 9.28E-24 | 3.00E-19 |
| Rpl23a | 0.38 | 3.87E-23 | 1.25E-18 |
| Ifngr1 | 0.38 | 3.32E-22 | 1.07E-17 |
| Csf2rb | 0.38 | 5.47E-21 | 1.77E-16 |
| Pim1 | 0.38 | 1.74E-19 | 5.61E-15 |
| Ccnd3 | 0.38 | 2.00E-17 | 6.46E-13 |

| | | | |
|---|---|---|---|
| Ifitm2 | 0.38 | 1.56E-16 | 5.03E-12 |
| Srrm1 | 0.37 | 3.21E-32 | 1.04E-27 |
| Rbbp6 | 0.37 | 3.68E-28 | 1.19E-23 |
| Cd244a | 0.37 | 7.24E-26 | 2.34E-21 |
| Mkrn1 | 0.37 | 4.17E-25 | 1.34E-20 |
| Baz2b | 0.36 | 3.10E-24 | 1.00E-19 |
| Jak2 | 0.36 | 6.33E-23 | 2.04E-18 |
| Gcnt2 | 0.36 | 2.08E-20 | 6.71E-16 |
| Rpl10 | 0.36 | 4.20E-19 | 1.35E-14 |
| March1 | 0.36 | 3.95E-16 | 1.27E-11 |
| Nop53 | 0.35 | 4.20E-33 | 1.36E-28 |
| Bach1 | 0.35 | 1.89E-24 | 6.09E-20 |
| Tet2 | 0.35 | 1.50E-23 | 4.85E-19 |
| Rnf169 | 0.35 | 3.66E-23 | 1.18E-18 |
| Fau | 0.35 | 1.35E-22 | 4.36E-18 |
| Rpl18a | 0.35 | 3.69E-19 | 1.19E-14 |
| Ssh2 | 0.35 | 3.78E-15 | 1.22E-10 |
| Son | 0.34 | 7.69E-36 | 2.48E-31 |
| Tspan13 | 0.34 | 4.60E-29 | 1.49E-24 |
| Ncf2 | 0.34 | 2.39E-27 | 7.72E-23 |
| Emilin2 | 0.34 | 5.39E-27 | 1.74E-22 |
| Mcemp1 | 0.34 | 2.22E-20 | 7.17E-16 |
| Klf9 | 0.34 | 1.36E-19 | 4.40E-15 |
| Rpl39 | 0.34 | 1.65E-16 | 5.31E-12 |
| Lsp1 | 0.34 | 1.31E-13 | 4.22E-09 |
| Cd52 | 0.34 | 6.86E-12 | 2.21E-07 |
| Id2 | 0.34 | 1.33E-09 | 4.30E-05 |
| Zfyve9 | 0.33 | 1.88E-24 | 6.08E-20 |
| Sell | 0.33 | 6.47E-24 | 2.09E-19 |
| Gm15283 | 0.33 | 7.84E-24 | 2.53E-19 |
| Arrdc4 | 0.33 | 1.06E-23 | 3.41E-19 |
| Rpl13a | 0.33 | 6.34E-21 | 2.05E-16 |
| St3gal4 | 0.33 | 4.58E-19 | 1.48E-14 |
| Runx1 | 0.33 | 1.30E-14 | 4.21E-10 |
| Zfp36l2 | 0.33 | 1.69E-14 | 5.44E-10 |
| Zfp52 | 0.32 | 5.77E-32 | 1.86E-27 |
| Fmnl1 | 0.32 | 4.13E-25 | 1.33E-20 |
| Spi1 | 0.32 | 6.82E-24 | 2.20E-19 |
| Diaph1 | 0.32 | 1.14E-23 | 3.69E-19 |
| H2-D1 | 0.32 | 3.56E-23 | 1.15E-18 |
| Aldh2 | 0.32 | 3.73E-22 | 1.20E-17 |
| 2410006H16I | 0.32 | 1.34E-19 | 4.34E-15 |
| Ms4a4a | 0.32 | 3.70E-19 | 1.20E-14 |
| Slfn2 | 0.32 | 2.98E-17 | 9.62E-13 |
| Rnf149 | 0.32 | 4.15E-17 | 1.34E-12 |
| Rpl37 | 0.32 | 3.54E-16 | 1.14E-11 |

| | | | |
|---|---|---|---|
| Rps23 | 0.32 | 7.47E-16 | 2.41E-11 |
| Cd80 | 0.32 | 2.79E-14 | 9.00E-10 |
| Gab2 | 0.32 | 7.42E-13 | 2.40E-08 |
| Rbm39 | 0.31 | 2.82E-26 | 9.10E-22 |
| Sbno2 | 0.31 | 2.54E-22 | 8.19E-18 |
| Man2b1 | 0.31 | 1.68E-21 | 5.43E-17 |
| Atp11b | 0.31 | 1.95E-21 | 6.30E-17 |
| Tacc1 | 0.31 | 3.13E-21 | 1.01E-16 |
| H2afj | 0.31 | 2.52E-20 | 8.13E-16 |
| Rara | 0.31 | 7.99E-20 | 2.58E-15 |
| Cebpb | 0.31 | 3.17E-18 | 1.02E-13 |
| Ap2a2 | 0.31 | 1.31E-17 | 4.22E-13 |
| Herc4 | 0.31 | 4.69E-17 | 1.51E-12 |
| Cd300a | 0.31 | 1.63E-16 | 5.27E-12 |
| Rps14 | 0.31 | 2.71E-16 | 8.74E-12 |
| Gpr141 | 0.31 | 2.35E-12 | 7.58E-08 |
| Gm46224 | 0.3 | 3.27E-25 | 1.05E-20 |
| Il10ra | 0.3 | 1.27E-21 | 4.11E-17 |
| Zfp91 | 0.3 | 3.42E-20 | 1.10E-15 |
| Jaml | 0.3 | 3.93E-19 | 1.27E-14 |
| Taldo1 | 0.3 | 9.75E-18 | 3.15E-13 |
| Cd47 | 0.3 | 2.46E-17 | 7.93E-13 |
| Ahr | 0.3 | 5.87E-17 | 1.89E-12 |
| Rpl28 | 0.3 | 2.75E-14 | 8.87E-10 |
| Eef2 | 0.29 | 8.07E-20 | 2.61E-15 |
| Abhd17b | 0.29 | 9.69E-20 | 3.13E-15 |
| Sik1 | 0.29 | 1.73E-19 | 5.59E-15 |
| Itgal | 0.29 | 2.95E-19 | 9.52E-15 |
| Fam49a | 0.29 | 5.06E-19 | 1.63E-14 |
| C3 | 0.29 | 1.86E-15 | 5.99E-11 |
| Rpl38 | 0.29 | 2.92E-15 | 9.44E-11 |
| Mindy3 | 0.29 | 5.61E-15 | 1.81E-10 |
| Rpl17 | 0.29 | 8.21E-14 | 2.65E-09 |
| Uck2 | 0.29 | 2.52E-13 | 8.14E-09 |
| Rpl34 | 0.29 | 9.00E-13 | 2.91E-08 |
| Lrp10 | 0.28 | 7.82E-26 | 2.53E-21 |
| Morf4l1 | 0.28 | 6.70E-24 | 2.16E-19 |
| Ptpn22 | 0.28 | 2.71E-22 | 8.76E-18 |
| Epc2 | 0.28 | 4.50E-20 | 1.45E-15 |
| Arglu1 | 0.28 | 6.73E-20 | 2.17E-15 |
| Stk24 | 0.28 | 7.51E-18 | 2.42E-13 |
| Rps15 | 0.28 | 2.58E-15 | 8.32E-11 |
| Rpl4 | 0.28 | 5.82E-15 | 1.88E-10 |
| Rpl31 | 0.28 | 7.60E-14 | 2.45E-09 |
| Rps21 | 0.28 | 1.19E-13 | 3.83E-09 |
| Tmem176b | 0.28 | 3.92E-13 | 1.27E-08 |

| | | | |
|---|---|---|---|
| Rpl35 | 0.28 | 2.00E-12 | 6.45E-08 |
| Rplp0 | 0.28 | 5.62E-12 | 1.81E-07 |
| AY036118 | 0.28 | 7.03E-09 | 2.27E-04 |
| Ralgps2 | 0.27 | 1.51E-24 | 4.89E-20 |
| Pnisr | 0.27 | 1.83E-19 | 5.90E-15 |
| Irf5 | 0.27 | 1.34E-18 | 4.32E-14 |
| Wapl | 0.27 | 3.22E-18 | 1.04E-13 |
| Fyn | 0.27 | 7.05E-17 | 2.28E-12 |
| Ddi2 | 0.27 | 7.36E-16 | 2.38E-11 |
| Rplp2 | 0.27 | 1.51E-15 | 4.88E-11 |
| Eif3f | 0.27 | 1.15E-14 | 3.70E-10 |
| Tgfb1 | 0.27 | 1.54E-13 | 4.99E-09 |
| Acer3 | 0.27 | 2.66E-13 | 8.59E-09 |
| Klra2 | 0.27 | 1.80E-12 | 5.82E-08 |
| Eef1b2 | 0.27 | 3.03E-12 | 9.79E-08 |
| Msrb1 | 0.27 | 3.96E-10 | 1.28E-05 |
| Esd | 0.27 | 1.13E-09 | 3.64E-05 |
| Samhd1 | 0.27 | 5.47E-09 | 1.77E-04 |
| Gm11808 | 0.26 | 2.28E-22 | 7.36E-18 |
| Pcif1 | 0.26 | 3.54E-22 | 1.14E-17 |
| Srrm2 | 0.26 | 3.57E-18 | 1.15E-13 |
| Ddx6 | 0.26 | 2.38E-17 | 7.69E-13 |
| Rock1 | 0.26 | 4.36E-17 | 1.41E-12 |
| Sys1 | 0.26 | 1.75E-16 | 5.64E-12 |
| Arhgap30 | 0.26 | 3.37E-16 | 1.09E-11 |
| Orai1 | 0.26 | 5.41E-16 | 1.75E-11 |
| Mia2 | 0.26 | 2.70E-15 | 8.72E-11 |
| Prkce | 0.26 | 1.62E-14 | 5.25E-10 |
| Cmip | 0.26 | 1.27E-13 | 4.09E-09 |
| Rps9 | 0.26 | 1.48E-13 | 4.76E-09 |
| Pfkp | 0.26 | 3.61E-13 | 1.16E-08 |
| Kdm7a | 0.26 | 6.16E-13 | 1.99E-08 |
| Rpl24 | 0.26 | 1.35E-12 | 4.37E-08 |
| Rpl27a | 0.26 | 5.05E-12 | 1.63E-07 |
| Ptprc | 0.26 | 1.48E-11 | 4.79E-07 |
| Rpl30 | 0.26 | 2.18E-11 | 7.04E-07 |
| Cirbp | 0.25 | 8.92E-24 | 2.88E-19 |
| Pdcd4 | 0.25 | 1.20E-21 | 3.89E-17 |
| Gps2 | 0.25 | 8.42E-21 | 2.72E-16 |
| Ppp2cb | 0.25 | 4.13E-19 | 1.33E-14 |
| Ccnl2 | 0.25 | 1.12E-18 | 3.60E-14 |
| Otulin | 0.25 | 8.11E-18 | 2.62E-13 |
| Hcls1 | 0.25 | 3.85E-17 | 1.24E-12 |
| Srsf11 | 0.25 | 4.34E-17 | 1.40E-12 |
| Rbm26 | 0.25 | 4.52E-17 | 1.46E-12 |
| Brd4 | 0.25 | 9.21E-17 | 2.97E-12 |

| | | | |
|---|---|---|---|
| Ikbkb | 0.25 | 1.95E-16 | 6.28E-12 |
| Il13ra1 | 0.25 | 2.85E-16 | 9.19E-12 |
| Rin3 | 0.25 | 4.55E-16 | 1.47E-11 |
| Tmem234 | 0.25 | 5.24E-16 | 1.69E-11 |
| Arid4b | 0.25 | 7.66E-16 | 2.47E-11 |
| Arid4a | 0.25 | 9.51E-16 | 3.07E-11 |
| Map2k4 | 0.25 | 1.16E-15 | 3.75E-11 |
| Usp3 | 0.25 | 1.56E-15 | 5.03E-11 |
| Rbm25 | 0.25 | 2.81E-15 | 9.06E-11 |
| Hnrnpdl | 0.25 | 1.22E-14 | 3.94E-10 |
| Nsf | 0.25 | 1.09E-13 | 3.52E-09 |
| Vsir | 0.25 | 1.71E-13 | 5.52E-09 |
| Lrrfip1 | 0.25 | 4.69E-12 | 1.52E-07 |
| Rps6 | 0.25 | 1.74E-11 | 5.61E-07 |
| Rps25 | 0.25 | 2.04E-11 | 6.58E-07 |
| Cdk2ap2 | 0.25 | 6.66E-11 | 2.15E-06 |
| Rpl26 | 0.25 | 1.09E-10 | 3.52E-06 |
| Gm9733 | 0.25 | 1.20E-10 | 3.88E-06 |
| Rps24 | 0.25 | 1.27E-10 | 4.12E-06 |
| Gm48678 | 0.24 | 1.27E-24 | 4.11E-20 |
| Cd33 | 0.24 | 1.82E-17 | 5.86E-13 |
| Stk40 | 0.24 | 3.58E-17 | 1.16E-12 |
| Rab2a | 0.24 | 1.47E-16 | 4.75E-12 |
| Pkn2 | 0.24 | 1.55E-16 | 5.01E-12 |
| Wbp11 | 0.24 | 3.02E-16 | 9.75E-12 |
| Smap2 | 0.24 | 2.56E-15 | 8.26E-11 |
| Lpin2 | 0.24 | 7.84E-15 | 2.53E-10 |
| Thrap3 | 0.24 | 1.96E-14 | 6.33E-10 |
| Prrc2c | 0.24 | 3.00E-14 | 9.68E-10 |
| Eif1 | 0.24 | 4.89E-14 | 1.58E-09 |
| Myh9 | 0.24 | 1.14E-13 | 3.67E-09 |
| Hnrnpll | 0.24 | 1.22E-13 | 3.95E-09 |
| Ythdc1 | 0.24 | 1.33E-13 | 4.31E-09 |
| Ndel1 | 0.24 | 4.51E-13 | 1.45E-08 |
| B2m | 0.24 | 4.60E-13 | 1.49E-08 |
| Fcho2 | 0.24 | 2.24E-12 | 7.22E-08 |
| Klhl2 | 0.24 | 7.51E-12 | 2.43E-07 |
| Eml4 | 0.24 | 8.52E-12 | 2.75E-07 |
| Cblb | 0.24 | 9.10E-12 | 2.94E-07 |
| Zyx | 0.24 | 1.10E-11 | 3.56E-07 |
| Nfil3 | 0.24 | 2.03E-11 | 6.55E-07 |
| Chd7 | 0.24 | 1.19E-10 | 3.85E-06 |
| Rps16 | 0.24 | 1.88E-10 | 6.08E-06 |
| Rpl6 | 0.24 | 2.75E-10 | 8.89E-06 |
| Rpl5 | 0.24 | 3.33E-10 | 1.08E-05 |
| Coro1a | 0.24 | 5.80E-10 | 1.87E-05 |

| | | | |
|---|---|---|---|
| Rps13 | 0.24 | 6.98E-10 | 2.25E-05 |
| Rps11 | 0.24 | 1.50E-09 | 4.84E-05 |
| Ncoa1 | 0.24 | 1.65E-09 | 5.31E-05 |
| Svil | 0.24 | 2.03E-09 | 6.55E-05 |
| Rps7 | 0.24 | 2.51E-09 | 8.09E-05 |
| Fbxl5 | 0.24 | 3.91E-08 | 1.26E-03 |
| Cd177 | 0.23 | 1.22E-20 | 3.94E-16 |
| Wdr45b | 0.23 | 6.67E-17 | 2.15E-12 |
| Cdkn1b | 0.23 | 1.78E-16 | 5.74E-12 |
| Dusp11 | 0.23 | 3.19E-16 | 1.03E-11 |
| Lysmd4 | 0.23 | 7.21E-16 | 2.33E-11 |
| Git2 | 0.23 | 1.02E-15 | 3.29E-11 |
| Eed | 0.23 | 1.04E-15 | 3.34E-11 |
| Cdc40 | 0.23 | 1.22E-15 | 3.94E-11 |
| Laptm5 | 0.23 | 1.29E-15 | 4.17E-11 |
| Pnn | 0.23 | 3.15E-15 | 1.02E-10 |
| Lbr | 0.23 | 4.70E-15 | 1.52E-10 |
| Snrnp70 | 0.23 | 6.72E-15 | 2.17E-10 |
| Acin1 | 0.23 | 9.27E-15 | 2.99E-10 |
| Tmod3 | 0.23 | 9.73E-15 | 3.14E-10 |
| Dhx40 | 0.23 | 2.71E-14 | 8.76E-10 |
| Snx20 | 0.23 | 6.81E-14 | 2.20E-09 |
| Crebbp | 0.23 | 1.06E-13 | 3.42E-09 |
| Cul3 | 0.23 | 1.88E-13 | 6.08E-09 |
| Tut7 | 0.23 | 2.00E-13 | 6.45E-09 |
| Ddit4 | 0.23 | 2.25E-13 | 7.28E-09 |
| Riok3 | 0.23 | 2.95E-13 | 9.52E-09 |
| Tsc22d4 | 0.23 | 3.12E-13 | 1.01E-08 |
| Eif3h | 0.23 | 7.21E-13 | 2.33E-08 |
| Btbd1 | 0.23 | 1.08E-12 | 3.49E-08 |
| Ctss | 0.23 | 1.23E-12 | 3.96E-08 |
| Tpt1 | 0.23 | 1.24E-12 | 4.02E-08 |
| Rb1cc1 | 0.23 | 1.41E-12 | 4.55E-08 |
| Runx2 | 0.23 | 1.67E-12 | 5.40E-08 |
| Il6ra | 0.23 | 2.88E-12 | 9.31E-08 |
| H3f3a | 0.23 | 2.28E-11 | 7.35E-07 |
| Klf3 | 0.23 | 2.71E-11 | 8.76E-07 |
| Rpl7 | 0.23 | 2.83E-11 | 9.14E-07 |
| Pld4 | 0.23 | 1.17E-10 | 3.77E-06 |
| Trem1 | 0.23 | 1.61E-10 | 5.19E-06 |
| Rack1 | 0.23 | 1.85E-10 | 5.99E-06 |
| Tpd52 | 0.23 | 5.04E-10 | 1.63E-05 |
| Lilrb4a | 0.23 | 8.64E-10 | 2.79E-05 |
| Rps3 | 0.23 | 1.67E-09 | 5.38E-05 |
| Spag9 | 0.23 | 4.92E-09 | 1.59E-04 |
| Wnk1 | 0.23 | 5.22E-09 | 1.69E-04 |

| | | | |
|---|---|---|---|
| Sirpb1c | 0.23 | 1.16E-08 | 3.75E-04 |
| Smpdl3a | 0.23 | 2.96E-08 | 9.57E-04 |
| Tgfbi | 0.23 | 1.24E-07 | 4.01E-03 |
| Slc25a28 | 0.22 | 2.08E-18 | 6.73E-14 |
| Tbc1d15 | 0.22 | 1.46E-16 | 4.70E-12 |
| Rassf5 | 0.22 | 1.55E-16 | 5.00E-12 |
| Pofut2 | 0.22 | 3.63E-16 | 1.17E-11 |
| Klhl24 | 0.22 | 7.43E-16 | 2.40E-11 |
| Dhx36 | 0.22 | 7.27E-15 | 2.35E-10 |
| Rbm27 | 0.22 | 1.64E-14 | 5.28E-10 |
| Cd53 | 0.22 | 1.82E-14 | 5.87E-10 |
| Adipor2 | 0.22 | 2.51E-14 | 8.10E-10 |
| Sf3b1 | 0.22 | 3.25E-14 | 1.05E-09 |
| Cggbp1 | 0.22 | 3.71E-14 | 1.20E-09 |
| Pstpip1 | 0.22 | 3.95E-14 | 1.27E-09 |
| Ppp4r3b | 0.22 | 6.15E-14 | 1.99E-09 |
| Elf4 | 0.22 | 6.17E-14 | 1.99E-09 |
| Pafah1b1 | 0.22 | 6.54E-14 | 2.11E-09 |
| Gnas | 0.22 | 8.14E-14 | 2.63E-09 |
| Taf1d | 0.22 | 1.14E-13 | 3.67E-09 |
| Clip1 | 0.22 | 1.14E-13 | 3.69E-09 |
| Mpp6 | 0.22 | 2.33E-13 | 7.52E-09 |
| Prpf4b | 0.22 | 6.69E-13 | 2.16E-08 |
| Tnks | 0.22 | 7.79E-13 | 2.52E-08 |
| Nfam1 | 0.22 | 9.30E-13 | 3.00E-08 |
| Ppp1r12a | 0.22 | 1.21E-12 | 3.92E-08 |
| Arl6ip5 | 0.22 | 1.76E-12 | 5.68E-08 |
| Luc7l2 | 0.22 | 1.89E-12 | 6.12E-08 |
| Map3k20 | 0.22 | 2.87E-12 | 9.26E-08 |
| Ptk2b | 0.22 | 3.61E-12 | 1.17E-07 |
| Map3k8 | 0.22 | 4.08E-12 | 1.32E-07 |
| Gpatch8 | 0.22 | 4.16E-12 | 1.34E-07 |
| Atrx | 0.22 | 7.76E-12 | 2.50E-07 |
| Itgb7 | 0.22 | 8.35E-12 | 2.70E-07 |
| Zfc3h1 | 0.22 | 1.18E-11 | 3.82E-07 |
| Serinc3 | 0.22 | 5.13E-11 | 1.66E-06 |
| Ndufb1-ps | 0.22 | 8.82E-11 | 2.85E-06 |
| Tmem176a | 0.22 | 9.08E-11 | 2.93E-06 |
| Sema4d | 0.22 | 1.08E-10 | 3.50E-06 |
| Stat3 | 0.22 | 1.67E-10 | 5.40E-06 |
| Picalm | 0.22 | 2.88E-10 | 9.29E-06 |
| Gas5 | 0.22 | 4.41E-10 | 1.42E-05 |
| Rpl37a | 0.22 | 5.33E-10 | 1.72E-05 |
| Jarid2 | 0.22 | 2.87E-09 | 9.26E-05 |
| Kdm6b | 0.22 | 5.59E-09 | 1.80E-04 |
| Rpl36 | 0.22 | 8.57E-09 | 2.77E-04 |

| | | | |
|---|---|---|---|
| Ccr1 | 0.22 | 1.76E-08 | 5.67E-04 |
| Fgr | 0.22 | 3.51E-08 | 1.13E-03 |
| Fam107b | 0.22 | 4.61E-08 | 1.49E-03 |
| Itga4 | 0.22 | 5.01E-07 | 1.62E-02 |
| Ms4a6c | 0.22 | 9.55E-07 | 3.08E-02 |
| Pnpla2 | 0.21 | 4.60E-20 | 1.48E-15 |
| Casp6 | 0.21 | 2.12E-17 | 6.83E-13 |
| Dna2 | 0.21 | 6.23E-17 | 2.01E-12 |
| Lrmp | 0.21 | 6.85E-17 | 2.21E-12 |
| Gmip | 0.21 | 1.17E-16 | 3.77E-12 |
| Dhx38 | 0.21 | 1.94E-16 | 6.28E-12 |
| Tor1a | 0.21 | 7.93E-16 | 2.56E-11 |
| Rprd2 | 0.21 | 6.08E-15 | 1.96E-10 |
| Tlk2 | 0.21 | 2.62E-14 | 8.46E-10 |
| Spen | 0.21 | 7.19E-14 | 2.32E-09 |
| Tle3 | 0.21 | 1.89E-13 | 6.11E-09 |
| Rtf1 | 0.21 | 1.98E-13 | 6.40E-09 |
| Smad4 | 0.21 | 4.64E-13 | 1.50E-08 |
| Phf3 | 0.21 | 9.58E-13 | 3.09E-08 |
| Csf2ra | 0.21 | 1.40E-12 | 4.50E-08 |
| Gatad2b | 0.21 | 1.75E-12 | 5.65E-08 |
| Hectd1 | 0.21 | 2.89E-12 | 9.32E-08 |
| Zc3h7a | 0.21 | 5.84E-12 | 1.89E-07 |
| Trem3 | 0.21 | 6.46E-12 | 2.09E-07 |
| Tut4 | 0.21 | 8.45E-12 | 2.73E-07 |
| Trip11 | 0.21 | 1.19E-11 | 3.83E-07 |
| Met | 0.21 | 1.62E-11 | 5.24E-07 |
| 07-Mar | 0.21 | 2.51E-11 | 8.10E-07 |
| Nipbl | 0.21 | 4.53E-11 | 1.46E-06 |
| Clint1 | 0.21 | 4.79E-11 | 1.55E-06 |
| Tln1 | 0.21 | 5.31E-11 | 1.71E-06 |
| Clk1 | 0.21 | 1.10E-10 | 3.55E-06 |
| Noct | 0.21 | 1.67E-10 | 5.38E-06 |
| H2-T23 | 0.21 | 3.55E-10 | 1.15E-05 |
| Cux1 | 0.21 | 4.92E-10 | 1.59E-05 |
| Rbm3 | 0.21 | 7.81E-10 | 2.52E-05 |
| Gmfg | 0.21 | 9.56E-10 | 3.09E-05 |
| Atf6 | 0.21 | 1.34E-09 | 4.32E-05 |
| Slc23a2 | 0.21 | 1.52E-09 | 4.92E-05 |
| Tm6sf1 | 0.21 | 2.41E-09 | 7.77E-05 |
| Map4k4 | 0.21 | 9.12E-09 | 2.94E-04 |
| Rpl35a | 0.21 | 9.27E-09 | 2.99E-04 |
| Tyrobp | 0.21 | 1.31E-08 | 4.24E-04 |
| Rps19 | 0.21 | 1.79E-08 | 5.78E-04 |
| Jmjd1c | 0.21 | 2.08E-08 | 6.73E-04 |
| Rnh1 | 0.21 | 3.78E-08 | 1.22E-03 |

| | | | |
|---|---|---|---|
| Rpl9 | 0.21 | 8.74E-08 | 2.82E-03 |
| Pip4k2a | 0.21 | 1.10E-07 | 3.56E-03 |
| Srgn | 0.21 | 1.44E-07 | 4.65E-03 |
| Lmnb1 | 0.21 | 1.48E-07 | 4.79E-03 |
| Fyb | 0.21 | 9.55E-07 | 3.08E-02 |
| Rps18 | 0.21 | 1.27E-06 | 4.10E-02 |
| Nedd9 | 0.21 | 1.46E-06 | 4.73E-02 |
| Gnpda1 | -0.21 | 9.15E-14 | 2.96E-09 |
| Mrpl14 | -0.21 | 3.01E-13 | 9.72E-09 |
| Colgalt1 | -0.21 | 1.23E-11 | 3.97E-07 |
| Mrps28 | -0.21 | 2.07E-11 | 6.69E-07 |
| Wdr83os | -0.21 | 5.27E-11 | 1.70E-06 |
| Cotl1 | -0.21 | 1.31E-09 | 4.24E-05 |
| Rpn1 | -0.21 | 2.60E-09 | 8.39E-05 |
| Cpd | -0.21 | 3.02E-09 | 9.75E-05 |
| Mcfd2 | -0.21 | 3.09E-09 | 9.98E-05 |
| Anxa4 | -0.21 | 3.30E-09 | 1.07E-04 |
| Adssl1 | -0.21 | 1.02E-08 | 3.30E-04 |
| Nceh1 | -0.21 | 3.13E-08 | 1.01E-03 |
| Exoc4 | -0.21 | 5.96E-08 | 1.92E-03 |
| Pnp | -0.21 | 8.36E-08 | 2.70E-03 |
| Stab1 | -0.21 | 1.98E-07 | 6.40E-03 |
| Rhob | -0.21 | 2.86E-07 | 9.23E-03 |
| Manf | -0.21 | 1.44E-06 | 4.66E-02 |
| Cdk4 | -0.22 | 2.19E-13 | 7.06E-09 |
| Plk2 | -0.22 | 6.52E-13 | 2.10E-08 |
| Dpysl2 | -0.22 | 4.45E-12 | 1.44E-07 |
| Tex14 | -0.22 | 5.16E-12 | 1.67E-07 |
| Tubb2a | -0.22 | 8.33E-12 | 2.69E-07 |
| Ran | -0.22 | 1.00E-09 | 3.23E-05 |
| P4ha1 | -0.22 | 5.96E-09 | 1.92E-04 |
| Tbc1d5 | -0.22 | 3.32E-08 | 1.07E-03 |
| Pgk1 | -0.22 | 3.75E-08 | 1.21E-03 |
| Tcf4 | -0.22 | 5.97E-08 | 1.93E-03 |
| Il1a | -0.22 | 3.48E-07 | 1.12E-02 |
| Selenop | -0.22 | 1.53E-06 | 4.93E-02 |
| Amdhd2 | -0.23 | 3.51E-17 | 1.13E-12 |
| Rab11a | -0.23 | 1.12E-13 | 3.61E-09 |
| mt-Cytb | -0.23 | 1.90E-13 | 6.14E-09 |
| Pepd | -0.23 | 1.22E-12 | 3.93E-08 |
| Airn | -0.23 | 1.35E-12 | 4.34E-08 |
| Lrp12 | -0.23 | 1.64E-12 | 5.31E-08 |
| Selenow | -0.23 | 7.80E-12 | 2.52E-07 |
| Pvt1 | -0.23 | 1.80E-11 | 5.80E-07 |
| Gm21188 | -0.23 | 5.79E-11 | 1.87E-06 |
| Bax | -0.23 | 1.04E-10 | 3.36E-06 |

| | | | |
|---|---|---|---|
| Creld2 | -0.23 | 1.75E-10 | 5.64E-06 |
| Tnfaip8 | -0.23 | 1.47E-09 | 4.75E-05 |
| Cd84 | -0.23 | 2.11E-08 | 6.81E-04 |
| Pycard | -0.23 | 5.59E-08 | 1.80E-03 |
| Tppp3 | -0.23 | 1.75E-07 | 5.66E-03 |
| Ccl12 | -0.23 | 4.77E-07 | 1.54E-02 |
| Rnase4 | -0.24 | 7.95E-15 | 2.57E-10 |
| Fkbp2 | -0.24 | 8.64E-13 | 2.79E-08 |
| Idh1 | -0.24 | 3.87E-12 | 1.25E-07 |
| Nav1 | -0.24 | 5.52E-11 | 1.78E-06 |
| Atp2a2 | -0.24 | 1.47E-10 | 4.73E-06 |
| Tmem37 | -0.24 | 1.55E-10 | 5.01E-06 |
| Atxn1 | -0.24 | 3.42E-09 | 1.11E-04 |
| Mertk | -0.24 | 6.85E-09 | 2.21E-04 |
| Diaph2 | -0.24 | 1.23E-08 | 3.96E-04 |
| Fcgr1 | -0.24 | 3.54E-08 | 1.14E-03 |
| Ednrb | -0.24 | 3.54E-08 | 1.14E-03 |
| Olr1 | -0.24 | 3.53E-07 | 1.14E-02 |
| Sept9 | -0.25 | 1.90E-16 | 6.15E-12 |
| Tbca | -0.25 | 3.65E-14 | 1.18E-09 |
| BC005537 | -0.25 | 8.31E-14 | 2.68E-09 |
| Adgrg6 | -0.25 | 9.94E-13 | 3.21E-08 |
| Specc1 | -0.25 | 5.05E-12 | 1.63E-07 |
| Odc1 | -0.25 | 3.10E-11 | 1.00E-06 |
| Kctd12 | -0.25 | 1.32E-10 | 4.27E-06 |
| Anxa1 | -0.25 | 2.73E-07 | 8.81E-03 |
| Sbf2 | -0.26 | 3.63E-16 | 1.17E-11 |
| Dock1 | -0.26 | 4.36E-16 | 1.41E-11 |
| St6gal1 | -0.26 | 1.49E-14 | 4.82E-10 |
| Dock7 | -0.26 | 5.11E-14 | 1.65E-09 |
| Fam20c | -0.26 | 2.51E-12 | 8.09E-08 |
| Aprt | -0.26 | 3.48E-12 | 1.13E-07 |
| Pdia3 | -0.26 | 2.23E-11 | 7.21E-07 |
| Ldha | -0.26 | 3.15E-10 | 1.02E-05 |
| Prdx1 | -0.26 | 5.59E-10 | 1.81E-05 |
| Pde7b | -0.26 | 2.47E-09 | 7.98E-05 |
| Ptgs2 | -0.26 | 8.58E-07 | 2.77E-02 |
| Rhoc | -0.27 | 2.69E-20 | 8.68E-16 |
| Vps13c | -0.27 | 1.21E-15 | 3.89E-11 |
| Canx | -0.27 | 2.60E-15 | 8.38E-11 |
| Magt1 | -0.27 | 5.12E-15 | 1.65E-10 |
| Uqcc2 | -0.27 | 6.37E-15 | 2.06E-10 |
| Jpt1 | -0.27 | 6.63E-15 | 2.14E-10 |
| Mpp1 | -0.27 | 1.07E-13 | 3.45E-09 |
| Ophn1 | -0.27 | 6.10E-13 | 1.97E-08 |
| Bst2 | -0.27 | 4.13E-12 | 1.33E-07 |

| | | | |
|---|---|---|---|
| Ppp1r14b | -0.27 | 2.03E-11 | 6.55E-07 |
| Igf2r | -0.28 | 2.84E-22 | 9.18E-18 |
| Calm3 | -0.28 | 1.11E-16 | 3.59E-12 |
| Fam162a | -0.28 | 1.02E-14 | 3.31E-10 |
| Ncf1 | -0.28 | 1.36E-12 | 4.38E-08 |
| Man1a | -0.28 | 7.32E-12 | 2.36E-07 |
| Myo1e | -0.28 | 5.77E-10 | 1.86E-05 |
| Tmem106a | -0.29 | 1.69E-19 | 5.45E-15 |
| Timm13 | -0.29 | 2.43E-19 | 7.83E-15 |
| Park7 | -0.29 | 3.09E-17 | 9.97E-13 |
| Sdcbp | -0.29 | 5.95E-16 | 1.92E-11 |
| Ltc4s | -0.29 | 1.16E-15 | 3.75E-11 |
| Nme1 | -0.29 | 2.20E-15 | 7.12E-11 |
| Ranbp1 | -0.29 | 2.12E-14 | 6.84E-10 |
| Dok2 | -0.29 | 2.73E-14 | 8.83E-10 |
| Gapdh | -0.29 | 1.04E-11 | 3.34E-07 |
| Nr4a1 | -0.29 | 3.90E-10 | 1.26E-05 |
| Adgre1 | -0.3 | 4.17E-12 | 1.35E-07 |
| Aldoa | -0.3 | 4.48E-12 | 1.45E-07 |
| Nfkbia | -0.3 | 4.40E-08 | 1.42E-03 |
| Uap1l1 | -0.31 | 6.83E-26 | 2.21E-21 |
| Cyfip1 | -0.31 | 7.77E-16 | 2.51E-11 |
| Hspe1 | -0.31 | 2.91E-13 | 9.41E-09 |
| Rgl1 | -0.31 | 6.11E-13 | 1.97E-08 |
| Sgms1 | -0.31 | 2.31E-11 | 7.45E-07 |
| Hivep2 | -0.31 | 3.15E-11 | 1.02E-06 |
| Hspa5 | -0.31 | 4.26E-11 | 1.37E-06 |
| Arhgap24 | -0.31 | 4.81E-10 | 1.55E-05 |
| Basp1 | -0.31 | 2.12E-07 | 6.84E-03 |
| Cacna1d | -0.31 | 3.26E-07 | 1.05E-02 |
| Bcar3 | -0.32 | 7.27E-27 | 2.35E-22 |
| Tpp1 | -0.32 | 1.15E-25 | 3.71E-21 |
| F7 | -0.32 | 1.11E-24 | 3.59E-20 |
| Pfn1 | -0.32 | 2.07E-18 | 6.68E-14 |
| Tubb5 | -0.32 | 7.34E-16 | 2.37E-11 |
| Tuba1b | -0.32 | 1.74E-15 | 5.62E-11 |
| Capg | -0.32 | 2.14E-14 | 6.91E-10 |
| Phlda1 | -0.32 | 4.68E-11 | 1.51E-06 |
| Dock2 | -0.33 | 1.64E-15 | 5.29E-11 |
| Lgmn | -0.33 | 2.12E-11 | 6.86E-07 |
| Uba52 | -0.34 | 4.46E-19 | 1.44E-14 |
| Psd3 | -0.34 | 4.11E-18 | 1.33E-13 |
| Slc9a9 | -0.34 | 1.12E-12 | 3.63E-08 |
| Pde4d | -0.34 | 1.13E-11 | 3.65E-07 |
| Fabp5 | -0.34 | 8.60E-07 | 2.78E-02 |
| Vat1 | -0.35 | 1.53E-22 | 4.93E-18 |

| | | | |
|---|---|---|---|
| Itsn1 | -0.35 | 6.49E-22 | 2.09E-17 |
| Grn | -0.35 | 5.16E-20 | 1.66E-15 |
| Dhrs3 | -0.35 | 1.11E-19 | 3.60E-15 |
| Arhgap10 | -0.35 | 1.50E-19 | 4.83E-15 |
| Frmd4b | -0.35 | 1.70E-19 | 5.48E-15 |
| Ctsc | -0.35 | 5.76E-15 | 1.86E-10 |
| Gm17268 | -0.35 | 1.85E-11 | 5.96E-07 |
| Klf2 | -0.35 | 6.92E-10 | 2.23E-05 |
| Tnf | -0.36 | 1.89E-19 | 6.09E-15 |
| Nrp1 | -0.36 | 4.32E-19 | 1.39E-14 |
| Mitf | -0.36 | 2.85E-14 | 9.21E-10 |
| Sdc3 | -0.37 | 4.41E-25 | 1.42E-20 |
| Anxa5 | -0.37 | 8.93E-23 | 2.88E-18 |
| Mthfs | -0.37 | 1.02E-21 | 3.31E-17 |
| Hdac9 | -0.37 | 4.58E-17 | 1.48E-12 |
| C1qc | -0.37 | 3.75E-10 | 1.21E-05 |
| 4930430E12F | -0.38 | 1.42E-26 | 4.60E-22 |
| Slc7a8 | -0.38 | 1.08E-20 | 3.48E-16 |
| Trem2 | -0.38 | 2.78E-20 | 8.98E-16 |
| Sdf2l1 | -0.38 | 1.72E-18 | 5.56E-14 |
| Myof | -0.39 | 3.67E-21 | 1.18E-16 |
| Cd93 | -0.39 | 2.21E-20 | 7.15E-16 |
| Dennd1a | -0.39 | 8.81E-19 | 2.84E-14 |
| Pdia6 | -0.39 | 4.29E-18 | 1.38E-13 |
| Calr | -0.39 | 6.62E-18 | 2.14E-13 |
| Abca1 | -0.39 | 1.34E-13 | 4.33E-09 |
| Gm10076 | -0.4 | 5.58E-31 | 1.80E-26 |
| Fmn1 | -0.4 | 6.63E-25 | 2.14E-20 |
| Cxcl1 | -0.4 | 1.33E-19 | 4.30E-15 |
| Pitpnc1 | -0.4 | 1.03E-15 | 3.32E-11 |
| Lrmda | -0.4 | 9.33E-14 | 3.01E-09 |
| Sash1 | -0.41 | 4.37E-20 | 1.41E-15 |
| C5ar1 | -0.41 | 4.14E-17 | 1.34E-12 |
| Ifrd1 | -0.41 | 1.36E-14 | 4.40E-10 |
| Emp1 | -0.41 | 3.18E-13 | 1.03E-08 |
| Hsp90b1 | -0.42 | 1.67E-22 | 5.39E-18 |
| Cfp | -0.42 | 4.20E-21 | 1.35E-16 |
| Ctsl | -0.42 | 3.35E-12 | 1.08E-07 |
| C1qa | -0.42 | 3.11E-11 | 1.00E-06 |
| Anxa3 | -0.43 | 2.00E-23 | 6.47E-19 |
| Ccrl2 | -0.43 | 4.44E-18 | 1.43E-13 |
| Dynll1 | -0.44 | 4.91E-24 | 1.58E-19 |
| Tpi1 | -0.44 | 4.62E-22 | 1.49E-17 |
| Lpp | -0.44 | 1.96E-21 | 6.34E-17 |
| Gadd45b | -0.45 | 4.69E-27 | 1.52E-22 |
| Tagln2 | -0.45 | 7.47E-20 | 2.41E-15 |

| | | | |
|---|---|---|---|
| Ctsb | -0.46 | 4.90E-25 | 1.58E-20 |
| Ecm1 | -0.46 | 9.62E-24 | 3.10E-19 |
| Dab2 | -0.46 | 2.56E-15 | 8.26E-11 |
| Dip2c | -0.48 | 2.14E-23 | 6.90E-19 |
| Cstb | -0.48 | 7.53E-20 | 2.43E-15 |
| C3ar1 | -0.49 | 2.55E-26 | 8.23E-22 |
| Egr1 | -0.49 | 5.79E-16 | 1.87E-11 |
| C1qb | -0.49 | 1.39E-11 | 4.49E-07 |
| Lmna | -0.5 | 6.32E-21 | 2.04E-16 |
| Pdia4 | -0.51 | 1.60E-31 | 5.18E-27 |
| Rasgef1b | -0.51 | 1.62E-30 | 5.23E-26 |
| Maf | -0.51 | 1.96E-19 | 6.34E-15 |
| Pdpn | -0.52 | 8.77E-23 | 2.83E-18 |
| Tbxas1 | -0.54 | 3.67E-24 | 1.19E-19 |
| Fn1 | -0.54 | 2.75E-18 | 8.87E-14 |
| Bcl2a1b | -0.56 | 9.86E-21 | 3.18E-16 |
| Fosb | -0.59 | 1.15E-29 | 3.72E-25 |
| Ccl9 | -0.6 | 3.04E-23 | 9.81E-19 |
| Mrc1 | -0.61 | 2.05E-21 | 6.62E-17 |
| Ppbp | -0.61 | 1.56E-15 | 5.02E-11 |
| Cxcl2 | -0.62 | 1.72E-10 | 5.54E-06 |
| Tnfaip3 | -0.63 | 6.65E-36 | 2.15E-31 |
| Mif | -0.65 | 5.12E-39 | 1.65E-34 |
| Lgals1 | -0.67 | 1.30E-32 | 4.19E-28 |
| Ctsd | -0.68 | 4.55E-36 | 1.47E-31 |
| Nfkbiz | -0.71 | 7.36E-40 | 2.38E-35 |
| Ccl6 | -0.8 | 5.02E-29 | 1.62E-24 |
| Ccl24 | -0.86 | 1.70E-33 | 5.50E-29 |
| Ccl2 | -0.92 | 1.21E-26 | 3.89E-22 |
| Ccl7 | -1.11 | 2.07E-41 | 6.68E-37 |
| Arg1 | -1.21 | 7.94E-36 | 2.56E-31 |
| Spp1 | -1.23 | 1.43E-42 | 4.61E-38 |
| Pf4 | -1.51 | 7.41E-46 | 2.39E-41 |
